# Identifying and quantifying conflicts between humans and terrestrial mammals in Great Britain

**DOI:** 10.1101/2025.07.02.662702

**Authors:** Kate L. Palphramand, Daniel A. Warren, Graham C. Smith, Dave Cowan

## Abstract

1. A literature review was conducted to identify human-wildlife conflicts associated with British terrestrial mammal species. Conflicts were classified as economic, health, environmental or social, divided into 32 subcategories, and ranked using a Generic Impact Scoring System (GISS).
2. We identified 48 species associated with 200 conflicts. Sika deer were involved in the most conflicts. The highest ranked conflicts were measurable on an economic scale (involving rabbits, badgers, brown rats, grey squirrels), with the total estimated cost of all economic conflicts exceeding £0.5 billion.
3. The most common conflicts were reservoirs of disease and zoonotic disease, with non-native species scoring statistically higher than native species for the latter. Generally, we scored these conflicts low, deemed localised and mild, but highlighted the importance of surveillance to monitor and control disease spread.
4. Most species caused minimal conflict, likely due to their limited distribution. We identified potential impact increased as a function of biomass and population size, therefore a GISS may help identify species capable of expanding beyond their current range, such as recently reintroduced beavers.
5. A GISS is useful for identifying conflict species but there is also the need to understand the value of British wildlife. We identified costs-to-benefits trade-offs for several high impact species, including rabbits, deer, badgers and foxes, which underlie human-conflict and/or coexistence and are critical for informed decision-making. As one in four British mammal species face local extinction, the emphasis should encourage focus away from conflict resolution to acceptance and/or tolerance where wildlife and people coexist.

## Introduction

Interactions between humans and wildlife form an integral part of many ecosystems worldwide, and are often varied and complex (Hill 2021). Whilst the presence of wildlife may have beneficial impacts on human interests, there is a continued focus on conflict narratives that risks conceptualising how human-wildlife interactions are viewed (Hill 2021). Such negative terminology implies deliberate action by wildlife species and ignores the conflict between interested groups of people about how a situation should be resolved (Peterson et al. 2010; IUCN 2022). Shifting the focus of human-wildlife relationships towards one based on coexistence acknowledges the idea that interactions are not solely negative and, rather, conflict should be considered as only one aspect of human-wildlife coexistence (Hill 2021).

Human-wildlife conflict is defined as the struggles that emerge from the presence or behaviour of wildlife that pose an actual or perceived, direct, and recurring threat to human interests or needs, leading to disagreements between groups of people and negative impacts on people and/or wildlife (IUCN SSC HWCTF 2020). There are many factors that can cause conflict wherever wildlife and human populations overlap, including human population growth, agricultural expansion, infrastructure development, climate change and other drivers of habitat loss (Zabel et al. 2019; IUCN 2022). Wildlife damage to crops is often cited as a major global conflict (Woodroffe, Thirgood, and Rabinowitz 2005; Pérez and Pacheco 2006; Fang et al. 2021), which will likely worsen as natural habitat loss due to the expansion of human activities precipitates an increase in human-wildlife encounters (Seoraj-Pillai and Pillay 2017; König et al. 2020). Population growth and habitat encroachment may also increase the risk of disease transmission, which is of particular concern where wildlife pose zoonotic risks. Human activities, such as modification of natural habitat and expanding urbanisation, not only risk increasing interactions with wildlife, but they also risk extirpating native species from an area whilst promoting the establishment of non-native species that may be better adapted to urban and suburban conditions (McKinney 2006; Borden and Flory 2021).

Invasive non-native species are a major threat to biodiversity worldwide, severely impacting areas of high endemism (Jeschke et al. 2014; Doherty et al. 2016; Dueñas et al. 2021) whilst also posing a serious threat to public health and economic activities. As of 2020, there were 3,248 non-native species in Great Britain; 300 of these are designated as having a negative ecological or human impact (NNSS 2020), costing the UK economy almost £4 billion annually (Eschen et al. 2023). Invasive mammalian species are considered to be one of the most damaging groups worldwide, particularly cats, rats, foxes and stoats (Medina et al. 2011; Szabo et al. 2012; Bellard, Genovesi, and Jeschke 2016; Doherty et al. 2016; Nentwig et al. 2018). Terrestrial mammal species include some of Britain’s most iconic and well-loved species and yet many of them are among the most depleted, with one in four species facing local extinction; the reasons for these declines vary between species, but include persecution, habitat loss and the introduction of non-native species (Mammal Society 2024b). Categorising British mammal species as invasive can be an emotive subject, with people’s perceptions largely dependent on the species threatened with control (Baker et al. 2020), resulting in debates on achievability, efficiency, social fairness and ethical implications (Crowley, Hinchliffe, and McDonald 2017). For example, the removal of hedgehogs from South Uist, where they were introduced to control garden pests but consequently contributed significantly to the decline of wading shorebirds (Jackson and Green 2000) resulted in widespread criticism and opposition, given that hedgehogs are a popular icon of British wildlife and are thought to be declining nationally (Crowley, Hinchliffe, and McDonald 2017). Invasive species management frequently leads to disputes and conflicts among people, and is often driven by the focus of interest, e.g. biodiversity conservation, economic, ecosystem services, animal and human health (Crowley, Hinchliffe, and McDonald 2017).

Much of the available literature on wildlife conflicts in the UK, both in terms of economic and biodiversity impacts, focuses on invasive non-native species (e.g. Manchester and Bullock 2000; Baker 2010; Williams et al. 2010; Eschen et al. 2023). However, there is evidence to suggest that the most invasive non-native species are not necessarily the most damaging (Ricciardi and Cohen 2007; Ricciardi et al. 2013; Cassini 2022), therefore there is the need to consider also the impacts of native species, some of which can cause considerable human-wildlife conflict and associated controversies. An example of such complexity is the badger as a known reservoir host of *Mycobacterium bovis*, the causative agent of bovine tuberculosis (bTB), which poses a risk to livestock health and is a serious economic problem for the UK cattle industry (Swift et al. 2021). Despite costing the taxpayer over £100 million annually (Defra 2024), public opinion on approaches to managing this conflict, especially with respect to culling, is deeply divided and polarised (Enticott 2015; Naylor et al. 2017; Cassidy 2019). Issues such as this highlight the challenge of conflict resolution for iconic British species, which can be both controversial and associated with complex legislative considerations. It is necessary, however, to consider all species as part of any wildlife management or conservation and development efforts (Gross et al. 2024).

The emphasis of legislation associated with wildlife in the UK has moved from protecting human economic interests (e.g. Prevention of Damage by Pests Act 1949 and Pests Act 1954), to promoting animal welfare (e.g. Wildlife and Countryside Act 1981 and Animal Welfare Act 2006) and protecting biodiversity (e.g. Habitats Directive 1992, Environment Act 2021). There is a general consensus that the negative effects created by some wildlife species, particularly invasive non-native species, have to be reduced (Nentwig, Kühnel, and Bacher 2010). Tackling invasive non-native species is a key part of achieving the UK government’s aims under its 25-year Environment plan (HM Government 2018) as well as the UK’s obligation as a signatory of the Convention on Biological Diversity (https://www.cbd.int/cop). The 2022 Global Biodiversity Framework (GBF) identified managing human-wildlife interactions to minimise human-wildlife conflict for coexistence as one of the targets for reducing threats to biodiversity, which in turn would feed into other elements of the framework, such as enhancing sustainability in agriculture, fisheries and forestry (The Wildlife Trust 2024). Where conflicts arise between wildlife and human interests, there is often the need to intervene to reduce or mitigate the impacts and therefore identifying what conflicts are occurring and what their impacts are is essential for informing decision-making and resource allocation (Eschen et al. 2023). However, identifying which species are causing conflict and subsequently which should be targeted first can be challenging, as it is difficult to compare the damage caused by different species, both within the same and different taxonomic groups (Nentwig, Kühnel, and Bacher 2010).

Whilst policy frameworks such as the GBF are essential for providing goals and targets to reverse biodiversity loss, they do not necessarily incorporate a process by which these targets, such as managing human-wildlife conflict, can be identified in the first place, nor which should be prioritised. Assessing conflicts between species is further complicated by the fact that impacts are context dependent (Kumschick et al. 2015), which requires that the measure of impact is standardised to quantify and compare differences between species, taxonomic groups and localities (Nentwig, Bacher, and Pyšek 2016). When addressing the biodiversity impacts of invasive non-native species, Nentwig, Kühnel, and Bacher (2010) suggested prioritising actions against the alien (non-native) invasive species with a high impact potential of becoming established or spreading, using a method to quantify impacts in a way that could be compared among alien species within a taxonomic group. They devised a generic impact scoring system (GISS) to compare relevant impact categories that had been reported in the scientific literature, applying a scoring system to quantify impacts within the categories and rank species according to their impact scores. The GISS has since been modified and applied to a variety of classifications including alien and invasive fish (Van der Veer and Nentwig 2014), alien terrestrial arthropods (Vaes-Petignat and Nentwig 2014), alien invasive aquatic invertebrates (Laverty et al. 2015), plants (Novoa et al. 2016; Sohrabi et al. 2021) and alien birds (Evans et al. 2020).

The main aim of our study was to identify and categorise human-wildlife conflicts associated with terrestrial mammal species in Great Britain in order to summarise and rank their magnitude and, where possible, their economic costs. The research augmented several reviews of human-wildlife conflicts produced over a decade ago (APHA, unpublished data) in response to a now withdrawn Defra policy framework on Wildlife Management in England (Defra 2010). The principal questions we aimed to address included: (1) are human-wildlife conflicts featuring terrestrial mammal species in Great Britain represented in the literature; (2) what conflicts have been reported; (3) can the evidence be used to assess the level of risk to human interests or the welfare or conservation status of an affected species? To determine the magnitude of conflicts, a modified GISS was applied to all conflicts so that species could be ranked according to their level of impact. Identifying species with the highest impact scores can assist policy development by enabling potential prioritisation of the most damaging species. In addition, the compilation of a comprehensive audit of conflicts associated with British mammal species highlights where further areas of research might be required and identifies the early emergence of potential issues.

## Materials and methods

### Design

This review was based on the PRISMA extension for scoping reviews (PRISMA-ScR) checklist (Tricco et al. 2018), to explore the extent and nature of conflict between humans and mammal species in Great Britain. The concept explored compiling and analysing conflicts with negative ecological or human impacts to identify mammal species for informed decision-making, and presenting the data in a way that could be reviewed and updated in future research. The context included terrestrial mammal species only within Great Britain as they involve some of the most controversial and sensitive issues as well as complex legislative considerations. The types of sources eligible for the review included peer-reviewed articles, conference proceedings, theses, published books/chapters, government and non-governmental reports and relevant websites. No date range was applied to the searches.

### Literature searches

Searches were undertaken using Web of Science and Scopus covering the review for the Defra policy framework (25^th^ November 2009 to 8^th^ April 2010 and 17^th^ December 2013 to 23^rd^ February 2014) and the augmented review (4^th^ October 2023 to 24^th^ Jan 2025). Separate searches were carried out for 64 terrestrial mammal species found in Great Britain; this included 58 resident species (see Mathews et al. (2018)), plus more recently established species (beavers, Greater white-toothed shrews) and feral species (e.g. feral cats, feral goats, feral ferrets, Exmoor/Dartmoor ponies). The following search terms were used: [{common name} AND {Latin name}] AND [{England} OR {Scotland} OR {Wales} OR {Great Britain} OR] AND [{conflict} OR {pest} OR {damage} OR {impact} OR {management} OR {disease} OR {zoono*} OR {animal health} OR {health} OR {conservation} OR {predat*}]. Due to the large number of outputs generated, reports were exported to Microsoft Excel where an initial screening of the title for any likely relevance to the species concerned and potential conflicts was carried out by two reviewers (KP and DC). The minimum requirements for a literature resource to be included in the analysis were the presence of an accessible abstract and that the conflict occurred in Great Britain. Only articles written in English were included.

### Data extraction and analysis

Extraction spreadsheets were developed in Microsoft Excel to assist with data collection. Data were extracted by two reviewers (KP and DC) by examining the abstracts in more detail and the full article. Data regarding study location, source type, date, conflict, species, duplication (i.e. the article was applicable to multiple conflicts and/or more than one species) were recorded. Articles that did not fit the criteria (i.e. the conflict was not occurring in Great Britain) but which were considered useful for compiling the review were retained but only conflicts that were reported as occurring in Great Britain were used in the analyses. Additional literature that was not captured in our initial searches was sourced through screening the reference lists of eligible articles and relevant websites. A full list of screened references is included in Supplementary Table S1.

The total number of species and conflicts identified using the extraction spreadsheets were used to compile a detailed report of the human-wildlife conflicts in Great Britain (Supplementary Material S2).

### Categorising and scoring conflicts

Conflicts were categorised as either economic, health, environmental or social impact. Within these four main categories, conflicts were further assigned to 32 subcategories (economic=15, health=6, environmental=7, social=4: Table 1). In the absence of objective scientific or social criteria to consider one impact category more important than the other, subcategories were not weighted and therefore considered of equal importance, as per Nentwig, Kühnel, and Bacher (2010). Some species were involved in multiple conflicts within multiple categories, which were captured as distinct entities. An impact scale scoring system modified from Nentwig, Kühnel, and Bacher (2010) was applied to estimate the magnitude of each conflict at the national level for Great Britain, rated independently by three reviewers (KP, DC and GS). The level of impact was quantified on a scale of 1 (minimal impact known) to 5 (the highest possible impact: Table 2), with a value of confidence (low=1, medium=2 or high=3: Table 3) also assigned to each conflict (taken from Hawkins et al. 2015). Species for which no data were available, no impacts were known or impacts were not detectable, were excluded from the analysis. The median of the three independent assignments of impact scale and confidence rating are reported for each conflict.

**Table 1.** Conflict subcategories identified within the four main conflict areas.

| Conflict category | Subcategory |
| --- | --- |
| Economic | Damage to forestry<br>Damage to agricultural interests<br>Damage to vehicles (RTAs)<br>Competition with livestock<br>Predation of game species<br>Predation / impact on commercial fisheries<br>Predation of poultry<br>Damage to property<br>Damage to amenities and domestic grassland<br>Conflicts with construction, infrastructure and development<br>Predation of livestock<br>Conflicts with transport<br>Sustaining predators harmful to game<br>Damage to property as a result of poaching<br>Conflicts with utilities |
| Health | Health risks from RTAs<br>Reservoir of disease<br>Bovine tuberculosis (bTB) transmission<br>Reservoir of zoonotic disease<br>Threat to livestock where ground is undermined<br>Threat of branches falling |
| Environment | Damage to conservation habitats<br>Threat to biodiversity<br>Threat to genetic integrity<br>Competition with or predation of wildlife (including protected or conservation species)<br>Predation of important ground-nesting birds and island ecosystem damage<br>Sustaining predators harmful to important conservation species<br>Damage to island ecosystems |
| Social | Public nuisance<br>Domestic nuisance<br>Predation of domestic pets<br>Domestic nuisance and hygiene |

**Table 2.** Criteria used to describe the scale of impacts in each of the four categories of conflict.

| Scale of impact | A. Economic impact | B. Health impact | C. Environmental impact | D. Social impact |
| --- | --- | --- | --- | --- |
| 1 | <£10K / yr | Local mild, short-term, reversible effects to individuals | Local, short-term, population loss, no significant ecosystem effect | Minimal social disruption |
| 2 | £10K – 100K / yr | Mild short-term, reversible effects to identifiable groups, localised | Some ecosystem impact, reversible changes, localised | Significant concern expressed at local level |
| 3 | £100K – 1M / yr | Minor irreversible effects and/or larger numbers covered by reversible effects, localised | Measurable long-term damage to populations and ecosystem, but little spread, no extinction | Temporary changes to normal activities at local level |
| 4 | £1M – 10M / yr | Significant irreversible effects locally or reversible effects over a large area | Long-term irreversible ecosystem change, spreading beyond local area | Some permanent change of activity locally, concern over a wider area |
| 5 | >£10M / yr | Widespread, severe, long-term, irreversible health effects | Widespread, long-term population loss or extinction, affecting several species with serious ecosystem effects | Long-term social change, significant loss of employment, migration from area |

**Table 3.** Criteria used for the confidence ratings of the impact scale assignments.

| Confidence level | Example evidence |
| --- | --- |
| **High (3)** (approx. 90% chance of impact scale assignment being correct) | There is direct relevant observational evidence to support the assessment; **and**<br>Impacts are recorded at the typical spatial scale over which the conflict can be characterised; **and**<br>There are reliable / good quality data sources on the impacts of the conflict; **and**<br>The interpretation of the data / information is straightforward; **and**<br>Data / information is not controversial or contradictory |
| **Medium (2)** (approx. 65-75% chance of impact scale assignment being correct) | There is some direct observational evidence to support the assessment but some information is inferred; **and/or**<br>Impacts are recorded at a spatial scale, which may not be relevant to the scale over which conflict can be characterized, but extrapolation or downscaling of the data to relevant scales is considered reliable, or to embrace little uncertainty; **and/or**<br>The interpretation of the data is to some extent ambiguous or contradictory |
| **Low (1)** (approx. 35% chance of impact scale assignment being correct) | There is no direct observational evidence to support the assessment, e.g. only inferred data have been used as supporting evidence; **and/or**<br>Impacts are recorded at a spatial scale, which is unlikely to be relevant to the scale over which conflict can be characterized; extrapolation or downscaling of the data to relevant scales is considered unreliable potentially leading to significant uncertainties; **and/or**<br>Evidence is poor and difficult to interpret, e.g. because it is strongly ambiguous; **and/or**<br>The information sources are considered to be of low quality or contain information that is unreliable |

Various species characteristics were also included in the analyses to determine whether they were associated with a conflict impact, modified from Nentwig, Kühnel, and Bacher (2010). Impact scores were considered as the highest impact for a particular species, irrespective of their distribution and the area they occupy within Great Britain. Overall impacts were compared between species by summing the median scores of all conflicts and within each category (“maximum impact”). To consider the distribution of a species, maximum impact values were multiplied by the proportion of hectads (10 kilometre square resolution) each species occupies in Great Britain (derived from Crawley et al. (2020). The maximum impact scores, corrected for the area occupied can be considered as the “potential impact” a species may have in Great Britain. Estimated population sizes were derived mostly from Mathews et al. (2018), typical body weights were taken mostly from Harris and Yalden (2008). Where information was missing, relevant online sources were used (see Supplementary Table S3). The overall biomass for each species was calculated by multiplying population estimates and body mass. Population estimates and/or distribution data could not be reliably established for three species (feral cat, feral ferret, greater white-toothed shrew), therefore they were excluded from some of the statistical analyses.

### Statistical analysis

All statistical analyses were performed using the R statistical programme version 4.2.2 (R Core Team 2022) and the RStudio Integrated Development Environment (IDE: version 2022.12.0, Posit Software, PBC). Initial analyses (to assess whether impact/confidence scores assigned by the three reviewers differed overall and in relation to the conflict species assessed) were performed using an ordinal logistic regression model fitted using the *ordinal::clm* function version 2023.12-4. (Christensen 2023). Species were aggregated based on status (native/non-native/feral) with reviewer ID included as an interaction term. Additional analyses were performed to assess whether impact scores assigned by the reviewers differed across impact categories (i.e. economic, environmental, health and social), across subcategories and between native/non-native species, including interaction terms. (N.B. feral species were removed from further analyses due to their low number of conflicts). Data were expanded to account for the disparity in the number of subcategories described for each species from the literature, so that for each species, a total of 32 subcategories were present, with an impact and confidence score of zero (0), representing no available data, assigned to absent subcategories.

Differences in maximum and potential impact scores were analysed using generalised linear models (GLM), using the *glmmTMB* package version 1.1.10 (Brooks et al. 2017). Diagnostic testing of these analyses was performed using the *DHARMa* package version 0.4.7 (Hartig 2024). Prior to analysis, data were modified to account for species whereby a potential impact score could not be reliably established, with a score of zero (0) assigned to indicate insufficient evidence of impact. Feral ferret was subsequently excluded from the analyses, given a lack of data regarding population size and geographic distribution. Analyses assessed whether estimates of impact (maximum or potential) differed in relation to native/non-native status when modelled independently. Several covariate predictors were also specified independently in each model, including population size, body mass (kg) and biomass (kg) (see Supplementary Table S4). Models consisted of GLMs, fitted with a gaussian error distribution and “identity” link function, selected given that model residuals conformed to key assumptions (i.e. linearity, normality, homogeneity of variance), when total impact estimates were logarithmically transformed [log(*n*+1)]. For each model, a two-way interaction term was specified between native/non-native status and each covariate predictor (population size, body mass, biomass). Due to issues with model performance, population size, body mass and biomass were also log-transformed [log(*n*+1)].

Additional analyses examined how estimates of impact (maximum or potential) might differ across the main impact categories (economic, health, environment, social); these were performed using a GLM fitted with a Tweedie distribution, which is appropriate when analysing a continuous response variable consisting of a combination of zeroes and positive values(Jorgenson 1987; Ma, Yan, and Hasan 2018). These models compared impact across categories in relation to native/non-native status as well as population size, body mass and biomass. Interaction terms involving impact category, species status and each covariate predictor were included with respect to native/non-native species only.

## Results

### General literature searches

A search of Web of Science and Scopus yielded 12,938 results for 64 terrestrial mammal species in Great Britain. For 16 of the species, no articles focused on conflict were returned (1,017 results). A further 11,691 articles were removed prior to full article screening as they were deemed irrelevant. A total of 230 database articles and 185 articles identified from citations within eligible articles were retrieved and underwent full screening. A further 10 internal reports and 41 relevant websites were also identified (see Supplementary Table S1). Of these, 295 met the eligibility criteria (mammal species conflicts specific to Great Britain); 79 were relevant to multiple species and/or conflicts resulting in a total of 564 articles used in the analyses. A PRISMA flow chart detailing the selection process is shown in Figure 1.

**Figure. 1.**
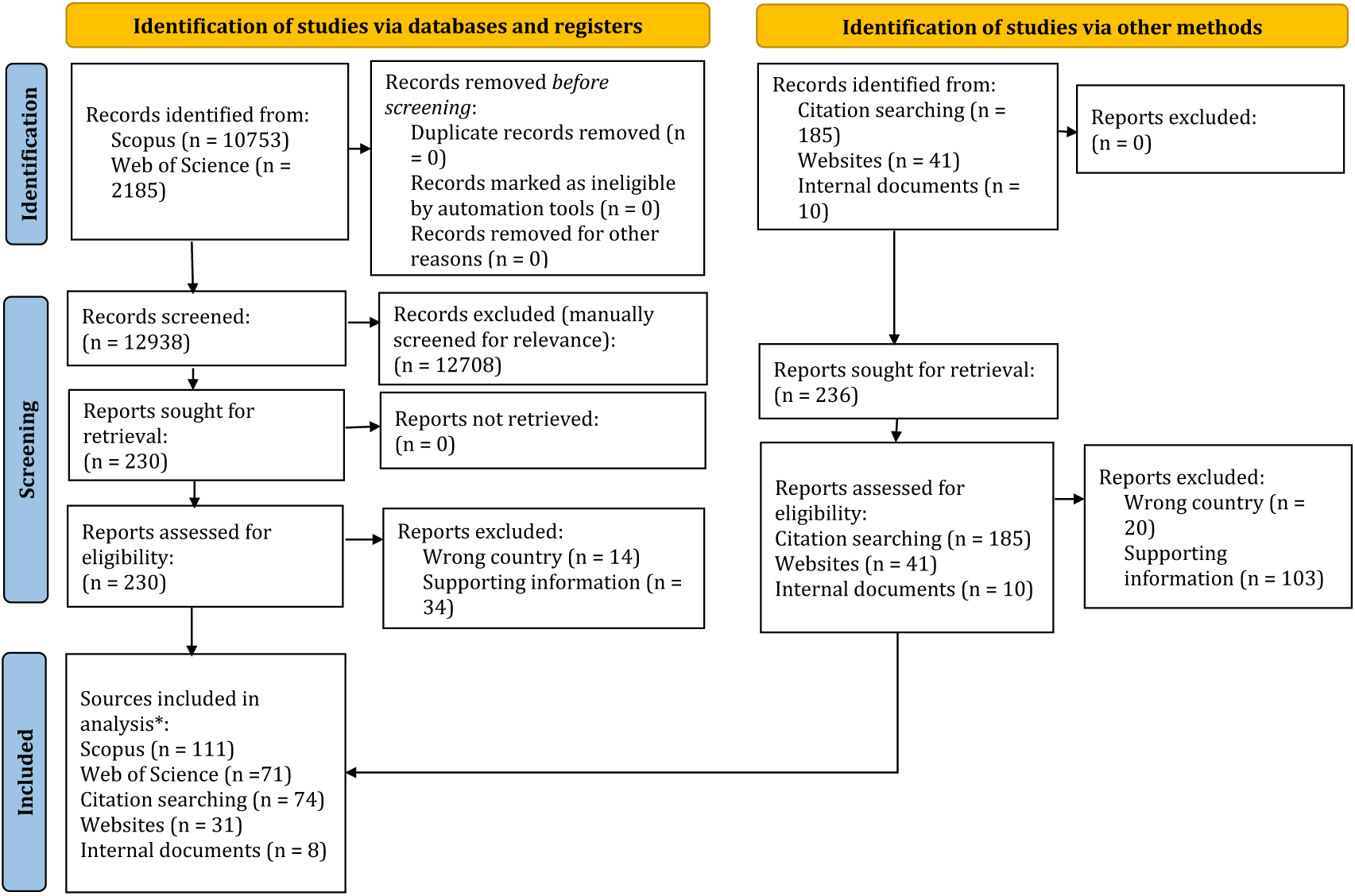
PRISMA flow chart of the scoping review showing the number of records selected after each step. * Seventy-nine articles were relevant to more than one species and/or conflict.

The 48 mammal species identified as causing conflict belonged to seven taxonomic groups (Rodentia: n=15; Carnivora: n=11; Artiodactyla: n=8; Chiroptera: n=6; Eulipotyphla: n=4; Lagomorpha: n=3; Perissodactyla: n=1). The first available record was from 1947 and the latest was from 2024. The majority of articles (n=367, 91%) were published within the last three decades (2014-2024: n=176, 44%; 2004-2013: n=110, 27%; 1994-2003: n=81, 20%).

A detailed description of all of the conflicts is in Supplementary Material S2. A summary of characteristics (native/non-native/feral status, population size, body mass, biomass) and conflicts for each species is in Supplementary Table S3. The periods during which conflicts were reported in the scientific literature (i.e. identified during the original reviews only (“pre-2014”), during the original and current reviews (“pre-2014 and 2014 to present”) and during the current review only (“2014 to present”) are also shown. The median impact scores and confidence ratings are shown after each conflict in parentheses.

### Overview of conflicts

#### Conflicts and status

Twenty-nine of the 48 mammal species (61%) are native to the British Isles, 17 (35%) are non-native (which includes 10 that have been introduced in recent times and seven that are considered naturalised) and two (4%) are feral (species that are considered wild but are either managed or may have human dependence). Of the 48 species, 40 (83%) caused at least one economic conflict, 36 (75%) caused at least one health conflict, 23 (48%) caused at least one environmental conflict and eight (17%) caused at least one social conflict. When compared by status (Figure 2), 86% (25/29) of native species caused economic conflict, 66% (19/29) caused health conflict and 35% (10/29) caused environmental conflict. In comparison, 82% (14/17) of non-native species caused economic conflict, 88% (15/17) caused health conflict and 71% (12/17) caused environmental conflict. Social conflicts accounted for approximately 17% for both native (5/29) and non-native (3/17) species. The proportion of feral species causing conflict was either 50% (economic and environment) or 100% (health), although this was attributable to only two species (feral cat, Dartmoor/Exmoor pony).

**Figure 2.**
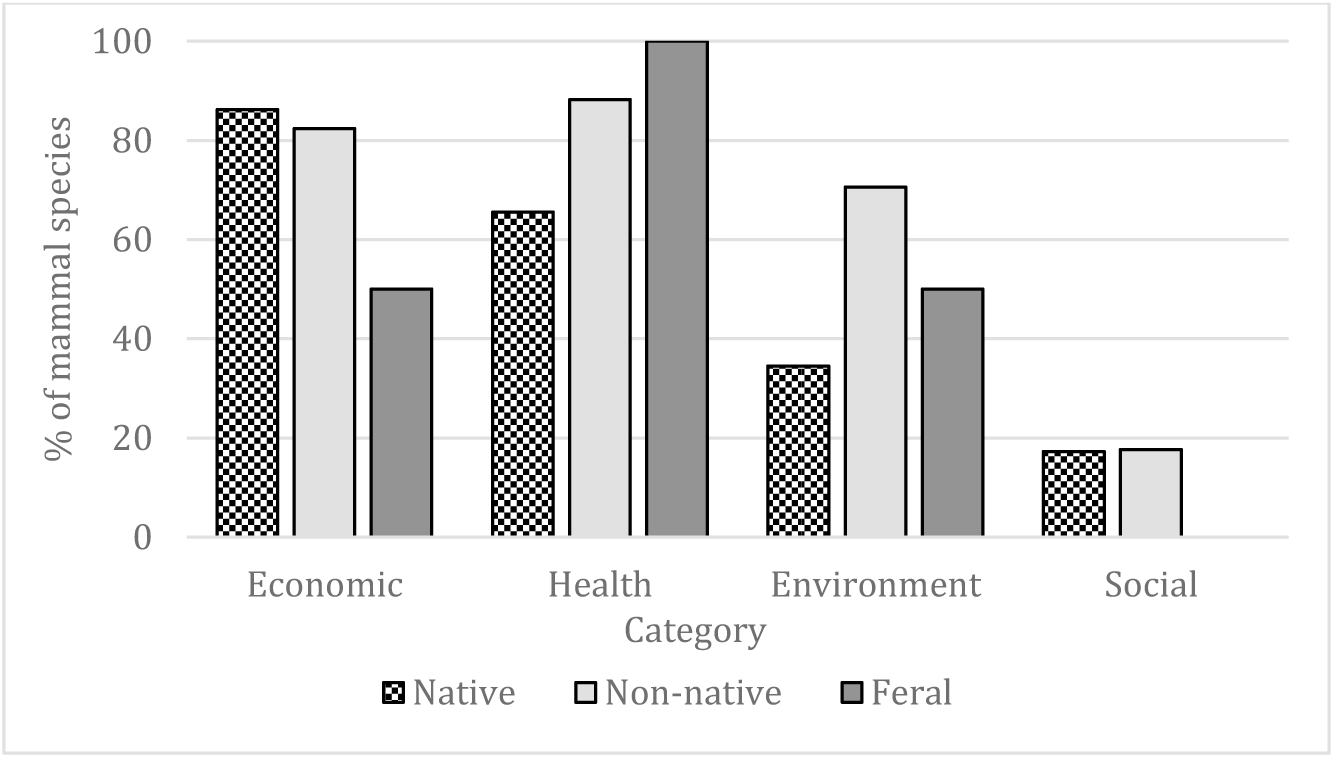
Percentage of species within each status causing conflict within the four main categories.

#### Frequency of conflicts

In total, 200 separate conflicts were identified, with 103 attributable to native species (52%), 92 to non-native species (46%) and five to feral species (3%). Ninety-one of the 200 conflicts (45%) were economic impacts (Supplementary Table 5a), 71 (35%) were health impacts (Supplementary Table 5b), 29 (15%) were environmental impacts (Supplementary Table 5c) and nine (5%) were social impacts (Supplementary Table 5d). A similar proportion of conflicts were recorded for both native and non-native species within all four categories (Figure 3). For native species, economic impacts accounted for 47% (48/103), health impacts accounted for 36% (37/103), environmental impacts accounted for 11% (12/103) and social impacts accounted for 6% (6/103). For non-native species, economic impacts accounted for 45% (42/92), health impacts accounted for 36% (32/92), environmental impacts accounted for 16% (15/92) and social impacts accounted for 3% (3/92). Feral species were involved in considerably less conflicts, mostly attributable to health and environmental impacts.

**Figure 3.**
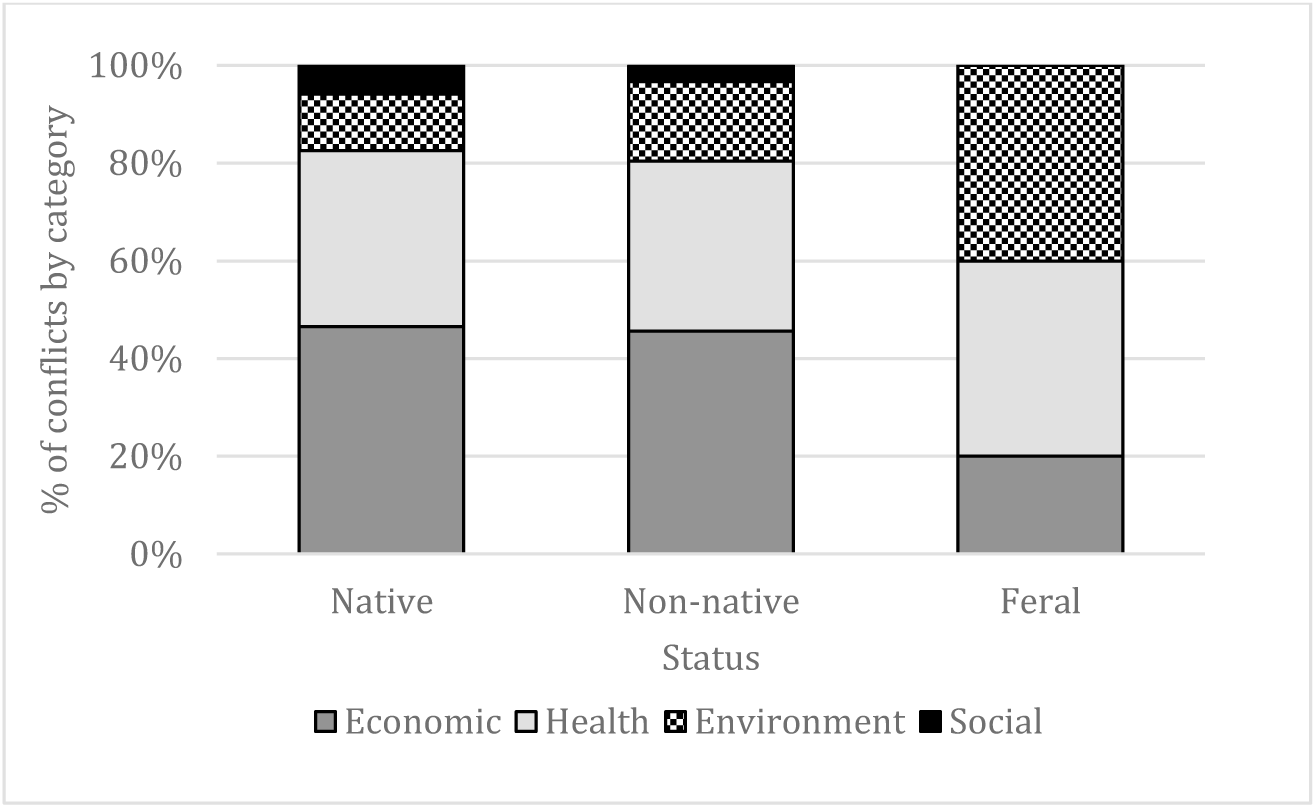
The percentage of conflicts within each category depending on status. Native species: n=103; non-native species: n=92; feral species: n=5.

The total number of mammal species causing conflict within a particular subcategory ranged from one to 30 (Supplementary Tables 5a-5d), with the most common conflicts being reservoir of zoonotic disease (n=30) and reservoir of disease (n=22).

### Impact scores and ranking of conflicts

#### Reproducibility of the scoring

Of the 200 conflicts, the three reviewers agreed on the impact score in 92 cases (46%); there was a maximum deviation of one score in 89 cases (44%), a maximum deviation of two scores in 18 cases (9%) and a maximum deviation of three scores in 1 case (1%). Overall, there was no statistically significant difference between the independently assigned impact scores (*χ*^2^(2)=3.057, *p*=0.217). However, confidence scores were found to differ statistically between reviewers (*χ*^2^(2)=144.54, *p*<0.001); all three agreed in 33 of the 200 conflicts (17%), two agreed in 131 cases (65%), with a deviation of one or two assigned by the third reviewer, and all three assigned a different confidence score in 36 cases (18%).

#### Conflict scoring and ranking

Using the median impact scores across the three reviewers, seven of the 200 conflicts (4%) were assessed as impact score 5, 20 (10%) as impact score 4, 30 (15%) as impact score 3, 88 (44%) as impact score 2 and 55 (27%) as impact score 1 (Supplementary Table 5a-5d). Six of the seven conflicts assessed as having the highest impact score of 5 (all with confidence rating of high) were economic and one was a health conflict. These were (in order of highest approximate economic cost – see Supplementary Material S2 for full detail): (1) rabbit damage to agricultural interests (£120 million), (2) badger involvement in the transmission of bTB to cattle (>£100 million), (3) brown rat damage to agricultural interests (£54 million), (4) rabbit damage to forestry (£43 million) (5) badger damage to agricultural interests (£39 million) (6) grey squirrel damage to forestry (£28 million, not including carbon capture estimates) and (7) brown rat damage to property (£14 million, based on local authority costs only). In all four categories, the majority of conflicts were scored as impact 2 (economic: 35% (32/91); health: 46% (33/71); environmental: 55% (16/29); social: 78% (7/9)). Despite two health conflicts being the most common amongst all mammal species, generally, health conflicts were scored low, with over 90% (64/71) scoring either a 1 or 2.

Analysis of impact scores in relation to native/non-native status and conflict category, found a statistically significant effect of category, when considered as a main order effect (*χ*^2^(3)=62.90, *p*<0.001) but not status (*χ*^2^(1)=0.421, *p*=0.516). Reviewers assigned a greater score in relation to economic impacts, followed by environmental, health and then social impacts (Figure 4). When conflict category was considered in relation to status, there was no statistically significant interaction term between the two (*χ*^2^(3)=4.839, *p*=0.184)

**Figure 4.**
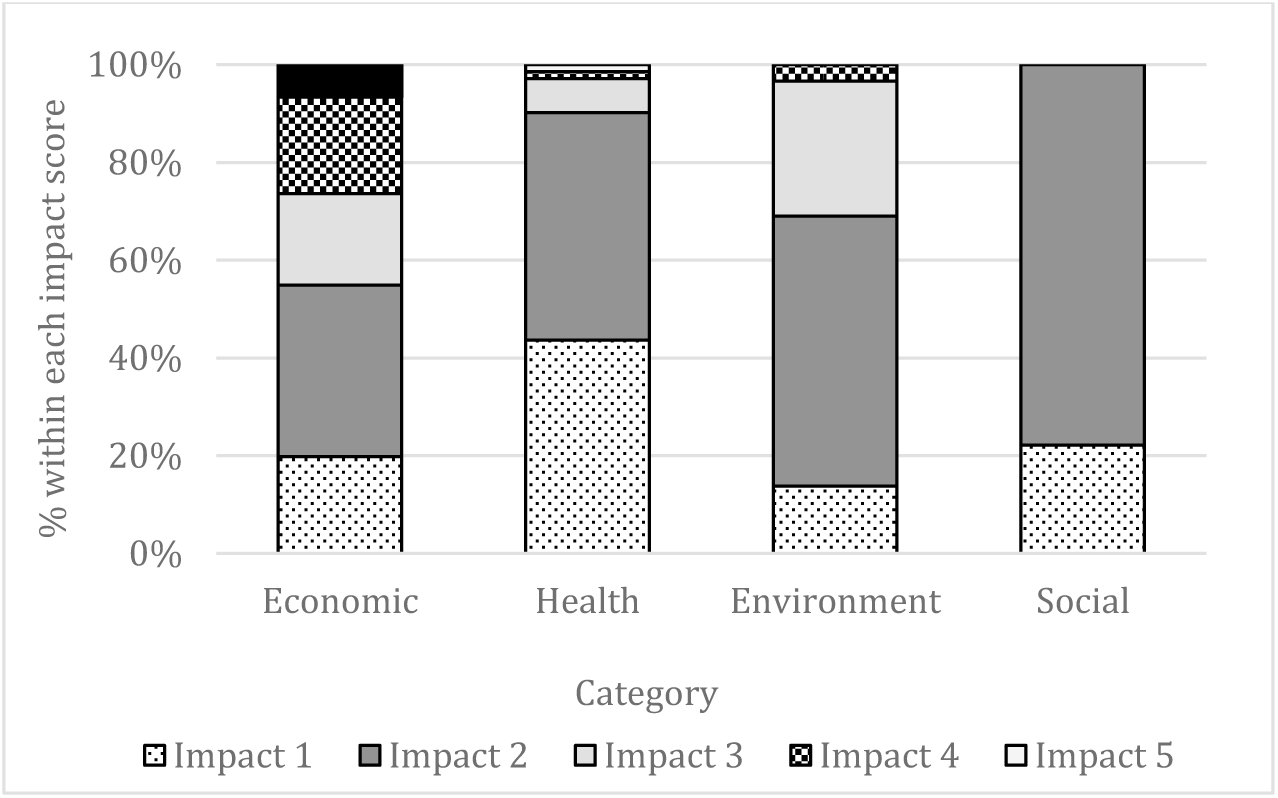
The proportion of conflicts depending on impact score (1-5) within the four categories. Economic: n=91, health: n=71, health: n=29, social: n=9.

Analysis of impact scores, in relation to native/non-native status and conflict subcategories, showed both as having a statistically significant effect (*χ*^2^(1)=33.63, *p*<0.001; *χ*^2^(31)=640.18, *p*<0.001, respectively), with a significant two-way interaction term also identified (*χ*^2^(31)=155.56, *p*<0.001). Across all economic impacts, non-native species received a statistically significantly greater impact score than natives in relation to damage to woodland, damage to agricultural interests, damage to property, conflict with transport and conflict with utilities (all *p*<0.05). Conversely, native species received a significantly higher score due to conflicts with construction and development and the predation of livestock (both *p*<0.001). Across health impacts, the model predicted a statistically greater impact score for non-native species as reservoirs of zoonotic diseases only (*p*=0.008). With regards to the environment, non-native species received a statistically significant greater impact score for damage to conservation habitats and their threat to biodiversity and genetic integrity (all *p*<0.05). In terms of social impacts, non-native species received a greater impact than natives with respect to public nuisance only (*p*<0.01).

#### Maximum impact

The number of separate conflicts per species ranged from one to eleven (Table 4), with sika deer involved in the most conflicts. Species scoring highest in maximum total impact (i.e. the sum of all median conflict scores for a particular species) were badger, rabbit, roe deer, fallow deer and red deer. Generally, these species also scored highly within the separate categories (except for social conflicts), with badgers scoring highest in both maximum economic and health impacts. The highest maximum environmental impact was produced by sika deer, followed by roe deer, red deer and feral cat. The highest maximum social impact was observed in the red fox, followed by rabbit, wild boar, pine marten, Natterer’s and serotine bat.

**Table 4.** Distribution (i.e. the proportion of Great Britain each mammal species is found, based on presence in 10×10km squares), total number of conflicts and impact scores (maximum and potential) in total and within each of the four categories (economic, health, environmental, social) for all conflict species. Species are listed in order of highest total potential impact. – denotes missing data.

| Species | Distribution | No. conflicts | Maximum impact |  |  |  |  | Potential impact |  |  |  |  |
| --- | --- | --- | --- | --- | --- | --- | --- | --- | --- | --- | --- | --- |
|  |  |  | Total maximum impact | Economic | Health | Environment | Social | Total potential impact | Economic | Health | Environment | Social |
| Badger | 0.746 | 10 | 33 | 21 | 9 | 3 | 0 | 24.624 | 15.670 | 6.716 | 2.239 | 0 |
| Rabbit | 0.833 | 8 | 26 | 17 | 5 | 2 | 2 | 21.649 | 14.155 | 4.163 | 1.665 | 1.665 |
| Roe deer | 0.726 | 9 | 25 | 12 | 8 | 5 | 0 | 18.151 | 8.713 | 5.808 | 3.630 | 0 |
| Red fox | 0.800 | 8 | 19 | 8 | 5 | 3 | 3 | 15.207 | 6.403 | 4.002 | 2.401 | 2.401 |
| Grey squirrel | 0.583 | 7 | 20 | 11 | 5 | 4 | 0 | 11.653 | 6.409 | 2.913 | 2.331 | 0 |
| Brown rat | 0.529 | 7 | 21 | 14 | 4 | 3 | 0 | 11.105 | 7.403 | 2.115 | 1.586 | 0 |
| Red deer | 0.412 | 10 | 23 | 11 | 7 | 5 | 0 | 9.480 | 4.534 | 2.885 | 2.061 | 0 |
| Fallow deer | 0.347 | 9 | 24 | 12 | 8 | 4 | 0 | 8.317 | 4.158 | 2.772 | 1.386 | 0 |
| American mink | 0.533 | 6 | 14 | 9 | 2 | 3 | 0 | 7.457 | 4.794 | 1.065 | 1.598 | 0 |
| Mole | 0.807 | 3 | 9 | 7 | 2 | 0 | 0 | 7.266 | 5.651 | 1.615 | 0 | 0 |
| Hedgehog | 0.824 | 4 | 8 | 2 | 3 | 3 | 0 | 6.592 | 1.648 | 2.472 | 2.472 | 0 |
| Brown hare | 0.680 | 5 | 9 | 7 | 2 | 0 | 0 | 6.119 | 4.759 | 1.360 | 0 | 0 |
| Muntjac | 0.334 | 9 | 17 | 7 | 8 | 2 | 0 | 5.673 | 2.336 | 2.669 | 0.667 | 0 |
| Field vole | 0.564 | 5 | 9 | 5 | 4 | 0 | 0 | 5.075 | 2.819 | 2.256 | 0 | 0 |
| Wood mouse | 0.560 | 3 | 6 | 2 | 4 | 0 | 0 | 3.360 | 1.120 | 2.240 | 0 | 0 |
| Otter | 0.782 | 2 | 4 | 2 | 0 | 2 | 0 | 3.126 | 1.563 | 0 | 1.563 | 0 |
| House mouse | 0.235 | 4 | 13 | 8 | 5 | 0 | 0 | 3.060 | 1.883 | 1.177 | 0 | 0 |
| Stoat | 0.597 | 3 | 5 | 2 | 1 | 2 | 0 | 2.983 | 1.193 | 0.597 | 1.193 | 0 |
| Natterer's | 0.411 | 3 | 7 | 5 | 0 | 0 | 2 | 2.880 | 2.057 | 0 | 0 | 0.823 |
| Sika deer | 0.134 | 11 | 19 | 8 | 5 | 6 | 0 | 2.540 | 1.069 | 0.668 | 0.802 | 0 |
| Bank vole | 0.446 | 3 | 5 | 2 | 2 | 1 | 0 | 2.229 | 0.892 | 0.892 | 0.446 | 0 |
| Common pipistrelle | 0.674 | 1 | 3 | 3 | 0 | 0 | 0 | 2.021 | 2.021 | 0 | 0 | 0 |
| Soprano pipistrelle | 0.619 | 1 | 3 | 3 | 0 | 0 | 0 | 1.858 | 1.858 | 0 | 0 | 0 |
| Brown long-eared bat | 0.600 | 1 | 3 | 3 | 0 | 0 | 0 | 1.799 | 1.799 | 0 | 0 | 0 |
| Serotine | 0.246 | 3 | 7 | 3 | 2 | 0 | 2 | 1.721 | 0.738 | 0.492 | 0 | 0.492 |
| Daubenton's | 0.491 | 2 | 3 | 0 | 3 | 0 | 0 | 1.472 | 0 | 1.472 | 0 | 0 |
| Pine marten | 0.171 | 4 | 6 | 2 | 0 | 2 | 2 | 1.025 | 0.342 | 0 | 0.342 | 0.342 |

Table 4 cont.
| Species | Distribution | No. conflicts | Maximum impact |  |  |  |  | Potential impact |  |  |  |  |
| --- | --- | --- | --- | --- | --- | --- | --- | --- | --- | --- | --- | --- |
|  |  |  | Total maximum impact | Economic | Health | Environment | Social | Total potential impact | Economic | Health | Environment | Social |
| Mountain hare | 0.162 | 4 | 6 | 5 | 1 | 0 | 0 | 0.973 | 0.811 | 0.162 | 0 | 0 |
| Water vole | 0.425 | 1 | 2 | 2 | 0 | 0 | 0 | 0.850 | 0.850 | 0 | 0 | 0 |
| Polecat | 0.269 | 3 | 3 | 2 | 1 | 0 | 0 | 0.807 | 0.538 | 0.269 | 0 | 0 |
| Weasel | 0.530 | 1 | 1 | 1 | 0 | 0 | 0 | 0.530 | 0.530 | 0 | 0 | 0 |
| Common shrew | 0.502 | 1 | 1 | 0 | 1 | 0 | 0 | 0.502 | 0 | 0.502 | 0 | 0 |
| Red squirrel | 0.247 | 2 | 2 | 0 | 2 | 0 | 0 | 0.494 | 0 | 0.494 | 0 | 0 |
| Feral goat | 0.023 | 6 | 13 | 4 | 7 | 1 | 1 | 0.209 | 0.070 | 0.047 | 0.047 | 0.047 |
| Hazel dormouse | 0.199 | 1 | 2 | 2 | 0 | 0 | 0 | 0.397 | 0.397 | 0 | 0 | 0 |
| Chinese water deer | 0.042 | 7 | 9 | 5 | 3 | 1 | 0 | 0.375 | 0.208 | 0.125 | 0.042 | 0 |
| Wild boar | 0.031 | 7 | 9 | 3 | 2 | 2 | 2 | 0.402 | 0.124 | 0.216 | 0.03 | 0.031 |
| Beaver | 0.018 | 6 | 9 | 7 | 2 | 0 | 0 | 0.166 | 0.129 | 0.037 | 0 | 0 |
| Yellow-necked mouse | 0.129 | 1 | 1 | 0 | 1 | 0 | 0 | 0.129 | 0 | 0.129 | 0 | 0 |
| Edible dormouse | 0.008 | 4 | 8 | 7 | 1 | 0 | 0 | 0.064 | 0.056 | 0.008 | 0 | 0 |
| Wildcat | 0.039 | 1 | 1 | 1 | 0 | 0 | 0 | 0.039 | 0.039 | 0 | 0 | 0 |
| Exmoor/Dartmoor pony | 0.005 | 2 | 3 | 2 | 1 | 0 | 0 | 0.016 | 0.011 | 0.005 | 0 | 0 |
| Black rat | 0.006 | 2 | 2 | 0 | 1 | 1 | 0 | 0.011 | 0 | 0.006 | 0.006 | 0 |
| Skomer vole | 0.004 | 1 | 1 | 0 | 1 | 0 | 0 | 0.004 | 0 | 0.004 | 0 | 0 |
| Orkney vole | 0.003 | 1 | 1 | 1 | 0 | 0 | 0 | 0.003 | 0.003 | 0 | 0 | 0 |
| Feral cat | - | 3 | 7 | 0 | 2 | 5 | 0 | - | - | - | - | - |
| Feral ferret | - | 3 | 4 | 2 | 0 | 2 | 0 | - | - | - | - | - |
| Greater white-toothed shrew | - | 2 | 3 | 0 | 1 | 2 | 0 | - | - | - | - | - |

Analyses identified a statistically significant difference between native/non-native species (*χ*^2^(1)=4.235, *p*=0.040), with the latter predicted as having a higher maximum impact overall (*β*=0.513, *z* ratio=2.058, *p*=0.046). When specified as main order effects, total maximum impact was found to increase as a function of body mass (*β*=0.335, *t* ratio=3.598, *p*<0.001) and biomass (*β*=0.184, *t* ratio=4.706, *p*<0.001) but not population size (*β*=0.040, *t* ratio=1.012, *p*=0.311). However, no statistically significant interaction term was identified (*p*>0.05), suggesting that differences in the rate of change for total maximum impact were comparable between native and non-native species, relative to changes in population size, body mass and biomass.

#### Potential impact

Species scoring the highest potential impact were badger, rabbit, roe deer and red fox (Table 4). Generally, these species also scored highly within the separate categories, with badgers scoring highest in both potential economic and health impacts. Roe deer scored the highest potential environmental impact, followed by hedgehog, red fox, grey squirrel, badger and red deer. The highest potential social impact was observed in the red fox, followed by rabbit and Natterer’s bat.

Analysis of potential impact estimates showed no statistically significant difference between native/non-native species (*χ*^2^(1)=0.056, *p*=0.813). When considered as main order effects, total potential impact was predicted to increase significantly in relation to population size (*β*=0.094 *t* ratio=1.980, *p*=0.048) and biomass (*β*=0.215, *t* ratio=4.691, *p*<0.001). When considered in relation to status, there was no statistically significant interaction for biomass (*χ*^2^(1)=0.298, *p*=0.585) suggesting that the rate of increase was comparable across native and non-native species. Body mass was found to have no statistically significant effect on total potential impact when considered as a main order effect (*χ*^2^(1)=1.741, *p*=0.187) or through interaction with status (*χ*^2^(1)=0.088, *p*=0.767). A statistically significant interaction term was identified between status and population size, with total potential impact predicted to increase, relative to population size, at a significantly higher rate for non-native species when compared to natives (*β*=0.179, *t* ratio=2.120, *p*=0.034).

#### Conflict change over time

For many mammal species, a conflict occurring before 2014 (identified during reviews for the Defra policy framework) was still being reported in the scientific literature within the last decade (Supplementary table S3). Twenty-four of the 39 (62%) conflicts reported only before 2014 were economic, followed by eight (21%) health, five environment (13%) and two social (5%). In contrast, 19 of the 39 (49%) conflicts reported from 2014 to 2024 were health, mostly attributable to reservoir of disease or reservoir of zoonotic disease, followed by eight environment (21%), seven economic (18%) and five social (13%).

## Discussion

Human-wildlife conflict is one of the most important sustainable development challenges globally, particularly where ecologically and economically important wildlife impact human livelihood (Braczkowski et al. 2023). It is difficult to robustly value everything in economic terms, wildlife being a particular challenge, therefore alternative methods are often required to determine the threats to people and biodiversity (HM Government 2018). In this paper we carried out a review of the literature to identify conflicts associated with British mammal species and applied a Generic Impact Scoring System to rank their impacts. Our results, both the comprehensive audit of conflicts associated with British mammals (Supplementary Material S2), and ranking tables depicting species’ maximum impact and potential impact based on geographic distribution overall and in the different category areas, are insightful visual tools that may be used to inform future management priorities and/or policy design. We found that the highest proportion of conflicts reported in the literature were economic, with the majority (83%) of mammal species identified as causing conflict in Great Britain associated with at least one economic conflict. A similarly high proportion of both native and non-native species (86% and 82%, respectively) caused economic damage. All of the seven conflicts assessed as having the highest impact score of 5, involving rabbits, badgers, brown rats and grey squirrels, were measurable on an economic scale, suggesting that conflicts associated with a monetary value are inherently easier to quantify compared with those that are not.

Where conflicts were associated with an economic value, we estimated a conservative total cost of human-wildlife conflicts in Great Britain (as reported in Supplementary Material S2) in excess of £0.5 billion. Wildlife interactions with humans, however, are not always adverse and can result in neutral or even positive effects (Peterson et al. 2010; Nyhus 2016). For example, rabbits caused the highest economic damage (estimated at £120 million), but they are considered a keystone species in certain UK grass and heathland habitats (Manchester and Bullock 2000; Bell, Endean, and Mountjoy 2021). Rabbit populations have also declined significantly in the UK in recent years, mostly driven by outbreaks of myxomatosis and Rabbit Haemorrhagic Disease Virus (RHDV1, RHDV2) (Petrovan, Ward, and Wheeler 2011; Bell et al. 2019), but also as a result of habitat degradation (Bell, Endean, and Mountjoy 2021). This has led to concerns regarding the impact of such population reductions on other species, such as carnivores, that rely on them as a food source (Sainsbury et al. 2019). One such carnivore is the red fox, which has been widely persecuted in Great Britain due to its perceived status as a pest species (Macdonald and Johnson 2015). Their impact on agriculture has been estimated to cost £20 million annually, most of which is attributable to direct and indirect effects of livestock predation (Baker, Harris, and White 2006). However, fox predation on rabbits, which have a higher impact on agriculture, is estimated to create an indirect economic benefit to farmers of at least £11 million annually and may even be approaching an economically neutral impact on agriculture (Baker, Harris, and White 2006). Human and wildlife interactions can be complex, and in the case of foxes and rabbits, the latter of which is considered an invasive non-native species in Great Britain, paradoxical. Understanding the costs-to-benefits trade-offs that underlie conflict and/or coexistence in human and wildlife systems are critical for informing conservation and development efforts, in terms of sustaining on-going human-wildlife coexistence or intervening to reconcile human activities with the needs of wildlife and vice versa (Balasubramaniam et al. 2021).

Categorising British mammal species as invasive or “problematic”, is an emotive subject and often polarised depending on their interaction with different groups of people. The mammal species we ranked highest for maximum and potential impact overall (including in both economic and health categories), was the badger, which is largely viewed by the public as an iconic British species but considered a significant problem in parts of the farming community (Cassidy 2019). Like many of the higher scoring impact species, badgers are widely distributed throughout Great Britain; although they naturally inhabit woodland and pasture (Gorman 2008), expanding urbanisation and habitat fragmentation has led to them becoming increasingly more common in urban areas, such as gardens and allotments (Fung et al. 2024). In the UK, badgers cause significant economic conflict, both in terms of the transmission of bTB to cattle (estimated >£100 million) and damage to agricultural interests (estimated £39 million). Strategies to eradicate bTB have focussed on badger culling, which can result in complex epidemiological outcomes, with both positive and negative effects on the incidence of bTB in cattle (Griffin et al. 2005; Donnelly et al. 2006) and has created a schism in public opinion (Enticott 2015; Naylor et al. 2017). As such, the most recent UK government’s policy for controlling bTB in badgers aims to phase out culling by 2026, focussing on badger vaccination, along with a number of other measures. These include an acceleration of work to develop a cattle vaccine, a major survey of the badger population and a wildlife surveillance programme to provide an understanding of the prevalence of bTB in the remaining badger populations, which is largely unknown (Defra 2024). Part of the UK government’s 25-year Environment plan for enhancing biodiversity is to reduce the impact of endemic disease, therefore understanding such population dynamics are key factors in establishing how much intervention may be required in order to move from human-wildlife conflict to coexistence.

Often, wildlife policies are implemented in order to move from actual undesired to desired population sizes, which are typically motivated by the species imposing an external cost of some kind (e.g. to agriculture or forestry), or benefit (e.g. threatened or considered highly valuable) (Gren et al. 2018). All six deer species in Great Britain cause damage to crops and woodlands, including conservation habitats, to varying degrees, but are also considered a valuable resource in terms of recreational hunting and food; venison is the main income each year from the overall management of wild deer in Scotland with over 90% of all deer culled on National Forest Estate land destined for the food industry (Pepper, Barbour, and Glass 2019). Deer are protected by the Deer Act 1991 and Hunting Act 2004, which criminalises various activities including poaching, using certain ammunition, hunting with dogs and shooting during closed seasons, but does allow legal hunting at certain times of the year. In some cases, a different approach may be required, particularly with respect to non-native species; for example, in response to a joint Defra and Forestry Commission consultation on managing deer populations in England, the Mammal Society (2024a) supported recommendations to remove the close season for sika stags, to limit their further spread and hybridisation with red deer. Managing deer population sizes to desired levels, in order to balance negative impacts with revenue retention, is key for determining policy design so that it adjusts for the asymmetric allocation of costs to benefits among stakeholders that constitute a threat to wildlife (Gren et al. 2018).

One other aspect of the government’s 25-year plan for enhancing biosecurity aims to tackle invasive non-native species (HM Government 2018). Overpopulations of deer and grey squirrels are cited as a major threat to woodlands; grey squirrel abundance and distribution has expanded rapidly since they were introduced to Great Britain in the late 19th century (Gurnell and Hare 2008), causing an estimated direct cost of £28 million to forestry each year. They have also contributed to the decline of the native red squirrel, which is an iconic conservation species in Great Britain. Whilst red squirrels are legally protected in the UK (e.g. Wildlife and Countryside Act 1981; Environment Act 2021), grey squirrels are an invasive non-native species and as such, are afforded no protection. In England, the action plans for red and grey squirrels are interlinked, with the focus on red squirrel conservation alongside the need for grey squirrel management (UKSA 2023). The 25-year plan states that where it is not feasible to eradicate species that have become too widely established, their threat should be neutralised by managing them effectively. However, drivers of human-wildlife conflict, such as urbanisation, may favour non-native species such as the grey squirrel and brown rat, which are able to thrive and coexist with humans (Fung et al. 2024); whilst urban and rural grey squirrels select similar habitats, urban populations can occur at much higher densities than their rural equivalents (Merrick, Evans, and Bertolino 2016). Inner urban areas may provide refugia for non-native species (Boon et al. 2023) potentially offering protection from natural predators (e.g. pine marten predation of grey squirrels) (Twining, Montgomery, and Tosh 2020), thus hindering efforts to manage overall populations and negatively impacting on conservation efforts.

Urbanisation and anthropogenic factors, amongst others, are thought to have greatly influenced the emergence and distribution of zoonoses (Rahman et al. 2020), with more than 60% of human pathogens considered zoonotic in origin (Taylor, Latham, and Woolhouse 2001). Although we found the highest proportion of reported conflicts were economic, the two most common conflicts were health related (reservoir of zoonotic disease and reservoir of disease). Generally, though, we deemed the impact score of health conflicts to be low (mostly 1 or 2), therefore we assumed that most health impacts were likely local, mild, short-to mid-term and with reversible effects (see Table 1). We found that non-native species were predicted to have a greater impact score as reservoirs of zoonotic disease compared with native species, highlighting the importance of surveillance, which is crucial to preventing and controlling the spread of disease, in particular zoonotic disease (Rahman et al. 2020). Portals such as the Non-native species secretariat (www.nonnativespecies.org) are useful in this respect, to provide current information on the status of a range of non-native species within Great Britain. We found that the majority of “new” conflicts (i.e. identified in our literature searches after 2014 only) were disease related, which may reflect a shift of focus leading to increased reporting in this area, possibly in light of several international public health emergencies of zoonotic origin within the last ten years, such as Middle East respiratory syndrome (MERS), Ebola and Covid-19 pandemics (Sun et al. 2024). Whilst there is extensive research on the pathology and epidemiology of disease in a wide range of British mammal species, as reported in Supplementary Material S2, it is evident that for some species, there is an inference of disease presence and/or zoonotic potential or a general lack of information (e.g. some hantaviruses (Vapalahti et al. 2003) and Ljungan virus (Salisbury et al. 2014; Rossi et al. 2021) in rodents; Babesia spp. in badgers (Guardone et al. 2020)), which suggests further research may be warranted. Understanding disease dynamics in wildlife is intrinsically linked to biodiversity conservation, as anthropogenic activities such as urbanisation and habitat destruction, are likely to lead to increased interaction between humans and wildlife (Bedenham et al. 2022).

Human-wildlife conflict has long been associated with negative impacts on human interests, although it is evident that interactions between people and wildlife are more varied and complex (Hill 2021). More recently, the emphasis has shifted from conflict between humans and wildlife and its mitigation, to a coexistence approach (Gross et al. 2024), or at least one that constrains conflict or adapts shared spaces to mitigate conflict. Foxes are one of the most globally widespread carnivores (Soulsbury et al. 2010; Castan eda et al. 2022) and are highly adapted to living in modified urban and agricultural areas where they exploit anthropogenic foods (Castan eda et al. 2022). Although reports of social disruption, such as bin-raiding, is evident, Adaway et al. (2025) found that public attitudes towards foxes were not predicted by urbanisation type (i.e. urban, suburban or rural) or geographic region and may in fact be shaped by broader societal narratives about foxes amongst the general public. Socio-psychological research shows that people are unlikely to change their behaviour unless it aligns with what they already know, value or believe (Toomey 2023). As species richness is generally lower in heavily urbanised areas (McKinney 2002, 2006; Ramadhan 2024) there is less opportunity for people to physically and emotionally connect with nature (Soga and Gaston 2016; Cox and Gaston 2018; Richardson et al. 2022), a factor considered to impact on mental health and wellbeing (Sibthorpe and Brymer 2020; White et al. 2021), and conservation initiatives (Richardson et al. 2020; Barrows et al. 2022). Understanding local perceptions and incorporating them into communication strategies is key to fostering deeper connections with nature and support for biological conservation efforts in general.

In many cases, species that caused the most conflict were amongst the most widely distributed throughout the British Isles, e.g. badger, rabbit, roe deer and red fox. Generally, though, we determined that the majority of British mammals do not cause major conflict, which for some species may be due, in part, to their limited geographical distribution. However, our analyses identified that total potential impact tended to increase as a function of biomass and population size, therefore a GISS approach may be particularly beneficial for monitoring existing or recently introduced species that have the potential to expand beyond their current range. For example, beavers were reintroduced to Great Britain around 20 years ago and are considered keystone species due to their ability to alter ecosystem structure and function through their engineering activities (Puttock et al. 2017; Auster, Puttock, and Brazier 2019). Whilst beavers are generally considered as having a positive impact on biodiversity, we identified six conflicts of interest with humans; although they ranked 38^th^ overall based on potential impact, likely owing to their current limited distribution, they ranked joint 12^th^ overall based on maximum impact. Whilst the reintroduction of beavers across Great Britain is currently carefully managed, highlighting and monitoring potential issues at this stage is essential for ensuring the status of this species remains positive rather than possible future perception as an invasive species. Currently, beavers are one of the eight conflict species classified as being at imminent risk of extinction in Great Britain (wildcat, beaver, red squirrel, water vole, hedgehog, hazel dormouse, Orkney vole and serotine bat), with mountain hares also classified as “near threatened” (Mammal Society 2024b). Where conflict involves a vulnerable species, the emphasis should encourage a shift of focus from negative aspects of human-wildlife relationships, as often represented within a conservation conflicts framing (Marchini et al. 2019) to one that acknowledges, and incorporates the idea, that human-wildlife relations are not inherently or solely negative in nature, with wildlife species having significant value that is tolerated where they coexist with people (Hill 2021).

The findings reported here must be viewed in light of several limitations. Although an extensive literature search was conducted to identify conflicts associated with British wildlife, it is likely that some conflicts were not identified or cost estimates were not available, and therefore not reported. Conflicts that were identified before 2014 only are more than likely still occurring or may not be reported as frequently in the scientific literature. Likewise, conflicts reported after 2014 only may have been occurring prior to this and again, either missed in the original literature searches or less frequently reported. Whilst the conflict descriptions given in Supplementary Material S2 are not exhaustive and based only on the searches returned, they likely represent reasonable overviews and are sufficiently robust to provide an overall audit of the current national importance of conflicts between human and wildlife interests in Great Britain. However, individual assignments of scale may need to be revised as further information emerges. A further limitation is the discrepancy in confidence ratings assigned to conflicts by the three reviewers; whilst they were largely consistent in the impact scoring of mammal species, they were less confident of the assignments, which is subjective and likely reflects individual interpretation based on the depth of evidence presented.

Another limitation of our findings is the lack of data for some species. Feral species were excluded from some of the analyses due to the small number of conflicts in Great Britain, although their impacts can be significant. Data regarding feral cats are limited due to the difficulty in differentiating between feral and domestic cats (Hand 2019), the uncertainties of feral cat population estimates (based mostly on urban populations) and the lack of known distribution within Great Britain. Cats are one of the most ubiquitous mammal species globally (see McDonald and Skillings (2021)) and one of the most destructive, with free-ranging cats suggested as having a major impact on the mortality of several bird species in the UK (Baker et al. 2008). They are also the biggest threat to the survival of Scottish wildcats due to introgression (Fredrikson 2016). Given their domesticated origins, feral cats have no native range and their ubiquitous adverse impacts on native wildlife around the world has led to them being labelled amongst the world’s worst invasive species (Lowe, Browne, and Boudjelas 2000). From a biodiversity conservation perspective, a wealth of evidence suggests that feral cats should be controlled and ideally removed from places where their impact is detrimental, often as an urgent priority (Trouwborst, McCormack, and Camacho 2019).

## Conclusions

The GISS approach we applied here is a useful tool for identifying individual conflicts for individual species but also provides a broader assessment of where human-wildlife impacts may be affecting socio-economic and biodiversity interests. It is clear that some mammal species cause considerable damage and intervention is required. However, mitigation measures should be carefully considered and designed, weighing all of the costs and benefits associated with a particular species, to achieve the desired outcome (e.g. population reduction, eradication). Wildlife is a valuable commodity, both physically and emotionally, and with many British mammals facing local extinction, we should perhaps accept and/or tolerate some negative interactions rather than seek absolute conflict resolution. Efforts should be made to ensure a balance, where possible, between the need to conserve and enhance biodiversity, the interests of the general public and the interests of those directly affected (Defra 2010), to ensure we progress from the negative association with wildlife interactions towards one where humans and wildlife can coexist.

## Data availability statement

The data that support the findings of this study are available as supplementary material.

## Supporting information

**Supplementary Table S1**. Comprehensive list of articles included in the conflict review and analyses, including study location, source type, date, conflict, species, duplication (i.e. the article was applicable to multiple conflicts and/or more than one species).

**Supplementary Material S2**. Detailed description of all of the conflicts identified in the scientific literature for 48 mammal species of the British Isles.

**Supplementary Table S3**. List of species causing conflict, including order, status, population size, body mass, biomass and type of conflicts recorded over time (pre-2014 only; both pre-2014 and 2014 to present; 2014 to present only). The median impact score (1=minimal known impact to 5=highest known impact) and confidence rating (1=low; 2=medium; 3=high) are shown after each conflict [x,x]. The highest impact scores are shown in bold.

**Supplementary Table S4**. Summary of model formulae applied when analysing the effect of non-native status and several independent covariates (population size (*circa* 2018), body mass (kg), and biomass (kg) on the impact scores (maximum or potential) of mammal species in Great Britain. Analyses were performed using generalised linear models (GLM) fitted using the *glmmTMB::glmmTMB* function (version 1.1.10; Brooks et al., 2017). Models were fitted with a gaussian error distribution, with “identity” link function, with impact scores log-transformed (log (n + 1)) to conform with model assumptions. Covariates were also log-transformed.

**Supplementary Table S5a-d**. Total number of mammal species causing conflict within the (5a) 15 economic subcategories; (5b) Six health subcategories; (5c) Seven environmental subcategories; (5d) Four social subcategories, and their median impact score. Status categories: Na=native; NN=non-native; Fe=feral.

