## Supplementary Material S2 for "Identifying and quantifying conflicts between humans and terrestrial mammals in Great Britain"

Detailed description of all of the conflicts identified in the scientific literature for 48 mammal species of the British Isles. Conflicts are represented by a two-character notation representing the category (A = economic; B = health; C = environment; D = social) and a number as a unique identifier of that conflict in that category.

### Artiodactyla

Roe deer - *Capreolus capreolus*

Introduction

The roe deer is native to Britain arriving approximately 11,000 years ago (Yalden 1998). They can be found in a variety of habitat types but are particularly associated with open mixed, deciduous or coniferous woodland with higher densities in mosaic habitats which incorporate agricultural land (Putman 2008). This is likely to reflect the relatively high proportion of woodland edge found in such habitats that roe deer can exploit as cover close to food resources. Despite being selective browsers, with deciduous trees and shrubs forming an important part of their diet, roe deer remain versatile and opportunistic, varying their diets according to the foods that are available (e.g. seasonally abundant fruits, seeds or mushrooms) (Putman 2008).

The roe deer is widely distributed throughout Britain, particularly in Scotland and northern England, with a patchy distribution in Wales and central and southeast England (Mammal Society 2023; British Deer Society 2023b). The geographical range and population size of roe deer has increased over the last 20 years with an estimated 265,000 roe deer living wild in Britain: 120,000 in England, 122,000 in Scotland and 22,000 in Wales (Mathews et al. 2018).

A.1. Damage to forestry

Roe deer will take the buds, shoots and leaves of deciduous shrubs and trees; they can browse to heights of 120cm by standing on their hind legs to reach vegetation above their usual browsing height of approximately 75cm (Putman 2008). Deer browse young trees and can affect the way in which trees grow, thus reducing their economic value as timber (White, Ward, et al. 2004). Bark damage can also result in fungal infection and subsequent branch or tree death, which causes further economic losses to foresters (Arnold et al. 2018). Bucks may mark territories in spring and summer by fraying or rubbing bark from saplings using their antlers (Putman 2008).

Plantation forests in the UK commonly suffer from tree browsing by roe deer (Welch et al. 1991; Gill 1992). Under heavy and repeated leader browsing, the entire height of annual growth can be continually lost thus requiring a longer establishment period for plantations (Ward et al. 2004). This, in turn, leads to increased costs for foresters from factors such as increased need for weeding and will also subsequently increase the rotation period of plantations (Ward et al. 2004). In addition, leader browsing may cause trees to develop multiple leading shoots and resulting in multiple stems being present at harvest (Gill 1992). Although the same volume of wood may be produced by the tree, the stems which are produced will be of a smaller diameter and will therefore be of lower value (Ward et al. 2004; Pepper, Barbour, and Glass 2019).

In the east of England, White, Ward, et al. (2004) estimated that the total cost of deer to forestry and plantation woodlands was between £616.6K and £862.2K. In Scotland, Forestry and Land Scotland (FLS) reported up to 150 million young trees in national forests and land were vulnerable to damage from deer with an estimated cost in the region of £3M annually (Forestry and Land Scotland 2021). FLS estimated that the total costs to forestry for the four species found there (roe, red, sika and fallow) was £8.71M although revenue generated from deer management (e.g. from venison, sporting income) was £1.75M (Pepper, Barbour, and Glass 2019).

A.2. Damage to agricultural interests

Damage to cereals is principally associated with fallow, roe and red deer with most of the damage to roots and fruits being attributed to roe deer (White, Smart, et al. 2004). A questionnaire survey of 1,192 respondents conducted for the British Deer Society in 1995 found that 69% of farmers had deer on their holdings, of which roe and fallow deer were the most frequent (Doney and Packer 2002). Roe deer inhabiting agricultural landscapes are likely to take mostly cultivated crops throughout the year, particularly cereals, and may merely supplement their diets by browsing within nearby woodland

(Putman 2008). Damage to cereals by roe deer has been reported to be primarily restricted to the period from March to May as crops were able to recover from prior damage with no significant yield loss detected at harvest (Putman 2008).

White, Smart, et al. (2004) estimated the total annual cost of deer damage to agriculture (mostly cereal damage) in the east of England to be £1.92-£4.57M, with two thirds of the damage being caused by fallow and roe deer and red deer being the next most significant species. More recently, some individual landowners have reported losses of over £1M per year to deer damage (Tubby 2022).

A.3. Damage to vehicles (RTAs)

The increases in the numbers and ranges of deer in Britain, coupled with the increases in volumes of traffic on the roads, have led to understandable increases in the frequency of road traffic accidents (RTAs), often referred to as deer-vehicle collisions (DVCs) (Hothorn et al. 2015). Estimates of deer-related traffic accidents in the UK range from 42,000 to 74,000 annually (Deer Aware 2019; Deer Collisions 2023). In 2017, the national estimate of DVCs in Scotland was 4,000 to 12,000 per year (Pepper, Barbour, and Glass 2019); however, records from Scotland are only a representative sample of the total DVCs with data suggesting they are continuing to increase (Pepper, Barbour, and Glass 2019) with revised estimates of 8,000 to 14,000 per year in Scotland alone (Langbein 2018). Such collisions can be costly due to the generally large body size of deer causing high levels of damage to vehicles and the associated delay, repair, replacement and insurance costs. Damage to vehicles alone (private and commercial) is estimated to be at least £17M (Deer Collisions 2023).

As deer are not distributed evenly throughout Britain, the areas that combine high deer densities with high traffic densities will be those that are subject to the most collisions. The deer-vehicle collisions project (Deer Collisions 2023) estimated that of the total number of DVCs in Britain every year, approximately 80% would be likely to occur in England, particularly in the south-east, due to increased traffic density, the numbers of deer and their distributions. In contrast, Langbein (Langbein 2008) estimated that the risk of being involved in a DVC was twice as high per driven mile in Scotland than England or Wales. The roe deer is the second most commonly involved species for deer-vehicle collisions in England each year, accounting for 32% of all collisions (Deer Collisions 2023; RSPCA 2022). Fallow account for 40%, muntjac account for 25% (RSPCA 2022) and red, sika and Chinese water deer cumulatively account for the remaining 3%. This means that roe deer could be responsible for £5.44M worth of material damage to vehicles in England (compared to £4.32M in 2013: Langbein (2007)).

B.1. Health risks from RTAs

Details on the levels of human injuries sustained as a direct result of deer are very difficult to obtain, as the Department of Transport road accident injury statistics form (STATS19) does not require for the type of animal to be distinguished unless it is a ridden horse (Langbein 2007). However, it is estimated that human injury occurs in around 1% (Scotland: Pepper, Barbour, and Glass (2019)) and 1.5% (England: Deer Collisions (2023)) of all deer collisions. Estimates at a UK level suggest there are more than 450 DVCs a year resulting in human injury, with 10-20 fatalities (Pepper, Barbour, and Glass 2019) and hundreds more serious or slight injuries per year due to accidents involving deer, either through direct collisions or swerving to avoid deer (RSPCA 2022). In 2017, 70 of the 450 DVCs reported were estimated to be in Scotland, although the actual number may be in excess of 120 per year (Pepper, Barbour, and Glass 2019). National Travel Survey data indicate that only around 25%% of all injury accidents are reported to police (DfT 2024), therefore the actual number of human injury collisions involving deer in the UK may be as high as around 1,800.Up to 20% of all deer-vehicle collisions involving deer are expected to leave live casualties that require dispatch at the side of the road and many other deer are likely to run away from the scene of the accident and subsequently suffer or die of their injuries (Langbein 2007). Therefore, the overall toll of deer injured during collisions with vehicles that are not killed outright, based on the countrywide annual estimate of 42,000 to 74,000 DVCs, could be between 8,400 and 14,800 every year. Traffic collisions can represent an important cause of annual mortality for deer with losses to total populations being 4-7% for roe deer (Langbein 2007).

Of 1,646 deer carcasses collected in west and southwest England by Delahay et al. (2007), 14% were accessible to the public, many by way of being involved in vehicle collisions. Of these 3.4% cultured positive for *Mycobacterium bovis* (the causative agent of bovine tuberculosis) and 2.2% had visible tuberculous lesions (CSL 2006). In roe deer specifically, prevalence may be between 0.4-1.9% (Pepper, Barbour, and Glass 2019), therefore the risk of infection is clearly low. In a number of reviews of zoonotic bovine TB in the UK, there have been no reports of a single human case arising from contact with deer (de la Rua-Domenech 2006; Davidson et al. 2017).

B.2. Reservoir of disease

Deer represent a major vertebrate host for all feeding stages of the tick *Ixodes Ricinus* in the UK and may play a role in the persistence of tick-borne pathogens (Johnson, Golding, and Phipps 2021). In Europe, infection with the tick-transmitted bacteria, *Anaplasma phagocytophilum*, is common in livestock (Johnson, Golding, and Phipps 2021) and can cause anaplasmosis in humans, although cases are rare (Azagi et al. 2020). It has been detected in a several species in England; Robinson, Shaw, and Morgan (2009) reported that anaplasmosis was common in deer in the New Forest, with fallow, roe, red and sika deer all naturally infected. Duscher et al. (2020) also reported the presence of *A. phagocytophilum* in roe deer and muntjac in Thetford Forest, although at a low prevalence in both species (6.6% roe deer, 1% muntjac). Duscher et al. (2020) concluded that although deer may be locally important in maintaining tick populations, they probably have a limited reservoir role for anaplasmosis and the risk to public health is likely low.

Schmallenberg virus (SBV) affects domestic ruminants causing subclinical or mild disease in adult animals, although exposure of pregnant females can cause abortion, stillbirths and congenital malformations in offspring (Tarlinton et al. 2012). SBV was identified and confirmed in Britain for the first time in 2012 on four sheep farms in Norfolk, Suffolk and East Sussex (AHVLA 2012). It was suspected that wild ruminants would also be susceptible to SBV because other viruses of the Simbu serogroup (e.g. Akabane virus) can infect wild ungulates, although they do not typically cause clinical disease in wild species (Garigliany et al. 2012). In 2012, Barlow et al. (2013) confirmed that wild British red and fallow deer (from East Anglia) can be infected by SBV and thus could be a transient reservoir host for the virus. By 2019, it had been confirmed that SBV was also present in roe deer and muntjac (Southwell, Sherlock, and Baylis 2020). These authors reported that, of 53 viable roe deer samples submitted for testing (from a wide coverage of Great Britain), six (11.3%) were positive to detection of SBV antibodies (using virus neutralisation (VNT) testing, which is considered a more accurate test than ELISA). They concluded that the overall prevalence of SBV in wild British deer was 13.8% (using VNT) to 22.1% (ELISA), suggesting that as many as 440,000 deer (of an approximately 2 million UK wild deer population) may have been exposed to SBV. Seropositivity rates are likely to vary depending on latitude (e.g. SBV rates are expected to be lower in Scotland where more deer are present), which concurred with findings that deer in the south were more likely to be SBV-positive (Southwell, Sherlock, and Baylis 2020). The latter study highlighted the significantly extended area over which SBV has been reported since 2012, raising concerns for the welfare of wild and farmed deer, and the transmission risk to livestock.

Gastrointestinal (GI) nematodes are among the most important causes of production loss in farmed ruminants (Chintoan-Uta et al. 2014). Deer are known to harbour several nematodes potentially harmful to livestock although there is no direct experimental evidence for cross-transmission between the species (Chintoan-Uta et al. 2014). Alleles for benzimidazole (BZ) resistance were identified in *β-tubulin* of *Haemonchus contortus* of roe deer found in areas of extensive or intensive livestock farming, with phenotypic resistance confirmed *in vitro* by an egg hatch test (Chintoan-Uta et al. 2014). These authors concluded that nematode cross-transmission between deer and livestock could occur and further studies should be carried out to establish the presence of phenotypic BZ resistance *in vivo* and whether cross-transmission occurs in the field. *Mycobacterium avium paratuberculosis* (MAP), which is responsible for Johne's disease, has been found to be quite prevalent in the roe deer of eastern Scotland, often causing significant levels of mortality in localised areas (Fletcher 2002). In contrast, the prevalence of the gene for MAP was found to be low in roe deer (3%) in England (Pearce et al. 2023).

The roe deer has been found to be susceptible to foot-and-mouth disease under experimental conditions but there were no cases of infection reported during the outbreak of 2001 (Putman 2008).

B.3. bTB transmission

The spread of *M. bovis* (which causes bovine tuberculosis (bTB)) in cattle is of importance to both farmers, whose livelihoods are affected, and the national economy. The total cost to the British Government of bTB in cattle is estimated to cost taxpayers over £100M every year (Defra 2024). Delahay et al. (2007) reported the average prevalence of *M. bovis* in roe deer in cattle bTB hotspots in South-West England as 1.02%, therefore the risk of transmission is low. The risk of exposure of cattle to bTB by roe deer has been estimated to be an average of 25% of that posed by badgers, the putative main wildlife reservoir of *M. bovis* in England and Wales (Ward et al. 2009). Although the main bTB host system in the UK is currently cattle-badger-fallow deer, there is also a potential risk posed by a cattle-roe deer-red deer system (Hardstaff et al. 2014). In other countries, roe deer are suspected of being spillover hosts rather than reservoir hosts; Lambert et al. (2016) reported a low prevalence of 0.49% in an endemic area of France but concluded that the pathology of the disease in roe deer (tuberculous lesions) had the ability for bacterial excretion and therefore transmission to other species.

B.4. Reservoir of zoonotic disease

Tick-borne viruses, such as louping ill, have been detected at subclinical levels, both in wild and farmed examples of roe deer, but appear to mostly occur in the presence of other predisposing factors (Fletcher 2002). Antibodies to tick-borne encephalitis virus (TBEV) were found in approximately 4% of culled deer across England and Scotland; of the 1,323 deer sampled, 663 were roe deer of which 27 (4.1%) were ELISA-positive (Holding et al. 2020). The detection of TBEV is important as it can infect humans, although the prevalence in deer appears to be quite low and generally localised; for example, in the Norfolk/Suffolk (Thetford Forest) focal area, serological evidence has shown a high prevalence of TBEV exposure (47.7% in the deer species found there; red, roe, fallow, muntjac) (Holding et al. 2020). Antibodies to the agent involved in Lyme disease, which affects humans, have been found in roe deer in the New Forest, making them a likely reservoir for adult ticks but not a route of transmission to humans (Fletcher 2002).

Cryptosporidiosis in humans is caused by water contaminated with *Cryptosporidium parvum*. Wells et al. (2015) identified *C. parvum* in a small number of roe deer sampled above a Scottish Water public supply intake, with a prevalence of 33% (two of six deer). Comparatively, red deer prevalence was 80% for *Cryptosporidium* spp. (16 of 20 deer), of which 87.5% of the positive deer were infected with *C. parvum*. The authors concluded that the deer sampled in their study, alongside livestock also sampled, represented a significant risk to water quality and public health.

Deer have been linked to several enteric disease agents, including foodborne pathogens and diseases important to livestock health. An outbreak of *Escherichia coli* O157 in humans in Scotland in 2015 was linked to venison sourced from wild deer (Smith-Palmer et al. 2018). Shiga-toxin producing *E. coli* has been found in roe deer in the UK at a prevalence of 17.7% (stx1) and 41.2% (stx2), suggesting they may be acting as a reservoir of disease and pose a risk to public health (Pearce et al. 2023). The two species of *Campylobacter* most associated with gastrointestinal disease in humans, *C. jejuni* and *C. coli*, were also found in roe deer but at relatively low prevalences of 3% (Pearce et al. 2023). Several *Yersinia* spp. have also been detected in wild roe deer from across the UK at prevalences of 14.71% *Yersinia enterocolitica*) and 5.88% (*Y. pseudotuberculosis*), which may pose a risk to public health (Pearce et al. 2023).

C.1. Damage to conservation habitats

Impacts to conservation woodland are generated by all six species of deer to varying degrees, especially where deer densities are high (Quine, Shore, and Trout 2004). Browsing and grazing by deer can significantly alter the structure and development of natural forests and woodland even with a low proportion of primary production removal (White, Smart, et al. 2004). Light grazing can be beneficial for stimulating further productivity, but heavy grazing and browsing pressure can result in low species diversity, with only hardy and resistant species remaining. Sage et al. (2004) found roe deer browsing of understorey vegetation in small woodlands to cause significant changes to habitat structure and therefore species compositions. Woodlands providing a variable structure of understorey vegetation are more likely to provide suitable habitat for a greater number of species throughout a greater period of the year, such as small woodland mammals, butterfly and moth species, migrant woodland songbirds and game bird species (Sage et al. 2004). It is estimated that 8,000 hectares of woodland with Site of Special Scientific Interest status in the UK is currently in unfavourable condition or recovering due to the impacts of deer (Countryfile 2024). The cost of deer to conservation interests within the east of England was estimated at £265.8K annually and damage was found to be related to deer density in the area studied from an extrapolation of regression equations (White, Smart, et al. 2004); muntjac accounted for almost 60% of the estimated population, with roe and fallow deer accounting for considerably less (21% and 10%, respectively). No recent estimated costs were identified, therefore considering inflation (Bank of England 2024), current estimates would be around £493.4K.

C.2. Threat to biodiversity

In a review of the impact of deer on carbon sequestration in Scottish woodlands, Hirst (2021) reported that deer affect both above- and below-ground carbon stores. Browsing by deer results in less plant biomass available for photosynthesis; they change the plant diversity and amount of foliage available, which can alter the potential for carbon capture and density of carbon stored in standing biomass. Deer also have direct and indirect impacts on below ground carbon stores, by browsing and trampling plant species; this in turn can alter the composition of leaf litter and the subsequent decomposition of organic matter in the soil which stores carbon. Hirst (2021) concluded that there was growing evidence in Scotland that deer could hinder targets for woodland creation and harm the health of pre-existing woodland, thus reducing the ability of Scotland's woodlands to store carbon and off-set greenhouse gas emissions.

### Red deer - *Cervus elaphus*

Introduction

The red deer is native to the British Isles arriving approximately 11,000 years ago (Yalden 1998). During the 20th century, the ranges and numbers of red deer increased (Ward 2005), leading to impacts on both agriculture and biodiversity in sensitive areas (Putman and Moore 1998). In Britain, most red deer are found in Scotland; in England, they are relatively widespread with concentrations found in the south-west, Hampshire, East Anglia and the northwest and further scattered populations in between (Ward 2005). The red deer is highly adaptable and can be associated with a variety of habitats, although in Britain, the highest density of red deer are found in upland open moorland (Mathews et al. 2018). In the southwest of England, the highest densities are found in woodland and farmland margins especially where broadleaved woodland has actively been selected over upland heath and coniferous forest (Langbein 1997). The red deer is classified as an intermediate feeder between grazing and browsing with grasses forming the most important component of its diet; in winter, much of this grass is replaced by brambles, heather, holly and ivy as well as the shoots of both deciduous and coniferous trees (Putman 2008). Their distribution is patchy throughout its range, with records in England and Wales particularly scattered (Mathews et al. 2018). There are an estimated 346,000 red deer living wild in Britain: 79,700 in England, 256,000 in Scotland and 10,200 in Wales (Mathews et al. 2018).

A.1. Damage to forestry

Red deer browse young trees and strip bark from mature trees and can affect the ways in which trees grow, thus reducing their economic value as timber (White, Ward, et al. 2004). Bark damage can also result in fungal infection and subsequent branch or tree death, which causes further economic losses to foresters (Arnold et al. 2018). Stags may also thrash or fray trees in late summer when cleaning velvet (Putman 2008). Red deer can browse to heights of 180cm by rearing up onto their hind legs and will eat the bark of some trees, particularly rowans, willows, Norway spruce and lodgepole pine (Putman 2008).

Red deer along with roe deer will browse a variety of ground flora found in British woodlands (Kirby 2001). (See roe deer for estimated annual costs to forestry).

A.2. Damage to agricultural interests

Deer damage to cereals is principally caused by fallow, roe and red deer (Putman and Moore 1998) although in Scotland, red deer cause the most agricultural damage, along with fallow in some areas (Pepper, Barbour, and Glass 2019). Broader scale impacts are less significant in the southwest of England (Putman and Kjellander 2002; Rutter and Langbein 2005). However, many of the studies that have been conducted looking into the actual extent of damage caused by deer to agricultural crops have found little significant impact (Langbein and Rutter 2003; Wilson, Britton, and Symes 2009). The latter authors found that red and fallow deer grazing on grassland was likely to contribute to a 15% decrease in dry matter yield but that yield was also highly spatially variable so it is possible that not all yield loss was due to deer grazing. Annual use of cereal fields appears to vary so that damage may be high in certain years but not sustained over a long period. No apparent loss of winter wheat yield was found for red or fallow deer (Wilson, Britton, and Symes 2009). (See roe deer for estimated annual costs to agricultural interests).

A.3. Damage to vehicles (RTAs)

(See roe deer for information on damage to vehicles). Red deer, along with sika and Chinese water deer, cumulatively account for 3% of all deer-vehicle collisions in England each year (RSPCA 2022). As the cost of damage to vehicles alone (private and commercial) is estimated to be at least £17M (Deer Collisions 2023), red deer could be responsible for £510K shared between the three deer species (compared to £405K in 2013: Langbein (2007)).

A.4. Competition with livestock

Red deer are known to be a competitive species for hill sheep in Scotland (Clutton-Brock and Albon 1989), but can graze complimentarily with cattle (Gordon 1988).. No recent information identified.

B.1. Health risks from RTAs

(See roe deer for information on health risks from RTAs). Traffic collisions can represent an important cause of annual mortality for deer with losses to total populations being 1.5-2.6% for red deer (Langbein 2007). Of 1,646 deer carcasses collected in west and southwest England by Delahay et al. (2007), 14% were accessible to the public, many by way of being involved in vehicle collisions. Of these 3.4% cultured positive for *Mycobacterium bovis* (the causative agent of bovine tuberculosis) and 2.2% had visible tuberculous lesions (CSL 2006). In red deer specifically, prevalence may be between 0.1-3.5% and therefore the risk of infection is clearly low (Pepper, Barbour, and Glass 2019). In several reviews of zoonotic bovine TB in the UK, there have been no reports of a single human case arising from contact with deer (de la Rua-Domenech 2006; Davidson et al. 2017).

B.2. Reservoir of disease

(See roe deer for information on *Anaplasma phagocytophilum*). In north-west Scotland, *A. phagocytophilum* was the most prevalent tick-borne pathogen found in questing ticks; most strains (86%) were identified as zoonotic ecotype I, probably maintained by red deer, with the remaining 14% identified as non-zoonotic ecotype II, probably maintained by roe deer (Olsthoorn et al. 2021). *A. phagocytophilum* has also been reported in captive red deer (n=105) culled within geographically separate locations in Great Britain, with varying prevalence: 27.3% in Norfolk (3/11 deer), 1.5% in Dumfries & Galloway (1/68) and 3.7% in Lancashire (1/26: Johnson, Golding, and Phipps (2021)). These authors also reported the presence of other tick-borne pathogens, including *Babesia* spp. (prevalence: 18.2% in Norfolk (2/11), 2.9% in Dumfries & Galloway (2/68) and 3.8% in Lancashire (1/26)), most likely the deer-associated *Babesia odocoilei*. Gray et al. (2021) reported the first detection of *B. odocoilei* in the UK, which was found in 16% of red deer (n=24) culled near a farm in north-east Scotland. These authors also reported *B. divergens* in 11% of culled deer. The zoonotic potential of *B.*

*odocoilei* in the UK is unknown; Johnson, Golding, and Phipps (2021) reported that it caused sub-clinical infection in apparently healthy culled deer, therefore further monitoring is required to assess its potential to impact public health. *B. divergens* also has zoonotic potential, although it usually causes symptoms in immunosuppressed individuals and is primarily of concern to the livestock industry (Kjemtrup and Conrad 2000).

(See roe deer for information on Schmallenberg virus (SBV)). Barlow et al. (2013) confirmed that wild British red and fallow deer (from East Anglia) can be infected by SBV and thus could be a transient reservoir host for the virus. Southwell, Sherlock, and Baylis (2020) reported that, of 14 viable red deer samples submitted for testing (from a wide coverage of Great Britain), only 1 (7.1%) was positive to detection of SBV antibodies (using virus neutralisation (VNT) testing, which is considered a more accurate test than ELISA). Graham et al. (2017) found that 2 of 17 (11.8%) red deer were positive to detection of SBV antibodies. Southwell, Sherlock, and Baylis (2020) concluded that the overall prevalence of SBV in wild British deer was 13.8% (using VNT) to 22.1% (ELISA), suggesting that as many as 440,000 deer (of an approximately 2 million UK wild deer population) may have been exposed to SBV.

Gastrointestinal (GI) nematodes are among the most important causes of production loss in farmed ruminants (Chintoan-Uta et al. 2014). Red deer are known to harbour abomasal nematodes but Chintoan-Uta et al. (2014) found the burden to be less in red deer compared to roe deer (with the lowest burden found in fallow deer) in areas of south-west England. Unlike roe deer though, there was no evidence of anthelmintic-resistant GI nematodes in red deer as no *H. contortus* were recovered from the study samples. In Spain, significant cross-transmission of GI nematodes between red deer and cattle has been reported (Santin-Duran et al. 2004), therefore the transmission risk between deer and livestock in the UK warrants further investigation. Red deer are also common wild definitive hosts for liver fluke (*Fasciola hepatica*), which have detrimental effects on the health and productivity of livestock (Charlier et al. 2014). *M. avium paratuberculosis* (MAP), which is responsible for Johne's disease, has been detected at low levels in red deer with Pearce et al. (2023) reporting a prevalence of 3.7% (although most samples submitted for testing were from abattoirs, deer farms or parks).

B.3. bTB transmission

Delahay et al. (2007) reported the prevalence of bovine tuberculosis *M. bovis* in red deer in South-West England as 1.02%, therefore the risk of transmission is low. The risk of exposure of cattle to bTB by red deer has been estimated to be approximately 16% of that posed by badgers, the putative main wildlife reservoir of *M. bovis* in England and Wales (Ward et al. 2009). However, Collard (2023) reported a high prevalence of bTB positive red deer sampled from a large herd on Exmoor (28.3%), noting a significant correlation between the presence of deer with bTB and the number of farms reporting bTB positive cattle. Although the main bTB host system in the UK is currently cattle-badger-fallow deer, there is also a potential risk posed by a cattle-roe deer-red deer system (Hardstaff et al. 2014), which may be more prominent in localised areas.

B.4. Reservoir of zoonotic disease

Tick-borne viruses, such as louping ill, have been detected both in wild and farmed examples of red deer but do not appear to cause clinical signs of illness (Reid, Barlow, and Pow 1978; Holding et al. 2020). Red deer are also a reservoir for the ticks of Lyme disease but have not been directly linked to involvement in transmission of the disease to humans (Fletcher 2002). Several studies have recorded the presence of tick-borne encephalitis virus (TBEV) in UK deer. Johnson, Golding, and Phipps (2021) found TBEV in captive red deer in three locations across Great Britain (prevalence: 0% in Norfolk, 26.7% in Dumfries and Galloway (4/15) and 5.9% in Lancashire (1/17), concluding that the seropositive results for TBEV could provide further evidence of TBEV circulation in UK deer, although further surveillance was required. Holding et al. (2020) also found antibodies to tick-borne encephalitis virus (TBEV) in approximately 4% of deer culled across England and Scotland; of the 1,323 deer sampled, 242 were red deer of which 9 (3.7%) were ELISA-positive. The detection of TBEV is important as it can infect humans, although the prevalence in deer appears to be quite low and generally localised;

for example, in the Norfolk/Suffolk (Thetford Forest) focal area, serological evidence has shown a high prevalence of TBEV exposure (47.7% in the deer species found there, red, roe, fallow, muntjac) (Holding et al. 2020).

Cryptosporidiosis in humans is caused by water contaminated with *Cryptosporidium parvum*. Wells et al. (2015) identified *Cryptosporidium* spp. in a large proportion of red deer sampled above a Scottish Water public supply intake, with a prevalence of 80% (16 of 20 deer); of these, 87.5% were infected with *C. parvum*. The authors concluded that the deer sampled in their study, alongside livestock also sampled, represented a significant risk to water quality and public health. Deer have been linked to several enteric disease agents, including foodborne pathogens and diseases important to livestock health. An outbreak of *Escherichia coli* O157 in humans in Scotland in 2015 was linked to venison sourced from wild deer (Smith-Palmer et al. 2018). These authors found that although shiga-toxin producing *E. coli* was found in red and sika deer at very low prevalence (0.28%), STEC O157 isolates were shed at high levels from positive deer with potential to cause severe disease in humans. Pearce et al. (2023) reported a higher prevalence in red deer sampled from across the UK (from abattoir, farmed, park and wild deer) of 18.0% (stx1) and 17.8% (stx2), suggesting they may be acting as a reservoir of disease and pose a risk to public health. The two species of *Campylobacter* most associated with gastrointestinal disease in humans, *C. jejuni* and *C. coli*, were also found in red deer but at relatively low prevalences of 1.5% and 3.5% (Pearce et al. 2023). Several *Yersinia* spp. have also been detected in wild red deer from across the UK at prevalences of 4.5% (*Yersinia enterocolitica*) and 1.5% (*Y. pseudotuberculosis*), which may pose a low risk to public health (Pearce et al. 2023).

<u>C.1. Damage to conservation habitats</u>

(See roe deer for information on damage to conservation habitats). The West Country holds the largest population of red deer found throughout England;  some of the damage caused by the increasing deer numbers includes the possible prevention of natural tree regeneration in semi-natural woodland and damage to other sensitive vegetation in conservation areas (Langbein 1997). . Red deer appear to be attracted to areas where there are larger proportions of transitional heathland which have significant cover of European gorse and bracken, and broad-leaved woodland (Langbein 1997). Tree saplings above a height of 35cm, which protrude above the lower field layer, seem to be especially targeted with large seedling losses (both established and depletion of seed source due to uprooting) occurring during winter and spring (Langbein 1997). Other factors may also influence regeneration, such as the abundance of other vegetation, light availability, cover and soil fertility (Langbein 1997), so that control of deer populations may not give the desired results but could be expensive to pursue. Old oak coppice regeneration is unlikely to occur in areas with deer densities above 5/km$^2$ and might still be suppressed at lower densities (Langbein 1997). Red deer grazing on heather moorland is not considered to be a significant problem where high densities of sheep also graze the area but may be an issue in areas of localised high deer density (Langbein 1997). Problems are likely to occur in areas that are simultaneously stocked with high densities of sheep without taking deer densities into consideration and in areas where the conservation aim is to improve or increase heathland cover (Langbein 1997).

It is estimated that 8,000 hectares of woodland with Site of Special Scientific Interest status in the UK is currently in unfavourable condition or recovering due to the impacts of deer (Countryfile 2024). The cost of deer to conservation interests within the east of England were estimated at £265.8K annually and damage was found to be related to deer density in the area studied from an extrapolation of regression equations (White, Smart, et al. 2004). No recent estimated costs were identified, therefore considering inflation (Bank of England 2024), current estimates would be around £493.4K. The relatively high density of red deer in these areas means that they are likely to be a significant contributor to any costs.

<u>C.2. Threat to biodiversity</u>

(See roe deer for information on threat to biodiversity). Red deer may also have positive effects on biodiversity; Murray et al. (2016) found that Whinchats, a declining grassland bird, preferred to forage in areas that were grazed by red deer and sheep in upland areas of Scotland.

Sika deer - *Cervus nippon*
Introduction

. Since being introduced to the British Isles approximately 150 years ago, Japanese sika deer have expanded their distributional range and established many large free-ranging populations in Scotland and England (Putman and Pemberton 2022). Sika are primarily associated with acidic soils, with many populations being established in coniferous plantation or adjacent heathland areas and a general preference for wooded areas of varying structure (Putman 2008; Mathews et al. 2018). Sika residing in these conditions tend to consume high volumes of grasses and heather with the proportions varying according to seasons and availability (Mann and Putman 1989). The distribution of sika deer is patchy throughout the British Isles with an estimated 103,000 sika deer living wild in Britain: 45,300 in England, 54,000 in Scotland and 3,600 in Wales (Mathews et al. 2018).

A.1. Damage to forestry

Sika are considered to cause considerable damage to forestry interests by stripping the bark from mature trees (Putman and Moore 1998). Bucks may also thrash and fray trees in late summer (Putman 2008). Deer browse young trees and strip bark from mature trees and can affect the ways in which trees grow, thus reducing their economic value as timber (White, Ward, et al. 2004). Bark damage can also result in fungal infection and subsequent branch or tree death, which causes further economic losses to foresters (Arnold et al. 2018). In the New Forest, considerable quantities of both deciduous and coniferous browse can be consumed by sika, particularly during the winter months (Putman 2008). In addition, sika deer may also cause 'bole-scoring' damage to mature trees, particularly coniferous trees; this scoring of tree trunks with their antler points causes deeper, more serious damage to trees than bark stripping and can result in reduced economic value or death of the tree, although damage is likely to be localised (Putman and Moore 1998).

Sika tend to prefer smooth-barked trees in comparison to roe and red deer, such as rowan and yew, which may lead to more dominant trees in a stand being targeted by sika (Gill 1992). Based on the distribution of fallow and sika and the distribution of new plantations and assuming that 25% of plantations in Scotland suffer damage, 40% in England and 5% in Wales, Williams et al. (2010) estimated the cost of damage due to bark stripping by deer at £1.2M (Scotland £736.9K, England £477K and Wales £25.8K). The total cost to forestry by all non-native deer was estimated at £17.34M. The revised recent estimate of the total cost to forestry by all non-native deer is £26.5M (Eschen et al. 2023). The total estimated management costs for deer in Scotland in 2018/19 (roe, red, sika and fallow) was £6.96M (Pepper, Barbour, and Glass 2019).

A.2. Damage to agricultural interests

An estimate of the total annual cost of deer damage to agriculture (mostly cereal damage) in the east of England has been calculated as £1.92-4.57M (White, Smart, et al. 2004). Williams et al. (2010) estimated the total cost of non-native deer to agriculture (including culling costs) in the UK was £7.26M; this had increased significantly by 2023, with an estimated cost of £21M, presumably due to the 50% increase in the number of non-native deer reported between the two periods (Eschen et al. 2023). With two thirds of the damage caused by fallow and roe deer, followed by red deer (White, Smart, et al. 2004), sika are clearly not a primary issue for agricultural damage. As with all deer species though, damage is likely to be greatest where they are found at highest density.

A.3. Damage to vehicles (RTAs)

(See roe deer for information on damage to vehicles (RTAs)). Red deer, sika and Chinese water deer, cumulatively account for 3% of all deer-vehicle collisions in England each year (RSPCA 2022). As the cost of damage to vehicles alone (private and commercial) is estimated to be at least £17M (Deer Collisions 2023), sika deer could be responsible for £510K shared between the three deer species (compared to £405K in 2013: deer.collisions.co.uk). Eschen et al. (2023) produced estimates of £14.4M annually for all non-native species of deer.

A.4. Competition with livestock

Sika deer have the potential for high levels of competition with other ungulates, particularly the native red and roe deer species but there is little evidence that sika have any competitive effects on their populations (Putman 2008).. In the New Forest, analyses have shown there to be habitat use overlaps with both cattle and horses (Putman 2008). No recent information identified.

B.1. Health risks from RTAs

(See roe deer for information on health risks from RTAs). Traffic collisions can represent an important cause of annual mortality for deer with losses to total populations of 3.4-6% for sika deer (Langbein 2007). Of 1,646 deer carcasses collected in west and southwest England (Delahay et al. 2007), 14% were accessible by the public, many by way of being involved in vehicle collisions. Of these 3.4% cultured positive for *Mycobacterium bovis* (the causative agent of bovine tuberculosis) and 2.2% had visible tuberculous lesions (CSL 2006). In southern England and Wales, sika deer had a prevalence of 2.1% (Delahay et al. 2002) therefore the risk of infection is low. In several reviews of zoonotic bovine TB in the UK, there have been no reports of a single human case arising from contact with deer (de la Rua-Domenech 2006; Davidson et al. 2017).

B.2. Reservoir of disease

(See roe deer for information on Schmallenberg virus (SBV)). Little information was identified for SBV infection in sika deer in Britain; in a cross-sectional study of British wild deer, Southwell, Sherlock, and Baylis (2020) sampled only one viable sika sample submitted for testing, which was negative for SBV antibodies. In Ireland however, Graham et al. (2017) found that 25 of 213 (11.7%) sika deer were positive to detection of SBV antibodies, which suggests this species could pose a potential transmission risk to livestock.

B.3. bTB transmission

Of 240 sika deer killed in England in 1971-1996, five were found to be carrying bovine TB, showing that infection is possible but the rate of infection is low (Delahay et al. 2007). The transmission of bTB to cattle is also unlikely due to the limited distribution of sika in England. In Ireland, higher levels of bTB in cattle have been associated with higher local densities of sika deer, suggesting they may be acting as maintenance hosts (Kelly, Mullen, and Good 2021).

B.4. Reservoir of zoonotic disease

Antibodies to tick-borne encephalitis virus (TBEV) were found in approximately 4% of culled deer across England and Scotland; of the 1,323 deer sampled, 48 were sika deer of which 1 (2.1%) was ELISA-positive (Holding et al. 2020). The detection of TBEV is important as it can infect humans, although the prevalence in deer appears to be quite low and generally localised.

An outbreak of *E. coli* O157 in humans in Scotland in 2015 was linked to venison sourced from wild deer (Smith-Palmer et al. 2018). Although shiga-toxin producing *E. coli* was found in red and sika deer at very low prevalence (0.28%), STEC O157 isolates were shed at high levels from positive deer with potential to cause severe disease in humans. Pearce et al. (2023) reported a higher prevalence in 12 wild sika deer of 25.0% (stx1) and 8.3% (stx2), suggesting they may be acting as a reservoir of disease and pose a risk to public health (Pearce et al. 2023). Several *Yersinia* spp. have also been detected in wild sika deer from across the UK at prevalences of 25% (*Yersinia enterocolitica*) and 8.3% (*Y. pseudotuberculosis*), which may pose a risk to public health (Pearce et al. 2023).

C.1. Damage to conservation habitats

(See roe deer for information on damage to conservation habitats). So far, there has been little significant level damage reported as being caused by sika in England but increasing localised populations could become highly detrimental to areas of conservation importance if their densities increase (Putman 2008).

C.2. Threat to biodiversity

(See roe deer for information on threat to biodiversity).

C.3. Threat to genetic integrity

Native red deer and non-native sika can hybridise in areas where distribution overlap occurs, producing fertile young that threatens the genetic integrity of the native red deer (Abernethy 1994; Smith et al. 2018). Little hybridisation is thought to occur in wild populations when densities of each species are high, but once the first cross has occurred, possibly beginning in captive populations, further hybridisation takes place at a rapid pace (Lowe and Gardiner 2009). It is thought that the majority of sika populations currently found in Britain are in fact, already hybridised populations (Putman and Hunt 1994). Due to the difficulties in identifying hybrids, especially in parks where both species have been kept together, there are now restrictions on introducing either species to the wild under Schedule 9 of the Wildlife and Countryside Act, 1981, to preserve the red deer genotype (Putman 2008). Senn and Pemberton (2008) also found that 43% of deer in a population at West Loch Awe in Scotland were hybrids. This is a much higher level than previously thought as other areas showed generally low hybridization due to the rarity of hybridisation events and the limited time the two species have been in close contact with each other. The effects of habitat variability bringing the two species closer together has resulted in increased levels of hybridisation in West Loch Awe and has the potential to cause much greater levels of hybridisation across Scotland in the future, which could be a very significant problem for the native red deer (Senn and Pemberton 2008).

Fallow deer - *Dama dama*

Introduction

Fallow deer became extinct in Britain prior to the last glaciation and are thus regarded as non-native; they were re-introduced to Britain approximately 1,000-2,000 years ago, most likely from mainland Europe (Putman 2003) and are considered naturalised. The fallow deer is characteristically found in mature woodland, which is primarily used for shelter (Mathews et al. 2018). They prefer deciduous or mixed woodland with a good level of understorey growth but will also be found in coniferous plantations with some open areas (Putman 2008). Fallow deer are preferential grazers feeding on ground vegetation along woodland rides; they will often forage in adjacent open land, such as agricultural land, and are not wholly dependent on woodland food supplies (Thirgood 1995). Acorns, chestnuts, beech mast and fruits, such as brambles, are eaten in autumn-early winter (Putman 2008)Their distribution is patchy throughout its range, with isolated records in Scotland but is particularly concentrated in the south and east of England (Mathews et al. 2018). As with the other deer species, their numbers and ranges have expanded (Ward 2005) leading to increased impacts to a range of human interests (Putman and Moore 1998). There are an estimated 264,000 fallow deer living wild in Britain: 188,000 in England, 56,700 in Scotland and 19,000 in Wales (Mathews et al. 2018).

A.1. Damage to forestry

Fallow deer are considered to cause considerable damage to forestry interests by stripping the bark from mature trees, although to a lesser extent than sika; they will bark strip both coniferous and deciduous trees, with incidence increasing during hard winters (Putman and Moore 1998). The browsing of young trees and stripping of bark from mature trees can affect the ways in which trees grow, thus reducing their economic value as timber (White, Ward, et al. 2004). Bark damage can also result in fungal infection and subsequent branch or tree death, which causes further economic losses to foresters (Arnold et al. 2018). Bucks may also thrash trees in late summer and can cause considerable damage to trees (Putman 2008). Fallow deer are also prone to damaging young plantations and can prevent the regeneration of coppice at high densities (Gill 1992). (See sika deer for information relating to costs of damage to forestry).

A.2. Damage to agricultural interests

Fallow deer are regularly observed grazing in lowland winter and spring-sown cereals, and will return to cereal crops to feed directly on the ripening ears (Putman and Moore 1998). They are the species most associated with damage to arable agriculture, notably cereals and seed crop grasses (Putman, Culpin, and Thirgood 1993; Putman and Moore 1998). A questionnaire survey of 1,192 respondents conducted for the British Deer Society in 1995 found that 69% of farmers had deer on their holdings, of which roe and fallow deer were the most frequent (Doney and Packer 2002). Of the 69% reporting deer, 59% reported damage to cereals and crops but only 54 respondents claimed the annual damage cost exceeded £500 and overall, the study found little difference between yield in damaged and undamaged crops (Wilson, Britton, and Symes 2009). Despite these concerns, many of the studies that have been conducted looking into the actual extent of damage caused by deer to agricultural crops have found deer to have little significant effect (Wilson, Britton, and Symes 2009). The study by Wilson, Britton, and Symes (2009) found that red and fallow deer grazing on grassland were likely to contribute to a 15% decrease in dry matter yield but that yield was also highly spatially variable so it is possible that not all yield loss was due to deer grazing. Annual use of cereal fields appears to vary so that damage may be particularly high in certain years but not sustained over a long period. No apparent loss of winter wheat yield was found for red or fallow deer (Wilson, Britton, and Symes 2009).

(See sika deer for estimated annual costs to agricultural interests caused by non-native deer). As an estimated two thirds of damage caused by deer is attributable to fallow and roe deer (White, Smart, et al. 2004), fallow may be responsible for a large proportion of the estimated £21M (Eschen et al. 2023) damage caused by non-native deer in the UK.

A.3. Damage to vehicles (RTAs)

(See roe deer for information on damage to vehicles (RTAs)). Fallow deer are the most commonly involved species in deer-vehicle collisions in England each year, accounting for 40% of all collisions (Deer Collisions 2023; RSPCA 2022). Damage to vehicles alone (private and commercial) is estimated to be at least £17M (Deer Collisions 2023), which means that fallow deer could be responsible for £6.8M worth of material damage to vehicles in England (compared to £5.4M in 2013: Deer Collisions, 2023). Eschen et al. (2023) estimated that non-native deer caused £14.4M damage annually in the UK.

B.1. Health risks from RTAs

(See roe deer for information on health risks from RTAs). Traffic collisions can represent an important cause of annual mortality for deer with losses to total populations being 7.6-13.2% for fallow deer (Langbein 2007). Of 1,646 deer carcasses collected in west and southwest England (Delahay et al. 2007), 14% were accessible by the public, many by way of being involved in vehicle collisions. Of these 3.4% cultured positive for *Mycobacterium bovis* (the causative agent of bovine tuberculosis) and 2.2% had visible tuberculous lesions (CSL 2006). In fallow deer specifically, prevalence may be around 4.37% (Delahay et al. 2007), which is higher than some other deer species, but the overall risk of infection to public health is likely low. In several reviews of zoonotic bovine TB in the UK, there have been no reports of a single human case arising from contact with deer (de la Rua-Domenech 2006; Davidson et al. 2017).

B.2. Reservoir of disease

(See roe deer for information on Schmallenberg virus (SBV) and *Anaplasma phagocytophilum*). Southwell, Sherlock, and Baylis (2020) reported that, of five viable fallow deer samples submitted for testing (from a wide coverage of Great Britain), two (40%) were positive to detection of SBV antibodies (using virus neutralisation (VNT) testing, which is considered a more accurate test than ELISA). Although prevalence in this study was high, sample size was small and further testing would be required to determine how prevalent SBV is in the UK fallow population. In Ireland, Graham et al. (2017) found that 11 of 138 (8%) fallow deer were positive to detection of SBV antibodies, which suggests fallow deer could pose a potential transmission risk to livestock. Southwell, Sherlock, and Baylis (2020) concluded that the overall prevalence of SBV in wild British deer was 13.8% (using VNT) to 22.1% (ELISA), suggesting that as many as 440,000 deer (of an approximately 2 million UK wild deer population) may have been exposed to SBV.

Robinson, Shaw, and Morgan (2009) reported *A. phagocytophilum* infection in fallow deer, although prevalence was much lower (21%) compared to red deer (80%) and sika deer (50%). Fallow deer also show very little clinical sign of foot and mouth disease but they can become carriers of the disease which is problematic due to their widespread distribution (Putman 2008). *M. avium* paratuberculosis (MAP), which is responsible for Johne's disease, has been detected at low levels in fallow deer with Pearce et al. (2023) reporting a prevalence of 2.4% (although most samples submitted for testing were from parks, with a smaller number of wild deer samples).

<u>B.3. bTB transmission</u>

The bTB host system in the UK and Ireland is currently cattle-badger-fallow deer (Hardstaff et al. 2014) although they likely represent a localised source of infection to cattle predominantly in parkland situations where deer and cattle grazing areas are in close proximity (Delahay et al. 2007). (Delahay et al. 2007) reported the prevalence of *M. bovis* in fallow deer in South-West England as approximately 4.37%, which is higher than roe, red or sika deer. Ward et al. (2009) estimated the risk of exposure of cattle to bTB by fallow deer to be approximately 27% of that posed by badgers, the putative main wildlife reservoir of *M. bovis* in England and Wales.

<u>B.4. Reservoir of zoonotic disease</u>

(See roe deer for information on TBEV). Antibodies to tick-borne encephalitis virus (TBEV) were found in approximately 4% of culled deer across England and Scotland; of the 1,323 deer sampled, 246 were fallow deer of which 10 (4.1%) were ELISA-positive (Holding et al. 2020). The detection of TBEV is important as it can infect humans, although the prevalence in deer appears to be quite low and generally localised; for example, in the Norfolk/Suffolk (Thetford Forest) focal area, serological evidence has shown a high prevalence of TBEV exposure (47.7% in the deer species found there; red, roe, fallow, muntjac) (Holding et al. 2020).

Deer have been linked to several enteric disease agents, including foodborne pathogens and diseases important to livestock health. An outbreak of *Escherichia coli* O157 in humans in Scotland in 2015 was linked to venison sourced from wild deer (Smith-Palmer et al. 2018). Shiga-toxin producing *E. coli* has been found in fallow deer from across the UK at prevalences of 6.75% (stx1) and 14.72% (stx2), suggesting they may be acting as a reservoir of disease and pose a risk to public health, although prevalence is less than roe and red deer (Pearce et al. 2023). The two species of *Campylobacter* most associated with gastrointestinal disease in humans, *C. jejuni* and *C. coli*, were also found in fallow deer but at relatively low prevalences of 0.6% and 3.7% respectively (Pearce et al. 2023). Several *Yersinia* spp. have also been detected in fallow deer at prevalences of 18.4% (*Yersinia enterocolitica*) and 4.3% (*Y. pseudotuberculosis*), which may pose a risk to public health (Pearce et al. 2023).

<u>C.1. Damage to conservation habitats</u>

(See roe deer for information on damage to conservation habitats). Browsing by fallow deer has been shown to suppress growth and subsequent recruitment to the canopy in national nature reserves such as Castor Hanglands, Peterborough, with Ward, Clarke, and Cooke (1994) reporting that up to 40% of woody plants showed signs of browsing damage. Problems for heathland likely arise where the conservation aim is to improve or increase heather cover (Langbein 1997). Heathland browsing by deer is often welcomed where there is limited livestock grazing as it helps to prevent invasion of heathland with unwanted scrub (Putman and Moore 1998).

<u>C.2. Threat to biodiversity</u>

(See roe deer for information on threat to biodiversity).

Chinese water deer - *Hydropotes inermis*

Introduction

The Chinese water deer is native to eastern China and Korea but has become established in the UK having been introduced into parks during the 19th century (Cooke 2019). Escapes and releases from these parks have led to the establishment of a free-living population with the first reports of Chinese water deer being found in the wild in Bedfordshire during the 1950s (Whitehead 1993). As with the other deer species, their numbers and ranges have expanded (Ward 2005) leading to increased impacts on agriculture and biodiversity (Putman and Moore 1998). They are found near reed beds but also in woodland with mixed vegetation for feeding and for cover and occasionally in agricultural habitats (Putman 2008; Mathews et al. 2018). They are selective feeders, taking grasses, sedges, herbs and woody species, crops (Putman 2008). Chinese water deer have a very restricted geographical range, largely found in Cambridgeshire and Norfolk (Mathews et al. 2018) but is also found in Bedfordshire and Suffolk in a discontinuous distribution (Putman 2008). They have the lowest number of free-living individuals compared to the other five deer species resident in the UK with an estimated population in England in 2005 of around 1,500, which had risen to 3,600 by 2016 (Mathews et al. 2018).

A.1. Damage to forestry

Although all deer can browse young trees and strip bark from mature trees, Chinese water deer are not considered to cause significant impacts on forestry (Putman 2008; Williams et al. 2010). In a review of the economic cost of invasive non-native species in GB, Chinese water deer were not included due to their limited use of woodlands (Williams et al. 2010). In a revised estimate of the economic costs of non-native deer in GB, Chinese water deer were included in the overall cost of £26.5M (Eschen et al. 2023) although they probably contribute very little in terms of forestry damage.

A.2. Damage to agricultural interests

(See sika deer for information on damage to agricultural interests). The Chinese water deer feed on crops, such as the green tops of carrots, winter wheat and potatoes left over post-harvest but it mostly eats weeds when on arable land (Putman 2008). Most damage tends to be done when other food sources are scarce and root crops and sprouting grain are eaten, although it is not thought that this amounts to anything significant and remains localised (Cooke and Farrell 1987). They are believed to have a negligible impact on agriculture in the east of England (White, Smart, et al. 2004). Williams et al. (2010) estimated the total cost of non-native deer to agriculture (including culling costs) in the UK was £7.26M; this had increased significantly by 2023, with an estimated cost of £21M (Eschen et al. 2023), presumably due to the 50% increase in the number of non-native deer reported between the two periods. With two thirds of the damage caused by fallow and roe deer, followed by red deer, Chinese water deer are clearly not a primary issue for agricultural damage. As with all deer species though, damage is likely to be greatest where they are found at highest density.

A.3. Damage to vehicles (RTAs)

(See roe deer for information on damage to vehicles (RTAs)). Chinese water deer, along with red and sika deer, cumulatively account for 3% of all deer-vehicle collisions in England each year (RSPCA 2022). As the cost of damage to vehicles alone (private and commercial) is estimated to be at least £17M (Deer Collisions 2023), Chinese water deer could be responsible for £510K shared between the three deer species (compared to £405K in 2013). Eschen et al. (2023) produced estimates of £14.4M annually for all non-native species of deer. DVCs with Chinese water deer are likely to be localised due to their limited geographical distribution; they are also considerably smaller than other deer species therefore less likely to cause as much damage (Langbein 2007).

B.1. Health risks from RTAs

(See roe deer for information on health risks from RTAs). Traffic collisions can represent an important cause of annual mortality for deer with losses to total populations being 7.3-12.6% for Chinese water deer (Langbein 2007). There is little information relating to the prevalence of bTB in Chinese water deer but given their restricted distribution in mainly southeast England, this species is likely to be of less concern in terms of public health risk. In several reviews of zoonotic bovine TB in the UK, there have been no reports of a single human case arising from contact with deer (de la Rua-Domenech 2006; Davidson et al. 2017).

B.2. Reservoir of disease

Little is known about diseases of Chinese water deer in England. Given their small geographical distribution and small population size, they are less likely to come into contact with ruminant livestock compared with other deer species, such as roe, red and fallow (Hartley et al. 2012).

B.4. Reservoir of zoonotic disease

In a survey of enteric disease agents in UK deer, Pearce et al. (2023) reported several pathogens in Chinese water deer, although the sample size submitted for testing was very small (n=2). Shiga-toxin producing *E. coli* was found in one of the samples (prevalence 50% (stx2)) along with *Yersinia entercolitica* (prevalence 50%). Both disease agents pose a risk to public health and due to their presence in one of the two deer, further investigation into the disease carried by Chinese water deer is warranted.

C.1. Damage to conservation habitats

Chinese water deer do not cause significant impacts to conservation habitats like the muntjac because they do not fray trees, exist in relatively low densities, are a less extreme concentrate feeder and are generally found in more robust habitats (Putman 2008).

Muntjac - *Muntiacus reevesi*
Introduction

The Reeves' muntjac was introduced to parks in the 19th century (Cooke 2019). It is native to southeast China and Taiwan where it inhabits areas of scrub and dense forest (Putman 2008). Muntjac tend to prefer dense habitat with a variety of vegetation that is provided by deciduous woodland or coppiced woodland, undisturbed gardens and cemeteries and commercial coniferous woodlands with some deciduous trees, although some reports have been made of young muntjac being found in urban areas of England (Putman 2008). They feed on leaves, flowers and shoots of many species, and brambles where available (Putman 2008). Escapes and releases from the parks in the early and mid-20th century have resulted in the muntjac being the most abundant of the non-native free-living deer species (Whitehead 1993). They are concentrated in the south of England, but have spread into Yorkshire, with further isolated populations in county Durham (Mathews et al. 2018). There are an estimated 128,000 muntjac in Britain: 112,000 in England, 16,300 in Wales. There is no known wild population in Scotland (Mathews et al. 2018).

A.1. Damage to forestry

Deer browse young trees and can affect the ways in which trees grow, thus reducing their economic value as timber (White, Ward, et al. 2004). However, muntjac cause very little damage to conifers and do not tend to strip bark from mature trees like other deer species but may take the shoots and flowers from deciduous trees (Putman 2008). Fraying of bark on saplings is not easily distinguished from roe deer damage but occurs at low level (Putman 2008).

The cost of damage due to bark stripping by non-native deer was estimated at £1.2M (Scotland £736.8K, England £477K and Wales £25.9K), although muntjac are likely to contribute very little to this figure

(Williams et al. 2010) The total cost to forestry by all non-native deer was estimated at £17.34M. The revised recent estimate for non-native deer is £26.5M (Eschen et al. 2023).

A.2. Damage to agricultural interests

Due to its small mouth, muntjac deer do not directly cause damage to carrots, potatoes or sugar beets but may take pieces after harvest (Putman 2008). Muntjac may also take crops such as oilseed rape, field beans or more valuable market garden crops (Cooke and Farrell 2001). Leaves and shoots from a variety of deciduous trees and garden plants may also be taken, especially as exclusions from gardens can be very difficult (Putman 2008). Due to their small size and their tendency to stay within 50m of woodland, muntjac are unlikely to cause heavy damage to tall cereal crops once they have ripened, which is when the highest proportion of damage occurs (White, Smart, et al. 2004).

Williams et al. (2010) estimated the total cost of non-native deer to agriculture (including culling costs) in the UK was £7.26M; this had increased significantly by 2023, with an estimated cost of £21M (Eschen et al. 2023), presumably due to the 50% increase in the number of non-native deer reported between the two periods. With two thirds of the damage caused by fallow and roe deer, followed by red deer (White, Smart, et al. 2004), muntjac are clearly not a primary issue for agricultural damage.

A.3. Damage to vehicles (RTAs)

(See roe deer for information on damage to vehicles (RTAs)). Muntjac deer are the third most commonly involved species in deer-vehicle collisions in England each year, accounting for 25% of all collisions (Deer Collisions 2023; RSPCA 2022). Damage to vehicles alone (private and commercial) is estimated to be at least £17M (Deer Collisions 2023), which means that muntjac deer could be responsible for £4.3M worth of material damage to vehicles in England (compared to £3.4M in 2013: deer.collisions.co.uk). Eschen et al. (2023) estimated that non-native deer caused £14.4M damage annually in the UK.

B.1. Health risks from RTAs

(See roe deer for information on health risks from RTAs). Traffic collisions can represent an important cause of annual mortality for deer with losses to total populations being 7.8-13.4% for muntjac deer (Langbein 2007). Of 1,646 deer carcasses collected in west and southwest England (Delahay et al. 2007), 14% were accessible to the public, many by way of being involved in vehicle collisions. Of these 3.4% cultured positive for *Mycobacterium bovis* (the causative agent of bovine tuberculosis) and 2.2% had visible tuberculous lesions (CSL 2006). In muntjac deer specifically, prevalence was around 5.17% (Delahay et al. 2007), which was the highest observed prevalence of all the deer species (although confidence limits were wide due to small sample size). Although this is the highest prevalence of all the deer species, the overall risk of infection to public health is likely low. In several reviews of zoonotic bovine TB in the UK, there have been no reports of a single human case arising from contact with deer (de la Rua-Domenech 2006; Davidson et al. 2017).

B.2. Reservoir of disease

There have been some cases of disease reported in muntjac, including bTB, meningitis, wasting disease and some joint diseases but these have been very limited and are unlikely to be widespread within the British muntjac population (Putman 2008). From experimental work, muntjac can be infected with foot-and-mouth disease and can also potentially transmit the virus intraspecifically and to cattle and sheep, although the risk is considered negligible (Hartley et al. 2012). In addition, no cases have been reported in the wild (Putman 2008; FAWC 2013). Duscher et al. (2020) reported the presence of *A. phagocytophilum* in roe deer and muntjac in Thetford Forest, although at a low prevalence in both species (6.6% roe deer, 1% muntjac).

(See roe deer for information on Schmallenberg virus (SBV)). SBV was reported as being present in muntjac in 2019 (Southwell, Sherlock, and Baylis 2020). These authors found that of 8 viable muntjac deer samples submitted for testing (from a wide coverage of Great Britain), one (12.5%) was positive to detection of SBV antibodies (using virus neutralisation (VNT) testing, which is considered a more accurate test than ELISA). They concluded that the overall prevalence of SBV in wild British deer was 13.8% (using VNT) to 22.1% (ELISA), suggesting that as many as 440,000 deer (of an approximately 2 million UK wild deer population) may have been exposed to SBV. Seropositivity rates are likely to vary depending on latitude (e.g. SBV rates are expected to be lower in Scotland where more deer are present), which concurred with findings that deer in the south were more likely to be SBV-positive (Southwell, Sherlock, and Baylis 2020) where muntjac deer are mostly found. The latter study highlighted the significantly extended area over which SBV has been reported since 2012, raising concerns for the welfare of wild and farmed deer, and the transmission risk to livestock.

In Northern Ireland, a novel gammaherpesvirus (genus *Rhadinovirus*) considered a threat to livestock was detected in wild muntjac deer, although only two animals were submitted for testing and the virus was present at very low levels causing no obvious pathological symptoms in deer (McKillen et al. 2017). These authors concluded that the pathogenicity of the virus to muntjac was unknown; if muntjac become established in Northern Ireland, they could potentially interact with livestock posing a risk of disease transmission and should be monitored further. The prevalence of *Mycobacterium avium paratuberculosis* (MAP), which is responsible for Johne's disease, was 8.3% in wild muntjac (n=24) submitted for testing in the UK (Pearce et al. 2023). Prevalence for MAP was higher in muntjac than any of the other deer species sampled.

B.3. bTB transmission

Delahay et al. (2007) reported the prevalence of *M. bovis* in muntjac in cattle bTB hotspots in South-West England as 5.17%. Ward et al. (2009) estimated the risk of exposure of cattle to bTB by muntjac to be approximately 8% of that posed by badgers, the putative main wildlife reservoir of *M. bovis* in England and Wales. This was the lowest risk of the four deer species considered in this study, but the estimate was highly uncertain. Ward and Smith (2011) also suggested that muntjac deer were likely to act as spillover hosts at densities below 6/km$^2$ and maintenance hosts at densities above 56/km$^2$.

B.4. Reservoir of zoonotic disease

Antibodies to tick-borne encephalitis virus (TBEV) were found in approximately 4% of culled deer across England and Scotland; of the 1,323 deer sampled, 108 were muntjac deer of which six (5.5%) were ELISA-positive (Holding et al. 2020). The detection of TBEV is important as it can infect humans, although the prevalence in deer appears to be quite low and generally localised; for example, in the Norfolk/Suffolk (Thetford Forest) focal area, serological evidence has shown a high prevalence of TBEV exposure (47.7% in the deer species found there, red, roe, fallow, muntjac) (Holding et al. 2020).

Deer have been linked to several enteric disease agents, including foodborne pathogens and diseases important to livestock health. An outbreak of *Escherichia coli* O157 in humans in Scotland in 2015 was linked to venison sourced from wild deer (Smith-Palmer et al. 2018). Pearce et al. (2023) reported a prevalence of 33.3% (stx2) in wild muntjac deer from across the UK suggesting they may be acting as a reservoir of disease and pose a risk to public health (Pearce et al. 2023). Several *Yersinia* spp. have also been detected in wild muntjac deer from across the UK at prevalences of 33.3% (*Yersinia enterocolitica*) and 4.17% (*Y. pseudotuberculosis*), which may pose a risk to public health (Pearce et al. 2023).

C.1. Damage to conservation habitats

(See roe deer for damage to conservation habitats). Impacts to conservation woodland are generated by all six species of deer to varying degrees, especially where deer densities are high (Quine, Shore, and Trout 2004). Muntjac prefer broadleaved woodland, such as that found in ancient woodland sites (Cooke 2021) and are one of several deer species in Britain that have caused progressive damage in conservation woodland due to increasing numbers and range (Dolman et al. 2010). Muntjac have significantly impacted substantial proportions of conservation woodland in Cambridgeshire causing irreversible damage (Cooke 2021) and will denude the understorey at very high densities (Cooke, Green, and Chapman 1996). At high densities, they have been associated with reductions in the abundance of rare or nationally important flowering plants, such as bluebells, dog's mercury and oxlips (White, Smart, et al. 2004).

D.1. Public nuisance

Unlike some other deer species in Britain, muntjac do not cause significant damage to agricultural or timber crops but can be of important concern to gardeners and allotment owners (British Deer Society 2023a). This may be an issue in areas of localised high density of muntjac.

Feral goat - *Capra aegagrus hircus*

Introduction

There are relatively few small and discrete populations of feral goats scattered around the UK, mostly residing in the hilly and mountainous areas of Scotland, Wales, Ireland and northern England (Putman 2008). Feral goats were first introduced to Britain as domestic livestock and some domestic goats continue to escape and become feral in some areas, mostly limited to areas where there are cliffs or rocky outcrops (Putman 2008).. The most recent population size estimates, dating from 1990-99, suggested there are between 5,000-10,000 individuals (Harris and Yalden 2008; Mathews et al. 2018). Very little updated information in literature searches.

A.1. Damage to forestry

Although feral goat populations are restricted to a small number of sites, bark stripping is common where goats have access to woodland (Putman 2008). Bark stripping and browsing by goats may cause more economic damage to trees and prohibit regeneration to a greater extent than either sheep or deer at equivalent densities, although it is difficult to separate bark-stripping damage between the different species (Putman 2008). Goats can strip bark up to a height of 2m and may climb trees causing damage higher up (Putman 2008). In oak woods on base-poor soils, trees with a smaller girth (1-15cm) are the most vulnerable to stripping, with a variety of tree species subject to bark-stripping (holly, ash, willow, oak, alder and hazel ) (Putman 2008).. Billies are more responsible than nannies for bark stripping damage, especially on mature trees (Rooney and Hayden 2002). Bark stripping increases the vulnerability of trees to fungal infection and may otherwise reduce their value as timber due to scarred or malformed growth (Arnold et al. 2018).

A.4. Competition with livestock

Farmers have claimed that feral goats compete with sheep for resources and that livestock fencing used to exclude sheep from areas is not a substantial enough barrier to prevent feral goats from gaining access (Putman 2008).

B.2. Reservoir of disease

(See roe deer for information on *Anaplasma phagocytophilum*). *A phagocytophilum* in feral goats in the UK has previously been reported in New Galloway in Scotland (Foster and Greig 1969). In Northern Ireland, four of five (80%) feral goats screened for the presence of *A. phagocytophilum* were positive (Harrison, Brown, and Montgomery 2012). Although only a small number of animals were sampled in Northern Ireland, the prevalence of infection was high, suggesting that feral goats may be a significant wildlife reservoir of this disease in the region and potentially an important reservoir wherever feral goats occur.

B.4. Reservoir of zoonotic disease

Feral goats have shown positive reactions for louping ill, Q fever (*Coxiella burnetti*), leptospirosis (*Leptospira grippotyphosa, L. icterohamorrhagiae, L. autumnalis*), and toxoplasmosis (*Toxoplasma gondii*) (Putman 2008). Domestic goats have been recorded as carriers of *Mycobacterium bovis* in the UK but there are no published records of feral goat infection (Delahay et al. 2002; Delahay et al. 2007).

C.1. Damage to conservation habitats

Damage to semi-natural woodland has been reported by conservationist interests (Putman 2008).

D.1. Public nuisance

Feral goats can move into suburban areas causing browsing and physical damage to gardens and allotments. In Llandudno, feral Kashmiri goats from the Great Orme headland have caused widespread damage to residents' properties and gardens since Covid-19, when public movement restrictions were in place. Current plans to reduce the number of goats includes relocating them and fertility control (BBC 2023a).

Wild boar - *Sus scrofa*
Introduction

The wild boar was native to Britain and may have died out and been reintroduced for hunting purposes (Dutton, Clayton, and Evans 2015). The species became extinct in the 17th century but recently became re-established through escapes from wild boar farms (Putman 2008). There are only four known self-sustaining viable populations in England and several in Scotland, including Lochaber and Dumfries and Galloway (Wilson 2014; Mathews et al. 2018). There are an estimated 2,600 wild boar living wild in Britain: 500 in England, 2,000 in Scotland and 150 in Wales (Mathews et al. 2018). They are mainly associated with woodland and despite their low numbers, they are still a cause for concern especially regarding crop damage, particularly where agricultural land borders woodland and public safety (Putman 2008; Mathews et al. 2018).

A.2. Damage to agricultural interests

Foraging wild boar have been reported to cause damage to grassland and some cereal crops, mostly during January – March and on land adjacent to woodland (Wilson 2005; NatureScot 2022b).

A.3. Damage to vehicles (RTAs)

As a result of the large size of wild boar, any collisions with traffic will cause damage to vehicles and the incidence of road accidents involving wild boar has increased since their reintroduction to the UK (Mammal Society 2015). The occurrence of boar-vehicle collisions will be localised and of more significance where densities are higher. Of 531 respondents in the Forest of Dean to a survey on wild boar, 49 (9.2%) had experienced a BVC of varying severity (Dutton, Clayton, and Evans 2015).

B.1. Health risks from RTAs

Traffic collisions with wild boar represent a health risk to both the vehicle occupants and the wild boar involved. A human fatality in an RTA involving collision with a wild boar occurred in 2015 on the M4 motorway in Wiltshire (BBC 2015b). This occurred outside the normal range of the nearest known wild boar population and it is thus possible that the animal concerned might have been an escape from a farm. Nevertheless, the cost of this single incident will have exceeded £1M given that the average cost per fatal RTA casualty in 2011 was reported to be £1.7M (DfT 2011).

B.2. Reservoir of disease

Wild boar could be a route by which diseases are transmitted to domestic swine (Bacigalupo et al. 2022), although the likelihood of contact between wild boar and domestic pigs is considered low (Defra 2008). They were initially disease free, having come originally from farms, but may encounter diseases in the wild and become a reservoir from which diseases could spread to domestic pigs (Putman 2008). The most significant impact of wild boar on disease risk is likely associated with the incursion and maintenance of exotic notifiable diseases, although the risk of this occurring is considered low (Defra 2008) despite increases in population size, density and distribution. Wild boar could potentially act as the reservoir for several serious viral diseases, including foot and mouth, Aujeszky's, classical swine fever, African swine fever and bTB (Wilson 2005).

B.3. bTB transmission

Although *M. bovis* has not been detected in wild boar in Britain (Delahay et al. 2002; Delahay et al. 2007), it was identified in farmed boar in southwest England in 2000 and subsequently associated with a breakdown in cattle on contiguous land (Delahay et al. 2002). It is also present in wild populations elsewhere in Europe, where it is considered an important host species (Justus et al. 2024). Wild boar are considered part of bTB disease host systems in a number of European countries, alongside a number of other species including cattle, fallow deer, roe deer, red deer and badgers (Hardstaff et al. 2014). As badgers are considered the main wildlife reservoir host for bTB in the UK, there is potential for disease transmission between badgers and wild boar where they come into contact with each other. Wild boar will typically overlap temporally with badgers and foxes, which are also nocturnal/crepuscular and spatially overlap with fallow deer, badgers and foxes (Caravaggi et al. 2018; Ogurtsov, Zheltukhin, and Kotlov 2018). As the focus of bTB wildlife studies in the UK is on badgers, it leaves many other species under surveyed; multi-species sampling should become a more frequent practice, especially in the UK and Ireland, to acquire a better understanding of the disease ecology of bTB (Justus et al. 2024).

<u>C.1. Damage to conservation habitats</u>

As wild boar have been absent from Britain for the last few hundred years, they could have an impact on some semi-natural ecosystems which now have conservation value, with some fragile ecosystems being adversely affected by extensive rooting (NatureScot 2022b). Evidence concerning the feral populations and distribution suggests that their impact is likely to be minor, providing the populations do not reach significantly higher densities than those currently present (Wilson 2005). There is a potential conflict with the hazel dormouse, which is listed as being of conservation concern; (Rozycka et al. 2015) reported fewer dormice in woodlands which contained feral boar, possibly because boar root for food on the woodland ground where hazel dormice hibernate in fragile nests thus impacting their populations through predation.

<u>D.1. Public nuisance</u>

Wild boar are shy, nocturnal creatures that would normally choose to run away from people (Graves 1984), however, there is likely to be perceived safety risk to the public. In a survey of 531 people in the Forest of Dean, around 120 (23%) reported being chased by wild boar, particularly if accompanied by dogs (Dutton, Clayton, and Evans 2015). The Forestry Commission database (April 2013-May 2015) only reported one such incident (Dutton, Clayton, and Evans 2015). Reports of wild boar in some European cities, posing a risk to the inhabitants, are occasionally reported in the popular press and media; however, despite a wild boar population in Europe estimated at around 1.25 million, Wilson (2005) did not find any reports of attacks or serious injuries caused by wild boar in Europe in the literature. Given their wide distribution and substantial populations throughout much of their range the risk of attack and injury is clearly very small.

### Carnivora

Feral cat - *Felis catus*

<u>Introduction</u>

Feral cats in the UK are free-living domestic cats. In addition, feral cats have been deliberately introduced to some islands for the control of rabbits (Gorman 2008). The current population size and distribution is not well known, although it is believed that there were probably upwards of 9 million cats in Britain in 1980, of which approximately 20% could be classed as feral (Tabor 1981). It is likely that that figure has since grown. Feral cats can also be difficult to categorise as they exist with varying degrees of dependence on people and can be found in both urban and farmland habitats (Gorman 2008). Its ability to live in a diverse range of situations and habitats means that it comes into conflict with other species of human interest.

<u>B.2. Reservoir of disease</u>

Feral cats can carry endoparasites such as *Toxocara cati*, *Toxascaris leonina* and *Toxoplasma gondii* (Yamaguchi et al. 1996). According to Fisher (Fisher 2003), the role of *T. cati* as a zoonotic parasite has long been underestimated. It is a roundworm commonly found in cats that can cause ocular larval migrans and visceral larval migrans; in can infect a high proportion of cats, for example, 84.8% of stray cats in a survey conducted in Denmark, and 91% of feral farm cats in the UK were found to be infected. (Denmark: Takeuchi-Storm et al. (2015); UK: Yamaguchi et al. (1996)). Nichol, Ball, and Snow (1981) found that of 92 feral cats necropsied from the London and Sheffield areas, 49 (53.3%) were infected with *T. cati*, 32 (34.8%) with *Dipylidium caninum*, 11 (12.0%) with *Taenia taeniaeformis* and 1 (1.1%) with *T. leonina*. *Isospora felis* was found in 3 (4.3%) of 69 samples of faeces. Twenty-two cats (23.9%) were simultaneously infected with two helminth species. Sources of infection are likely to be environmental as the eggs are persistent (Yamaguchi et al. 1996), making children vulnerable from mud-eating habits.

<u>C.3. Threat to genetic integrity</u>

The genetic integrity of the wildcat is threatened by interbreeding with feral cat populations throughout Europe, with studies having been conducted in countries including Hungary, Italy and Scotland (Beaumont et al. 2001). The wildcat is only found in the northern half of Scotland and there is therefore risk of both local and national extinction. The distribution of wildcats is difficult to assess due to the presence of and changes in reporting of hybrids (Sainsbury et al. 2019), therefore there is a lack of clear baseline from which to evaluate genetic changes.

<u>C.4. Competition with or predation of wildlife (including protected or conservation species)</u>

Feral cats are perceived to threaten bird populations through predation (Mead 2008). However, domestic and feral cats are difficult to distinguish due to the varying degrees of dependence on humans (Gorman 2008)In the United States it has been estimated that free-ranging cats kill 1.3–4.0 billion birds and 6.3–22.3 billion mammals annually with feral cats, as opposed to domestic cats, causing the majority of this mortality (Loss, Will, and Marra 2013). Woods, McDonald, and Harris (2003) estimated that the British population of approximately 9 million domestic cats brought home, over a period of five months, in the order of 92 (85–100) million prey items in the period of this survey, including 57 (52–63) million mammals, 27 (25–29) million birds and 5 (4–6) million reptiles and amphibians.

In a review into the suitability of urban refugia for Eurasian red squirrels, Fingland et al. (2021) noted that many of the predation events that affect red squirrels appear to be due to free-ranging domestic and feral cats, although there is little evidence to suggest that predation limits urban red squirrel populations. On Jersey, cat predation accounted for 5% (17/337) of red squirrel mortalities (Blackett et al. 2018).

Wildcat - *Felis silvestris*
<u>Introduction</u>

The wildcat is a critically endangered species native to the UK but is now only found in the northern part of Scotland (Mathews et al. 2018). They were historically widespread throughout Britain but a population survey undertaken by the former Nature Conservancy Council found population densities to be low, even in areas of suitable habitat (Battersby 2005), typically a mosaic of habitat types with core of broadleaved or mixed forest. Their decline in numbers is likely to be multi-factorial, including destruction of habitat, accidental killing by dogs, snares and poison baiting, viral diseases, hybridisation with the feral or domestic cat and persecution resulting from attacks on game birds and livestock (McOrist and Kitchener 1994). The main increasing threat is hybridisation with feral or domestic cats (Kilshaw et al. 2015). There are estimated to be around 200 wildcats living in Scotland, although this figure is highly likely to include feral cats or hybrid wildcats (Mathews et al. 2018).

<u>A.5. Predation of game species</u>

Wildcats are regarded as vermin by gamekeeping interests (pheasant, red grouse) and many are killed every year, although the evidence for wildcats as a major predator of grouse is poor (Gorman 2008). Given the very low numbers of wildcats, this economic impact is likely to be negligible.

Otter - *Lutra lutra*
Introduction

The otter is a mustelid species native to Britain that has responded positively to conservation measures in terms of both abundance and distribution (Crawford 2003). It is widespread across the UK but remains scarce with a dispersed population after a serious decline in numbers in the 1950s due to habitat fragmentation and pollution (Chanin and Jefferies 1978). The European otter is protected under the Wildlife & Countryside Act (1981), Habitats Directive Annex IV as a European Protected Species and is generally considered to be an iconic species of British wildlife (Macmillan and Phillip 2008), but it can cause conflict with the inland fisheries industry. Applications can be made to Natural England for licences that would allow otherwise unlawful activities in order to resolve conflicts with this protected species (Natural England 2011). It is estimated that there are around 11,000 otters living wild in Britain: 2,900 in England, 7,100 in Scotland and 1,000 in Wales (Mathews et al. 2018).

A.6. Predation/impact on commercial fisheries

Fish constitute the most significant component of the otter's diet, drawing it into direct competition with inland commercial fisheries and recreational anglers (Allen et al. 2020). Recreational angling is worth an estimated £1.4B annually to the English economy (GOV.UK. 2018), with several thousand pounds of investment claimed to be at risk if an otter has chosen to den nearby (timesonline.co.uk). Specimen carp fisheries may be a particular problem due to the value of individual fish reaching several thousand pounds (Crawford 2003). In the Upper Thames Valley, lowland England, fish represented 46% of total otter diets with fish of "potential value" (e.g. commercial or sporting value) representing 19% (Grant and Harrington 2015). Although cyprinids were the most consumed fish of potential value, only a single otter spraint contained potentially valuable carp. Britton et al. (2005) also reported a similar finding with evidence of carp consumption in only two of 98 otter carcasses from across the UK. In Poland, serious damage was attributed primarily to otters on 56% of farms, with otter believed to be the only species responsible for losses of large "marketable" fish, i.e. stocks of the greatest economic value (Kloskowski 2010).

C.4. Competition with or predation of wildlife (including protected or conservation species)

Research on otter predation in smaller water bodies such as ponds, which have high levels of biodiversity compared to other inland waters in Europe, have shown otters display great plasticity in foraging behaviour, taking a wide range of prey species (Almeida et al. 2013). These authors reported birds as the most important prey category consumed at ponds in Norfolk, along with European eel, common frogs, common toads, northern pike and tench. Non-native carp were also an important prey item whereas native crucian carp were considered a minor prey item. Mammals are taken very rarely, with rabbits and water voles being the preferred mammalian prey species (Carss 1995). The otter has been found to prey on black guillemots in Shetland resulting in a 10.5% failure rate of occupied nests (Ewins 1985). The black guillemot is protected under the Wildlife and Countryside Act, 1981, although has recently moved from amber to green conservation status in the UK due to minimal change in its breeding range (Stanbury et al. 2024). Predation on wild salmon may also represent conflict between two species of conservation concern; Carrs, Kruuk, and Conroy (1990) found that a considerable number of adult salmon were killed by otters during the spawning period, although the majority were male and therefore unlikely to affect the breeding success of the salmon population.

**Small mustelids**

Pine marten – *Martes martes*
Stoat - *Mustela erminea*
Feral ferret - *Mustela furo*
Weasel - *Mustela nivalis*
Polecat - *Mustela putorius*

### Introduction

Pine martens, stoats, feral ferrets, weasels and polecats are small members of the Mustelidae family found in Great Britain. Of these five species, only the feral ferret is non-native, introduced to Britain around the 11th century and now widely kept as pets, escapes of which may have led to established populations throughout the UK (Gorman 2008). The stoat and weasel are both widespread throughout the UK, whereas the polecat and pine marten have more patchy distributions (Mathews et al. 2018). The pine marten is mainly found in northern Scotland with pockets in England and Wales, mostly confined to rugged upland areas although the viability of these populations is uncertain (Bright and Halliwell 1999; Mathews et al. 2018). The polecat is concentrated in Wales and the English midlands, despite being previously widespread, due to persecution during the late 19th century (Gorman 2008). Reintroductions in the 1970s have caused populations to become re-established in a few pockets of England, with consequential increases in number and range (Birks and Kitchener 1999; Mathews et al. 2018). According to Mathews et al. (2018), recent population estimates for free-living small mustelids in Britain are as follows: around 3,700 pine marten (3,700 in Scotland, 51 in Wales (based on a release-monitor programme between 2015 and 2017 by the Vincent Wildlife Trust); 438,000 stoat (260,000 in England, 140,000 in Scotland, 37,600 in Wales); unknown population estimate for feral ferret (although likely to have broad geographical range due to domestication); 83,300 polecat (66,400 in England, 16,800 in Wales). There are an estimated 450,000 weasels in Great Britain (www.ptes.org).

#### A.5. Predation of game species - pine marten, stoat, feral ferret, weasel, polecat

Conflicts with the management of game are arising due to the increase in range of the pine marten, but mostly in Scotland (Gorman 2008). The stoat can have effects on wild game bird populations but does not seem to cause substantial damage to reared game birds (Warren and Baines 2002). In a study of red grouse at two North Pennine moors between 2013-2015, stoats or raptors were responsible for killing around 40% of diseased birds and 30% of healthy birds (Baines et al. 2018). Red grouse numbers have also been observed to increase in response to experimentally reduced levels of key predators, including stoats and to a lesser extent weasels (Fletcher, Hoodless, and Baines 2013). Historically stoats have been considered a threat due to the predation on game birds; they are considered to have a lesser impact than foxes, feral cats and mink but a more serious impact compared to rats, hedgehogs, polecats, feral ferrets and weasels (Packer and Birks 2002).

#### A.7. Predation of poultry - pine marten, feral ferret, polecat

The pine marten and polecat will both readily prey on domestic poultry if given the opportunity but are also relatively easy to keep out of areas where poultry is housed (Gorman 2008). The impact of feral ferrets is considered low and less of an impact than on game birds.

#### B.2. Reservoir of disease – stoat

A case of *Angiostrongylus vasorum* infection was reported in a stoat found as an RTA in west Cornwall in 2010 (Simpson 2010). *A. vasorum* has a patchy but fairly widespread distribution through most of western Europe, infecting dogs and other canids, causing cardiorespiratory disease and coagulopathy. There are well documented hotspots of infection in southeast England and south Wales, with cases also reported in northern England and Scotland (Tappin 2014), with foxes considered as a reservoir for canine infection (Morgan et al. 2021). There is limited information for stoats, suggesting it poses little risk to other species.

B.4. Reservoir of zoonotic disease – polecat

*Toxoplasma gondii* DNA has been detected in polecats in Britain (Burrells et al. 2013). As polecats are known to live near farms and other human settlements, they may pose a risk for transmission to cats and livestock (Burrells et al. 2013; Veronesi, Deak, and Diakou 2023).

C.3. Threat to genetic integrity – feral ferret

Ferrets are bred in captivity as pets or for use in rabbit population management, but many have escaped or been released with consequent potential to interbreed with the native polecat (Birks and Kitchener 1999). Hybrids have also purposely been bred to increase aggressiveness for hunting rabbits and to improve appearance in polecat classes at ferret shows (McKay 1995). The existence of hybrids makes it difficult to determine the true status of the British polecat population and interbreeding could jeopardise the remaining individuals of the genetically pure breed, which are already not particularly widespread (Gorman 2008). Etherington et al. (2022) found that English polecats were highly introgressed with domestic ferrets, which may have contributed to their recent increase in population size.

C.4. Competition with or predation of wildlife (including protected or conservation species) – pine marten, stoat

Baines, Aebischer, and Macleod (2016) found that female capercaillie across 26 Scottish forests surveyed between 1991 and 2009 reared more chicks in forests with lower pine marten indices. They concluded that the reduction in breeding success and continuing decline of capercaillies within small, fragmented forests may be due to a number of factors, including predation of clutches and chicks by martens and crows, as well as climate change. Capercaillies are at risk of extinction in the UK and as such, are a priority species under the EU Birds Directive.

Stoats were introduced to the Orkney archipelago in 2010, posing a serious threat to native wildlife, such as the endemic Orkney vole, *Microtus arvalis orcadensis*, and ground-nesting birds (Fraser et al. 2015). A partnership was formed in 2016 between Scottish Natural Heritage (later NatureScot), the RSPB and the Orkney Islands Council with the aim of eradicating stoats across the Orkney archipelago to safeguard native biodiversity (Zub et al. 2022).

Pine martens have also been reported to have positive impacts on protected species in the UK, such as red squirrels. Recent research has demonstrated that as pine marten landscape use intensity increases, so does red squirrel occupancy, likely linked to parallel declines in grey squirrel occupancy (Bamber et al. 2020; Twining et al. 2022).

D.2. Domestic nuisance – pine marten

There have been cases reported of pine martens making dens and breeding in homes and buildings in Scotland and Ireland (Birks, Messenger, and Halliwell 2005). Furthermore, they may occasionally visit bird tables for food (Gorman 2008). This has created conflicts with householders and led to requests to relocate pine martens despite full protection under Schedule 5 of the Wildlife and Countryside Act, 1981 (Brown and Birks 2006).

Badger - *Meles meles*
Introduction

The badger is a native species which is widespread across Britain but is particularly abundant in the southwest of England and south Wales (Judge et al. 2014). Optimal badger habitat consists of deciduous woodland associated with earthworm-rich pasture in regions which have mild and wet climatic conditions, although they have also been seen to inhabit urban areas (Gorman 2008). Badgers are generalist omnivores, taking a wide range of food items. Predation on the eggs of ground-nesting birds is relatively low level as they are only an occasional food item and damage to significant proportions of nests is rare (Gorman 2008). The predation of birds themselves may be similar to the levels discussed in below regarding game birds, as it is difficult to identify bird species when they are present in badger diets (Hounsome and Delahay 2005). Badger predation is a known cause of hedgehog mortality and hedgehog abundance has been found to increase in areas where badgers are culled (Trewby et al. 2014). There are estimated to be between 562,000 and 760,000 badgers in Britain (variation in estimate is due to exclusion/inclusion of broadleaved woodland as a suitable habitat): 384,000 to 519,000 in England, 115,000 to 156,000 in Scotland, 62,900 to 85,000 in Wales (Mathews et al. 2018).

A.2. Damage to agricultural interests

A survey of 3,600 landowners (of which 1,982 responded) was conducted in 1997 to determine the extent of badger damage to agriculture in England and Wales with the most frequently reported damage being burrowing activities, such as under fences, and damage to crops such as wheat, forage maize and vines (Moore et al. 2001). Damage was widespread but the incidence was highly dependent on region and land use, and most was of little economic importance. Costs of over £1K were reported to have been incurred by 5% of respondents so using the respondents as a representative sample, the mean estimated national cost of badger damage was £41.5M per annum. Assuming that all non-respondents had no damage, the estimated cost was £21.5M per annum, with most of the damage being attributed to burrowing activities (Moore et al. 2001). Recent estimates allowing for inflation would be approximately £39.0M (Bank of England 2024). Approximately 25% of the 600-700 licence applications under the Protection of Badgers Act received by Natural England between 1994 and 2004 were for the purpose of preventing damage to agricultural land, which was the most commonly cited problem (Delahay et al. 2009). In Scotland, damage by badgers to growing crops is reported by farmers although the levels of concern are significantly lower than in England, probably due to lower numbers; in most cases, reported economic losses are small although may be significant enough to warrant the issue of a licence to remove them (Mitchell-Jones 2020).

A.3. Damage to vehicles (RTAs)

Roads represent a significant source of mortality for wildlife, with road-traffic collisions being the leading observed cause of mortality in European badgers (around 50,000 annually) in the UK (Harris et al. 1992). No recent information was identified. Badgers show a bimodal peak in road-kill records, peaking around early spring and autumn (Raymond et al. 2021), which coincide with the main breeding seasons. Damage can be caused to vehicles, given the large body size of badgers, but will not cause the same levels of damage as deer.

A.5. Predation of game species

Historical rates of game bird predation by badgers are low, affecting 2-3% of partridge and pheasant nests (Potts 1986; Hill and Robertson 1988). More recent rates appear to be between 10-13% for grey partridge and corn bunting nests (Reynolds et al. 1992), although birds are only present in approximately 8% of British badger diet samples (Hounsome and Delahay 2005). Predation of chickens has been reported although the only evidence is derived from poultry found in badger diets, which may be from scavenging rather than predation (Gorman 2008). Limited recent information.

A.8. Damage to property

Between 1994-2004, the number of licence applications received by Natural England rose significantly, with the reason for requiring the licence cited as prevention of damage in 64% of the cases (Delahay et al. 2009). Damage to buildings and gardens accounted for 10.9% and 10.7% of successful applications, respectively (Delahay et al. 2009). An increasing proportion of applications came from the east of England and from urban as opposed to rural areas; although licence applications are more common in the south and east of England, they have risen in every region of the country (Delahay et al. 2009). Davison et al. (2010) estimated a cost of between £5K to £10K for the exclusion of badgers from a modest-sized sett on residential property.

A.9. Damage to amenities and domestic grassland

Domestic grassland, such as parks and golf courses, are ideal foraging grounds for badgers, with reports of them devastating or even destroying large areas of turf on golf courses (GreenKeeping 2019). They are considered less damaging than other species, such as rabbits, Canada geese, moles and mink, with rabbits reported to cost more in terms of control (Williams et al. 2010).

A.10. Conflicts with construction, infrastructure and development

Badgers and their setts are protected by law, therefore activities such as construction or repair of flood defences or watercourses that risk disturbance require that a licence is obtained (GOV.UK 2015). Licence applications for development projects must involve expert input, such as surveying and identifying mitigation methods to offset the impacts of construction (GOV.UK 2015).

Badgers can excavate near to roads and cause economic damage as a result. In 2024, Lincolnshire County Council were granted a licence from Natural England to carry out essential repair work to a coastal road after badgers caused £100K of damage by tunnelling underneath it (Lincolnshire Live 2024).

B.1. Health risks from RTAs

Details on the levels of human injuries sustained as a direct result of badgers are very difficult to obtain, as the Department of Transport road accident injury statistics form (STATS19) does not require the type of animal involved to be reported unless it is a ridden horse (Langbein 2007). Given their nocturnal behaviour and size, they are likely to have less of an impact on human injuries or fatalities caused by road-traffic collisions than, for example, larger deer.

In a review of bTB infection in wild mammals in southwest England, Delahay et al. (2007) reported that badgers had the highest prevalence of infection (10.94%) compared to all other species sampled. In a separate study, Powell et al. (2024) reported an overall prevalence of 6.5% (ranging from 1.1 to 13.0%) from 525 badger carcasses collected from the southern edge of England's bTB epidemic (Oxfordshire, Berkshire, Buckinghamshire, Hampshire and East Sussex). Of these, 443 (84.3%) carcasses were from RTAs, mostly collected by farmers or farming groups (44.0%), conservation groups (25.1%), veterinarians (16.2%), government agencies or local authorities (3.4%) or other groups or individuals (11.2%). Given that a high number of badgers are killed annually on British roads and disease prevalence is high, badgers may pose a risk if the come into contact with the public, with the risk likely to be higher in badger bTB hotspot areas.

B.2. Reservoir of disease

Badgers are known to harbour a wide range of pathogens (Guardone et al. 2020). In particular, parasites belonging to the order Piroplasmida, such as badger-associated *Babesia* parasites (suspected to belong to the *B. microti* group), have been detected in several European countries (Lindhorst et al. 2024). Badger-associated *Babesia* and other *Babesia* spp. have been described in free-ranging canids and mustelids in southern Italy (e.g. (Santoro et al. 2019)) to the extent that these hosts may be playing an important role in the maintenance of the sylvatic cycle of these parasites. In the UK, badger-associated *Babesia* was confirmed in 50% (9 of 18) of badger carcasses collected as RTAs across Cheshire between 2016-2017 (Guardone et al. 2020). These authors concluded that the presence of this type of *Babesia*, which has been found in a range of canids and clinically affected domestic dogs elsewhere in Europe, highlighted the need to investigate the potential risk posed by this putative pathogen for other wild carnivore and domestic animals.

Other parasites have been identified in badgers, including *Trypanosoma* (*Megatrypanum*) *pestanai* (in the UK, France and Italy) and filarioid nematodes (*Diroflaria immitis* and *D. repens*), although the role of the badger as a definite host and a reservoir for *D. immitis* and *D. repens* remains unclear (Lindhorst et al. 2024). Infections with *Anaplasma phagocytophilum*, *Ehrlichia* sp. (Battisti et al. 2020) and *Candidatus* Neoehrlichia sp. (Hornok et al. 2017) have been identified in badgers in different European countries including, Italy and the Netherlands. Other pathogens that have been detected in badgers are *Bartonella* spp. (Gerrikagoitia et al. 2012), *Rickettsia* spp. (Jurczyk et al. 2022) and *Borrelia* spp. (Gern and Sell

2009). However, of the 18 badgers sampled from Cheshire between 2016-2017, there was no molecular evidence of *Anaplasma phagocytophilum*, *Ehrlichia canis*, *Coxiella burnetii*, *Francisella tularensis* or *Bartonella* spp. (Guardone et al. 2020). Given the small sample size of this study and the reported presence of several pathogenic parasites in badgers elsewhere, further investigations in the UK might be warranted.

B.3. bTB transmission

The badger is the focus of ongoing controversy regarding management options for controlling the spread of *M. bovis* to cattle, which is estimated to cost the taxpayer over £100M annually (Defra 2024). Badgers are considered to be a source of bTB in cattle from direct contact with cattle, contact with troughs and contamination of feed in farm buildings (Tolhurst et al. 2009). A 54% decrease in cattle bTB incidence was observed in the proactively culled areas of the Randomised Badger Culling Trial (RBCT) (Jenkins, Woodroffe, and Donnelly 2008). The estimated minimum contribution of badgers to confirmed tuberculosis in cattle in high-incidence areas in England is 38% (Donnelly and Nouvellet 2013). Powell et al. (2024) also reported that the prevalence of *M. bovis* in badgers (6.5%) was significantly and positively correlated with the number of incident cases observed in cattle at the county level (see Health risks from RTAs above). In Northern Ireland, Milne et al. (2020) found that the same *M. bovis* genetic types were spatially co-localised in cattle and RTA badger hosts, indicative of a shared epidemic. However, Swift et al. (2021) reported that there were variations in the *M. tuberculosis* complex (MTC) spoligotypes from badgers and cattle in areas on the spatial edge of the cattle epidemic (across six counties including Cheshire, Derbyshire, Nottinghamshire, Leicestershire, Warwickshire, Northamptonshire). In the northern counties, spoligotype SB0129 predominated in both cattle and badgers, but elsewhere there was a much wider range of spoligotypes found in badgers than cattle, in which infection was mostly with the regional cattle spoligotype. The authors concluded that with the exception of Cheshire, they found little evidence to link the expansion of the bTB epidemic in cattle in England to widespread badger infection.

C.4. Competition with or predation of wildlife (including protected or conservation species)

Badgers have been reported to compete with or predate on several wildlife species, including some of conservation concern. One of these species is the European hedgehog, which is facing a decline in numbers across parts of its range (Mathews et al. 2018). Badgers are considered as intraguild predators of hedgehogs, not only sharing food resources, such as earthworms and snails, but also predating on hedgehogs (Hof, Allen, and Bright 2019). Trewby et al. (2014) reported that the counts of hedgehogs in areas of preferred grassland (amenity grassland) more than doubled over a five-year period of badger culling during the Randomised Badger Culling Trial, concluding that the management of a top predator in this controlled experiment provided evidence of mesopredator release. In a separate study, Hof, Allen, and Bright (2019) investigated the role of the badger in the nationwide distribution of hedgehogs in England; they concluded that, while habitat-related factors explained more variation in the likelihood of hedgehog presence, badger presence was also negatively impacting the presence of hedgehogs. As hedgehogs were less likely to be found in areas where badgers were more likely to be found, intraguild predation by badgers may be influencing the distribution of hedgehogs in England.

Badgers are also well-known predators of bumblebees, which are of conservation concern due to the decline of some species across the UK (Goulson et al. 2010). Roberts (2019) found that badgers dug up 125 of 574 (21.8%) artificial bumblebee nests at Woodchester Park, Gloucestershire, with no other species observed on camera destroying nests. In a study of 100 bumblebee nests located mainly in rural locations around Stirling, in central Scotland, the biggest cause of mortality was caused by large animals, most likely badgers, destroying half (50%) of the nests surveyed (Goulson, O'Connor, and Park 2017). Although only one badger was observed at a nest during the study, dietary evidence from previous studies confirms that badgers regularly consume bees (Goulson, O'Connor, and Park 2017). As with many eusocial hymenopterans, each nest represents a single breeding female, hence the population trajectory of a species will depend on the frequency of success or failure of nests (Chapman and Bourke 2008). Badgers may therefore be contributing significantly to the decline of bumblebees.

Badgers have also been reported to predate birds of conservation concern, including lowland waders, some of which have undergone significant breeding population declines and range contractions in recent decades (Wilson, Ausden, and Milsom 2004; PECBMS 2012). Malpas et al. (2013) investigated the effect of predator-exclusion fencing at 10 sites across England, Wales and Northern Ireland, selected based on the presence and identification of mammalian predators as the major cause of low wader breeding success; they found foxes accounted for 63% of nest predation whilst badgers accounted for 13% (RSPB unpublished data). Badgers can also predate other nests of ground-nesting birds, such as the wood warbler, leading to breeding failure and implications for their population dynamics (Maag et al. 2022).

American mink – *Neovison vison*

Introduction

The American mink (hereafter referred to as mink) is a semi-aquatic mustelid that was introduced to the UK in 1929 for use in the fur trade (Thompson 1962). Populations of mink were first recorded in the wild during the 1950s, introduced either through escapes, or larger, purposeful releases (Thompson 1968), after which the populations increased and are now widely distributed throughout Britain (Strachan and Jefferies 1996; Mathews et al. 2018). Mink farming in the UK ended in 2003 (Fur Farming (Prohibition) Bill, 2000) and population estimates twenty years ago indicated a decline in numbers, which has been linked to re-colonisation by the otter (Jefferies 2003; McDonald, O'Hara, and Morrish 2007). Mink populations have been monitored as an invasive species as part of the National Otter Survey since the mid-1970s due to their predation on aquatic and semi-aquatic species including some species of conservation importance. Mink are generalist predators showing a strong preference for riparian habitats, feeding on a wide range of prey including waterfowl, fish and water voles (Mathews et al. 2018); their predation on such species has resulted in them becoming the focus of population management to try and reduce the levels of conflict they cause (Bonesi and Palazon 2007). It is estimated there are 122,000 free-living mink in Britain: 62,400 in England, 46,600 in Scotland and 12,900 in Wales (Mathews et al. 2018).

A.5. Predation of game species

Mink are known to prey on reared game birds. Mass kills of birds such as pheasants, sometimes of over 100 individuals have been reported (Harrison and Symes 1989). These reports are now outdated and do not provide any useful indication of how frequent or widespread such occurrences were. No recent information identified.

A.6. Predation/impact on commercial fisheries

Despite the reports of predation by mink in commercial fisheries, as well as damage to fish cages and the stressing of fish which can cause mass death of stocks (Harrison and Symes 1989; Moore, Robertson, and Aegerter 2000), there have been very few studies conducted of economic impact (Williams et al. 2010). In Wales, mink were perceived to be present and having a detrimental impact at 35% (15/43) of fisheries (Whitby et al. 2010). Trout farms are particularly at risk due to their similar requirements for high water quality; a study conducted in Scandinavia found that 16% of trout farmers questioned considered mink to be a significant problem (Harrison and Symes 1989). Although predation of fish was the most frequently reported problem, other issues, such as the scratching or stressing of fish, could have more economically important consequences through larger indirect fish losses (Harrison and Symes 1989). Moore, Robertson, and Aegerter (2000) reported a cost of £11.6K by one farm in the Western Isles, where mink had been blamed for the release of 14,500 smolts.

A.7. Predation of poultry

A study conducted in Denmark questioned 692 poultry keepers about the frequency of mink attacks; 27% of respondents reported attacks during a period of six years, with 24% of respondents reporting fewer than five attacks (Hammershoj and Asferg 2000), suggesting that mink attacks are unlikely to have large economic consequences for poultry keepers. Harrison and Symes (1989) also suggested that the economic impacts are likely to be small and concentrated on smallholdings, as most commercially reared poultry has been reared indoors in mink-proof buildings since the 1950s. Nevertheless, the Scottish National Heritage Hebridean Mink Project (SNH 2008) have recently stated that mink predation on domestic poultry has caused some businesses to cease the keeping of poultry altogether. The predation of poultry by mink could therefore have larger localised impacts where all businesses involved are small scale, such as the Western Isles. Williams et al. (2010) estimated the annual economic cost of mink predation on free-range poultry to be £77.5K in England, £37.9K in Scotland and £24.0K in Wales giving a total of £139.3K. Eschen et al. (2023) revised these estimates recently, with costs of £68.8K in England, £33.6K in Scotland and £21.2K in Wales.

A.9. Damage to amenities and domestic grassland

Mink have been identified as one of several species that can cause damage to grassland amenities, such as golf courses, accounting for approximately 5% of control costs (rabbits account for 50%, moles account for 25% and Canada geese account for 20%: (Williams et al. 2010)). Williams et al. (2010) estimated that the annual cost of controlling mink in Britain was £528.4K, with £384.8K in England, £35K in Wales and £108.6K in Scotland. Eschen et al. (2023) updated these costs recently, estimating mink costs to golf courses in Britain was approximately £800K, with £500K in England and £200K in Wales.

B.4. Reservoir of zoonotic disease

Mink are important reservoir hosts for many zoonotic pathogens, including parasites (Veronesi, Deak, and Diakou 2023). In the UK, mink have been found to be serologically and molecularly positive for *Toxoplasma gondii* (Burrells et al. 2013); in addition, infection with *Cryptosporidium* has been detected in mink from Ireland at a prevalence of 6.17% (Veronesi, Deak, and Diakou 2023). These authors concluded that, given both *T. gondii* and *Cryptosporidium* are often water transmitted and mink prefer habitats near to water, they may be important spreaders of the parasites.

C.4. Competition with or predation of wildlife (including protected or conservation species)

Mink are known to prey on seabird colonies and waterfowl, and are frequently reported to limit their prey (Roos et al. 2018). A study on the upper Thames found that waterfowl, such as coots and moorhens, were the fourth most selected prey after rabbits, fish and small mammals and the impact of mink predation during the bird breeding season was moderate to high (Ferreras and MacDonald 1999). Conversely, Halliwell and Macdonald (1996) found no evidence that mink were limiting moorhen abundance in the upper Thames catchment. Seabird species affected include the tern, common gull and black-headed gull and it is unusual for any chicks to fledge from a colony of less than 200 breeding pairs when a mink is present (Craik 1997). One important factor which leads to such large-scale impacts on breeding colonies of birds, in comparison to the size and requirements of the mink, is that the mink tend to remove all or most of the chicks or eggs and cache them (Craik 1991, 1990). Consequently, colonies have either moved or become greatly reduced in size and it is likely that similar impacts are being experienced in other areas (Craik 1997). The declines recorded by Craik (1997) amounted to 2.2 and 4.5% of common gull and common tern numbers respectively for the entire British Isles.

Mink are also strongly associated with the decline of native water vole populations in the UK. Between 1939 and 1998, water voles experienced a cumulative loss of occupied sites of 98.7% across all regions of England, Scotland and Wales (Moorhouse et al. 2015), with mink considered as a contributing factor (Lambin, Horrill, and Raynor 2019). Habitat destruction and fragmentation, disturbance and environmental contamination are other factors which have contributed to the decline of water voles (Halliwell and Macdonald 1996), but the feral mink has lessened their ability to cope by predating on this already vulnerable species. Barreto et al. (2006) found the presence of mink to be the most significant variable in determining the distribution of water voles.

The distribution of the native polecat is restricted in the UK due to persecution and the contamination of its habitat (Strachan and Jefferies 1996; Birks 2000). Unlike its counterparts in mainland Europe, the polecat in Britain does not show a preference for riparian habitats, possibly to avoid competition with the American mink (Mathews et al. 2018; Brzeziński et al. 2021). American mink are considered as possible drivers of polecat decline in Europe (Melero, Palazon, and Lambin 2014), but it is difficult to determine the long-term consequences of interspecific competition between the two species (Brzeziński et al. 2021). However, a dietary study conducted in France found that polecats and mink were able to coexist by temporal space segregation and the selection of alternative prey (Lodé 1993). This suggests that habitats in the UK with a diversity of prey types should be able to support both polecats and mink.

Red fox - *Vulpes vulpes*

Introduction

The red fox is a native British carnivore which is widespread across the UK, although with lower densities in the northwest of Scotland (Gorman 2008). The red fox is common in urban areas although colonisation of new towns and cities may be constrained by outbreaks of sarcoptic mange (Wilkinson and Smith 2008). It has been widely persecuted; fox populations were supplemented using imports from Europe in the 19th century to improve hunting in localised areas (Lever 1994), although hunting foxes with more than two dogs was banned in the early 2000's (Protection of Wild Mammals (Scotland) Act 2002 and the Hunting Act 2004). The red fox inhabits a wide variety of habitats, as it is a generalist and can take advantage of many types of cover and food (Gorman 2008). Baker, Harris, and White (2006)estimated there were approximately 250,000 adult foxes in the UK at the start of the breeding season (from three separate estimates). More recently, Mathews et al. (2018) suggested this estimate has increased, with 357,000 foxes in Britain: 255,000 in England, 74,000 in Scotland and 27,700 in Wales.

A.5. Predation of game species

There is strong evidence that fox predation has a significant effect on wild game birds and hare populations, but not as great as for other ground-nesting birds, making game conservation a major reason for controlling fox populations (Macdonald et al. 2000). Fox predation on pheasants in release pens is low and the most significant losses occur after the pheasants have been released and can no longer be directly protected so higher release rates may be required to compensate for losses to commercial shoots (Baker, Harris, and White 2006). Reports indicate that the red fox is a significant game predator particularly during the summer months as game birds become more available (Dell'Arte et al. 2007). Sage et al. (2018) analysed data over a 25-year period on radio-tracked released pheasants on lowland farmland in the UK, with seven sites during spring and summer reporting predation rates of between 20% and 71%, mainly by foxes. Foxes will eat both the birds and their eggs, targeting all sections of the game bird life cycle. At three of four sites with low predator control, between 5% and 22% of nest failures were caused by incubating hens being predated by foxes (Sage et al. 2018). The importance of particular prey species will depend upon the habitat, for example, predation on grouse appears to be most likely when there is a crash in the vole population cycle (Selås 2006). The impacts of fox predation may be long-term but have been shown to be reversible when fox numbers decline (Selås 2006). Ludwig et al. (2017) found that grouse moor management, which included control of generalist predators, had a positive effects on the abundance and breeding success of grouse, which were two-to three-fold higher when fox indices and crow abundance were reduced by 50-70%.

A.11. Predation of livestock

The predation of livestock by foxes tends to be low level but is widespread due to their distribution and can become locally important where specific farms are targeted (Macdonald et al. 2000). Foxes can take a variety of small livestock, such as lambs, poultry and piglets but will be far less likely to take animals that are housed (Baker, Harris, and White 2006). In a questionnaire survey of 220 farmers in Wiltshire, almost two-thirds did not consider the fox to be a personal pest, although most believed they should be controlled everywhere; considerably less believed foxes were actually taking domestic livestock (Baker and Macdonald 2000). However, a recent study in Scotland found fox DNA on 34 of 39 (87%) swabs collected from dead or injured lambs, gathered for potential predator/scavenger DNA recovery (George et al. 2024).

The overall impact of foxes on agriculture has been estimated to cost approximately £12M annually (Baker, Harris, and White 2006), most of which is likely to be from both direct and indirect effects from livestock predation. Allowing for inflation, these costs would currently be around £20.1M (Bank of England 2024). However, foxes also predate rabbits (Trout and Tittensor 2008), which are a significant agricultural pest of much higher economic impact, costing around £120M annually (Baker, Harris, and White 2006). Over its lifetime, a fox may be worth £150-900 in increased revenue due to the consumption of rabbits; using the lower estimate translates to an indirect economic benefit to farmers of at least £7M annually (£11.7M with inflation: Bank of England (2024)) from the adult fox population in Britain, thus foxes can be considered as economically neutral to agriculture (Baker, Harris, and White 2006).

B.2. Reservoir of disease

Foxes are known to harbour a number of diseases throughout their range (Veronesi, Deak, and Diakou 2023), including nematodes *Crenasoma vulpis*, *Eucoleus aerophilus* and *Angiostrongylus vasorum* (Taylor et al. 2015). *A. vasorum* is of particular concern due to the increased global reporting of this agent as a cause of disease in dogs. In a 2012-2013 study of 442 foxes sampled from across Britain, Taylor et al. (2015) reported *A. vasorum* prevalence at 18.3%, which had increased from 7.3% reported from a 2005-2006 survey. The increased prevalence observed in Britain compares with elsewhere in Europe, which has seen *A. vasorum* expand from localised pockets to a patchy but wide distribution across the continent, confirmed by a rising prevalence in foxes (Morgan et al. 2021).

B.3. bTB transmission

Although foxes have a lower disease prevalence of bTB (around 4%) than badgers and some deer species (Delahay et al. 2007; Justus et al. 2024), disease prevalence rates are high compared to other mammal species (see Delahay et al. (2007)). Foxes have been known to shed bacilli in saliva, faeces and urine, even without exhibiting macroscopic lesions (Michelet et al. 2018). Foxes that shed bacilli through multiple routes while presenting non-visible lesions may well be considered “super-shedders” since they are likely to shed many bacilli over a lengthy period (Michelet et al. 2018). However, the relatively small body mass of foxes may put a limit on the number of CFUs that can be excreted at any given time (Justus et al. 2024).

B.4. Reservoir of zoonotic disease

The red fox supports a variety of ectoparasites (fleas, lice and mites), as well as a range of helminths. The nematodes *Toxocara canis* and *Unicinaria stenocephala* are the most common species found in wild foxes (Gorman 2008). These pose a risk to other vulnerable wildlife species in addition to domestic dogs that may come into contact with foxes, their faeces or dens. Of 224 fox scats collected for gastrointestinal parasite analysis in Edinburgh in 2017, *T. canis* prevalence was 14.3% and 13.1% in summer and autumn respectively whilst *U. stenocephala* prevalence was 47.1% and 42.0%. Other potential zoonoses were also found, including *Eucoleus aerophilus* (prevalence 52.1% and 32.7%) and *Taenia* spp. (12.6% and 4.67%) (Gecchele, Pedersen, and Bell 2020). Urbanisation of foxes leads to closer proximity with humans, increasing the risk of cross-species or zoonotic transmission (Hassell et al. 2017).

Foxes can also transmit sarcoptic mange (*Sarcoptes scabiei*), which causes scabies in humans. It is highly genetically variable with numerous strains infecting different species, e.g. *S. scabiei* var. *vulpes* (fox) and highly contagious (Scott et al. 2020). Across all conurbations in Britain, mange in foxes is widespread, although Scott et al. (2020) identified a single cluster of high prevalence (37.1%) in northwest and central England, which exceeded double the mean prevalence overall (15.1%). The authors concluded this mirrored the northward expansion of urban fox distribution and highlights the potential risk foxes pose to public health.

The presence of *Trichinella psuedospiralis* has now been confirmed in this species in England (Learmount et al. 2015). The red fox is also a useful sentinel species with respect to the current absence of *Trichinella spiralis* and *Echinococcus multilocularis* in Great Britain (Learmount et al. 2012). The red fox is also a potential reservoir of rabies should the disease arrive in the UK (Smith and Wilkinson 2003).

C.4. Competition with or predation of wildlife (including protected or conservation species)

Although fox predation has been implicated as a contributory factor to the declines of birds such as corncrake, grey partridge, black grouse and capercaillie, other factors, such as habitat loss and degradation, increased use of pesticides and collisions with deer fencing are likely to be more important (Baker, Harris, and White 2006). However, other studies have suggested that fox predation is likely to have a limiting effect on their prey numbers (Roos et al. 2018). A long-term study in Scotland indicated that capercaillie breeding success was negatively related to the abundance of foxes, crows and raptors combined and that breeding success improved when some foxes were killed (Baines et al. 1995). Curlew are classified as globally Near Threatened on the IUCN Red List of Threatened Species with the UK population (an estimated 19-27% of the global breeding population) declining rapidly (Brown et al. 2015). Various studies have identified key predators, including red foxes; Fletcher et al. (2010) demonstrated that reducing the abundance of foxes and carrion crows on moorland in northern England increased the annual breeding numbers of curlew greater than three-fold. Zielonka et al. (2020) reported that predation accounted for 90.6% (29/32) of failed curlew nests in Norfolk, with foxes being the main predator. They concluded that curlews suffered unsustainably high rates of nest predation primarily attributable to the fox, which have significant impacts on curlew populations.

The predation of hares may also be of conservation concern due to the relatively low numbers in Britain and their conservation status (Vaughan et al. 2003).

D.2. Domestic nuisance

Damage by foxes in urban areas, such as knocking over bins and creating litter on streets, is not generally significant (Macdonald and Doncaster 1985) and could be confused with similar activities by domestic or feral cats. The management of these issues is sensitive because many foxes are fed by local residents and some consider their presence in urban areas not as invasive but as a welcome addition to urban wildlife (CIEH 2013).

D.3. Predation of domestic pets

There have been several stories in the media about attacks by urban foxes on children and pets. In Greater London, between 2016 and 2018, a spate of 32 mutilated and dead cats led to media speculation of a human serial cat killer pursuing domestic cats. However, DNA testing, computed tomography imaging and postmortem examination ruled out human intervention, but did find a significant association between cat carcass mutilation and the presence of fox DNA. The findings supported the theory that the cause of mutilation was postmortem scavenging by foxes; where the cause of death could be established (26/32), 10 were attributable to fox predation (Hull et al. 2022).

**Chiroptera**

Daubenton's bat - *Myotis daubentonii*
Natterer's bat - *Myotis nattereri*
Serotine bat - *Eptesicus serotinus*
Common pipistrelle bat - *Pipistrellus pipistrellus*
Soprano pipistrelle bat - *Pipistrellus pygmaeus*
Brown long-eared bat - *Plecotus auritus*

Introduction

Bats are small flying mammals of the order Chiroptera, which are mainly nocturnal (Racey 2008). Only two families of bat are present in the UK (Rhinolophidae and the Vespertilionidae), which is comprised of 18 species, 17 of which are known to be breeding here (www.bats.org.uk). They have varying distributions and populations but all UK bats and their roosts are protected by law, including the Wildlife and Countryside Act, 1981 and a variety of other protective legislation such as EU Habitats Directive (www.bats.org.uk). Of the bat species recorded as causing conflict, Mathews et al. (2018) recent population estimates in Britain are as follows: 1,030,000 Daubenton's bats (682,000 in England, 235,000 in Scotland, 108,000 in Wales, although there are very wide confidence intervals for this species); 414,000 Natterer's bats (based on mixed habitat estimates) or 973,000 (based on woodland estimates), mostly located in England; 136,000 serotine bats (117,000 in England, 18,700 in Wales); 3,040,000 common pipistrelle bats (1,870,000 in England, 875,00 in Scotland, 297,000 in Wales); 4,670,000 soprano pipistrelle bats (2,980,000 in England, 1,210,000 in Scotland, 478,000 in Wales); 934,000 brown long-eared bats (607,000 in England, 230,000 in Scotland, 96,600 in Wales).

A.8. Damage to property - Natterer's

Bats frequently roost in historic churches, with 60% of pre-16th century churches in England estimated to contain bat roosts, comprised of at least 10 species of bat (Marnell and Presetnik 2010). Whilst the presence of bats often causes no conflict, the deposition of bat droppings and urine can cause damage to the building itself and items of cultural significance (Zeale et al. 2016). In extreme cases, large quantities of droppings can restrict the use of the church for worship or community purposes; as bats and their roosts are protected by law, there are significant challenges balancing conservation and cultural needs (Zeale et al. 2016). Natterer's bats often roost in historic churches, with maternity colonies comprising over 100 bats causing acute problems between spring and autumn (Zeale et al. 2016).

A.10. Conflicts with construction, infrastructure and development - serotine, Natterer's, common pipistrelle, soprano pipistrelle, brown long-eared

Buildings often provide excellent roosting sites for bats, especially during hibernation periods (Racey 2008). As a result of the protection that bats are afforded, conflicts can arise where bat populations are present at sites where development is proposed. Serotine bats, Natterer's bats, the two pipistrelle species and brown-long eared bats are considered to roost particularly frequently in buildings (e.g. (Altringham 2003); this may make them more vulnerable to deliberate disruption and bring them into direct conflict with human activities (e.g. development, demolition or maintenance). Developments such as wind farms built in areas where bats forage can potentially be detrimental to bat populations, with large numbers of fatalities from barotraumas reported (Baerwald et al. 2008). Existing buildings and structures can also provide ideal habitats for bats, which may pose problems for any further construction work (Fenn 2002). However, the presence of bats or their roosts in buildings does not mean that building work, repairs or timber treatments cannot go ahead and advice needs to be sought from local statutory nature conservation organisations to deal with individual cases and licensing requirements (Natural England 2022a). The timing and methods of carrying out the work may need to be adapted to minimise bat disturbance (planning-applications.co.uk). In England, derogation from strict protection is allowed under licence, provided activities are undertaken to mitigate any potential negative effects on bat numbers although the post-development effectiveness of such mitigation is not necessarily monitored effectively (Stone, Jones, and Harris 2013). In 2023, the number of European Protected Species mitigation licences issued for bats, often in association with development activity, was 1,142 (Natural England 2023).

B.2. Reservoir of disease - Daubenton's

Since first being identified in 2006, White-nose syndrome (WNS) has spread rapidly with detection in 39 US states and seven Canadian provinces (Hoyt, Kilpatrick, and Langwig 2021). Counts of five bat species from over 200 sites across 27 US states and two Canadian provinces, estimated declines of winter counts in bat colonies of 36%, 79% and over 90% for three of the species (Cheng et al. 2021). In

July 2013 the fungus was isolated from a Daubenton's bat in the UK and from several environmental samples, although surveillance work indicated that although the fungus is present in the UK, it is not causing mortality (Barlow et al. 2015).

B.4. Reservoir of zoonotic disease - Daubenton's, serotine

European bat lyssavirus (EBLV), which causes rabies, is transmissible to humans through bites (Whitby et al. 2010). Bats are generally docile and will avoid contact with humans, hence the main risk is for those people who handle bats. Two species of particular concern as reservoirs for EBLV in the UK are the Daubenton's bat, which is widely distributed across the UK and the serotine bat, which is confined to southern England (Racey, Hutson, and Lina 2013). Passive bat rabies surveillance in the UK initially identified the presence of EBLV-2 only, found in Daubenton's bats (Schatz et al. 2013); in 2018, EBLV-1 was detected in a single serotine bat with this figure rising to 34 cases by May 2024 (Golding et al. 2024). These authors concluded that EBLV-1 is potentially established in the UK, as is the case in other European countries, such as the Netherlands where 20.6% (251 of 1,219) of bats tested during passive surveillance were found to be positive for the virus (Schatz et al. 2013).

D.4. Domestic nuisance & hygiene - serotine, Natterer's

In addition to net economic loss arising from roosts in buildings, which prevent or delay construction or demolition work or building damage (see above), bats roosting in buildings may represent further nuisance. They are not known to damage property by gnawing wood or cables but their droppings may cause unpleasant odours and will be found where bats roost (Zeale et al. 2016).

**Eulipotyphla**

Hedgehog - *Erinaceus europaeus*

Introduction

The hedgehog is an iconic native species with a widespread distribution in areas of suitable habitat (Churchfield 2008). They are found in most areas although they are increasingly associated with urban areas; hedgehogs are often observed in gardens and amenity grasslands (Mathews et al. 2018) and are found at higher densities in areas with amenity grassland compared with pasture (Young et al. 2006; Parrott, Etherington, and Dendy 2014). Hedgehogs have also been recorded on many of the UK islands, to which they were most likely introduced; some efforts have been made to eradicate them in the Hebrides, where they caused severe declines in the breeding success of waders due to predation on eggs (Jackson, Fuller, and Campbell 2004). Despite this, the hedgehog remains popular and is often fed by people in suburban areas, where they have been reported in 70% of gardens (Ansell, Baker, and Harris 2001). The hedgehog now has partial protection under Schedule 6 of the Wildlife and Countryside Act 1981 as well as the Wild Mammals (Protection) Act 1996 so that trapping and invasive actions require a licence (Churchfield 2008). Recent population estimates were around 522,000 free-living hedgehogs in Britain: 320,000 in England, 145,000 in Scotland, 56,800 in Wales (Mathews et al. 2018).

A.5. Predation of game species

The British hedgehog has undergone persecution for its role in depleting the eggs of game birds in the past, despite the numbers of eggs affected considered to be insignificant when compared with the numbers taken by foxes and crows (Churchfield 2008).

B.2. Reservoir of disease

European hedgehogs are susceptible to foot-and-mouth disease (FMD) virus, which can have devastating impacts on susceptible livestock (Riley and Chomel 2005). During an outbreak of FMD in Britain in 1946, a number of hedgehogs found on or near to infected premises were found to be affected by the disease; although transmission between hedgehogs and cattle and vice versa had been observed experimentally, the definitive transmission risk that hedgehogs posed under natural conditions could not be fully concluded (McLauchlan and Henderson 1947). No recent information identified.

B.4. Reservoir of zoonotic disease

*Salmonella* is the most common zoonotic disease associated with hedgehogs and is widespread in British hedgehogs (Keymer, Gibson, and Reynolds 1991). Lawson et al. (2018) investigated various tissue samples (liver and small intestinal contents) from 170 wild hedgehogs and faecal samples from 208 wild hedgehogs across Great Britain. Classical microbiological methods and Gram staining coupled with the determination of biochemical characteristics and histopathological examination were used to identify bacterial isolates and description of lesions. *S. enteritidis* multi-locus sequence-type (ST)183 was isolated from 46/170 (27%) tissue samples and from 6/208 (3%) faecal samples. The types of *Salmonella* found in hedgehogs were also investigated and compared with those found in humans. The results showed that infections in both species might have originated from a common population, with the authors concluding that hedgehogs might serve as a reservoir host and source of salmonellas for human (Lawson et al. 2018). Hedgehogs also play a major role in the transmission of *Salmonella* Tilene, a rarely encountered serotype of humans (Riley and Chomel 2005).

Zoonotic fungi that cause ringworm (Dermatophytes) have also been identified in hedgehogs in various locations of the UK, with prevalence estimated to be higher in urban areas (17%) than rural areas (9%); Morris and English (1969) concluded that the higher disease prevalence in urban areas was likely due to the higher density of hedgehogs in these areas. These studies are now outdated and do not provide any useful indication of how frequent or widespread such occurrences were. No recent information identified.

C.5. Predation of important ground-nesting birds and island ecosystem damage

The main areas where hedgehogs have a significant impact on ground-nesting birds are on the Scottish islands, where they have been introduced and predate eggs of shorebirds and threaten local extinction (Jackson and Green 2000). Calladine et al. (2017) found that where hedgehogs were rare on islands in the Outer Hebrides, clutch survival rates of five species of waders (dunlin, lapwing, redshank, snipe and ringer plover) were higher than where hedgehogs were relatively more abundant. From camera evidence, hedgehogs were identified as the most frequent nest predator, although other predators were also likely influencing the population sizes of breeding waders.

European mole - *Talpa europaea*

Introduction

The mole is a widespread native British mammal, which is highly adaptable and found in most habitats where invertebrate prey is present and the soil is sufficiently deep to allow tunnel construction (Mathews et al. 2018). The mole thrives in pasture and on arable land and has been widely persecuted as a pest of agricultural and amenity land as a result. There are an estimated 41,400,000 moles in Britain: 24,300,000 in England, 12,200,000 in Scotland, 4,900,000 in Wales (Mathews et al. 2018). Moles are not found on many of the British islands (Gorman and Stone 1990).

A.2. Damage to agricultural interests

A questionnaire study found that 64% of farmers considered moles to be pests, although damage believed to be caused by moles was slight with silage pollution by soil reported as the biggest problem (Atkinson, Macdonald, and Johnson 1994). The exposed soil excavated by moles also gives weeds species opportunities to colonise where there is no competition from grass (Quy and Poole 2004). Molehills are also damaging to agricultural machinery by blunting blades and catching stones brought up by moles (Quy and Poole 2004). As they tunnel, moles can cause damage to young plants and seedlings by uprooting them, either killing them directly or leaving the plant roots exposed to frost (Guedon 1998). Furthermore, tunnelling can cause soils to dry out resulting in crops that are liable to wilting. Conversely, their tunnelling activities can cause flooding because of disturbance to drainage systems (Quy and Poole 2004). Atkinson, Macdonald, and Johnson (1994) calculated the farm level costs to range between £126 and £800, with pasture and mixed farms suffering the greatest levels of damage and localised problems having the potential to be very significant. The cost of mole damage and measures taken to minimise these impacts in the UK were estimated to be less than £5M per year (Quy and Poole 2004). In a nationwide survey of farms in 2007, Baker et al. (2016) found no evidence that mole activity had increased since the 1992 survey carried out by Atkinson, Macdonald, and Johnson (1994). Based on 2024 prices, mole damage would be estimated to be around £6.8M (Bank of England 2024).

A.9. Damage to amenities and domestic grassland

The mole is also considered to damage domestic and amenity grassland where aesthetic or high-value amenities are important, e.g. racecourses, golf courses (Baker et al. 2016). Damage reported on amenities and gardens is often merely aesthetic, but mole activity on race courses and sports fields may present a risk of injury to people and animals; these types of conflict have not been fully economically assessed and are mostly considered to be an inconvenience (Quy and Poole 2004). It has been calculated that moles account for 25% of a typical golf course pest control bill, therefore approximately £2.7M (Williams et al. 2010). Allowing for inflation, these costs would be around £4.0M currently (Bank of England 2024).

B.5. Threat to livestock where ground is undermined

Tunnelling by moles and the resulting molehills can pose a threat of serious injuries to horses and riders on racecourses, either by tripping or falling into collapsed holes (Quy and Poole 2004). At the speeds racehorses travel, such injuries could be fatal.

Common shrew (*Sorex araneus*)
Introduction

The common shrew is found in most terrestrial habitats with low vegetation cover, although it is most abundant in thick grass, bushy scrub, hedgerows and broadleaved woodland (Mathews et al. 2018). It requires habitats with high invertebrate abundance due to its high energy requirements and may be negatively affected by changes in agricultural and/or pesticide use which reduce its prey availability (Mathews et al. 2018). It is widespread throughout Britain with an estimated population of 21,100,000: 11,000,000 in England, 7,690,000 in Scotland, 2,330,000 in Wales (Mathews et al. 2018).

B.4. Reservoir of zoonotic disease

Coronoviruses (CoV) are important pathogens affecting human and animal health, with the recent emergence of several serious disease outbreaks, including severe acute respiratory syndrome (SARS), Middle East respiratory syndrome (MERS) and global Covid-19, highlighting the risk animals can pose for cross-species transmission. Tsoleridis et al. (2016) detected novel alphacoronoviruses in a number of rodent and shrew species sampled from across the East Midlands region of the UK between 2008 and 2015 with 33.3% (1/3) of common shrews and 4.1% (5/123) brown rats testing PCR positive for CoVs. Many animal and human CoV spillover events can be traced back to a bat reservoir (Corman et al. 2014; Yang et al. 2015), although genetically distinct CoVs have been found recently in rodents in China (Wang et al. 2015), suggesting rodents may harbour unidentified CoVs with zoonotic potential (Tsoleridis et al. 2016). Phylogenetic analysis showed that the novel rodent/shrew CoVs found in the UK were mostly related to the rodent Lucheng *Rn* CoV, with Tsoleridis et al. (2016) concluding that it was highly unlikely that the novel rodent/shrew CoVs were a recent and direct epizootic event from bats. Whilst the prevalence of CoVs in commons shrews appears high, the sample size tested was very low; however, further investigation into potential cross-species transmission is warranted given these novel CoVs are phylogenetically separate from bats.

Greater white-toothed shrew (*Crocidura russula*)
<u>Introduction</u>

The greater white-toothed shrew is native to Europe and islands such as Guernsey, Alderney and Sark. It was discovered in Ireland in 2007 and more recently in Sunderland, northeast England, in 2021 (Bond et al. 2022). Greater white-toothed shrews are found in hedgerows, grassland, woodland and cultivated areas, with a preference for human settlements and farm buildings during colder months (Mammal Society 2024a). Estimated population sizes are currently unknown.

<u>B.4. Reservoir of zoonotic disease</u>

Pathogenic species of *Leptospira* cause leptospirosis, a bacterial zoonotic disease with a global distribution affecting over one million people annually (Costa et al. 2015). Of 18 greater white-toothed shrews trapped in Ireland, three (16.7%) were culture positive for an unidentified serovar of *Leptospira*, later indicated as being genetically related to pathogenic *L. alstonii* (Nally et al. 2016). These authors concluded that greater white-toothed shrews were acting as bridge vectors for a novel zoonotic pathogen. Although the distribution and abundance of greater white-toothed shrews in Britain is uncertain, they could potentially pose a risk to public health and further monitoring is warranted.

A number of viruses, which may significantly impact human and animal health, have been detected in greater white-toothed shrews in Europe (Haring et al. 2024). Between 2002 and 2021, greater white-toothed shrews sampled across Germany were found with co-infections of three viruses; a previously identified orthoparamyxovirus (Denwin virus) and orthonairovirus (Erve virus), plus a novel shrew-human hepatitis E virus (shrewHEV) (Haring et al. 2024). These authors reported that 9 of 16 (56%) shrews were positive for Denwin virus (which has also been detected in greater white-toothed shrews in Belgium: Horemans et al. (2023)) indicating a high prevalence and wide geographical distribution (Haring et al. 2024). The zoonotic potential of these viruses is unknown although their phylogenetic proximity to other zoonotic viruses within the same families (e.g. orthopoaramyxoviruses: Nipah virus, measles virus, Hendra virus and Langya virus; hepevirus: *Paslahepevirus balayani*, which causes human hepatitis E) warrants further investigation (Haring et al. 2024).

<u>C.2. Threat to biodiversity</u>

Initial data from Ireland showed that greater white-toothed shrews may limit the abundance and distribution of the pygmy shrew (*Sorex minutus*), Ireland's only shrew species (McDevitt et al. 2014). Upon its discovery in Sunderland in 2021, there was concern for its potential impact on other small mammal species, such as the pygmy shrew, although the small mammal community in Great Britain is more complex than Ireland (Bond et al. 2022). These authors concluded from photographic evidence that greater white-toothed shrews have been in northeast England since at least 2015 and potentially longer; the fact that they have gone for so long without being discovered reflects how small mammals are relatively under-recorded (Crawley et al. 2020). As pygmy shrews are probably the most under-recorded mammal in northeast England, baseline data of distribution and abundance must be obtained to ascertain what impact greater white-toothed shrews might have in the future (Bond et al. 2022).

### Lagomorpha

Rabbit – *Oryctolagus cuniculus*
<u>Introduction</u>

The European rabbit is an introduced species brought to England by the Normans and kept in managed warrens. Its populations showed large increases from c.1750 when agricultural practices began to provide more suitable habitats and predator control, to protect game interests, meant that the predation pressure was reduced (Cowan 2008). Rabbits are known to cause extensive damage, particularly within the agricultural industry and are also the reservoir of a variety of significant diseases and are therefore a large focus of conflict with human interests (Cowan 2008). The release of myxomatosis into the UK population in 1953 caused a substantial decline in numbers (Ross et al. 1986); although impacts are severe on a local level (Petrovan, Ward, and Wheeler 2011), its effect is poorly understood on a national scale (Mathews et al. 2018). Although rabbit populations have declined in recent years, mostly due to infection with myxomatosis and rabbit haemorrhagic disease virus RHDV1 and RHDV2 (Bell, Davis, et al. 2019), they remain widespread throughout the country, with an estimated 36,000,000 rabbits in Britain: 21,300,000 in England, 11,800,000 in Scotland, 2,910,000 in Wales (Mathews et al. 2018).

A.1. Damage to forestry

The European rabbit causes widespread damage to forestry. Rabbits are a particular issue for young trees where bark damage can result in tree death, although most damage is caused by browsing (Williams et al. 2010). Costs of controlling rabbits (i.e. fencing) was estimated to be around £8M annually in Britain (£3.9M in Scotland, £3.3M in England, £800K in Wales) (Williams et al. 2010). Although rabbit populations have declined in recent years, it is unclear if costs to control rabbits have also declined, therefore Eschen et al. (2023) revised the figures according to inflation, estimating annual costs of controlling rabbits to be around £11.4M in Britain (£5.6M in Scotland, £4.7M in England, £1.1M in Wales).

Although Williams et al. (2010) identified there is limited data to indicate the reduction in value of timber due to rabbit damage, they estimated total annual damage costs by rabbits of around £62.0M (£34.0M in Scotland, £21.1M in England, £7.0M in Wales). Due to the decline in rabbit populations and assumed reduction in rabbit grazing, Eschen et al. (2023) predicted damage costs by rabbits to be lower at around £31.4M annually in Britain (£21.8M in Scotland, £8.2M in England, £1.4M in Wales). When costs of control and damage were combined, rabbits were estimated to cost around £42.8M.

A.2. Damage to agricultural interests

Rabbits cause widespread damage to a range of agricultural crops (Cowan 2008), both at the growing stage and end marketable product, with winter wheat, barley and oats being the most vulnerable crops (Williams et al. 2010). These authors estimated the annual cost of damage to agricultural interests caused by rabbits in Britain, based on an estimated population size of 40,000,000 (Smith, Garthwaite, and Prickett 2006) to be approximately £183.3M (£111.4M in England, £15.0M in Wales and £57.0M in Scotland) (Williams et al. 2010). Due to the decline in rabbit numbers, Eschen et al. (2023) estimated annual costs to agriculture in Britain of around £119.8M (£73.0M in England, £35.1M in Scotland, £11.6M in Wales).

A.9. Damage to amenities and domestic grassland

Rabbits are frequently reported as pests on golf courses, accounting for the majority (50%) of the annual costs golf courses spend on pest management (Williams et al. 2010). These authors estimated that the annual cost of rabbits to golf courses in Britain was £5.3M (£3.8M in England, £1.1M in Scotland, £350K in Wales). Due to the decline in rabbit numbers, Eschen et al. (2023) estimated the costs to be around £3.4M (£2.5M in England, £700K in Scotland, £300K in Wales).

A.12. Conflicts with transport

Williams et al. (2010) reported that the Highways Agency spent £615K on rabbits across the whole network of trunk roads and motorways in England, due to direct control by contractors, vegetation management (e.g. removal of harbouring shrubs), litigation charges and vegetation protection (i.e. cost of installing and removing plastic shelter for trees, etc.). These costs were extrapolated for Scotland and Wales, £186K and £75K respectively. Eschen et al. (2023) revised these figure to reflect the reduction in rabbit numbers, with an estimated cost of £550K in England, £200K in Scotland, £100K in Wales.

B.2. Reservoir of disease

Rabbits have recently become increasingly recognised as a reservoir of livestock disease, in particular paratuberculosis, which is a widespread and difficult to control (Judge et al. 2007; Shaughnessy et al.

2013). Paratuberculosis has also been linked to Crohn's disease in humans, with rabbits identified as the key species in which paratuberculosis is able to persist and be transmitted to the wider host community (Judge et al. 2007). Shaughnessy et al. (2013) sampled 281 rabbits from 13 beef farms in east Scotland that were having difficulties controlling paratuberculosis in their cattle herds; 67 (24.8%) rabbits were culture positive for *Mycobacterium* species, confirmed as MAP by PCR. The authors concluded there was an association between the persistence of paratuberculosis in livestock despite the implementation of disease control strategies and MAP-infected sympatric wild rabbit populations.

Rabbit populations have declined significantly in recent years due to RHDV2, which has spread rapidly worldwide (Rouco et al. 2019). It has caused crashes in both domestic and wild rabbit populations (e.g. in France: (Le Gall-Recule et al. 2013) and has likely replaced the former RHDV1 variant in the wild (Rouco et al. 2019). RDHV2 has also been found in European brown hares in Spain, Italy (Velarde et al. 2017) and the UK, with cases in Essex and Dorset (Bell, Davis, et al. 2019), although in limited numbers in the UK and possibly representing a spill-over event from rabbits (Rocchi et al. 2019).

Adiaspiromycosis is a mycotic infection caused by thermally dimorphic fungi classified as *Emmonsia parva* and *E. crescens*. *E. crescens* infection is typically associated with wild rodents but has been diagnosed in a broad range of wildlife species in Great Britain, with mustelids considered to be particularly susceptible (Malatesta et al. 2014), and also domestic species, including canine (al-Doory, Vice, and Mainster 1971) and equine (Pusterla et al. 2002). Borman et al. (2009) reported that 27.8% (n=27/94) of animals found dead in Great Britain (mostly from southwest England) and submitted to the Wildlife Veterinary Investigation Centre, Truro between 2003 and 2005 were positive for *E. crescens* infection on either microscopy or histopathology, although none of these were rabbits. A wild rabbit submitted for testing at the University of Cambridge was found to have a severe case of adiaspiromycosis; although PCR amplification failed to identify *Emmonsia*-specific DNA, histopathological findings suggested the disease was most likely caused by *E. crescens* (Hughes and Borman 2018). The risk posed by rabbits is likely to be low.

Of 87 wild rabbits shot in Cambridgeshire in 2016-2017, *Eimeria stedae*, which is one of a number of parasites that causes white-spotted liver lesions, was detected in 21 (24%) rabbits (Bochynska et al. 2022). These authors found that gross lesions consistent with white-spotted liver were common in wild rabbits from this area; with an estimated 900,000 pet rabbits in the UK, they concluded there was a risk that wild rabbits may act as a reservoir of infection for this disease via contaminated vegetation.

<u>B.4. Reservoir of zoonotic disease</u>

Wild rabbits have been linked to VTEC *Escherichia coli* O157 which can be lethal in humans. Rabbits have been identified as possible vectors of the disease for transmission to humans (Bailey et al. 2002), particularly due to their close proximity with infected cattle excreting *E. coli* O157 (Scaife et al. 2006). However, there is no evidence as to the extent of human infection for which the rabbit is responsible.

Cryptosporidiosis is a faeco-orally transmitted diarrhoeal disease, caused by species of the protozoan genus *Cryptosporidium*. *C. parvum* and *C. hominis* are the most common species reported in humans worldwide and have been responsible for the majority of waterborne outbreaks to date, with the exception of a waterborne outbreak in Northamptonshire, UK, in 2008 caused by *C. cuniculus* from rabbits (Chalmers et al. 2009; Puleston et al. 2014). This highlighted the importance of wildlife in the dissemination of *Cryptosporidium* to drinking water sources and the associated risk to public health.

*Calodium hepaticum* is a globally distributed nematode that causes white-spotted liver lesions. It is transmissible to humans and can cause severe hepatic infections resulting in death if misdiagnosed (Fuehrer, Igel, and Auer 2011) although it is rarely reported in Great Britian (Barlow and Mullineaux 2018). Bochynska et al. (2022) reported the presence of *C. hepaticum* in seven of 87 (8%) wild rabbits shot in Cambridgeshire (see above); although *E. stedae* was found to cause the majority of cases of white-spotted liver in the sampled rabbits, there is still a risk that wild rabbits may transmit *C. hepaticum* to pet rabbits or humans.

<u>C.6. Sustaining predators harmful to important conservation species</u>

In the UK, rabbits are not only an important part of the prey base for native predators but also form an important prey species for non-native mink (Oliver, Luque-Larena, and Lambin 2009). Where rabbits are sustaining more abundant mink populations, particularly during periods when other prey are in short supply, conservation species such as water voles are further disadvantaged by the presence of consequently greater numbers of predators (Halliwell and Macdonald 1996; Oliver, Luque-Larena, and Lambin 2009).

D.1. Public nuisance

Rabbits graze on the shoots of herbaceous plants and can dig holes and scrapes in lawns and flower beds (RHS 2024).

Brown hare - *Lepus europaeus*
Introduction

The brown hare is found throughout Britain having been introduced probably in the Iron age (Cowan 2008). It is most common in lowland arable areas and prefers cultivated areas, although is not restricted to them and will feed on pastures (Cowan 2008). The brown hare is valued as a game animal and densities tend to be highest for shoots where fox control is in operation (Cowan 2008). Although hares have minimal legal protection due to them being a non-native game species, they are now included in the UK Biodiversity Action Plan due to its suspected decline. There are an estimated 579,000 hares in Britain: 454,000 in England, 87,700 in Scotland, 37,300 in Wales (Mathews et al. 2018).

A.1. Damage to forestry

Brown hares can damage trees by bark stripping, which can potentially lead to fungal infections and branch/tree death or scarring of wood, which will reduce its value (Gurnell and Hare 2008). They can also damage plantations of young trees. Reports from New Zealand state that hares, even in low numbers, can cause severe damage to young trees, cuttings, vegetables and plant nurseries with a single pair capable of destroying up to 100 trees in one night (hbrc.govt.nz). However, in Thetford Forest, eastern England, Zini, Wäber, and Dolman (2022) found that the intensity of Scots Pine leader damage was not affected by the presence of hares as predicted by multi-model inference, with negligible impact compared to deer.

A.2. Damage to agricultural interests

Brown hares can cause significant damage to crops such as sugar beet, cereals, horticultural crops and shrubs (Cowan 2008). Due to the relatively lowland range of the brown hare, compared to that of the mountain hare, it is likely that the brown hare causes greater levels of damage in arable areas, but may also be confused with damage caused by rabbits (pers. comm., D.P. Cowan).

A.14. Damage to property as a result of poaching

Poaching of wildlife is illegal. Poachers often use inhumane methods to kill their targets and can cause damage to property (kent.police.uk). Some farmers and gamekeepers accuse poachers of destroying crops, damaging vehicles and fences and releasing livestock onto roads (BBC 2023b)

B.2. Reservoir of disease

(See rabbit for information on RHDV2). RDHV2 has been found in European brown hares in Spain, Italy (Velarde et al. 2017) and the UK, with cases in Essex and Dorset (Bell, Davis, et al. 2019), although in limited numbers in the UK and possibly representing a spill-over event from rabbits (Rocchi et al. 2019).

B.4. Reservoir of zoonotic disease

(See rabbit for information on *Calodium hepaticum*). A case of *C. hepaticum* was recorded in a brown hare in Great Britian, which presented with severe extensive granulomatous hepatitis. The condition is considered a rare occurrence in brown hares but may be misdiagnosed grossly as hepatic coccidiosis (Barlow and Mullineaux 2018). Given the infrequent reporting of this zoonotic agent in hares, they likely represent a very low risk to public health.

In Europe, hares have been found to act as reservoirs of *Borrelia burgdorferi* in the absence of other small mammal reservoirs (e.g. voles, wood mice, yellow-necked mice, hedgehogs) (Sala and De Faveri 2016).

Mountain hare - *Lepus timidus*

Introduction

The mountain hare is a native species to Britain that has been widely introduced beyond its original range (Battersby 2005). It is widespread in Scotland, found primarily on heather moorland at an altitude of 300-900m (Flux 2009). In England, its distribution is currently restricted to the Peak District, where it is found primarily in areas of common heather and cotton grass (Harris and Yalden 2008). The mountain hare is listed in Annex V of the EC habitats directive (1992), as a species of community interest where taking from the wild and exploitation may be subject to management measures (Mammal Society 2024b). There are an estimated 135,000 mountain hares in Britain: 2,500 in England, 132,000 in Scotland (Mathews et al. 2018).

A.1. Damage to forestry

The mountain hare can cause damage to trees by browsing or bark stripping, although bark stripping appears to be relatively rare in the UK, often moving along a row and browsing each tree in turn (Gill 1992). Mountain hares prefer willow and rowan but will also browse most conifers in Britain (Hewson 2009). This can be very detrimental to forestry plantations, especially as younger trees are often browsed preferentially (Gill 1992).

A.2. Damage to agricultural interests

Mountain hares are known to feed on crops, although the extent to which they are responsible for crop losses is not well documented (Macdonald et al. 2000). They consume crops such as oilseed rape, turnip, grasses and cereals (Cowan 2008). They will also consume more expensive market garden crops, which will be significant to people on a more individual level, although the brown hare is more likely to be responsible for damage in more lowland areas (Macdonald et al. 2000).

A.14. Damage to property as a result of poaching

(See brown hare for information on damage to property as a result of poaching). Due to the distribution of mountain hares, damage to property may be significant for individual farmers and landowners but is likely localised.

B.4. Reservoir of zoonotic disease

Louping ill virus has been shown to spread amongst ticks co-feeding in close proximity on mountain hares (Jones et al. 1997). Louping ill is a virus which can also cause mortality in red grouse and sheep and is therefore undesirable to game keepers and farmers (gwct.org.uk). However, there is no clear evidence of the involvement of mountain hares in the spread of louping ill to red grouse and consequent monetary losses for gamekeepers.

**Perissodactyla**

Exmoor/Dartmoor pony - *Equus ferus caballus*

Introduction

Herds of wild horses roam free in the UK, mostly in the New Forest but also on Exmoor and Dartmoor. There are a number of free-ranging populations of relatively unimproved stock that are managed to some degree, and are therefore not truly feral and self-sustaining (Putman 2008). Due to their free-roaming status, these animals are not contained and are therefore at risk of road traffic accidents (RTAs) with consequent damage to vehicles and associated health risks for both humans and horses. There are an estimated 350 free-roaming Exmoor ponies (www.visit-exmoor.co.uk) and 1,500 Dartmoor ponies (www.visitdartmoor.co.uk).

A.3. Damage to vehicles (RTAs)

No information for economic costs of RTAs caused by feral ponies found. Several incidents of ponies damaging parked vehicles has been reported, including a small herd of Dartmoor ponies causing £1,200 damage to a parked car by licking the paintwork. Dartmoor Search and Rescue Team reported further incidents of vehicles being chewed (www.horseandhound.co.uk).

B.1. Health risks from RTAs

The number of ponies killed on roads in the New Forest varies annually, with 34 reported in 2022 and 27 in 2023 (https://www.newforestnpa.gov.uk). Over a 10-day period in September 2023, four ponies died on a one-mile stretch of the B3054, suggesting their occurrence is likely localised. Where ponies have been found with severe wounds, broken limbs or dead, road traffic has been blamed although not all of the incidents were verified (Green 2013). No information was identified on the risks posed to human health.

**Rodentia**

Red squirrel - *Sciurus vulgaris*

Introduction

The red squirrel is native to Great Britain, with just over 80% of the remaining population found in Scotland (Mathews et al. 2018). They occur in both broadleaved and conifer woodland, as well as mixed forests, parks and gardens (Harris and Yalden 2008). They eat a wide range of food, mainly tree seeds and fruits, but also tree shoots, buds, flowers, berries and lichen (Gurnell et al. 2015). Forest habitat management in key stronghold areas is often focussed on conserving red squirrels, with Sitka spruce dominated plantations containing a larch and pine element offering the best long-term opportunity to sustain populations, where the competitive advantage of grey squirrels is reduced (Shuttleworth et al. 2012). Since at least the mid-1940's, red squirrels have experienced a sustained decline in numbers and distribution, while the non-native grey squirrel has significantly expanded, colonising the majority of England, Wales and parts of Scotland and Ireland (Gurnell et al. 2004). Alongside the expansion of grey squirrels, outbreaks of squirrelpox virus (SQPV) are thought to have caused the extinction of local populations of red squirrels (Chantrey et al. 2014). There are an estimated 287,000 red squirrels in Britain: 38,900 in England, 239,000 in Scotland, 9,190 in Wales (Mathews et al. 2018).

B.2. Reservoir of disease

Fatal exudative dermatitis (FED) is a significant cause of death of red squirrels in Jersey, where it is associated with a virulent clone of *Staphylococcus aureus* ST49. *S. aureus* ST49 has been found in other hosts such as small mammals, pigs and humans (Fountain et al. 2021). These authors tested red squirrels from several locations across Britain that were both healthy (Jersey, Isle of Arran, Brownsea Island) and displaying signs of FED (Jersey, Isle of Wight). *S. aureus* was frequently carried by red squirrels from the Isle of Arran, predominantly strains associated with small ruminants. From Brownsea, *S. aureus* was less frequent involving strains associated with birds, small ruminants and humans. Fountain et al. (2021) concluded that *S. aureus* carriage in red squirrels largely reflected frequent but facile acquisitions of strains carried by other hosts (spillover), possibly including *S. aureus* ST188, acquired from humans.

B.4. Reservoir of zoonotic disease

Blackett et al. (2018) reported that 116 of 337 (34.4%) red squirrels examined post-mortem in Jersey between 2007 and 2014 had died due to disease. The most prevalent disease was hepatic capillariasis (31.5%, 106/337), which is caused by *Capillaria hepatica* (syn. *Calodium hepaticum*). It commonly infects wild rodents and less frequently, other species, including humans (Fuehrer, Igel, and Auer 2011), but it is rarely reported in Great Britian (Barlow and Mullineaux 2018). Toxoplasmosis was also reported in 7/337 (2.1%) red squirrels, which was lower than detected by Simpson et al. (2013) in red squirrels on the Isle of Wight (15.6%, 12/77). The possible range of red squirrels into suburban areas and potential contact with the large domestic cat populations found in Jersey and on the Isle of Wight may increase the risk of red squirrels acting as intermediate hosts for *Toxoplasma gondii*.

The presence of leprosy in red squirrels, initially identified as *Mycobacterium lepromatosis*, was first described in Scotland in 2014 (Meredith et al. 2014). An additional causative agent, *M. leprae*, was discovered in red squirrels from Brownsea Island (Avanzi et al. 2016) and has since been found elsewhere in the UK (e.g. Isle of Arran; Schilling et al. (2019)). It is currently unknown if *M. lepromatosis* has ever caused human disease in Europe, but the *M. leprae* strain is highly related to strains causing leprosy in medieval Europe (Avanzi et al. 2016). Although red squirrels are a potential reservoir for leprosy in Britain (Avanzi et al. 2016), their distribution is limited and incidence appears to be low; no cases of human leprosy have been reported since the 1950's (Fulton et al. 2016), therefore red squirrels pose minimal risk to human health (Schilling et al. 2019).

Grey squirrel - *Sciurus carolinensis*
Introduction

The grey squirrel was introduced from the USA in 1876, to approximately 30 sites within England and Wales, from where its distribution expanded rapidly and extensively into a broad range of habitats, although particularly in mature broadleaved forests (Gurnell and Hare 2008). They are also found in mixed and coniferous forests, urban and suburban areas, and parks and gardens, capable of surviving in highly fragmented, functionally isolated landscapes (Stevenson-Holt et al. 2014). The expansion of the grey squirrel population has been associated with the decline or extinction of the native red squirrel (*Sciurus vulgaris*) across much of its original range in Britain (Gurnell et al. 2004; Chantrey et al. 2014). The grey squirrel also causes damage in buildings (Gurnell and Hare 2008) and has an important role in threatening the development of new woods (Gill, Gurnell, and Trout 1995). There are an estimated 2,700,000 grey squirrels in Britain: 1,940,000 in England, 478,000 in Scotland, 283,000 in Wales (Mathews et al. 2018).

A.1. Damage to forestry

Grey squirrels cause substantial damage to the timber industry through bark stripping that can kill trees, affect their growth, increase their susceptibility to various pathogens and affect timber value (Nichols et al. 2016; Eschen et al. 2023). Trees between 10 and 40 years old are particularly vulnerable, especially those species with thin bark (Williams et al. 2010). Scars and distorted growth caused by the damage to both broadleaved and coniferous trees can reduce the quality and thus value of timber (Gurnell and Hare 2008). In Great Britain, approximately 166,000ha of woodland stands are estimated to have bark stripping damage observable from the ground; the percentage area of woodland stands with bark stripping damage is highest in England, with approximately 11% of stands showing damage, 6% in Wales and 0.3% in Scotland (National Forest Inventory 2020). Williams et al. (2010) estimated the cost of squirrel damage to broadleaved trees at £413K per annum and the total yield loss to forestry estimated at £685K. With the inclusion of estimated control costs the total economic loss to forestry attributable to the grey squirrel was estimated at £6.1M annually. Based on the area of at-risk woodland and the squirrel population in each country, 65% of all costs were incurred in England (£4.0M), 20% in Scotland (£1.2M) and 15% in Wales (£915K). Revised costs by Eschen et al. (2023) were significantly higher, with the direct costs of grey squirrels (including lost timber value, grey squirrel control, restocking costs of trees that have died due to grey squirrel damage but not carbon capture estimates) to the forest sector in Great Britain estimated to cost £28.7M (£23.3M in England, £0.9M in Scotland, £4.5M in Wales).

A.2. Damage to agricultural interests

Some damage of economic importance is caused to market gardens, orchards and arable crops if they are located near favourable grey squirrel habitat, particularly when other food sources are in short supply (Gurnell and Hare 2008). No recent information found.

A.8. Damage to property

Grey squirrels cause problems in homes and buildings including damage to thatch, gnawing electrical cable, water pipes and roof structures they create noise and leave faeces and urine, which are a nuisance to human occupants (Eschen et al. 2023). Grey squirrels are also frequently seen taking food from bird feeders and occasionally take bulbs from gardens and public parks (Gurnell 1987). Control costs typically vary between £162 and £240 among councils or companies offering grey squirrel control (Eschen et al. 2023). Williams et al. (2010) estimated the annual cost of damage by grey squirrels to houses was £5.1M, with £4.7M in England, £118K in Scotland and £311K in Wales. Revised costs estimated the damage done to houses by grey squirrels was £7.3M, with £6.7M in England, £200K in Scotland and £400K in Wales (Eschen et al. 2023)

B.2. Reservoir of disease

Compelling evidence exists that grey squirrels are reservoir hosts of squirrelpox virus (SQPV), which has exacerbated the decline and extinction of the native red squirrel across much of its original range (Sainsbury et al. 2008; Chantrey et al. 2014). Prevalence of SQPV is high in English and Welsh grey squirrel populations; they have a high seroprevalence of antibodies to SQPV (Bruemmer et al. 2010) and show few clinical symptoms of being infected, whereas red squirrels are highly susceptible to the virus (Schuchert et al. 2014). Rates of decline of red squirrels are 17-25 times higher in areas where the SQPV is present in grey squirrels (Rushton et al. 2006). Chantrey et al. (2014) concluded that SQPV-infected grey squirrels potentially initiate outbreaks of squirrelpox disease in red squirrels.

Evidence has emerged of adenovirus infection in both squirrel species (Everest et al. 2009; Wernike et al. 2018). These authors reported that squirrel adenovirus 1(SqAdV-1) was widespread in the Scottish red and grey squirrel populations, highlighting the need for continuous wildlife surveillance; other studies in the UK have observed high mortality in red squirrels (Everest, Holmes, and Shuttleworth 2017) indicating that SqAdV infections have the potential to be a significant mortality factor, whilst grey squirrels have been identified as subclinical reservoirs (Everest, Holmes, and Shuttleworth 2017).

B.4. Reservoir of zoonotic disease

Grey squirrels in the UK have been found to carry relatively high numbers of immature *I. Ricinus*; they are commonly infected with *Borrelia burgdorferi sensu lato* (s.l.) species complex and infected individuals are able to transmit the pathogen to feeding larvae (Millins et al. 2015). In Scotland, grey squirrels have been found to be infected with all genospecies found in questing ticks, most commonly *B. garinii,* a genospecies normally associated with bird hosts (Millins et al. 2015). These authors found that the prevalence of *B. burgdorferi* s.l. in grey squirrels was seasonal reflecting changes in tick activity, indicating that infection in grey squirrels might be relatively short-lived. They concluded that grey squirrels were more likely to be a spillover host than a maintenance host for *B. burgdorferi* s.l., and that removing grey squirrels could result in a reduction in Lyme borreliosis risk given they host immature ticks and can transmit locally circulating strains of *B. burgdorferi* s.l.

Wild rodents have been implicated as a source of zoonotic pathogens, including *Yersinia enterocolitica* (Keesing and Ostfield 2024), which is transmitted via the faecal-oral route. Given their potential transmission risk at the livestock-wildlife interface, they represent a major food-borne public health hazard (Arden et al. 2022). Of 36 grey squirrels sampled from across the UK, only one (2.78%) was confirmed as *Y. entercolitica* positive, which in turn was deemed to be a non-pathogenic biotype (Arden et al. 2022) therefore posing minimal risk to public health.

B.6. Threat of branches falling

Grey squirrels cause considerable damage to trees by stripping the bark, which prevents the flow of water and food leading to significant stress, weakening and possible death of the tree (Gurnell and Hare 2008; Forestry Commission 2023).

C.2. Threat to biodiversity

Even in the absence of SQPV and other disease in some Scottish grey squirrel populations, red squirrels are still susceptible to replacement by greys through resource competition (Shuttleworth et al. 2012). On mainland Britain, red squirrel populations have become extinct in the southern counties of England, although they still exist on some off-shore islands, and fragmented populations persist in northern English counties, Wales and Scotland (Mathews et al. 2018). Preventing the establishment of sympatric grey squirrel populations and reducing their competitive ability is central to conservation strategies for red squirrels, which is achieved by proactive planting and forest management regimes within extensive Sitka spruce dominated plantations (Shuttleworth et al. 2012).

Grey squirrels may also impact negatively on urban bird assemblages, either through nest predation (Moller 2008) or competition for supplementary food (Bonnington et al. 2013), although empirical evidence is limited. Bonnington, Gaston, and Evans (2014) found that grey squirrels appeared to have some influence on the structure of urban bird assemblages, but only species most sensitive to nest predation by grey squirrels and not species less sensitive to nest predation. They also found no evidence that grey squirrels reduced the abundance or species richness of wintering avian assemblages in highly urbanised regions due to food competition, most likely because of the high abundance of supplementary feeding stations in UK gardens (approximately half of UK households feed birds: (Davies et al. 2009)). In contrast, Hanmer, Thomas, and Fellowes (2018) found that experimental bird feeders in suburban gardens were utilised by grey squirrels for over 44% of the time, virtually excluding birds from feeding at the same time (>99.99%). Feeders where grey squirrels were dominant were less likely to be visited by birds, even when grey squirrels were absent. The authors concluded that grey squirrels reduced the availability of supplementary food to wild birds.

Eurasian beaver - *Castor fiber*

Introduction

Beavers are native to Great Britain and were once widespread throughout freshwater habitats before being hunted to extinction around 500 years ago (Halley, Saveljev, and Rosell 2020). Reintroductions and natural spread have since restored the species to large areas of its original range with estimates increasing from around 1,200 animals over a century ago to a minimum 1,500,000 presently (Halley, Saveljev, and Rosell 2020).. Interest in beaver reintroductions has been increasing in Britain due to their ability to alter ecosystem structure and function through their engineering activities, most notably the building of dams (Puttock et al. 2017) and it has made a considerable recovery in many areas due to conservation efforts, reintroductions and legal protection (Halley, Rosell, and Saveljev 2012). Beavers inhabit riparian broadleaf woodland or scrub, requiring year-round access to standing fresh water or slow-moving watercourses with suitable herbaceous vegetation for dam-building and foraging (Macdonald et al. 1995). They eat a wide variety of aquatic and terrestrial vegetation including aspen, willow, poplar, alder, grasses and forbs (Gaywood et al. 2015). Known populations of beavers are currently limited to two areas in Scotland (Tayside and Knapdale) and three areas in England (the River Otter and River Tamar in Devon, the River Stour in Kent) (Common, Donald, and Sainsbury 2024). Population estimates (of both free-living and fenced beavers) have increased rapidly over a short period of time to around 2,000 currently (Rosell and Campbell-Palmer 2022), although this figure is likely to be higher. In Scotland alone, the number of beavers more than doubled in three years to around 1,000-1,500 individuals (Campbell-Palmer et al. 2021; NatureScot 2022a).

A.1. Damage to forestry

Beavers are ecosystem engineers that extensively affect the surrounding environment, with tree-felling a key part of this process (Puttock et al. 2017). Much of the opposition to the presence of beavers arise from fear of socio-economic impacts that their activity may have on forested land by destroying vegetation or causing flooding (NatureScot 2015). In addition, they may cause conflict by felling trees of commercial value (Campbell-Palmer et al. 2015). In Scotland, most forestry relies on conifers and therefore beavers are unlikely to have much impact through felling; however, if conifers are present in beaver-flooded riparian areas, they are prone to death as most coniferous species are intolerant of prolonged flooding (Gaywood 2017). Campbell, Dutton, and Hughes (2007) reported that the cost of conflicts by beavers, according to 57 experts involved in research or management from mainland Europe, was higher for forestry activities than all other land-use types, although still relatively low given they represented annual costs (median cost of €101-1,000 (approximately £120-1,185) per beaver population). Most conflict was due to foraging activity (60% reported felling of commercial trees) or loss of forestry land due to flooding. They concluded that conflicts were generally localised and small scale, rarely rising above €10K (approximately £11.8K) per year.

A.2. Damage to agricultural interests

Beaver activities may affect agricultural land due to burrowing and canal construction, damming of smaller water courses leading to flooding and direct foraging of crops (Campbell-Palmer et al. 2015). In Scotland, concerns are greatest where beaver activities affect areas of intensive agriculture (NatureScot 2015) such as Tayside, where agriculture represents a major land use of which the majority is prime agricultural land (Coz, Young, and Gibbs 2020). Campbell, Dutton, and Hughes (2007) reported that the cost of conflicts by beavers on agricultural activities, according to 57 experts from mainland Europe, appeared to be small with a median range of €1-101 (approximately £120-1,185) per population annually. Most conflict was due to beavers foraging on crops (53% reported this activity) or loss of agricultural land due to flooding. They concluded that conflicts were generally localised and small scale, rarely rising above €10K (approximately £11.8K) per year.

A.6. Predation/impact on commercial fisheries

Beavers do not eat fish but there are concerns they potentially impact fish stocks by hindering fish movements (Gaywood 2017), such as wild migratory Atlantic salmon, and within inland salmon and trout fisheries (Coz, Young, and Gibbs 2020). Armstrong et al. (2015) looked at the overlap between potential beaver habitat and salmon habitat in six study catchments in Scotland and found an overlap of 47-73%. As the value of Atlantic salmon fishery is estimated to be over £73.5M per year (Radford et al. 2004), beavers could potentially cause substantial economic conflict, although it is uncertain exactly what impact they have on local fisheries (Gaywood 2017). Conversely, beavers have been found to have positive effects on local fish populations; Needham et al. (2021) found that of two streams entering the same loch in northern Scotland, the one modified by beavers promoted higher abundances of larger sizes of brown trout. Campbell, Dutton, and Hughes (2007) reported that the cost of beaver conflicts with fisheries, according to 57 experts from mainland Europe, appeared to be negligible with most respondents reporting no conflict. Incidents involving greater costs were isolated, rarely rising above €10K (approximately £11.8K) per year.

A.10. Conflicts with construction, infrastructure and development

There is potential for beaver activities, such as damming, burrowing and tree-felling, to impact a range of infrastructure including flooding of roads and tracks, culverts, weirs, flood banks, canals and water treatment plants (Campbell-Palmer et al. 2015). Impacts are likely to be local but could become more widespread.

B.2. Reservoir of disease

Eurasian beavers are potential hosts for a range of infectious diseases and parasites, including those typically associated with common European rodents (Girling et al. 2019). These authors found that the highest-risk non-zoonotic pathogens identified from previous literature on beavers (both the Eurasian beaver and North American beavers, *Castor canadensis*) were *Chrysosporium parvum* (*Emmonsia parva* and *E. crescens*) and *Mycobacterium avium* subspecies *avium*. These diseases were reported separately in single beavers found dead in Sweden and the Netherlands respectively but have currently not been reported in any beavers in Britain. *Emmonsia* spp. (formerly *Chrysosporium* spp.), specifically *E. crescens*, is considered an environmental pathogen found in a broad range of wildlife species in Britain (Common, Donald, and Sainsbury 2024); it has been found in several rodent species, including wood mice and bank voles (Kruckemeier et al. 2024) and therefore beavers have the potential to become infected with this pathogen through environmental transmission. Given that *E. crescens* is probably a ubiquitous organism, there is a low likelihood that spores released from infected beavers that have died increase the environmental burden of infective conidia (Common, Donald, and Sainsbury 2024).

Common, Donald, and Sainsbury (2024) also reported that *Eimeria* spp. oocysts were detected in a single live-trapped beaver in Tayside during health screening of the population between 2013 and 2019, which presented with no associated signs of disease. *Eimeria* spp., which cause coccidiosis, is common in many rodents but they show a high degree of host specificity and cause subclinical infection (Yarto-Jaramillo 2015), therefore the risk beavers pose is minimal.

<u>B.4. Reservoir of zoonotic disease</u>

Girling et al. (2019) identified a number of high-risk zoonotic pathogens from previous literature on beavers, including *Cryptosporidium parvum*, *Echinococcus multilocularis*, *Fasciola hepatica*, *Giardia* spp., *Trichinella britovi*, *Escherichia coli*, *Franciscella tularensis*, *Mycobacterium avium*, *Salmonella* spp., *Yersinia* spp., and terrestrial rabies virus. The process of translocating beavers for release in Great Britain poses the greatest risk to public health, particularly if beavers originating from countries where zoonotic parasites are endemic or free-living beavers from Great Britain of unknown origin are translocated (Common, Donald, and Sainsbury 2024). In a disease risk analysis carried out for the conservation translocation of beavers in England, Common, Donald, and Sainsbury (2024) identified five zoonotic pathogens of importance that presented a varying risk of disease to people. The highest risk pathogen was *E. multilocularis*, which could cause severe biological and economic consequences from its incursion given the UK's current status of being infection-free from this cestode. As a large proportion of beavers in captivity (in enclosures) in Great Britain originate from areas where *E. multilocularis* is endemic, it is possible they harbour the parasite and pose a risk if translocated. This risk is reduced if second-generation beavers are used for translocations, as the parasite cannot be transferred from parent to offspring, or if beavers proven to have been born in Great Britain are translocated (Common, Donald, and Sainsbury 2024).

*Trichinella* spp. and *Taenia martis* were identified as medium-risk if free-living beavers from Great Britain were translocated to England and high-risk if beavers in captivity from known endemic areas were translocated (Common, Donald, and Sainsbury 2024). Like *E. multilocularis*, maintaining the UK's infection-free status for these parasites is important given the severity of disease in people and the high economic cost of preventing these diseases should they become endemic in the UK, particularly *Trichinella* spp. (Common, Donald, and Sainsbury 2024). The risk is reduced if second-generation beavers or beavers born in the Great Britain are translocated (Common, Donald, and Sainsbury 2024).

*Brucella* spp. and zoonotic hantaviruses (Puumala orthohantovirus, PUUV, which is the etiological agent of the human disease nephropathia epidemica and Saaremaa virus, SAAV) were also identified as medium-risk (Common, Donald, and Sainsbury 2024). *Brucella* spp. have a worldwide distribution, but the UK is currently free from the species causing brucellosis in humans and livestock (Common, Donald, and Sainsbury 2024). PUUV and SAAV are endemic in some European countries, such as Norway and Germany respectively. Like the aforementioned zoonotic pathogens, the risk of introducing these pathogens to the UK is higher if beavers from endemic areas or beavers of unknown origin from the UK are used for translocation (Common, Donald, and Sainsbury 2024).

Hazel dormouse - *Muscardinus avellanarius*
Introduction

The hazel dormouse is native to the UK and found primarily in broadleaved woodland, generally associated with early successional stages of woodland or coppice (Bright and Mitchell-Jones 2006) but also scrub, coniferous plantations and hedges (Chanin and Woods 2003). They have an omnivorous diet including high-quality plant foods (flowers, buds, seeds, fruits) and insects (Mathews et al. 2018). They are protected in the UK under the Wildlife and Countryside Act, 1981 and are a Priority Species under the UK Post-2010 Biodiversity Framework. They are also listed as a European Protected Species under the European Habitats Directive (www.wildlifetrusts.org). They are mostly found in southern England and Wales, but are absent from Scotland (Mathews et al. 2018). The population is in decline and had reduced by 72% between 1993 and 2014 (Goodwin et al. 2017). There are an estimated 930,000 hazel dormice in Britain: 757,000 in England, 172,000 in Wales (Mathews et al. 2018).

A.10. Conflicts with construction, infrastructure and development

Limited information, but because of the protection hazel dormice are afforded, conflicts likely arise where they are present at sites where development is proposed. In England, guidance must be consulted when assessing planning applications if hazel dormice are found on or near to a proposed development site (Natural England 2022b).

Edible dormouse - *Glis glis*
Introduction

The edible dormouse was introduced to the Chiltern Hills in England by Walter Rothschild in 1902 (Morris 2008). The dormice multiplied rapidly and caused damage to crops, forestry and buildings and were therefore the subject of an unsuccessful campaign to eradicate them (Long 2003). The expansion of their range has been very slow and they are still confined to the Chilterns area (Morris 2008), with habitat fragmentation and barriers thought to limit its spread (Mathews et al. 2018). Where they are present, they can be a source of conflict between human and wildlife interests, causing an unacceptable nuisance and damage to houses (Morris 2008). The edible dormouse is found in coniferous, broadleaved and mixed woodland, as well as orchards and gardens (Harris et al. 1995); it shows a dietary preference for oak acorns and beech nuts (Ruf et al. 2006) but also eats seeds, fruits, berries, buds, bark, fungi, insects, eggs and carrion (www.mammal.org.uk). There are an estimated 23,000 in England only (Mathews et al. 2018).

A.1. Damage to forestry

Edible dormice strip patches of bark from trees, especially pine, spruce and beech, which leaves them vulnerable to fungal infection and crown death (Gurnell and Hare 2008). Although forestry damage is limited to the Chilterns area, it can be locally significant. A survey of 14,000 trees in Forestry Commission woods (Brady, unpublished; cited by Jackson (1994)) in areas occupied by edible dormice found 14.6% of the trees had had patches of bark removed, representing a 25% loss of revenue equivalent to approximately 220 trees per hectare. Up to 70% of trees in other stands were affected, estimated at 2,000 trees per hectare. The Forestry Commission has estimated the cost of damage by edible dormice at approximately £25K per year on land managed by them, in addition to £170K-400K losses on privately-owned woodland, based on the assumption that approximately 50% of trees are affected. In addition to this tree damage, Forest Research and Forest Enterprise have spent approximately £61K on research, advice and information on edible dormouse, giving an estimated total cost of edible dormouse to forestry of £250K per year (entirely attributable to England) (Williams et al. 2010). As no recent costs associated with ring barking and damage to forestry were found, Eschen et al. (2023) adjusted this figure for inflation to £300K, excluding associated research costs.

A.2. Damage to agricultural interests

Fruit crops from orchards have been the subject of damage by edible dormice in the past but this is now of less importance as the extent of orchards in the Chiltern Hills has decreased (Morris 1997). No recent information found.

A.8. Damage to property

The edible dormouse has been found to cause nuisance in houses where it will deposit faeces and urine, cause noise during the night, and gnaw on wood, insulation, water pipes and electrical wires (Büchner, Trout, and Adamík 2018). Damage to property is localised and tends to be recurrent, and likely more of a nuisance than a serious economic impact. The total cost of damage caused by edible dormice in England was estimated at £114K, which included £62K of associated property damage and £52K of estimated control costs (Williams et al. 2010); when adjusted for inflation, this figure was estimated to be £200K (Eschen et al. 2023).

B.4. Reservoir of zoonotic disease

The edible dormouse has been reported as a competent reservoir for particular genospecies of the tick-borne agent of Lyme disease (Fietz et al. 2014) and was associated with the transmission of *Borrelia burgdorferi* to humans in a study conducted in central Europe (Matuschka et al. 1994). As the likelihood of encountering infected ticks increases over time and thus the age of the host, edible dormice may play an important role in the transmission cycle of Lyme disease given their longevity (lifespan of 13-14 years: Trout, Brooks, and Morris (2015)). They may also enhance their capacity as a vector of the disease through nesting behaviour, which exposes them to nymphal ticks developed from larvae previously detached in the same nest (Matuschka et al. 1994). As edible dormice will often enter domestic properties, there is a risk of disease transmission to people; however, this risk is likely to be low given the limited distribution of edible dormice in England and the seasonality of spirochete prevalence in this species (Fietz et al. 2014).

**Rodentia – small field rodents**

Bank vole - *Myodes glareolus*
Field vole - *Microtus agrestis*
Wood mouse - *Apodemus sylvaticus*
Yellow-necked mouse - *Apodemus flavicollis*
Orkney vole - *Microtus arvalis orcadensis*
Skomer vole - *Myodes glareolus skomerensis*

Introduction

There are numerous small mammals native to the British Isles, inhabiting a range of different habitats. Mathews et al. (2018) reported that several vole species (bank, Orkney and Skomer) and wood mice are found in a wide variety of habitats, such as hedgerows, conifer plantations, broadleaved and mixed woodland, with wood mice also found in heather, blanket bog, sand dunes and urban areas. Yellow-necked mice are primarily associated with mature broadleaved woodland and hedgerows, whilst field voles are normally found in low productivity grassland. They are all native to Britain, except Orkney and Skomer voles (although they are considered naturalised); Orkney voles were introduced to the Orkney archipelago approximately 5,000 years ago, whereas the time of introduction of Skomer voles is unknown (Mathews et al. 2018). According to Mathews et al. (2018), the current estimates of small field rodent population sizes are as follows: 27,400,000 bank voles (19,100,000 in England, 5,390,000 in Scotland, 2,930,000 in Wales); 59,900,000 field voles (28,600,000 in England, 21,500,000 in Scotland, 9,760,000 in Wales); 39,600,000 wood mice (22,700,000 in England, 12,300,000 in Scotland, 4,600,000 in Wales); 1,500,000 yellow-necked mice (1,360,000 in England, 140,000 in Wales); between 1,000,000 and 2,000,000 Orkney voles. There are thought to be approximately 2,000 Skomer voles (www.animalia.bio/skomer-vole).

A.1. Damage to forestry - bank vole, field vole

The bank vole is sometimes considered a threat to forestry as it has been reported to eat seeds and seedlings, gnaw on roots and strip bark from trees (Gurnell and Hare 2008). It has been reported that bank voles can remove buds, particularly of newly planted pine trees (Forestry Commission 2023). However, whereas most damage occurs in winter in mainland Europe, when alternative food availability is relatively low, this is not the case in Britain with the majority of bank vole diet comprising of dead leaves (Hansson 1985). The field vole can cause significant damage to plantations of young trees but only causes problems in deciduous woodland (Gurnell and Hare 2008). Damage caused by field voles can occur all year round (Forestry Commission 2023) and due to the widespread distribution of these vole species and confirmed damage, it is likely they cause a degree of economic loss.

A.2. Damage to agricultural interests - field vole, wood mouse

In arable farmland, the field vole causes damage by cutting stems near to the base and then eating the stem and leaves (Gurnell and Hare 2008). The wood mouse is known to feed on stored and sown grain and seed, with a particular preference for sugar beet, but this has not been found to cause any significant economic loss (Pelz 1989). It will also feed on the grass weeds commonly found in cereal crop plantations, and therefore has low significance as a seed predator of cereal crops themselves (Westerman et al. 2003). No recent information found.

A.13. Sustaining predators harmful to game - field vole, Orkney vole

Predators often respond to reductions in preferred prey species by switching to alternative prey resources (Francksen et al. 2017). However, several studies have reported that a higher abundance of some vole species may also increase predation on other prey species. Francksen et al. (2017) found that buzzards predated red grouse incidentally while hunting for field voles, which may increase when vole abundances are high through promoting foraging in heather moorland habitats where grouse are more numerous. Similarly, Ludwig et al. (2017) found that higher field vole abundance was associated with increased predation of red grouse chicks, most likely by rodent-hunting raptors.

Orkney voles have also been reported as a frequent prey component of hen harriers in Orkney, which have been heavily persecuted in Great Britain due to its predation of red grouse (Nota, Downing, and Iyengar 2019). It is estimated that the game shooting is worth £1.6 billion to the UK economy, with approximately 12% attributed to grouse shooting (Sotherton, Tapper, and Smith 2009), therefore potential conflict may arise given the protected status of hen harriers under the Wildlife and Countryside Act, 1981 and the UK Birds of Conservation Concern (red listed).

B.2. Reservoir of disease - field vole, wood mouse

Field voles are known maintenance hosts of *Mycobacterium microti*, which belongs to the *M. tuberculosis* complex (van Soolingen et al. 1997). It causes tuberculosis and has been identified in a range of other mammalian species, most notably cats, with sporadic cases reported in humans (Kipar et al. 2014). Of 327 field voles sampled from Kielder Forest, Northumberland in 2003, 43 (13.2%) were identified as being infected with *M. microti*, by histopathology and/or culture (Kipar et al. 2014). These authors concluded that the morphology of the lesions was consistent with active disease; the presence of mycobacteria in open skin lesions, airways and salivary glands indicated bacterial shedding from these areas, suggesting the environment, direct contact and consumption of field voles as likely sources of infection.

Wood mice, potentially infected with murine adenovirus, have been suggested as potentially increasing grey squirrel adenovirus infection rates through (as yet unconfirmed) inter-species transmission (Greenwood and Sanchez 2002), which may transfer to sympatric red squirrel populations (Everest et al. 2018). They have also been identified as a risk factor to captive red squirrel collections, particularly those bred for conservation translocation within the UK (Everest et al. 2018).

B.4. Reservoir of zoonotic disease - bank vole, field vole, Skomer vole, wood mouse, yellow-necked mouse

Rodent species harbour a large proportion of zoonotic parasites and are one of the taxa with the highest zoonotic potential (Olival et al. 2017). They present a potential health risk from leptospirosis through contact with the urine of infected animals (Gurnell and Hare 2008). Ball (2014) analysed 283 rodent samples from rural and urban sites in the UK, identifying infection with the pathogenic species *Leptospira interrogans* in 10.5% (n=16/152) wood mice, 10.6% (n=5/47) bank voles and 20% (n=2/10) field voles. Ball (2014) concluded that several wild rodent species in England were capable of maintaining and shedding pathogenic strains know to infect humans and dogs.

In woodland habitats in Ireland, wood mice have been indicated as putative reservoir hosts of Lyme borreliosis from bites of infected ticks (Smith, Gray, and McKenzie 1991). However, during eco-epidemiological screening of rodent species in rural west Wales between 2015 and 2017, no *Borrelia burgdorferi* positive samples were detected in 358 rodents (299 bank voles (of which 163 were Skomer voles), 59 wood mice: Occhibove et al. (2022). In the same study, *Anaplasma phagocytophilum* and *Babesia microti* were detected in a single pooled sample of ticks collected from bank voles, suggesting they may play a role in maintaining tick populations. *Bartonella* spp. were also detected in 14 fleas collected from 12 of 299 (4.0%) bank voles, one of nine (11.1%) field voles, one of 59 (1.7%) wood mice.

The bank vole is the rodent host of Puumala hantavirus (PUUV), which causes the human disease nephropathia epidemica (NE), considered a mild form of hemorrhagic fever with renal syndrome (HFRS) (Kallio et al. 2009). In Finland, Kallio et al. (2009) studied a large and long-term dataset (14 years, 2,583 human cases of NE and 4,751 trapped bank voles) and found that the number of human infections was influenced by the phase of vole cycle and time of year, and vole abundance. A vole-associated hantavirus (Tatenale virus) was discovered in an individual field vole in northern England during surveillance between 2009 and 2011 (Pounder et al. 2013). Subsequent screening of the extensive population of field voles in Kielder Forest, detected Tatenale virus-like lineage at a prevalence of 17% in liver tissue (Thomason et al. 2017). These authors concluded that Tatenale-like hantavirus lineages are widespread and common in northern England, raising the possibility that Tatenale-like virus could have been responsible for some of the human cases of HFRS, as PUUV has never been reported in the UK (Chappell et al. 2020). A third lineage of Tatenale-like virus has since been detected in central England (Chappell et al. 2020) suggesting this virus could be widespread in the field vole population. Wood mice are also reservoirs of hantavirus elsewhere in Europe, although the UK position is unclear (Vapalahti et al. 2003). Likewise, the yellow-necked wood mouse is a known reservoir of the Dobrava hantavirus in Europe, which causes a severe form of haemorrhagic fever with renal syndrome in humans (Sibold et al. 2001).

Wild rodents have been implicated as a source of zoonotic pathogens, including *Yersinia enterocolitica* (Keesing and Ostfield 2024), which is transmitted via the faecal-oral route. Given their potential transmission risk at the livestock-wildlife interface, they represent a major food-borne public health hazard (Arden et al. 2022). Of 139 wild rodents sampled from across the UK, 8 (5.8%) were confirmed as *Y. entercolitica* positive (2.7% (2/75) wood mice, 2.1% (1/47) bank voles, 29.4% (5/17) field voles), which in turn were all deemed to be a non-pathogenic biotype (Arden et al. 2022) therefore posing minimal risk to public health. During eco-epidemiological screening of rodent species in west Wales between 2015 and 2017, *E. coli* was identified in 1.7% (6/358) individuals: 1.7%(1/59) wood mice and 1.7% (5/299) bank voles (Occhibove et al. 2022). These authors concluded that the prevalence of *E. coli* was low in wild rodents and that they do not represent a reservoir host for pathogenic strains of *E. coli*, which has been found elsewhere (e.g. (Kilonzo et al. 2013)).

Ljungan virus is a recently identified member of the genus Parechovirus within the family Picornaviridae that was isolated from bank voles in Sweden (Niklasson, Hornfeldt, and Lundman 1998). It has been suggested that it has zoonotic potential, as the other member of the Picornaviridae family is human parechovirus, which is commonly found in children with diarrhoea and gastroenteritis (Harvala and Simmonds 2009). In a study of 209 small rodents sampled from within and near to Kielder Forest,

24.4% (51/209) were PCR positive for Ljungan virus: 27.0% (10/37) of bank voles, 26.5% (22/83) of field voles, 19.7% (13/66) of wood mice, 26.1% (6/23) of house mice (Salisbury et al. 2014). These authors concluded that the prevalence of Ljungan virus in the rodent population highlighted the need to confirm the zoonotic potential of this pathogen.

Herpesvirus was detected by PCR in Skomer voles at a low prevalence (2.5%, 4/163), with sequences showing overall similarity greater than 98% with human alphaherpesvirus 3 varicella-zoster (three samples), and human alphaherpesvirus 2 herpes simplex (one sample: Occhibove et al. (2022)).

(See common shrew for information on alphacoronoviruses). Tsoleridis et al. (2016) detected novel alphacoronoviruses in a number of rodent and shrew species sampled from across the East Midlands region of the UK between 2008 and 2015 with100% (1/1) of bank voles and 27.3% (3/11) of field voles testing PCR positive for CoVs. During the same study, 300 bank voles from Poland were also tested with only one (0.3%) testing PCR positive for CoVs. As the novel rodent/shrew CoVs discovered in the UK were phylogenetically separate from bats, further investigation is warranted to determine the zoonotic potential of these CoVs.

C.1. Damage to conservation habitats - bank vole

Some plant communities of conservation concern, such as montane willow scrub, are under threat globally due to increased herbivory by both large wild and domestic mammalian herbivores (Brookshire et al. 2002). Montane willow scrub comprises up to eight characteristic willow species, of which four are scarce in the UK (Shaw et al. 2013). Removal or management of large herbivores can alter the biomass and structure of vegetation, which may favour small mammals that feed on juvenile plant stages and in turn impact the plant community composition (Shaw et al. 2013). As natural regeneration of montane willow scrub is slow, nursery-grown saplings may be planted to augment extant threatened populations or to establish new populations (McBride 2002). Shaw et al. (2013) found that bank voles damaged approximately 40% of juvenile nursery-grown saplings planted in enclosures in Scotland (where large herbivores had been excluded), resulting in smaller increases in height, although mortality due to predation was very low (between 0.7 to 1.8% depending on willow species). These authors concluded that while direct mortality was low, it is conceivable that the impact of damage on growth of saplings may interact with other causes of mortality such as further herbivory or competition from other plants.

Water vole - *Arvicola amphibius*

Introduction

Water voles are primarily riparian in Great Britain, usually found within 2m of slow-flowing rivers, streams and marshes with tall, dense vegetation that provides cover from avian predators (Lawton and Woodroffe 1991). They are protected under the Wildlife and Countryside Act 1981 and are a priority species under the UK Post-2010 Biodiversity Framework. They are also listed as endangered on both the GB and England Red List for Mammals. Water voles are widespread throughout mainland UK, although their range and numbers have significantly declined due to habitat loss or alteration and predation by American mink (MacPherson and Bright 2011; Mathews et al. 2018). There are an estimated 132,000 water voles in Britain: 77,200 in England, 50,000 in Scotland, 4,500 in Wales (Mathews et al. 2018).

A.10. Conflicts with construction, infrastructure and development

Due to the water voles' protected status, which includes burrows and other structures used for shelter, conflict can arise where individuals are present in riparian areas requiring maintenance or development, e.g. to regrade eroding banksides or installing pipelines (Gelling et al. 2018). Activities used to displace water voles safely prior to development can include habitat manipulation, such as vegetation removal and/or water draw-down, to encourage the relocation of water voles from within the affected area, up to a maximum bank length of 50m (Gelling et al. 2018) or complete relocation. In 2015, the Environment Agency spent £135K removing 55 water voles from the Somerset Levels prior to dredging work, which included a wildlife survey, winter storage and relocation costs to Hampshire and Cornwall (BBC 2015a).

House mouse - *Mus musculus*

Introduction

Since its introduction to the UK in the Iron Age, the house mouse has become widely distributed, including on many small islands, although on the mainland the species generally lives commensally with human activities in and around buildings in both rural and urban settings (Gurnell and Hare 2008). Due to these widespread commensal populations, the house mouse poses potential largescale conflict with human interests. Mouse populations can reach very high densities of up to 7/m$^2$ (Berry 1991) in some types of farm building (e.g. indoor livestock units) in the absence of competitors such as rats. They are widely distributed in England, with a patchier distribution across Wales and Scotland. There are an estimated 5,203,000 house mice in Britain: 4,340,00 in England, 523,000 in Scotland, 339,000 in Wales (Mathews et al. 2018).

A.2. Damage to agricultural interests

The primary damage that the house mouse causes across the UK is consumption of stored food products and contamination with faeces and urine (Gurnell and Hare 2008). Despite their small size and relatively low individual food intake, problems can be exacerbated as comparatively large amounts of grain can be fragmented and rendered unsuitable due to the distribution and commensal habits of house mice (Gurnell and Hare 2008). Wildey (2002) documented that rodents (mainly house mice and rats) infested approximately 70% of grain stores in the UK irrespective of type.

A.8. Damage to property

House mice cause considerable hygiene problems in buildings, producing over 50 droppings per individual per day and extensive urine marking (Gurnell and Hare 2008). Further damage is caused by gnawing through electrical cables and insulating material (Gurnell and Hare 2008; Stejskal, Rodl, and Aulicky 2016) with associated maintenance costs and fire risks. A survey based on the English House Condition Survey revealed that 1.83% of domestic dwellings in England have house mice living indoors (Langton, Cowan, and Meyer 2001). The annual cost of house mouse control in homes was estimated at £17.9M (Williams et al. 2010). Revised costs of house mouse control are considerably lower (£6.3M) (Eschen et al. 2023).

B.2. Reservoir of disease

House mice can be a reservoir and vector of a variety of human and animal diseases (Becker et al. 2007). Pocock et al. (2001) found the house mouse was not a significant role for the transmission of *Yersinia* or *Salmonella* to livestock; in contrast, Davies and Wales (2013) reported both *Salmonella* Derby and *Salmonella* Bovismorbificans in mice removed from a pig farm in South England, with mice excreting both serovars for 10 months after capture.

Ljungan virus is a recently identified member of the genus Parechovirus within the family Picornaviridae that was isolated from bank voles in Sweden (Niklasson, Hornfeldt, and Lundman 1998). It has been suggested that it has zoonotic potential, as the other member of the Picornaviridae family is human parechovirus, which is commonly found in children with diarrhoea and gastroenteritis (Harvala and Simmonds 2009). In a study of 209 small rodents sampled from within and near to Kielder Forest, 24.4% (51/209) were PCR positive for Ljungan virus: 27.0% (10/37) of bank voles, 26.5% (22/83) of field voles, 19.7% (13/66) of wood mice, 26.1% (6/23) of house mice (Salisbury et al. 2014). These authors concluded that the prevalence of Ljungan virus in the rodent population highlighted the need to confirm the zoonotic potential of this pathogen.

B.4. Reservoir of zoonotic disease

Cats are the definitive hosts of *Toxoplasma gondii*, which is transmitted when contaminated prey is consumed. Several species can act as intermediate hosts, including humans and house mice in infested dwellings (Murphy et al. 2008; Smith et al. 2021). Out of 200 house mice trapped in 27 infested properties in Manchester, UK, 59% (118) tested positive for *Toxoplasma* infection (Murphy et al. 2008). These authors also found evidence of vertical transmission to foetuses possibly maintaining the infection in urban areas. They concluded that high mouse densities in urban areas pose a significant risk of transmission of *Toxoplasma gondii* to humans.

Brown rat - *Rattus norvegicus*

Introduction

The brown rat is found throughout Britain, with the exception of exposed mountainous regions and a few offshore islands; it is more common in the east and southeast of England where cereal crops are grown most extensively (Gurnell and Hare 2008). It was accidentally introduced to Britain during the 18th century and has become very well established as a versatile commensal rodent, although it is generally found at lower densities in areas where competing species occur (Gurnell and Hare 2008). Typical habitats where the brown rat is found are farms, sewers, landfill sites, urban waterways and warehouses but they can also occupy hedgerows and ditches near cereal plantations or in coastal habitats (Gurnell and Hare 2008; Mathews et al. 2018). There are an estimated 7,070,000 brown rats living in Britain: 4,730,000 in England, 1,060,000 in Scotland, 1,280,000 in Wales (Mathews et al. 2018).

A.2. Damage to agricultural interests

Brown rats are mostly found in rural areas and have been recorded as being present on 42% of agricultural properties (Meyer 1994). Rats cause several types of costs to farms, including yield loss, damage to electrical wiring and spoilt stored crops (Eschen et al. 2023). Although the brown rat will consume stored crops and animal feed, the primary source of damage to stored food is contamination with hair, droppings and urine (Meyer 1994). One study reported that 70% of a tonne of wheat was spoilt by 10 to 26 rats during a 12 to 28-week period, when only 4.4% had been eaten (Buckle 2007). Damage to the electrical wiring of farm vehicles and equipment is a common cause of breakdown and the need for expensive repairs, while rats are said to cause 50% of electrical fires (Richards 1989), which can result in significant costs (Kelly et al. 2013). In recent years, the cost of farm fires due to electrical faults, lightning strikes and arson is reported to have increased considerably by 37% (Farmers Weekly 2024). Buckle (2007) estimated the cost of damage caused by rats to the UK farming industry at £21M per annum, which included costs due to consumed and spoilt stored crops and animal feed, damage due to electrical fires as well as field crop losses. Williams et al. (2010) provided a similar estimate of £21.8M (£10.9M in England, £4.4M in Wales and £6.5M in Scotland). Revised estimated costs of rat damage in Britain were significantly higher, totalling £54.4M, which included £14.9M for pre-harvest yield loss, £12.4M for rat control and £27.1M for farm fires (Eschen et al. 2023).

A.5. Predation of game species

There is limited published information on rat predation of game species but it is cited as an issue by gamekeepers. Evidence from other bird groups, such as seabirds, suggest that rat predation can take place at various life stages as rats will take both chicks and adult birds, as well as eggs (Jones et al. 2008), which may also apply to game birds. However, it is likely that game birds will attract fewer rats than seabirds as they do not breed in large accessible colonies but are sometimes aggregated in rearing pens, which may be attractive to rats.

A.8. Damage to property

A survey, based on the English Housing Condition Survey, revealed that 1.6% of domestic dwellings in England had brown rats living outdoors and 0.23% had brown rats living indoors (Langton, Cowan, and Meyer 2001). Brown rats can gnaw through any material softer than the enamel on their teeth, including most building woods, aluminium sheeting, soft mortar and poor-quality concrete (Williams et al. 2010). A British Pest Control Association 2016 survey found that rats accounted for 46% of pest control call-outs (Eschen et al. 2023). Williams et al. (2010) estimated the annual cost of rat control in homes in Britain to be £31.6M (£27.1M in England, £2.9M in Scotland, £1.6M in Wales). Eschen et al. (2023) revised these costs for local authorities only (excluding private control companies) with an estimated cost to the British economy of £14.8M (£13.0M in England, £700K in Scotland, £1.1M in Wales).

A.12. Conflicts with transport

Brown rats do not cause major problems to the railway network, although isolated incidents have generated substantial reactive costs and delays (Williams et al. 2010). Damaged cables and subsequent impacts on signalling are the main issues caused by rat damage (Battersby 2005). Williams et al. (2010) estimated potential costs to railways and passengers of around £5.1M (£3.3M in England, £900K each in Scotland and Wales). Using 2021 data from Network Rail, Eschen et al. (2023) estimated considerably lower costs to Britain's railway network of £600K (£400K in England, £100K in Scotland, £100K in Wales).

A.15. Conflicts with utilities

Brown rats can cause conflict with the water industry, more specifically the sewage network. Baiting to control rats across the national sewerage system has been estimated to cost approximately £3.2M in Britain (£2.6M in England, £450K in Scotland, £154K in Wales) (Williams et al. 2010). Eschen et al. (2023) revised these costs due to inflation with rats costing the British water industry £4.6M (£3.8M in England, £600K in Scotland, £200K in Wales).

B.4. Reservoir of zoonotic disease

The brown rat is an important vector for a variety of diseases affecting humans of which leptospirosis is a particularly high risk for people coming into contact with contaminated water (Gurnell and Hare 2008). However, a study of 67 rats sampled from Liverpool found no evidence of *Leptospira* infection, although it should be noted that none of the rats were trapped in rural or semi-rural areas (Ball 2014). A study of the prevalence of Q fever (*Coxiella burnetii*) in the brown rat found that rats are an important reservoir for the disease, which can be transmitted to humans. It can also be transmitted to cats preying on infected rats, which in turn may come into even closer contact with people (Webster and Macdonald 1995).

Hantaviruses are also of concern regarding the brown rat as they can cause hemorrhagic fever with renal syndrome and hantavirus pulmonary syndrome in humans. They are transmitted via the inhalation of contaminated aerosols of rodent excretions and are therefore of most concern where rats are occupying domestic buildings in urban areas (Schmaljohn and Hjelle 1997). Seoul hantavirus has been detected in wild brown rats in the UK and implicated as the source of established human disease (Jameson et al. 2013).

The brown rat has also been found to be a significant reservoir of *Cryptosporidium* spp. which causes diarrheal illnesses in humans and livestock (Quy et al. 1999). This disease is a particular danger to individuals with low immunological defences, for whom it can be fatal (Webster and Macdonald 1995).

In the UK, the number of cases of hepatitis E virus (HEV) infections in humans has been increasing since 2010 (PHE 2020). Although most cases are considered to be linked to the consumption of pork or pork products infected with *Orthohepevirus* A genotype 3, the dominant subgroup circulating in British pigs differs from the one found in people, suggesting infections have been acquired from imported pork products or an alternative zoonotic source (Murphy et al. 2019). As brown rats have been shown to carry both swineHEV and ratHEV, Murphy et al. (2019) tested 307 rodents from pig farms (n=12) and other locations (n=10) across Northern England, Wales and Scotland in 2014-2016. Of 61 rat livers sampled, 13% (8) were positive for ratHEV at four different locations (two in Cheshire, one each in West and North Yorkshire). No other rodents (house mouse, wood mouse, bank vole, field vole, grey squirrel, red squirrel) tested positive for either swineHEV or ratHEV. As several human cases of ratHEV infections have been reported recently in Asia, infected rats may pose a risk to public health (Murphy et al. 2019).

(See common shrew for information on coronaviruses (CoVs). Although the prevalence of CoVs in rats in the UK appears low (4.1% (5/123)), further investigation into potential cross-species transmission is warranted given these novel CoVs are phylogenetically separate from bats (Tsoleridis et al. 2016).

C.5. Predation of important ground-nesting birds and island ecosystem damage

Introduced rats are responsible for a relatively large number of extinctions from islands across the world (Howald et al. 2007). The brown rat can have devastating ecological impacts on islands where there were previously no natural predators for certain species, particularly seabirds (e.g.Hilton and Cuthbert 2010), causing them to become very vulnerable to predation pressure. Evidence from other bird groups, such as seabirds, suggest that rat predation can take place at various life stages as rats will take both chicks and adult birds, as well as eggs (Jones et al. 2008).Predation on species of conservation concern by the brown rat, such as some ground-nesting birds, has been reported, although direct evidence can be difficult to obtain, especially via the traditional dietary analysis methods (Stapp 2002). Island populations can be impacted severely and damage by rats has been noted to occur in some of the Scottish isles, where eradication is desired to protect the seabird populations (Stapp 2002). On the Scottish Island of Ailsa Craig, burrow-nesting seabirds, such as Atlantic puffins and petrels, are particularly vulnerable to rat predation as they have evolved to avoid avian predators and will leave eggs and chicks unattended when out at sea (Zonfrillo 2000). UK islands support approximately one-fifth of the global Manx shearwater breeding population and several recent projects have focussed on the removal of brown rats to conserve these burrow-nesting seabirds (e.g. Bell et al. (2011)). On the islands of St Agnes and Gugh, Isles of Scilly, a community-based eradication project resulted in the successful eradication of brown rats from the islands (Bell, Floyd, et al. 2019).

The impact of brown rats on seabird populations may differ depending on rat density. O'Hanlon and Lambert (2016) baited seabird nests with domestic quail and hen eggs on the Calf of Man during 2011 and observed rats visiting nests at only one of four locations on 7.5% (19/255) observation nights. No evidence of rat predation was observed. The authors concluded that although predation by rats was likely to occur at higher densities, where pressures on food and resources were greater, where rats occur at low density, predation of eggs is unlikely.

Black rat - *Rattus rattus*

Introduction

The black rat was introduced in Roman times and was historically abundant in the UK until it was displaced by the brown rat in the 18th century (Gurnell and Hare 2008). It subsequently became limited to urban areas, such as dockside warehouses with populations declining further after the containerisation of dockyards and the use of grain silos (Symes and Yalden 2002). Local island populations also existed on Lundy and the Shiants until they were eradicated in 2006 and 2016 respectively by the Seabird Recovery Project (Mathews et al. 2018). There are limited records of black rats, although it is plausible that small populations may still exist (Mathews et al. 2018) with an estimated 1,300 estimated in Britain (British Wildlife Centre 2024).

B.4. Reservoir of zoonotic disease

Black rats can be major reservoirs of pathogens and vectors for diseases that affect humans and animals, including *Babesia* spp., *Toxoplasma gondii* (Zanet et al. 2023), *Angiostrongylus* spp. and *Cryptosporidium* spp. (Simpson et al. 2024). However, as the number of black rats in the UK is thought to be very low it is unlikely they pose a significant risk to public health.

C.7. Damage to island ecosystems

The black rat can have devastating ecological impacts on islands where there were previously no natural predators for certain species, particularly seabirds (e.g. Hilton and Cuthbert 2010), causing them to become very vulnerable to predation pressure. Introduced rats are responsible for a relatively large number of extinctions from islands across the world (Howald et al. 2007), with introduced brown and black rats thought to be responsible for the loss of burrow-nesting seabird populations on Lundy (Lock 2006). Although the number of black rats in the UK is thought to be low, it is possible there are extant small island populations, which may pose a risk to the surrounding ecosystem.

AHVLA. 2012. "Schmallenberg virus: results of UK testing." www.defra.gov.uk/ahvla/2012/12/14/schmallenberg-virus-updated-testing-results-figures.

al-Doory, Y., T.E. Vice, and M.E. Mainster. 1971. "Adiaspiromycosis in a dog." *Journal of the American Veterinary Medical Association* 59:87-90.

Allen, D., J. Davoile, A. Nobajas, S. Pemberton, D. Webb, and L. Wright. 2020. "Fenced fisheries, Eurasian otters (*Lutra lutra*) and licenced trapping: an impact assessment." *IUCN Otter Specialist Group Bulletin* 37 (1):38-52.

Almeida, David, Nahuel Rodolfo, Carl D. Sayer, and Gordon H. Copp. 2013. "Seasonal use of ponds as foraging habitat by Eurasian otter with description of an alternative handling technique for common toad predation." *Folia Zoologica* 62 (3):214-221. doi: 10.25225/fozo.v62.i3.a7.2013.

Altringham, J.D. 2003. *New Naturalist: British Bats*: HarperCollins.

Ansell, R., P. Baker, and S. Harris. 2001. "The value of gardens for wildlife – lessons from mammals and herpetofauna." *British Wildlife* 13:1205-1215.

Arden, K., K. Gedye, O. Angelin-Bonnet, E. Murphy, and D. Antic. 2022. "Yersinia enterocolitica in wild and peridomestic rodents within Great Britain, a prevalence study." *Zoonoses Public Health* 69 (5):537-549. doi: 10.1111/zph.12943.

Armstrong, J.D., H.L. Anderson, S.L. Dugan, and R. Gardiner. 2015. A mapping study of the overlap of potential Eurasian beaver (*Castor fiber*) habitat and Atlantic Salmon (*Salmo salar*) distribution in Scotland. In *Scottish Marine and Freshwater Science*.

Arnold, J.M., P. Gerhardt, S.M.J.G. Steyaert, E. Hochbichler, and K. Hackländer. 2018. "Diversionary feeding can reduce red deer habitat selection pressure on vulnerable forest stands, but is not a panacea for red deer damage." *Forest Ecology and Management* 407:166–173. doi: https://doi.org/10.1016/j.foreco.2017.10.050.

Atkinson, R.P.D., D.W. Macdonald, and P.J. Johnson. 1994. "The status of the European Mole *Talpa europaea* L. as an agricultural pest and its management." *Mammal Review,* 24:73-90.

Avanzi, C., J. Del-Pozo, A. Benjak, K. Stevenson, V. R. Simpson, P. Busso, J. McLuckie, C. Loiseau, C. Lawton, J. Schoening, D. J. Shaw, J. Piton, L. Vera-Cabrera, J. S. Velarde-Felix, F. McDermott, S. V. Gordon, S. T. Cole, and A. L. Meredith. 2016. "Red squirrels in the British Isles are infected with leprosy bacilli." *Science* 354 (6313):744-747. doi: 10.1126/science.aah3783.

Azagi, T., D. Hoornstra, K. Kremer, J. W. R. Hovius, and H. Sprong. 2020. "Evaluation of Disease Causality of Rare Ixodes ricinus-Borne Infections in Europe." *Pathogens* 9 (2). doi: 10.3390/pathogens9020150.

Bacigalupo, S. A., L. K. Dixon, S. Gubbins, A. J. Kucharski, and J. A. Drewe. 2022. "Wild boar visits to commercial pig farms in southwest England: implications for disease transmission." *Eur J Wildl Res* 68 (6):69. doi: 10.1007/s10344-022-01618-2.

Baerwald, E. F., G. H. D'Amours, B. J. Klug, and R. M. Barclay. 2008. "Barotrauma is a significant cause of bat fatalities at wind turbines." *Curr Biol* 18 (16):R695-6. doi: 10.1016/j.cub.2008.06.029.

Bailey, J.R., L. Warner, G.C. Pritchard, S. Williamson, T. Carson, G. Willshaw, and T. Cheasty. 2002. "Wild rabbits--a novel vector for Vero cytotoxigenic *Escherichia coli* (VTEC) O157." *Communicable Disease and Public Health* 5:74-75.

Baines, David, Nicholas J. Aebischer, and Allan Macleod. 2016. "Increased mammalian predators and climate ch1ange predict declines in breeding success and density of Capercaillie tetrao urogallus, an old stand specialist, in fragmented Scottish forests." *Biodiversity and Conservation* 25 (11):2171-2186. doi: 10.1007/s10531-016-1185-8.

Baines, David, Helen Allinson, James Paul Duff, Harriet Fuller, David Newborn, and Michael Richardson. 2018. "Lethal and sub-lethal impacts of respiratory cryptosporidiosis on Red Grouse, a wild gamebird of economic importance." *Ibis* 160 (4):882-891. doi: 10.1111/ibi.12573.

Baines, R., S. Baker, J. Hallett, and D. Macdonald. 1995. The impact of foxes and fox hunting on the management of Wiltshire county farms estate. Report to Wiltshire County Council, Occasional Paper No. 22, Royal Agricultural College & University of Oxford, Oxford.

Baker, P., S. Harris, and P.C.L. White. 2006. After the hunt: the future for foxes in Britain. London: International Fund for Animal Welfare; https://mknhs.org.uk/wp-content/uploads/2015/08/IFAW-after-the-hunt.pdf.

Baker, S. E., S. A. Ellwood, P. J. Johnson, and D. W. Macdonald. 2016. "Moles and Mole Control on British Farms, Amenities and Gardens after Strychnine Withdrawal." *Animals (Basel)* 6 (6). doi: 10.3390/ani6060039.

Baker, Sandra E., and David W. Macdonald. 2000. "Foxes and foxhunting on farms in Wiltshire: a case study." *Journal of Rural Studies* 16 (2):185-201. doi: 10.1016/s0743-0167(99)00051-0.

Ball, C. 2014. "Leptospirosis in UK vet visiting dogs, wild rodents and the pathogenomics of Leptospira species."PhD thesis, University of Liverpool.

Bamber, Jack A., Craig M. Shuttleworth, Matthew W. Hayward, and David J. Everest. 2020. "Reinstating trophic cascades as an applied conservation tool to protect forest ecosystems from invasive grey squirrels (Sciurus carolinensis)." *Food Webs* 25. doi: 10.1016/j.fooweb.2020.e00164.

Bank of England. 2024. http://www.bankofengland.co.uk. Date accessed: 26th November 2024.

Barlow, A., P. Green, T. Banham, and N. Healy. 2013. "Serological confirmation of SBV infection in wild British deer." *Vet Rec* 172 (16):429. doi: 10.1136/vr.f2438.

Barlow, A. M., L. Worledge, H. Miller, K. P. Drees, P. Wright, J. T. Foster, C. Sobek, A. M. Borman, and M. Fraser. 2015. "First confirmation of *Pseudogymnoascus destructans* in British bats and hibernacula." *Vet Rec* 177 (3):73. doi: 10.1136/vr.102923.

Barlow, A., and L. Mullineaux. 2018. "Zoonotic disease in a European brown hare in England." *Vet Rec* 183 (5):163. doi: 10.1136/vr.k3316.

Barreto, Guillermo R., Steve P. Rushton, Rob Strachan, and David W. Macdonald. 2006. "The role of habitat and mink predation in determining the status and distribution of water voles in England." *Animal Conservation* 1 (2):129-137. doi: 10.1111/j.1469-1795.1998.tb00020.x.

Battersby, J. 2005. UK Mammals: Species status and population trends. First report by the Tracking Mammals Partnership. JNCC/Tracking Mammals Partnership, Peterborough.

Battisti, E., S. Zanet, S. Khalili, A. Trisciuoglio, B. Hertel, and E. Ferroglio. 2020. "Molecular Survey on Vector-Borne Pathogens in Alpine Wild Carnivorans." *Front Vet Sci* 7:1. doi: 10.3389/fvets.2020.00001.

BBC. 2015a. "Moving 55 water voles from Somerset Levels cost £135K. ." https://www.bbc.co.uk/news/uk-england-somerset-32127454.

BBC. 2015b. "'Wild boar' causes Wiltshire M4 fatal crash. https://www.bbc.co.uk/news/uk-england-wiltshire-30691090."

BBC. 2023a. "Llandudno: Great Orme goats to be moved after damaging property.". https://www.bbc.co.uk/news/uk-wales-65353062.

BBC. 2023b. "Mendips: Illegal poachers cause 'extensive damage' to farmland. https://www.bbc.co.uk/news/uk-england-somerset-67100632."

Beaumont, M., E. M. Barratt, D. Gottelli, A. C. Kitchener, M. J. Daniels, J. K. Pritchard, and M. W. Bruford. 2001. "Genetic diversity and introgression in the Scottish wildcat." *Mol Ecol* 10 (2):319-36. doi: 10.1046/j.1365-294x.2001.01196.x.

Becker, S. D., M. Bennett, J. P. Stewart, and J. L. Hurst. 2007. "Serological survey of virus infection among wild house mice (Mus domesticus) in the UK." *Lab Anim* 41 (2):229-38. doi: 10.1258/002367707780378203.

Bell, D. J., J. P. Davis, M. Gardner, A. M. Barlow, M. Rocchi, M. Gentil, and R. J. Wilson. 2019. "Rabbit haemorrhagic disease virus type 2 in hares in England." *Vet Rec* 184 (4):127-128. doi: 10.1136/vr.l337.

Bell, E., D. Boyle, K. Floyd, P. Garner-Richards, B. Swann, R. Luxmoore, A. Patterson, and R. Thomas. 2011. "The ground-based eradication of Norway rats (*Rattus norvegicus*) from the Isle of Canna, Inner Hebrides, Scotland." In *Island invasives: eradication and management*, edited by C.R.C. Veitch, M.N. Clout and D.R. Towns, 269-274. Switzerland: IUCN Gland.

Bell, E., K. Floyd, D. Boyle, J. Pearson, L. Lock, P. Buckley, S. Mason, R. McCarthy, W. Garratt, K. Sugar, and J. Pearce. 2019. "The Isles of Scilly seabird restoration project: the eradication of brown rats (*Rattus norvegicus*) from the inhabited islands of St Agnes and Gugh, Isles of Scilly." In *Island invasives: scaling up to meet the challenge*, edited by C.R. Veitch, M.N. Clout, A.R. Martin, J.C. Russell and C.J. West, 88-94. Occasional paper SSC no. 62., IUCN Gland, Switzerland.

Berry, R.J. 1991. "House Mouse *Mus domesticus*." In *Handbook of British Mammals*, edited by G.B. Corbet and S. Harris, 239-247. Blackwell Scientific, Oxford.

Birks, J.D.S. 2000. "The recovery of the polecat, *Mustela putorius*, in Britain." In *Mustelids in a modern world: Management and conservation aspects of small carnivore: human interaction*, edited by H.I. Griffiths. Backhuys Publishers, Leiden, The Netherlands.

Birks, J.D.S., and A.C. Kitchener. 1999. The distribution and status of the polecat, *Mustela putorius*, in Britain in the 1990s. Vincent Wildlife Trust, London.

Birks, J.D.S., J.E. Messenger, and E.C. Halliwell. 2005. "Diversity of den sites used by pine martens *Martes martes*: a response to the scarcity of aboreal cavities?" *Mammal Review* 35:313-320.

Blackett, T. A., V. R. Simpson, S. Haugland, D. J. Everest, C. F. Muir, K. C. Smith, and A. C. Mill. 2018. "Mortalities, amyloidosis and other diseases in free-living red squirrels (Sciurus vulgaris) on Jersey, Channel Islands." *Vet Rec* 183 (16):503. doi: 10.1136/vr.104779.

Bochynska, D., S. Lloyd, O. Restif, and K. Hughes. 2022. "Eimeria stiedae causes most of the white-spotted liver lesions in wild European rabbits in Cambridgeshire, United Kingdom." *J Vet Diagn Invest* 34 (2):199-205. doi: 10.1177/10406387211066923.

Bond, Ian, Emily Gilford, Allan McDevitt, Melissa Young, and Frazer Coomber. 2022. "First records of the greater white-toothed shrew Crocidura russula from Great Britain." *Mammal Communications* 8. doi: 10.59922/aaci6532.

Bonesi, Laura, and Santiago Palazon. 2007. "The American mink in Europe: Status, impacts, and control." *Biological Conservation* 134 (4):470-483. doi: 10.1016/j.biocon.2006.09.006.

Bonnington, C., K. J. Gaston, and K. L. Evans. 2014. "Relative roles of grey squirrels, supplementary feeding, and habitat in shaping urban bird assemblages." *PLoS One* 9 (10):e109397. doi: 10.1371/journal.pone.0109397.

Bonnington, Colin, Kevin J. Gaston, Karl L. Evans, and Colleen Downs. 2013. "Assessing the potential for Grey Squirrels Sciurus carolinensis to compete with birds at supplementary feeding stations." *Ibis* 156 (1):220-226. doi: 10.1111/ibi.12107.

Borman, A. M., V. R. Simpson, M. D. Palmer, C. J. Linton, and E. M. Johnson. 2009. "Adiaspiromycosis due to Emmonsia crescens is widespread in native British mammals." *Mycopathologia* 168 (4):153-63. doi: 10.1007/s11046-009-9216-6.

Bright, P., Morris, P., and T. Mitchell-Jones. 2006. *The dormouse conservation handbook*: English Nature.

Bright, P.W., and E.C. Halliwell. 1999. Species recovery programme for the pine marten in England: 1996-1998. In *Research Reports No. 306, English Nature, Peterborough*.

British Deer Society. 2023a. "Muntjac deer." https://bds.org.uk/information-advice/about-deer/deer-species/muntjac-deer/. Date accessed: 11th November 2024.

British Deer Society. 2023b. "Roe deer." https://bds.org.uk/information-advice/about-deer/deer-species/roe-deer/. Date accessed: 30th April 2024.

British Wildlife Centre. 2024. "Black rat – *Rattus rattus*." https://britishwildlifecentre.co.uk/planyourvisit/animals/black-rat/.

Britton, J. R., J. S. Shepherd, S. Toms, and V. Simpson. 2005. "Presence of carp, *Cyprinus carpio*, in the diet of the otter, *Lutra lutra*." *Fisheries Management and Ecology* 12 (3):221-223. doi: 10.1111/j.1365-2400.2005.00430.x.

Brookshire, J. E., B. J. Kauffman, D. Lytjen, and N. Otting. 2002. "Cumulative effects of wild ungulate and livestock herbivory on riparian willows." *Oecologia* 132 (4):559-566. doi: 10.1007/s00442-002-1007-4.

Brown, D., J. Wilson, D. Douglas, P. Thompson, S. Foster, N. McCulloch, J. Phillips, D. Stroud, S. Whitehead, N. Crockford, and R. Sheldon. 2015. "The Eurasian Curlew – the most pressing bird conservation priority in the UK?" *British Birds* 108:660-668.

Brown, H., and J.D.S. Birks. 2006. "Resolving conflicts generated by pine martens denning in buildings in Scotland." In, edited by M. Santos-Reis, J.D.S. Birks, E.C. O'Doherty and G Proulx, 125-133. Alpha Wildlife Publications, Sherwood Park, Alberta, Canada.

Bruemmer, C. M., S. P. Rushton, J. Gurnell, P. W. Lurz, P. Nettleton, A. W. Sainsbury, J. P. Duff, J. Gilray, and C. J. McInnes. 2010. "Epidemiology of squirrelpox virus in grey squirrels in the UK." *Epidemiol Infect* 138 (7):941-50. doi: 10.1017/S0950268810000816.

Brzeziński, Marcin, Aleksandra Zarzycka, Tom A. Diserens, and Andrzej Zalewski. 2021. "Does the American mink displace the European polecat? A need for more research on interspecific competition between invasive and native species." *European Journal of Wildlife Research* 67 (4). doi: 10.1007/s10344-021-01511-4.

Büchner, Sven, Roger Trout, and Peter Adamík. 2018. "Conflicts with Glis glis and Eliomys quercinus in households: a practical guideline for sufferers (Rodentia: Gliridae)." *Lynx, new series* 49 (1):19-26. doi: 10.2478/lynx-2018-0003.

Buckle, A. 2007. "Rat control 1: Problems caused on farms, The Campaign for Responsible Rodenticide Use (CRRU)." Farmers Weekly. http://www.fwi.co.uk/academy/article/106448/rat-control-2-problems-caused-on-farms.html.

Burrells, A., P. M. Bartley, I. A. Zimmer, S. Roy, A. C. Kitchener, A. Meredith, S. E. Wright, E. A. Innes, and F. Katzer. 2013. "Evidence of the three main clonal Toxoplasma gondii lineages from wild mammalian carnivores in the UK." *Parasitology* 140 (14):1768-76. doi: 10.1017/S0031182013001169.

Calladine, John, Elizabeth M. Humphreys, Lucy Gilbert, Robert W. Furness, Robert A. Robinson, Robert J. Fuller, Nick A. Littlewood, Robin J. Pakeman, Johanne Ferguson, and Charles Thompson. 2017. "Continuing influences of introduced hedgehogs Erinaceus europaeus as a predator of wader (Charadrii) eggs four decades after their release on the Outer Hebrides, Scotland." *Biological Invasions* 19 (7):1981-1987. doi: 10.1007/s10530-017-1422-4.

Campbell-Palmer, R., A. Puttock, R.N. Needham, K. Wilson, H. Graham, and R.E. Brazier. 2021. "Survey of the Tayside Area Beaver Population 2020-2021. NatureScot Research Report 1274."

Campbell-Palmer, R., G. Schwab, S. Girling, S. Lisle, and D. Gow. 2015. Managing wild Eurasian beavers: a review of European management practices with consideration for Scottish application. Scottish Natural Heritage Commissioned Report No. 812, Inverness, UK.

Campbell, R.D., A. Dutton, and J. Hughes. 2007. Economic impacts of the beaver. Report for the Wild Britain Initiative.

Caravaggi, A., M. Gatta, M. C. Vallely, K. Hogg, M. Freeman, E. Fadaei, J. T. A. Dick, W. I. Montgomery, N. Reid, and D. G. Tosh. 2018. "Seasonal and predator-prey effects on circadian activity of free-ranging mammals revealed by camera traps." *PeerJ* 6:e5827. doi: 10.7717/peerj.5827.

Carrs, D.N., H. Kruuk, and J.W.H. Conroy. 1990. "Predation on adult Atlantic salmon, *Salmo salar* L., by otters, *Lutra lutra* (L.), within the River Dee system, Aberdeenshire, Scotland " *Journal of Fish Biology* 37:935-944.

Carss, D.N. 1995. "Foraging behaviour and feeding ecology of the otter Lutra lutra: A selective review." *Hystrix (n.s.)* 7:179-194. doi: 10.4404/hystrix-7.1-2-4069.

Carter, N.A. 1984. "Bole scoring by sika deer (*Cervus nippon*) in England." *Deer* 6:77-78.

Chalmers, R. M., G. Robinson, K. Elwin, S. J. Hadfield, L. Xiao, U. Ryan, D. Modha, and C. Mallaghan. 2009. "*Cryptosporidium* sp. rabbit genotype, a newly identified human pathogen." *Emerg Infect Dis* 15 (5):829-30. doi: 10.3201/eid1505.081419.

Chanin, P. R. F., and D. J. Jefferies. 1978. "The decline of the otter Lutra lutra L. in Britain: an analysis of hunting records and discussion of causes." *Biological Journal of the Linnean Society* 10 (3):305-328. doi: 10.1111/j.1095-8312.1978.tb00018.x.

Chanin, P., and M. Woods. 2003. Surveying dormice using nest tubes: Research Report 524. Peterborough.

Chantrey, J., T. D. Dale, J. M. Read, S. White, F. Whitfield, D. Jones, C. J. McInnes, and M. Begon. 2014. "European red squirrel population dynamics driven by squirrelpox at a gray squirrel invasion interface." *Ecol Evol* 4 (19):3788-99. doi: 10.1002/ece3.1216.

Chapman, Roselle E., and Andrew F. G. Bourke. 2008. "The influence of sociality on the conservation biology of social insects." *Ecology Letters* 4 (6):650-662. doi: 10.1046/j.1461-0248.2001.00253.x.

Chappell, J. G., T. Tsoleridis, O. Onianwa, G. Drake, I. Ashpole, P. Dobbs, W. Edema, F. Kumi-Ansah, M. Bennett, R. E. Tarlinton, J. K. Ball, and C. P. McClure. 2020. "Retrieval of the Complete Coding Sequence of the UK-Endemic Tatenale Orthohantavirus Reveals Extensive Strain Variation and Supports Its Classification as a Novel Species." *Viruses* 12 (4). doi: 10.3390/v12040454.

Charlier, J., J. Vercruysse, E. Morgan, J. van Dijk, and D. J. Williams. 2014. "Recent advances in the diagnosis, impact on production and prediction of Fasciola hepatica in cattle." *Parasitology* 141 (3):326-35. doi: 10.1017/S0031182013001662.

Cheng, T. L., J. D. Reichard, J. T. H. Coleman, T. J. Weller, W. E. Thogmartin, B. E. Reichert, A. B. Bennett, H. G. Broders, J. Campbell, K. Etchison, D. J. Feller, R. Geboy, T. Hemberger, C. Herzog, A. C. Hicks, S. Houghton, J. Humber, J. A. Kath, R. A. King, S. C. Loeb, A. Masse, K. M. Morris, H. Niederriter, G. Nordquist, R. W. Perry, R. J. Reynolds, D. B. Sasse, M. R. Scafini, R. C. Stark, C. W. Stihler, S. C. Thomas, G. G. Turner, S. Webb, B. J. Westrich, and W. F. Frick. 2021. "The scope and severity of white-nose syndrome on hibernating bats in North America." *Conserv Biol* 35 (5):1586-1597. doi: 10.1111/cobi.13739.

Chintoan-Uta, C., E.R. Morgan, P.J. Skuce, and G.C. Coles. 2014. "Wild deer as potential vectors of anthelmintic-resistant abomasal nematodes between cattle and sheep farms." *Proc Biol Sci* 281 (1780):20132985. doi: 10.1098/rspb.2013.2985.

Churchfield, S. 2008. "Insectivores: Orders Erinaceomorpha and Soricomorpha." In *Mammals of the British Isles (4th edn.).* edited by S. Harris and D.W. Yalden, 240-291. Southampton: The Mammal Society.

CIEH. 2013. "Urban foxes: Guidelines on their management." Chartered Institute of Environmental Health. http://www.cieh.org/WorkArea/showcontent.aspx?id=49460.

Clutton-Brock, T.H., and S.D. Albon. 1989. *Red deer in the highlands*. Oxford: BSP Professional Books.

Collard, K.J. 2023. "A study of the incidence of bovine tuberculosis in the wild red deer herd of Exmoor." *Eur J Wildl Res* 69 (1):14. doi: 10.1007/s10344-022-01638-y.

Common, S., H. Donald, and A. Sainsbury. 2024. Revised Disease Risk Analysis for the conservation translocation of the Eurasian beaver (*Castor fiber*) to England [NECR540]. Natural England.

Cooke, A. . 2019. *Muntjac and water deer: natural history, environmental impact and management*. Exeter, UK: Pelagic Publishing.

Cooke, A.S., and L. Farrell. 1987. The utilisation of neighbouring farmland by Chinese water deer (*Hydropotes inermis*) at Woodwalton Fen National Nature Reserve. Huntingdonshire Fauna & Flora Society Report 1986.

Cooke, A.S., and L. Farrell. 2001. "Impact of Muntjac deer (*Muntiacus reevesi*) at Monks Wood National Nature Reserve, Cambridgeshire, eastern England." *Forestry* 74:241-250.

Cooke, A.S., P. Green, and N.G. Chapman. 1996. "Mortality in a feral population of Muntjac *Muntiacus reevesi* in England." *Acta Theriologica* 41:277-286.

Cooke, Arnold S. 2021. "Colonisation, impacts in conservation woodland and management of Reeves' muntjac (*Muntiacus reevesi*) in an English county." *European Journal of Wildlife Research* 67 (3). doi: 10.1007/s10344-021-01480-8.

Corman, V. M., N. L. Ithete, L. R. Richards, M. C. Schoeman, W. Preiser, C. Drosten, and J. F. Drexler. 2014. "Rooting the phylogenetic tree of middle East respiratory syndrome coronavirus by characterization of a conspecific virus from an African bat." *J Virol* 88 (19):11297-303. doi: 10.1128/JVI.01498-14.

Costa, F., J. E. Hagan, J. Calcagno, M. Kane, P. Torgerson, M. S. Martinez-Silveira, C. Stein, B. Abela-Ridder, and A. I. Ko. 2015. "Global Morbidity and Mortality of Leptospirosis: A Systematic Review." *PLoS Negl Trop Dis* 9 (9):e0003898. doi: 10.1371/journal.pntd.0003898.

Countryfile. 2024. "Deer culling in Britain: what's the problem and why are deer culled? ." https://www.countryfile.com/wildlife/mammals/deer-culling-in-britain. Date accessed: 22nd October 2024.

Cowan, D.P. 2008. "Order Lagomorpha: Rabbits and Hares." In *Mammals of the British Isles (4th edn.)*, edited by S. Harris and D.W Yalden. Southampton: The Mammal Society.

Coz, Deborah M., Juliette C. Young, and Leah Gibbs. 2020. "Conflicts over wildlife conservation: Learning from the reintroduction of beavers in Scotland." *People and Nature* 2 (2):406-419. doi: 10.1002/pan3.10076.

Craik, J.C.A. 1990. "The price of mink." *Scottish Bird News* 19:4-5.

Craik, J.C.A. 1991. "More serious than Shetland?" *Seabird Group Newsletter* 60:2-4.

Craik, J.C.A. 1997. "Long-term effects of North American Mink *Mustela vison* on seabirds in western Scotland." *Bird Study* 44:303-309.

Crawford, A. 2003. Fourth Otter Survey of England 2000-2002. The Environment Agency, UK.

Crawley, D., F. Coomber, L. Kubasiewicz, C. Harrower, P. Evans, J. Waggitt, B. Smith, and F. Mathews. 2020. *Atlas of the mammals of Great Britain and Northern Ireland*. The Mammal Society: Pelagic Publishing, UK.

CSL. 2006. A quantitative risk assessment on the role of wild deer in the perpetuation of TB in cattle. In *Report to the Department of Environment, Food and Rural Affairs, Project Code: SE3036*.

Davidson, J. A., M. G. Loutet, C. O'Connor, C. Kearns, R. M. Smith, M. K. Lalor, H. L. Thomas, I. Abubakar, and D. Zenner. 2017. "Epidemiology of Mycobacterium bovis Disease in Humans in England, Wales, and Northern Ireland, 2002-2014." *Emerg Infect Dis* 23 (3):377-386. doi: 10.3201/eid2303.161408.

Davies, R. H., and A. D. Wales. 2013. "Salmonella contamination of cereal ingredients for animal feeds." *Vet Microbiol* 166 (3-4):543-9. doi: 10.1016/j.vetmic.2013.07.003.

Davies, Zoe G., Richard A. Fuller, Alison Loram, Katherine N. Irvine, Victoria Sims, and Kevin J. Gaston. 2009. "A national scale inventory of resource provision for biodiversity within domestic gardens." *Biological Conservation* 142 (4):761-771. doi: 10.1016/j.biocon.2008.12.016.

Davison, John, Timothy J. Roper, Charles J. Wilson, Matthew J. Heydon, and Richard J. Delahay. 2010. "Assessing spatiotemporal associations in the occurrence of badger–human conflict in England." *European Journal of Wildlife Research* 57 (1):67-76. doi: 10.1007/s10344-010-0400-2.

de la Rua-Domenech, R. 2006. "Human Mycobacterium bovis infection in the United Kingdom: Incidence, risks, control measures and review of the zoonotic aspects of bovine tuberculosis." *Tuberculosis (Edinb)* 86 (2):77-109. doi: 10.1016/j.tube.2005.05.002.

Deer Aware. 2019. "The deer on our roads website.". www.deeraware.com. Date accessed: 22nd October 2024.

Deer Collisions. 2023. "National deer-vehicle collisions project. ." www.deercollisions.co.uk. Date accessed: 22nd October 2024.

Defra. 2008. Feral wild boar in England: an action plan. London: Department for Environment, Food and Rural Affairs.

Defra. 2024. "Government to end badger cull with new TB eradication strategy.". https://www.gov.uk/government/news/government-to-end-badger-cull-with-new-tb-eradication-strategy.

Delahay, R. J., J. Davison, D. W. Poole, A. J. Matthews, C. J. Wilson, M. J. Heydon, and T. J. Roper. 2009. "Managing conflict between humans and wildlife: trends in licensed operations to resolve problems with badgers Meles meles in England." *Mammal Review* 39 (1):53-66. doi: 10.1111/j.1365-2907.2008.00135.x.

Delahay, R. J., A. N. De Leeuw, A. M. Barlow, R. S. Clifton-hadley, and C. L. Cheeseman. 2002. "The status of Mycobacterium bovis infection in UK wild mammals: a review." *Vet J* 164 (2):90-105. doi: 10.1053/tvjl.2001.0667.

Delahay, R. J., G. C. Smith, A. M. Barlow, N. Walker, A. Harris, R. S. Clifton-Hadley, and C. L. Cheeseman. 2007. "Bovine tuberculosis infection in wild mammals in the South-West region of England: a survey of prevalence and a semi-quantitative assessment of the relative risks to cattle." *Vet J* 173 (2):287-301. doi: 10.1016/j.tvjl.2005.11.011.

Dell'Arte, Graziella Lucia, Toni Laaksonen, Kai Norrdahl, and Erkki Korpimäki. 2007. "Variation in the diet composition of a generalist predator, the red fox, in relation to season and density of main prey." *Acta Oecologica* 31 (3):276-281. doi: 10.1016/j.actao.2006.12.007.

DfT. 2011. Reported Road Casualties in Great Britain: 2011. Annual Report.

DfT. 2024. National Travel Survey.

Dolman, P., R. Fuller, R. Gill, D. Hooton, and R. Tabor. 2010. "Escalating ecological impacts of deer in lowland woodland." *British Wildlife* 21:242–254.
Doney, J., and J.J. Packer. 2002. "The impact of deer on agriculture: results of a questionnaire survey and subsequent validation." *Deer* 12:98-104.
Donnelly, C. A., and P. Nouvellet. 2013. "The contribution of badgers to confirmed tuberculosis in cattle in high-incidence areas in England." *PLoS Curr* 5. doi: 10.1371/currents.outbreaks.097a904d3f3619db2fe78d24bc776098.
Duscher, G.G., E. Battisti, A. Hodžić, K. Wäber, P. Steinbach, M. Stubbe, and M. Heddergott. 2020. "First detection and molecular identification of *Anaplasma phagocytophilum* in an introduced population of Reeve's muntjac (*Muntiacus reevesi*) in United Kingdom." *Molecular and Cellular Probes* 52:101582.
Dutton, J.S., H.T. Clayton, and S.M. Evans. 2015. The social aspects of wild boar in the Forest of Dean. Unpublished report for the Forestry Commission by the University of Worcester.
Eschen, R., M. Kadzamira, S. Stutz, A. Ogunmodede, D. Djeddour, R. Shaw, C. Pratt, S. Varia, K. Constantine, and F. Williams. 2023. "An updated assessment of the direct costs of invasive non native species to the United Kingdom." *Biological Invasions* 25:3265-76. doi: https://doi.org/10.1007/s10530-023-03107-2.
Etherington, G. J., A. Ciezarek, R. Shaw, J. Michaux, E. Croose, W. Haerty, and F. Di Palma. 2022. "Extensive genome introgression between domestic ferret and European polecat during population recovery in Great Britain." *J Hered* 113 (5):500-515. doi: 10.1093/jhered/esac038.
Everest, D. J., S. S. Grierson, M. F. Stidworthy, and C. Shuttleworth. 2009. "PCR detection of adenovirus in grey squirrels on Anglesey." *Vet Rec* 165 (16):482. doi: 10.1136/vr.165.16.482.
Everest, D.J., J.P. Holmes, and C. Shuttleworth. 2017. "Virus presence within Welsh red squirrel." *Veterinary Record* 180:520–521.
Everest, David J., Craig M. Shuttleworth, Sylvia S. Grierson, Akbar Dastjerdi, Mark F. Stidworthy, J. Paul Duff, Robert J. Higgins, Aileen Mill, and Julian Chantrey. 2018. "The implications of significant adenovirus infection in UK captive red squirrel (Sciurus vulgaris) collections: How histological screening can aid applied conservation management." *Mammalian Biology* 88:123-129. doi: 10.1016/j.mambio.2017.10.003.
Ewins, P.J. 1985. "Otter predation on black guillemots." *British Birds* 78:663-664.
Farmers Weekly. 2024. "Farm fire costs hit £110M in 2023." https://www.fwi.co.uk/news/farm-fire-costs-hit-110m-in-2023.
FAWC. 2013. "Opinion on the welfare of farmed and park deer. Farm and Animal Welfare Committee." www.defra.gov.uk/fawc.
Fenn, P. 2002. An investigation of the issues raised for the construction industry in the United Kingdom by bats. Research Papers. 4(11) RICS Foundation.
Ferreras, P., and D.W. MacDonald. 1999. "The impact of American mink *Mustela vison* on water birds in the upper Thames." *Journal of Applied Ecology* 36 (5):701-708.
Fietz, J., J. Tomiuk, F. R. Matuschka, and D. Richter. 2014. "Seasonal prevalence of Lyme disease spirochetes in a heterothermic mammal, the edible dormouse (Glis glis)." *Appl Environ Microbiol* 80 (12):3615-21. doi: 10.1128/AEM.00251-14.
Fingland, Kathryn, Samantha J. Ward, Adam J. Bates, and Samantha Bremner-Harrison. 2021. "A systematic review into the suitability of urban refugia for the Eurasian red squirrel *Sciurus vulgaris*." *Mammal Review* 52 (1):26-38. doi: 10.1111/mam.12264.
Fisher, M. 2003. "*Toxocara cati*: an underestimated zoonotic agent." *Trends Parasitol* 19 (4):167-70. doi: 10.1016/s1471-4922(03)00027-8.
Fletcher, Kathy, Nicholas J. Aebischer, David Baines, Robin Foster, and Andrew N. Hoodless. 2010. "Changes in breeding success and abundance of ground-nesting moorland birds in relation to the experimental deployment of legal predator control." *Journal of Applied Ecology* 47 (2):263-272. doi: 10.1111/j.1365-2664.2010.01793.x.
Fletcher, Kathy, Andrew N. Hoodless, and David Baines. 2013. "Impacts of predator abundance on red grouse Lagopus lagopus scotica during a period of experimental predator control." *Wildlife Biology* 19 (3):248-256. doi: 10.2981/12-009.

Fletcher, T.J. 2002. Diseases of deer relevant to Scotland. Contract report to the Deer Commission for Scotland. Inverness.

Flux, John E.C. 2009. "Life history of the Mountain hare (Lepus timidus scoticus) in north-east Scotland." *Journal of Zoology* 161 (1):75-123. doi: 10.1111/j.1469-7998.1970.tb02171.x.

Forestry and Land Scotland. 2021. "FLS launch five-year deer management tender." https://forestryandland.gov.scot/news-releases/fls-launch-five-year-deer-management-tender/. Date accessed: 30th April 2024.

Forestry Commission. 2023. "Grey squirrel and deer damage to trees: Visual guide." https://assets.publishing.service.gov.uk/media/67337357c10bb403d96bf2fa/WMM_leaflet_Nov_24_WEB.pdf.

Foster, W.N., and J.C. Greig. 1969. "Isolation of tick-borne fever from feral goats in New Galoway." *Vet Rec* 85 (21):585-6. doi: 10.1136/vr.85.21.585.

Fountain, K., T. Blackett, H. Butler, C. Carchedi, A. K. Schilling, A. Meredith, M. J. Gibbon, D. H. Lloyd, A. Loeffler, and E. J. Feil. 2021. "Fatal exudative dermatitis in island populations of red squirrels (*Sciurus vulgaris*): spillover of a virulent *Staphylococcus aureus* clone (ST49) from reservoir hosts." *Microb Genom* 7 (5). doi: 10.1099/mgen.0.000565.

Francksen, Richard M., Mark J. Whittingham, Sonja C. Ludwig, Staffan Roos, and David Baines. 2017. "Numerical and functional responses of Common Buzzards *Buteo buteo* to prey abundance on a Scottish grouse moor." *Ibis* 159 (3):541-553. doi: 10.1111/ibi.12471.

Fraser, E.J., X. Lambin, R.A. McDonald, and S.M. Redpath. 2015. Stoat (*Mustela erminea*) on the Orkney Islands - assessing risks to native species. Scottish Natural Heritage Commissioned Report No. 871.

Fuehrer, H.P., P. Igel, and H. Auer. 2011. "Capillaria hepatica in man--an overview of hepatic capillariosis and spurious infections." *Parasitol Res* 109 (4):969-79. doi: 10.1007/s00436-011-2494-1.

Fulton, N., L.F. Anderson, J.M. Watson, and I. Abubakar. 2016. "Leprosy in England and Wales 1953-2012: surveillance and challenges in low incidence countries." *BMJ Open* 6 (5):e010608. doi: 10.1136/bmjopen-2015-010608.

Garigliany, M.M., C. Bayrou, D. Kleijnen, D. Cassart, S. Jolly, A. Linden, and D. Desmercht. 2012. "Schmallenberg virus: a new Shamonda/Sathuperi-like virus on the rise in Europe." *Antiviral Research* 95:82-87.

Gaywood, M., A. Stringer, D. Blake, J. Hall, M. Hennessy, A. Tree, D. Genney, I. Macdonald, A. Tonhasca, C. Bean, J. McKinnell, S. Cohen, R. Raynor, P. Watkinson, D. Bale, K. Taylor, J. Scott, and S. Blyth. 2015. Beavers in Scotland: a report to the Scottish Government, Edinburgh, Scottish Natural Heritage.

Gaywood, Martin J. 2017. "Reintroducing the Eurasian beaver Castor fiber to Scotland." *Mammal Review* 48 (1):48-61. doi: 10.1111/mam.12113.

Gecchele, L.V., A.B. Pedersen, and M. Bell. 2020. "Fine-scale variation within urban landscapes affects marking patterns and gastrointestinal parasite diversity in red foxes." *Ecol Evol* 10 (24):13796-13809. doi: 10.1002/ece3.6970.

Gelling, M., A.L. Harrington, M. Dean, E.C. Haddy, C.E. Marshall, and D.W. Macdonald. 2018. "The effect of using 'displacement' to encourage the movement of water voles *Arvicola amphibius* in lowland England." *Conservation Evidence* 15:20-25.

George, S.C., S. Campbell, C. Hale, J. Kaden, and L.I.M. Webster. 2024. Investigating the role of badgers (*Meles meles*) in lamb predation on Scottish farms 2022-23. NatureScot Research Report 1345.

Gern, L., and K. Sell. 2009. "Isolation of *Borrelia burgdorferi* sensu lato from the skin of the European badger (*Meles meles*) in Switzerland." *Vector Borne Zoonotic Dis* 9 (2):207-8. doi: 10.1089/vbz.2008.0050.

Gerrikagoitia, X., H. Gil, C. Garcia-Esteban, P. Anda, R.A. Juste, and M. Barral. 2012. "Presence of *Bartonella* species in wild carnivores of northern Spain." *Appl Environ Microbiol* 78 (3):885-8. doi: 10.1128/AEM.05938-11.

Gill, R.M.A. 1992. "A Review of Damage by Mammals in North Temperate Forests. 2. Small Mammals." *Forestry* 65 (3):281-308. doi: 10.1093/forestry/65.3.281.

Gill, R.M.A., J. Gurnell, and R.C. Trout. 1995. "Do woodland mammals threaten the development of new woods?" In *The ecology of woodland creation*, edited by R. Ferris-Kaan, 201-224. London: John Wiley & Sons.

Girling, Simon J., Adam Naylor, Mary Fraser, and Róisín Campbell-Palmer. 2019. "Reintroducing beavers *Castor fiber* to Britain: a disease risk analysis." *Mammal Review* 49 (4):300-323. doi: 10.1111/mam.12163.

Golding, M.E., G. Wu, R. Wilkie, E. Picard-Meyer, A. Servat, D.A. Marston, J.N. Aegerter, D.L. Horton, and L.M. McElhinney. 2024. "Investigating the emergence of a zoonotic virus: phylogenetic analysis of European bat lyssavirus 1 in the UK." *Virus Evol* 10 (1):veae060. doi: 10.1093/ve/veae060.

Goodwin, Cecily E. D., David J. Hodgson, Nida Al-Fulaij, Sallie Bailey, Steve Langton, and Robbie A. McDonald. 2017. "Voluntary recording scheme reveals ongoing decline in the United Kingdom hazel dormouse *Muscardinus avellanarius* population." *Mammal Review* 47 (3):183-197. doi: 10.1111/mam.12091.

Gordon, I.J. 1988. "Facilitation of Red Deer Grazing by Cattle and Its Impact on Red Deer Performance." *The Journal of Applied Ecology* 25 (1). doi: 10.2307/2403605.

Gorman, M. 2008. "Carnivores: Order Carnivora." In *Mammals of the British Isles (4th edn.)*, edited by S. Harris and D.W. Yalden, 392-525. Southampton: The Mammal Society.

Gorman, M.L., and R.D. Stone. 1990. *The natural history of moles*. London, UK: Christopher Helm.

Goulson, D., S. O'Connor, and K. J. Park. 2017. "Causes of colony mortality in bumblebees." *Animal Conservation* 21 (1):45-53. doi: 10.1111/acv.12363.

Goulson, Dave, Pippa Rayner, Bob Dawson, and Ben Darvill. 2010. "Translating research into action; bumblebee conservation as a case study." *Journal of Applied Ecology* 48 (1):3-8. doi: 10.1111/j.1365-2664.2010.01929.x.

GOV.UK. 2015. "Badgers: protection and licences. https://www.gov.uk/guidance/badgers-protection-surveys-and-licences#:~:text=defences%20or%20watercourses-,When%20you%27ll%20need%20a%20licence,possible%20to%20avoid%20affecting%20badgers."

GOV.UK. 2018. "Recreational angling puts £1.4bn into English economy." https://www.gov.uk/government/news/recreational-angling-puts-14bn-into-english-economy.

Graham, D.A., C. Gallagher, R.F. Carden, J.M. Lozano, J. Moriarty, and R. O'Neill. 2017. "A survey of free-ranging deer in Ireland for serological evidence of exposure to bovine viral diarrhoea virus, bovine herpes virus-1, bluetongue virus and Schmallenberg virus." *Ir Vet J* 70:13. doi: 10.1186/s13620-017-0091-z.

Grant, Kathryn R., and Lauren A. Harrington. 2015. "Fish selection by riverine Eurasian otters in lowland England." *Mammal Research* 60 (3):217-231. doi: 10.1007/s13364-015-0223-3.

Graves, H.B. 1984. "Behaviour and ecology of wild and feral swine (*Sus scrofa*)." *Journal of Animal Science of the Total Environment* 58:482-492.

Gray, A., P. Capewell, R. Zadoks, M.A. Taggart, A.S. French, F. Katzer, B.R. Shiels, and W. Weir. 2021. "Wild deer in the United Kingdom are a potential reservoir for the livestock parasite *Babesia divergens*." *Current Research in Parasitology & Vector-Borne Diseases* 1:100019. doi: https://doi.org/10.1016/j.crpvbd.2021.100019.

Green, P. 2013. "A report to the Exmoor moorland landscape partnership. Exmoor National Park Authority."

GreenKeeping. 2019. "This shows the level of damage badgers can do to a course." https://www.greenkeepingeu.com/shows-level-damage-badgers-can-course/.

Greenwood, A.G., and S. Sanchez. 2002. "Serological evidence of murine pathogens in wild grey squirrels (Sciurus carolinensis) in North Wales." *Vet Rec* 150 (17):543-6. doi: 10.1136/vr.150.17.543.

Guardone, L., V.V. Ebani, R. Verin, S. Nardoni, A. Consolazione, M. Bennett, and F. Mancianti. 2020. "Molecular Detection of Arthropod-Borne Pathogens in Eurasian Badgers (Meles meles) from the United Kingdom." *Animals (Basel)* 10 (3). doi: 10.3390/ani10030446.

Guedon, G. 1998. "Mole control in farming: how to avoid the use of strychnine." *Phytoma* 510:10-16.

Gurnell, J. 1987. *The natural history of squirrels*. London, UK: Christopher Helm.

Gurnell, J., T. Blackett, H. Butler, P.W.W. Lurz, L. Magris, and C.M. Shuttleworth. 2015. "British red squirrel strongholds: challenges for conservation." In *Red Squirrels: Ecology, Conservation & Management in Europe*, edited by C. Shuttleworth, Lurz, P. & Haywood, M. Woodbridge, Suffolk, UK: European Squirrel Initiative.

Gurnell, J., and E.J. Hare. 2008. "Rodents: Order Rodentia." In *Mammals of the British Isles*, edited by S. Harris and D.W. Yalden, 55-199. Southampton: The Mammal Society.
Gurnell, John, Luc A. Wauters, Peter W. W. Lurz, and Guido Tosi. 2004. "Alien species and interspecific competition: effects of introduced eastern grey squirrels on red squirrel population dynamics." *Journal of Animal Ecology* 73 (1):26-35. doi: 10.1111/j.1365-2656.2004.00791.x.
Halley, D., F. Rosell, and A. Saveljev. 2012. "Population and distribution of Eurasian beaver (*Castor fiber*)." *Baltic Forestry* 18 (1):168-175.
Halley, D.J., A.P. Saveljev, and F. Rosell. 2020. "Population and distribution of beavers *Castor fiber* and *Castor canadensis* in Eurasia." *Mammal Review* 51:1-24.
Halliwell, E. C., and D. W. Macdonald. 1996. "American mink Mustela vison in the upper Thames catchment: Relationship with selected prey species and den availability." *Biological Conservation* 76 (1):51-56. doi: 10.1016/0006-3207(95)00072-0.
Hammershoj, M., and T. Asferg. 2000. "Mink and polecats in Denmark: status, impacts and damage to poultry." *Mammal Review* 30:227-232.
Hanmer, Hugh J., Rebecca L. Thomas, and Mark D. E. Fellowes. 2018. "Introduced Grey Squirrels subvert supplementary feeding of suburban wild birds." *Landscape and Urban Planning* 177:10-18. doi: 10.1016/j.landurbplan.2018.04.004.
Hansson, L. 1985. "The food of bank voles, wood mice and yellow-necked mice." *Symposia of the Zoological Society of London* 55:141-168.
Hardstaff, J.L., G. Marion, M.R. Hutchings, and P.C.L. White. 2014. "Evaluating the tuberculosis hazard posed to cattle from wildlife across Europe." *Res Vet Sci* 97 Suppl:S86-93. doi: 10.1016/j.rvsc.2013.12.002.
Haring, V.C., B. Litz, J. Jacob, M. Brecht, M. Bauswein, J. Sehl-Ewert, M. Heroldova, C. Wylezich, D. Hoffmann, R.G. Ulrich, M. Beer, and F. Pfaff. 2024. "Detection of novel orthoparamyxoviruses, orthonairoviruses and an orthohepevirus in European white-toothed shrews." *Microb Genom* 10 (8). doi: 10.1099/mgen.0.001275.
Harris, S., W. Cresswell, P. Reason, and P. Cresswell. 1992. "An integrated approach to monitoring badger (*Meles meles*) population changes in Britain." In *Wildlife 2001: Population*, edited by D.R. McCullough and R.H. Barrett, 945–53. London: Elsevier Applied Science.
Harris, S., P. Morris, S. Wray, and D. Yalden. 1995. A review of British mammals: population estimates and conservation status of British mammals other than cetaceans. Peterborough: JNCC.
Harris, S., and D.W. Yalden. 2008. *Mammals of the British Isles: handbook 4th edition*. Southampton: The Mammal Society.
Harrison, A., K.J. Brown, and W.I. Montgomery. 2012. "*Anaplasma phagocytophilum* in feral goats in Northern Ireland." *Veterinary Record* 170:602-603.
Harrison, M.D.K., and R.G. Symes. 1989. "Economic damage by feral American mink (*Mustela vison*) in England and Wales." In *Mammals as pests*, edited by R.J. Putman. London: Chapman and Hall.
Hartley, M., F. Voller, T. Murray, and H. Roberts. 2012. "Qualitative veterinary risk assessment of the role of wild deer in the likelihood of incursion and the impact on effective disease control of selected exotic notifiable diseases in England." *European Journal of Wildlife Research* 59 (2):257-270. doi: 10.1007/s10344-012-0674-7.
Harvala, H., and P. Simmonds. 2009. "Human parechoviruses: biology, epidemiology and clinical significance." *J Clin Virol* 45 (1):1-9. doi: 10.1016/j.jcv.2009.03.009.
Hassell, J. M., M. Begon, M. J. Ward, and E. M. Fevre. 2017. "Urbanization and Disease Emergence: Dynamics at the Wildlife-Livestock-Human Interface." *Trends Ecol Evol* 32 (1):55-67. doi: 10.1016/j.tree.2016.09.012.
Hewson, Raymond. 2009. "Browsing by Mountain hares *Lepus timidus* on trees and shrubs in north-east Scotland." *Journal of Zoology* 182 (2):168-171. doi: 10.1111/j.1469-7998.1977.tb04152.x.
Hill, D.A., and P.A. Robertson. 1988. *The pheasant: ecology, management and conservation*. Oxford: Blackwell Scientific Publications.
Hilton, G.M., and R.J. Cuthbert. 2010. "The catastrophic impact of invasive mammalian predators on birds of the UK Overseas Territories: a review and synthesis." *Ibis* 152:443-458.
Hirst, C. 2021. Deer in a changing climate – how do wild deer affect carbon sequestration in Scottish woodlands? Edinburgh: University of Edinburgh.

Hof, A. R., A. M. Allen, and P. W. Bright. 2019. "Investigating the Role of the Eurasian Badger (*Meles meles*) in the Nationwide Distribution of the Western European Hedgehog (*Erinaceus europaeus*) in England." *Animals (Basel)* 9 (10). doi: 10.3390/ani9100759.

Holding, M., S. D. Dowall, J. M. Medlock, D. P. Carter, S. T. Pullan, J. Lewis, R. Vipond, M. S. Rocchi, M. Baylis, and R. Hewson. 2020. "Tick-Borne Encephalitis Virus, United Kingdom." *Emerg Infect Dis* 26 (1):90-96. doi: 10.3201/eid2601.191085.

Horemans, M., J. Van Bets, T. Joly Maes, P. Maes, and B. Vanmechelen. 2023. "Discovery and genome characterization of six new orthoparamyxoviruses in small Belgian mammals." *Virus Evol* 9 (2):vead065. doi: 10.1093/ve/vead065.

Hornok, S., K. Trauttwein, N. Takacs, A. Hodzic, G. G. Duscher, and J. Kontschan. 2017. "Molecular analysis of Ixodes rugicollis, Candidatus Neoehrlichia sp. (FU98) and a novel Babesia genotype from a European badger (*Meles meles*)." *Ticks Tick Borne Dis* 8 (1):41-44. doi: 10.1016/j.ttbdis.2016.09.014.

Hothorn, T., J. Müller, L. Held, L. Möst, and A. Mysterud. 2015. "Temporal patterns of deer–vehicle collisions consistent with deer activity pattern and density increase but not general accident risk." *Accident, Analysis and Prevention* 81:143-152. doi: https://doi.org/10.1016/j.aap.2015.04.037.

Hounsome, T.I.M., and Richard Delahay. 2005. "Birds in the diet of the Eurasian badger *Meles meles*: a review and meta-analysis." *Mammal Review* 35 (2):199-209. doi: 10.1111/j.1365-2907.2004.00061.x.

Howald, G., C. J. Donlan, J. P. Galvan, J. C. Russell, J. Parkes, A. Samaniego, Y. Wang, D. Veitch, P. Genovesi, M. Pascal, A. Saunders, and B. Tershy. 2007. "Invasive rodent eradication on islands." *Conserv Biol* 21 (5):1258-68. doi: 10.1111/j.1523-1739.2007.00755.x.

Hoyt, J.R., A.M. Kilpatrick, and K.E. Langwig. 2021. "Ecology and impacts of white-nose syndrome on bats." *Nature Reviews* 19:196-210.

Hughes, K., and A. M. Borman. 2018. "Adiaspiromycosis in a wild European rabbit, and a review of the literature." *J Vet Diagn Invest* 30 (4):614-618. doi: 10.1177/1040638718772631.

Hull, K.D., S. Jeckel, J.M. Williams, S.A. Ciavaglia, L.M.I. Webster, E. Fitzgerald, Y.M. Chang, and H.M. Martineau. 2022. "Fox (*Vulpes vulpes*) involvement identified in a series of cat carcass mutilations." *Vet Pathol* 59 (2):299-309. doi: 10.1177/03009858211052661.

Jackson, D.B., and R.E. Green. 2000. "The importance of the introduced hedgehog (*Erinaceus europaeus*) as a predator of the eggs of waders (Charadrii) on machair in South Uist, Scotland." *Biological Conservation* 93:333–48. doi: https://doi.org/10.1016/S0006-3207(99)00135-4.

Jackson, Digger B., Robert J. Fuller, and Steve T. Campbell. 2004. "Long-term population changes among breeding shorebirds in the Outer Hebrides, Scotland, in relation to introduced hedgehogs (*Erinaceus europaeus*)." *Biological Conservation* 117 (2):151-166. doi: 10.1016/s0006-3207(03)00289-1.

Jackson, J.E. 1994. "The edible or fat dormouse (*Glis glis*) in Britain." *Quarterly Journal of Forestry* 88:119-125.

Jameson, L., C. Logue, B. Atkinson, N. Baker, S. Galbraith, M. Carroll, T. Brooks, and R. Hewson. 2013. "The continued emergence of hantaviruses: isolation of a Seoul virus implicated in human disease, United Kingdom, October 2012." *Euro Surveillance* 18:4-7.

Jefferies, D.J. 2003. The water vole and mink survey of Britain 1996–1998 with a history of the long term changes in the status of both species and their causes. Ledbury, UK: The Vincent Wildlife Trust.

Jenkins, H. E., R. Woodroffe, and C. A. Donnelly. 2008. "The effects of annual widespread badger culls on cattle tuberculosis following the cessation of culling." *Int J Infect Dis* 12 (5):457-65. doi: 10.1016/j.ijid.2008.04.001.

Johnson, N., M. Golding, and L. P. Phipps. 2021. "Detection of Tick-Borne Pathogens in Red Deer (*Cervus elaphus*), United Kingdom." *Pathogens* 10 (6). doi: 10.3390/pathogens10060640.

Jones, H.P., B.R. Tershy, E.S. Zavaleta, D.A. Croll, B.S. Keitt, M.E. Finkelstein, and G.R. Howald. 2008. "Severity of the effects of invasive rats on seabirds:

a global review." *Conservation Biology* 22:16–26.

Jones, L.D., M. Gaunt, R.S. Hails, K. Laurenson, P.J. Hudson, H. Reid, P. Henbest, and E.A. Gould. 1997. "Transmission of louping ill virus between infected and uninfected ticks co-feeding on mountain hares." *Med Vet Entomol* 11 (2):172-6. doi: 10.1111/j.1365-2915.1997.tb00309.x.

Judge, J., G.J. Wilson, R. Macarthur, R.J. Delahay, and R.A. McDonald. 2014. "Density and abundance of badger social groups in England and Wales in 2011-2013." *Scientific Reports* 4:3809.

Judge, Johanna, Ross S. Davidson, Glenn Marion, Piran C. L. White, and Michael R. Hutchings. 2007. "Persistence of *Mycobacterium avium* subspecies paratuberculosis in rabbits: the interplay between horizontal and vertical transmission." *Journal of Applied Ecology* 44 (2):302-311. doi: 10.1111/j.1365-2664.2007.01282.x.

Jurczyk, K., K. Bunkowska-Gawlik, A. Perec-Matysiak, and J. Hildebrand. 2022. "Rickettsiae among mustelids - new data from south-west Poland." *Ann Parasitol* 68 (2):617-622. doi: 10.17420/ap6803.468.

Justus, William, Simon Valle, Owain Barton, Amy Gresham, and Graeme Shannon. 2024. "A review of bovine tuberculosis transmission risk in European wildlife communities." *Mammal Review* 54 (3):325-340. doi: 10.1111/mam.12347.

Kallio, E.R., M. Begon, H. Henttonen, E. Koskela, T. Mappes, A. Vaheri, and O. Vapalahti. 2009. "Cyclic hantavirus epidemics in humans predicted by rodent host dynamics." *Epidemics* 1 (2):101-7. doi: 10.1016/j.epidem.2009.03.002.

Keesing, F., and R.S. Ostfield. 2024. "Emerging patterns in rodent-borne zoonotic diseases." *Science* 385 (6715):1305-1310. doi: https://doi.org/10.1126/science.adq7993.

Kelly, D.J., E. Mullen, and M. Good. 2021. "Bovine Tuberculosis: The Emergence of a New Wildlife Maintenance Host in Ireland." *Front Vet Sci* 8:632525. doi: 10.3389/fvets.2021.632525.

Kelly, J., D. Tosh, K. Dale, and A. Jackson. 2013. The economic cost of invasive and non-native species in Ireland and Northern Ireland. A report prepared for the Northern Ireland Environment Agency and National Parks and wildlife Service as part of Invasive Species Ireland

Keymer, I.F., E.A. Gibson, and D.J. Reynolds. 1991. "Zoonoses and other findings in hedgehogs (*Erinaceus europaeus*): a survey of mortality and review of the literature." *Vet Rec* 128 (11):245-9. doi: 10.1136/vr.128.11.245.

Kilonzo, C., X. Li, E.J. Vivas, M.T. Jay-Russell, K.L. Fernandez, and E.R. Atwill. 2013. "Fecal shedding of zoonotic food-borne pathogens by wild rodents in a major agricultural region of the central California coast." *Appl Environ Microbiol* 79 (20):6337-44. doi: 10.1128/AEM.01503-13.

Kilshaw, Kerry, Robert A. Montgomery, Ruairidh D. Campbell, David A. Hetherington, Paul J. Johnson, Andrew C. Kitchener, David W. Macdonald, and Joshua J. Millspaugh. 2015. "Mapping the spatial configuration of hybridization risk for an endangered population of the European wildcat (*Felis silvestris silvestris*) in Scotland." *Mammal Research* 61 (1):1-11. doi: 10.1007/s13364-015-0253-x.

Kipar, A., S. J. Burthe, U. Hetzel, M. A. Rokia, S. Telfer, X. Lambin, R. J. Birtles, M. Begon, and M. Bennett. 2014. "*Mycobacterium microti tuberculosis* in its maintenance host, the field vole (*Microtus agrestis*): characterization of the disease and possible routes of transmission." *Vet Pathol* 51 (5):903-14. doi: 10.1177/0300985813513040.

Kirby, K.J. 2001. "The impact of deer on the ground flora of British broadleaved woodland." *Forestry* 74 (3):219-229. doi: 10.1093/forestry/74.3.219.

Kjemtrup, A.M., and P.A. Conrad. 2000. "Human babesiosis: an emerging tick-borne disease." *Int J Parasitol* 30 (12-13):1323-37. doi: 10.1016/s0020-7519(00)00137-5.

Kloskowski, Janusz. 2010. "Human–wildlife conflicts at pond fisheries in eastern Poland: perceptions and management of wildlife damage." *European Journal of Wildlife Research* 57 (2):295-304. doi: 10.1007/s10344-010-0426-5.

Kruckemeier, S., M. Ramon, E. Vidal, L. Martino, J. Burgaya, M. P. Ribas, A. Dias-Alves, L. Lobato-Bailon, B. Perez de Val, O. Cabezon, and J. Espunyes. 2024. "Adiaspiromycoses in Wild Rodents from the Pyrenees, Northeastern Spain." *J Wildl Dis* 60 (2):526-530. doi: 10.7589/JWD-D-23-00100.

Lambert, Sébastien, Jean Hars, Edouard Réveillaud, Jean-Louis Moyen, Hélène Gares, Thomas Rambaud, Eric Gueneau, Eva Faure, Maria-Laura Boschiroli, and Céline Richomme. 2016. "Host status of wild roe deer in bovine tuberculosis endemic areas." *European Journal of Wildlife Research* 63 (1). doi: 10.1007/s10344-016-1071-4.

Lambin, X., J.C. Horrill, and R. Raynor. 2019. "Achieving large scale, long-term invasive American mink control in northern Scotland despite short term funding." In *Island invasives: scaling up to meet the challenge. Occasional Paper SSC no. 62*, edited by C.R. Veitch, M.N. Clout, A.R. Martin, J.C. Russell and C.J. West, 651–657. Gland, Switzerland: IUCN.

Langbein, J. 1997. The ranging behaviour, habitat-use and impact of deer in oak woods and heather moors of Exmoor and the Quantock Hills. Fordingbridge, UK: British Deer Society.

Langbein, J. 2007. National deer-vehicle collisions project: England 2003-2005. Highways Agency Research Report 07/1.

Langbein, J. 2008. DVCs in peri-urban areas: a risky life for deer. Presentation at the British Deer Society conference, London.

Langbein, J. 2018. Deer-vehicle collisions in Scotland: National data overview, regional hotspots, trends, mitigation. Presentation to Lowland Deer Network, June 2018.

Langbein, J., and S.M. Rutter. 2003. Quantifying the damage wild deer cause to agricultural crops and pastures. Proceedings of the Future for Deer Conference, March 2003. Peterborough, UK: English Nature Research Report no. 548.

Langton, S.D., D.P. Cowan, and A.N. Meyer. 2001. "The occurrence of commensal rodents in dwellings as revealed by the 1996 English House Condition Survey." *Journal of Applied Ecology* 38 (4):699-709. doi: 10.1046/j.1365-2664.2001.00631.x.

Lawson, B., L. H. V. Franklinos, J. Rodriguez-Ramos Fernandez, C. Wend-Hansen, S. Nair, S. K. Macgregor, S. K. John, R. Pizzi, A. Nunez, P. M. Ashton, A. A. Cunningham, and M. de Pinna E. 2018. "*Salmonella Enteritidis* ST183: emerging and endemic biotypes affecting western European hedgehogs (*Erinaceus europaeus*) and people in Great Britain." *Sci Rep* 8 (1):2449. doi: 10.1038/s41598-017-18667-2.

Lawton, John H., and Gordon L. Woodroffe. 1991. "Habitat and the Distribution of Water Voles: Why are there Gaps in a Species' Range?" *The Journal of Animal Ecology* 60 (1). doi: 10.2307/5446.

Le Gall-Recule, G., A. Lavazza, S. Marchandeau, S. Bertagnoli, F. Zwingelstein, P. Cavadini, N. Martinelli, G. Lombardi, J. L. Guerin, E. Lemaitre, A. Decors, S. Boucher, B. Le Normand, and L. Capucci. 2013. "Emergence of a new lagovirus related to Rabbit Haemorrhagic Disease Virus." *Vet Res* 44 (1):81. doi: 10.1186/1297-9716-44-81.

Learmount, J., V. Boughtflower, P.C. Allanson, K.M. Hartley, A.B. Gutierrez, N.A. Stephens, G. Marucci, and G.C. Smith. 2015. "First report of Trichinella pseudospiralis in a red fox in mainland Britain." *Vet Parasitol* 208 (3-4):259-62. doi: 10.1016/j.vetpar.2015.01.011.

Learmount, J., I. Zimmer, C. Conyers, V. Boughtflower, C. Morgan, and G. Smith. 2012. " A diagnostic study of *Echinococcus multilocularis* in red foxes (*Vulpes vulpes*) from Great Britain." *Veterinary Parasitology* 190:447-453.

Lever, C. 1994. *Naturalized animals: the ecology of successfully introduced species*. London: Poyser.

Lincolnshire Live. 2024. "Badgers cause £100K of damage in Mablethorpe leading to emergency road closure." https://www.lincolnshirelive.co.uk/news/local-news/badgers-cause-100k-damage-mablethorpe-9689319. Date accessed: 14th November 2024.

Lindhorst, Z.T.L., S. Brandstetter, M.S. Unterkofler, B. Eigner, J. Spergser, M. Colyn, P. Steinbach, D. Cirovic, N. Sprem, T. Dumic, V. Veneziano, F. Muller, J. Harl, G. Deak, A.M. Ionica, M. Heddergott, and H.P. Fuehrer. 2024. "Molecular analysis of vector-borne pathogens in Eurasian badgers (Meles meles) from continental Europe." *Parasit Vectors* 17 (1):451. doi: 10.1186/s13071-024-06515-y.

Lock, J. 2006. "Eradication of brown rats *Rattus norvegicus* and black rats *Rattus rattus* to restore breeding seabird populations on Lundy Island, Devon, England." *Conservation Evidence* 3:111-113.

Lodé, Thierry. 1993. "Diet composition and habitat use of sympatric polecat and American mink in western France." *Acta Theriologica* 38:161-166. doi: 10.4098/AT.arch.93-14.

Long, J.L. 2003. *Introduced mammals of the world: Their history, distribution and influence*. Victoria, Australia: CSIRO Publishing.

Loss, S.R., T. Will, and P.P. Marra. 2013. "The impact of free-ranging domestic cats on wildlife of the United States." *Nat Commun* 4:1396. doi: 10.1038/ncomms2380.

Lowe, V.P.W., and A.S. Gardiner. 2009. "Hybridization between Red deer (*Cervus elaphus*) and Sika deer (*Cervus nippon*) with particular reference to stocks in N.W. England." *Journal of Zoology* 177 (4):553-566. doi: 10.1111/j.1469-7998.1975.tb02259.x.

Ludwig, Sonja C., Staffan Roos, Damian Bubb, and Dave Baines. 2017. "Long-term trends in abundance and breeding success of red grouse and hen harriers in relation to changing management of a Scottish grouse moor." *Wildlife Biology* 2017 (SP1). doi: 10.2981/wlb.00246.

Lurz, P.W.W., N. Geddes, A.J. Lloyd, M.D.F. Shirley, S.P. Rushton, and B. Burlton. 2003. "Planning a red squirrel conservation area: using a spatially explicit population dynamics model to predict the impact of felling and forest design plans." *Forestry* 76:95–108.

Maag, N., J.W. Mallord, M.D. Burgess, S. Lupold, A. Cristinacce, R. Arlettaz, S. Carlotti, T. M. Davis, A. Grendelmeier, C.J. Orsman, M. Riess, P. Stelbrink, and G. Pasinelli. 2022. "Accounting for predator species identity reveals variable relationships between nest predation rate and habitat in a temperate forest songbird." *Ecol Evol* 12 (10):e7411. doi: 10.1002/ece3.9411.

Macdonald, D.W., and P. Doncaster. 1985. Foxes in your neighbourhood? Horsham, W Sussex: RSPCA.

Macdonald, D.W., F. Tattersall, P. Johnson, C. Carbone, J. Reynolds, J. Langbein, S. Rushton, and M. Shirley. 2000. Managing British mammals: case studies from the hunting debate. Oxford: Wildlife Conservation Research Unit.

Macdonald, D.W., F.H. Tattersall, E.D. Brown, and D. Balharry. 1995. "Reintroducing the European beaver to Britain: Nostalgic meddling or restoring biodiversity?" *Mammal Review* 25 (4):161-200.

Macmillan, D.C., and S. Phillip. 2008. "Consumptive and non-consumptive values of wild mammals in Britain." *Mammal Review* 38 (2-3):189-204. doi: 10.1111/j.1365-2907.2008.00124.x.

MacPherson, Jenny L., and Paul W. Bright. 2011. "Metapopulation dynamics and a landscape approach to conservation of lowland water voles (*Arvicola amphibius*)." *Landscape Ecology* 26 (10):1395-1404. doi: 10.1007/s10980-011-9669-0.

Malatesta, D., V.R. Simpson, L. Fontanesi, R. Fusillo, M. Marcelli, L. Bongiovanni, M. Romanucci, C. Palmieri, and L. Della Salda. 2014. "First description of adiaspiromycosis in an Eurasian otter (*Lutra lutra*) in Italy." *Vet Ital* 50 (3):199-202. doi: 10.12834/VetIt.40.1916.8.

Malpas, Lucy R., Rosalind J. Kennerley, Graham J. M. Hirons, Rob D. Sheldon, Malcolm Ausden, Joanne C. Gilbert, and Jennifer Smart. 2013. "The use of predator-exclusion fencing as a management tool improves the breeding success of waders on lowland wet grassland." *Journal for Nature Conservation* 21 (1):37-47. doi: 10.1016/j.jnc.2012.09.002.

Mammal Society. 2015. "Re-establishment of Wild Boar in Britain." https://mammal.org.uk/position-statements/re-establishment-of-wild-boar-in-britain. Date Accessed: 30th April 2024.

Mammal Society. 2023. "Roe deer." https://mammal.org.uk/british-mammals. Date accessed: 30th April 2024.

Mammal Society. 2024a. "Greater white-toothed shrew." https://mammal.org.uk/british-mammals/greater-white-toothed-shrew. Date accessed: 27th November 2024.

Mammal Society. 2024b. "Mountain hare." https://mammal.org.uk/british-mammals/mountain-hare. Date accessed: 27th November 2024.

Mann, J.C.E., and R.J. Putman. 1989. "Diet of British sika deer (*Cervus nippon* Temminck) in contrasting environments." *Acta Theriologica* 34:97-110.

Marnell, F., and P.  Presetnik. 2010. Protection of overground roosts for bats (particularly roosts in buildings of cultural heritage importance). Bonn: Eurobats.

Mathews, F., L.M. Kubasiewicz, J. Gurnell, C.A. Harrower, R.A. McDonald, and R.F. Shore. 2018. A review of the population and conservation status of British mammals: technical summary. A report by the Mammal Society under contract to Natural England, Natural Resources Wales and Scottish Natural Heritage. Peterborough: Natural England.

Matuschka, F.R., H. Eiffert, A. Ohlenbusch, and A. Spielman. 1994. "Amplifying role of edible dormice in Lyme disease transmission in central Europe." *J Infect Dis* 170 (1):122-7. doi: 10.1093/infdis/170.1.122.

McBride, A. 2002. "Restoration action planning." In *Guidance for the restoration of montane scrub*, edited by D. Gilbert, 7-17. Munlochy, UK: Highland Birchwoods.

McDevitt, A. D., W. I. Montgomery, D. G. Tosh, J. Lusby, N. Reid, T. A. White, C. D. McDevitt, J. O'Halloran, J. B. Searle, and J. M. Yearsley. 2014. "Invading and expanding: range dynamics and ecological

consequences of the greater white-toothed shrew (Crocidura russula) invasion in Ireland." *PLoS One* 9 (6):e100403. doi: 10.1371/journal.pone.0100403.
McDonald, Robbie A., Kevin O'Hara, and Daniel J. Morrish. 2007. "Decline of invasive alien mink (Mustela vison) is concurrent with recovery of native otters (Lutra lutra)." *Diversity and Distributions* 13 (1):92-98. doi: 10.1111/j.1366-9516.2006.00303.x.
McKay, J. 1995. *Complete guide to ferrets*. Shrewsbury: Swan Hill Press.
McKillen, J., K. Hogg, P. Lagan, C. Ball, S. Doherty, N. Reid, L. Collins, and J. T. A. Dick. 2017. "Detection of a novel gammaherpesvirus (genus Rhadinovirus) in wild muntjac deer in Northern Ireland." *Arch Virol* 162 (6):1737-1740. doi: 10.1007/s00705-017-3254-z.
McLauchlan, J.D., and W.M. Henderson. 1947. "The occurrence of foot-and-mouth disease in the hedgehog under natural conditions." *J Hyg (Lond)* 45 (4):474-9. doi: 10.1017/s0022172400014194.
McOrist, S., and A.C. Kitchener. 1994. "Current threats to the European Wildcat, *Felis silvestris*, in Scotland." *Ambio* 23:243-245.
Mead, C.J. 2008. "Ringed birds killed by cats." *Mammal Review* 12 (4):183-186. doi: 10.1111/j.1365-2907.1982.tb00014.x.
Melero, Y., S. Palazon, and X. Lambin. 2014. "Invasive crayfish reduce food limitation of alien American mink and increase their resilience to control." *Oecologia* 174 (2):427-34. doi: 10.1007/s00442-013-2774-9.
Meredith, A., J. Del Pozo, S. Smith, E. Milne, K. Stevenson, and J. McLuckie. 2014. "Leprosy in red squirrels in Scotland." *Vet Rec* 175 (11):285-6. doi: 10.1136/vr.g5680.
Meyer, A.N. 1994. "Rodent control in practice: food stores." In *Rodent pests and their control*, edited by A.P. Buckle and R.H. Smith, 273-290. Wallingford: CAB International.
Michelet, L., K. De Cruz, S. Henault, J. Tambosco, C. Richomme, E. Reveillaud, H. Gares, J. L. Moyen, and M. L. Boschiroli. 2018. "*Mycobacterium bovis* Infection of Red Fox, France." *Emerg Infect Dis* 24 (6):1150-1153. doi: 10.3201/eid2406.180094.
Millins, C., A. Magierecka, L. Gilbert, A. Edoff, A. Brereton, E. Kilbride, M. Denwood, R. Birtles, and R. Biek. 2015. "An Invasive Mammal (the Gray Squirrel, Sciurus carolinensis) Commonly Hosts Diverse and Atypical Genotypes of the Zoonotic Pathogen Borrelia burgdorferi Sensu Lato." *Appl Environ Microbiol* 81 (13):4236-45. doi: 10.1128/AEM.00109-15.
Milne, G., A. Allen, J. Graham, R. Kirke, C. McCormick, E. Presho, R. Skuce, and A.W. Byrne. 2020. "Mycobacterium bovis Population Structure in Cattle and Local Badgers: Co-Localisation and Variation by Farm Type." *Pathogens* 9 (7). doi: 10.3390/pathogens9070592.
Mitchell-Jones, A.J. 2020. Badger impacts on biodiversity and agriculture in Scotland: a literature review. NatureScot Research Report No. 1205.
Moller, H. 2008. "Foods and foraging behaviour of Red (Sciurus vulgaris) and Grey (Sciurus carolinensis) squirrels." *Mammal Review* 13 (2-4):81-98. doi: 10.1111/j.1365-2907.1983.tb00270.x.
Moore, N.P., P.A. Robertson, and J.N. Aegerter. 2000. Feasibility study into the options for management of mink in the Western Isles. Report to Scottish Natural Heritage.
Moore, Niall, Anne Whiterow, Paul Kelly, David Garthwaite, Julie Bishop, Steve Langton, and Chris Cheeseman. 2001. "Survey of badger Meles meles damage to agriculture in England and Wales." *Journal of Applied Ecology* 36 (6):974-988. doi: 10.1046/j.1365-2664.1999.00454.x.
Moorhouse, T.P., D.W. Macdonald, R. Strachan, and X. Lambin. 2015. "What Does Conservation Research Do, When Should it Stop, and What Do we Do Then? Questions Answered with Water Voles." In *Wildlife Conservation on Farmland Volume 1. Managing for Nature on Lowland Farms*, edited by D.W. Macdonald and R.E. Feber, 269–290. Oxford: : Oxford University Press.
Morgan, E.R., D. Modry, C. Paredes-Esquivel, P. Foronda, and D. Traversa. 2021. "Angiostrongylosis in Animals and Humans in Europe." *Pathogens* 10 (10). doi: 10.3390/pathogens10101236.
Morris, P. 2008. "Edible dormouse *Glis glis*." In *The handbook of British mammals, 4th edn*, edited by Harris S. and Yalden D.W, 82–85. Southampton: The Mammal Society, .
Morris, P., and Mary P. English. 1969. "*Trichophyton mentagrophytes* var. *erinacei* in British hedgehogs." *Medical Mycology* 7 (2):122-128. doi: 10.1080/00362177085190221.
Morris, P.A. 1997. "A review of the fat dormouse (*Glis glis*) in Britain." *Natura Croatica* 6:163–176.

Murphy, E.G., N.J. Williams, D. Jennings, J. Chantrey, R. Verin, S. Grierson, L.M. McElhinney, and M. Bennett. 2019. "First detection of Hepatitis E virus (Orthohepevirus C) in wild brown rats (*Rattus norvegicus*) from Great Britain." *Zoonoses Public Health* 66 (6):686-694. doi: 10.1111/zph.12581.

Murphy, R.G., R.H. Williams, J.M. Hughes, G. Hide, N.J. Ford, and D.J. Oldbury. 2008. "The urban house mouse (*Mus domesticus*) as a reservoir of infection for the human parasite *Toxoplasma gondii*: an unrecognised public health issue?" *International Journal of Environmental Health Research* 18 (3):177 – 185.

Murray, Christopher, Jeroen Minderman, James Allison, and John Calladine. 2016. "Vegetation structure influences foraging decisions in a declining grassland bird: the importance of fine-scale habitat and grazing regime." *Bird Study* 63 (2):223-232. doi: 10.1080/00063657.2016.1180342.

Nally, J.E., Z. Arent, D. O' Bayles, R.L. Horsnby, C. Gilmore, S. Regan, A.D. McDevitt, J. Yearsley, S. Fanning, and B.J. McMahon. 2016. "Emerging infectious disease implications of invasive mammalian species: the greater white-toothed shrew (*Crocidura russula*) is associated with a novel serovar of pathogenic Leptospira in Ireland." *PLOS Neglected Tropical Diseases* 10:e0005174.

National Forest Inventory. 2020. https://www.data.gov.uk/dataset/6e3126bd-fb2c-4cac-b2c4-d521f006b87a/national-forest-inventory-woodland-england-20202021. Date accessed: 10th December 2024.

Natural England. 2011. Natural England Species Information Note SIN006 Otter: European protected species.

Natural England. 2022a. "Bats: advice for making planning decisions.". https://www.gov.uk/guidance/bats-advice-for-making-planning-decisions.

Natural England. 2022b. "Hazel dormice: advice for making planning decisions. ." https://www.gov.uk/guidance/hazel-dormice-advice-for-making-planning-decisions.

Natural England. 2023. "Summary of wildlife licences issued by Natural England in 2023. ." https://naturalengland.blog.gov.uk/2024/03/06/natural-england-wildlife-licensing-statistics-for-2023/.

NatureScot. 2015. Tayside Beaver Study Group. Final report.

NatureScot. 2022a. "Beaver Management Report for 2022.". www.nature.scot/doc/beaver-management-report-2022.

NatureScot. 2022b. Updated non-native species risk assessment of feral pigs in Scotland. NatureScot Research Report 1288.

Needham, Robert J., Martin Gaywood, Angus Tree, Nick Sotherton, Dylan Roberts, Colin W. Bean, and Paul S. Kemp. 2021. "The response of a brown trout (*Salmo trutta*) population to reintroduced Eurasian beaver (*Castor fiber*) habitat modification." *Canadian Journal of Fisheries and Aquatic Sciences* 78 (11):1650-1660. doi: 10.1139/cjfas-2021-0023.

Nichol, S., S.J. Ball, and K.R. Snow. 1981. "Prevalence of intestinal parasites in feral cats in some urban areas of England." *Vet Parasitol* 9 (2):107-10. doi: 10.1016/0304-4017(81)90028-5.

Nichols, Christopher P., Julian A. Drewe, Robin Gill, Nigel Goode, and Neville Gregory. 2016. "A novel causal mechanism for grey squirrel bark stripping: The Calcium Hypothesis." *Forest Ecology and Management* 367:12-20. doi: 10.1016/j.foreco.2016.02.021.

Niklasson, B., B. Hornfeldt, and B. Lundman. 1998. "Could myocarditis, insulin-dependent diabetes mellitus, and Guillain-Barre syndrome be caused by one or more infectious agents carried by rodents?" *Emerg Infect Dis* 4 (2):187-93. doi: 10.3201/eid0402.980206.

Nota, Kevin, Stephen Downing, and Arati Iyengar. 2019. "Metabarcoding-based dietary analysis of hen harrier (*Circus cyaneus*) in Great Britain using buccal swabs from chicks." *Conservation Genetics* 20 (6):1389-1404. doi: 10.1007/s10592-019-01215-y.

O'Hanlon, Nina J., and Mark S. Lambert. 2016. "Investigating brown rat *Rattus norvegicus* egg predation using experimental nests and camera traps." *European Journal of Wildlife Research* 63 (1). doi: 10.1007/s10344-016-1063-4.

Occhibove, F., N.J. McKeown, C. Risley, and J.E. Ironside. 2022. "Eco-epidemiological screening of multi-host wild rodent communities in the UK reveals pathogen strains of zoonotic interest." *Int J Parasitol Parasites Wildl* 17:278-287. doi: 10.1016/j.ijppaw.2022.02.010.

Ogurtsov, S.S., A.S. Zheltukhin, and I.P. Kotlov. 2018. "Daily activity patterns of large and medium-sized mammals based on camera traps data in the Central Forest Nature Reserve, Valdai Upland, Russia." *Nature Conservation Research,* 3:68-88.

Olival, K.J., P.R. Hosseini, C. Zambrana-Torrelio, N. Ross, T.L. Bogich, and P. Daszak. 2017. "Host and viral traits predict zoonotic spillover from mammals." *Nature* 546 (7660):646-650. doi: 10.1038/nature22975.

Oliver, M., J.J. Luque-Larena, and X. Lambin. 2009. "Do rabbits eat voles? Apparent competition, habitat heterogeneity and large-scale coexistence under mink predation." *Ecol Lett* 12 (11):1201-9. doi: 10.1111/j.1461-0248.2009.01375.x.

Olsthoorn, F., H. Sprong, M. Fonville, M. Rocchi, J. Medlock, L. Gilbert, and J. Ghazoul. 2021. "Occurrence of tick-borne pathogens in questing Ixodes ricinus ticks from Wester Ross, Northwest Scotland." *Parasit Vectors* 14 (1):430. doi: 10.1186/s13071-021-04946-5.

Packer, James J., and Johnny D.S. Birks. 2002. "An assessment of British farmers' and gamekeepers' experiences, attitudes and practices in relation to the European Polecat Mustela putorius." *Mammal Review* 29 (2):75-92. doi: 10.1046/j.1365-2907.1999.00039.x.

Parrott, Dave, Thomas R. Etherington, and Julie Dendy. 2014. "A geographically extensive survey of hedgehogs (Erinaceus europaeus) in England." *European Journal of Wildlife Research* 60 (2):399-403. doi: 10.1007/s10344-014-0795-2.

Pearce, S.J., K. Turner, T.A. Cogan, and A.P. Foster. 2023. "A survey of enteric disease agents in UK deer populations." *Res Vet Sci* 154:113-123. doi: 10.1016/j.rvsc.2022.12.007.

PECBMS. 2012. *Population trends of common European breeding birds 2012*. Prague: CSO/RSPB.

Pelz, H.J. 1989. "Ecological aspects of damage to sugar beet seeds by *Apodemus sylvaticus*." In *Mammals as pests*, edited by R.J. Putman, 34-48. London: Chapman & Hall.

Pepper, S., A. Barbour, and J. Glass. 2019. The management of wild deer in Scotland. Report of the Deer Working Group.

Petrovan, S.O., A.I. Ward, and P. Wheeler. 2011. "Detectability counts when assessing populations for biodiversity targets." *PLoS One* 6 (9):e24206. doi: 10.1371/journal.pone.0024206.

PHE. 2020. "Hepatitis E: Symptoms, transmission, treatment and prevention – GOV.UK. ." https://www.gov.uk/government/publications/hepatitis-e-symptoms-transmission-prevention-treatment/hepatitis-e-symptoms-transmission-treatment-and-prevention.

Pocock, M.J., J.B. Searle, W.B. Betts, and P.C.L. White. 2001. "Patterns of infection by Salmonella and Yersinia spp. in commensal house mouse (*Mus musculus domesticus*) populations." *J Appl Microbiol* 90 (5):755-60. doi: 10.1046/j.1365-2672.2001.01303.x.

Potts, G.R. 1986. *The partridge: pesticides, predation and conservation*. London: Collins.

Pounder, K.C., M. Begon, T. Sironen, H. Henttonen, P.C. Watts, L. Voutilainen, O. Vapalahti, B. Klempa, A.R. Fooks, and L.M. McElhinney. 2013. "Novel Hantavirus in Wildlife, United Kingdom." *Emerg Infect Dis* 19 (4):673-5. doi: 10.3201/eid1904.121057.

Powell, Sian, Nicola Dessi, Malcolm Bennett, Belinda Wang, Andrew Robertson, Elisabeth Waller, Graham Smith, and Richard Delahay. 2024. "Tuberculosis in found dead badgers at the edge of the expanding bovine tuberculosis epidemic." *Research Square Preprint*. doi: 10.21203/rs.3.rs-5193016/v1.

Puleston, R. L., C. M. Mallaghan, D. E. Modha, P. R. Hunter, J. S. Nguyen-Van-Tam, C. M. Regan, G. L. Nichols, and R. M. Chalmers. 2014. "The first recorded outbreak of cryptosporidiosis due to *Cryptosporidium cuniculus* (formerly rabbit genotype), following a water quality incident." *J Water Health* 12 (1):41-50. doi: 10.2166/wh.2013.097.

Pusterla, N., P.A. Pesavento, C.M. Leutenegger, J. Hay, L.J. Lowenstine, M.M. Durando, and K.G. Magdesian. 2002. "Disseminated pulmonary adiaspiromycosis caused by Emmonsia crescens in a horse." *Equine Vet J* 34 (7):749-52. doi: 10.2746/042516402776250342.

Putman, R., and J. Pemberton. 2022. "Sika in the British Isles: Population Ecology, Spread and Impacts of an Introduced Species." In *Sika Deer: Life History Plasticity and Management. Ecological Research Monographs*, edited by K. Kaji, H. Uno and H. Iijima. Singapore: Springer.

Putman, R.J. 2003. *The deer manager's companion*. Shrewsbury: Swan Hill Press.

Putman, R.J. 2008. "Ungulates: Orders Perissodactyla and Artiodactyla." In *Mammals of the British Isles (4th edn.)*, edited by S. Harris and D. Yalden, 556-653. Southampton: The Mammal Society.

Putman, R.J., S. Culpin, and S.J. Thirgood. 1993. "Dietary differences between male and female fallow deer in sympatry and in allopatry." *Journal of Zoology* 229 (2):267–275. doi: https://doi.org/10.1111/j.1469-7998.1993.tb02635.x.
Putman, R.J., and E.J. Hunt. 1994. "Patterns of hybridisation and introgression between red and sika deer in different populations of the North of Scotland and Argyl." *Deer* 9:104-110.
Putman, R.J., and P. Kjellander. 2002. "Deer damage to cereals: economic significance and predisposing factors." In *Conservation and conflict – Mammals and farming in Britain*, edited by F. Tattersall and W. Manley, 186-197. London: Linnean Society Occasional Publication 4.
Putman, R.J., and N.P. Moore. 1998. "Impact of deer in lowland Britain on agriculture, forestry and conservation habitats." *Mammal Review* 28 (4):141-164.
Puttock, A., H.A. Graham, A.M. Cunliffe, M. Elliott, and R.E. Brazier. 2017. "Eurasian beaver activity increases water storage, attenuates flow and mitigates diffuse pollution from intensively-managed grasslands." *Sci Total Environ* 576:430-443. doi: 10.1016/j.scitotenv.2016.10.122.
Quine, C.P., R. Shore, and R. Trout. 2004. Managing woodlands and their mammals: proceedings of a joint Mammal Society/Forestry Commission Symposium. Edinburgh: Forestry Commission.
Quy, R., and D. Poole. 2004. A review of the methods used within the European Union to control the European Mole, *Talpa europaea*. London: Defra.
Quy, R.J., D.P. Cowan, P.J. Haynes, A.P. Sturdee, R.M. Chalmers, A.T. Bodley-Tickell, and S.A. Bull. 1999. "The Norway rat as a reservoir host of *Cryptosporidium parvum*." *J Wildl Dis* 35 (4):660-70. doi: 10.7589/0090-3558-35.4.660.
Racey, P.A. 2008. "Bats: Order Chiroptera." In *Mammals of the British Isles (4th edn.)*, edited by S. Harris and D. Yalden, 292-391. Southampton: The Mammal Society.
Racey, P.A., A.M. Hutson, and P.H.C. Lina. 2013. "Bat Rabies, Public Health and European Bat Conservation." *Zoonoses and Public Health* 60:58-68. doi: doi: 10.1111/j.1863-2378.2012.01533.x.
Radford, A., G. Riddington, J. Anderson, and H. Gibson. 2004. The Economic impact of game and coarse angling in Scotland. Edinburgh, UK: Scottish Executive Environment and Rural Affairs Department.
Raymond, S., A.L.W. Schwartz, R.J. Thomas, E. Chadwick, and S.E. Perkins. 2021. "Temporal patterns of wildlife roadkill in the UK." *PLoS One* 16 (10):e0258083. doi: 10.1371/journal.pone.0258083.
Reid, H.W., R.M. Barlow, and I. Pow. 1978. "Isolation of louping-ill virus from red deer (*Cervus elaphus*)." *Vet Rec* 102 (21):463-4. doi: 10.1136/vr.102.21.463.
Reynolds, J., S. Dowell, M. Brockless, K. Blake, and N. Boatman. 1992. "Tracking partridge predation." *Game Conservancy Annual Review* 23:60-62.
RHS. 2024. "Rabbits." https://www.rhs.org.uk/biodiversity/rabbits. Date accessed: 2nd December 2024.
Richards, C. 1989. "The pest status of rodents in the United Kingdom." In *Mammals as pests*, edited by R.J. Putman. London: Chapman & Hall Ltd.
Riley, P.Y., and B.B. Chomel. 2005. "Hedgehog zoonoses." *Emerging Infectious Diseases* 11:1-5.
Roberts, B.R. 2019. "The hidden ecology of bumblebees: using classical and new methods to explore nest searching, floral resources and badger predation." PhD thesis, University of Exeter.
Robinson, Matthew T., Susan E. Shaw, and Eric R. Morgan. 2009. "*Anaplasma phagocytophilum* infection in a multi-species deer community in the New Forest, England." *European Journal of Wildlife Research* 55 (4):439-442. doi: 10.1007/s10344-009-0261-8.
Rocchi, M., M. Maley, M. Dagleish, and B. Boag. 2019. "Rabbit haemorrhagic disease virus type 2 in hares in Scotland." *Vet Rec* 185 (1):23. doi: 10.1136/vr.l4481.
Rooney, S., and T.J. Hayden. 2002. Forest mammals – management and control. Dublin: COFORD.
Roos, S., J. Smart, D.W. Gibbons, and J.D. Wilson. 2018. "A review of predation as a limiting factor for bird populations in mesopredator-rich landscapes: a case study of the UK." *Biol Rev Camb Philos Soc* 93 (4):1915-1937. doi: 10.1111/brv.12426.
Rosell, F., and R. Campbell-Palmer. 2022. *Beavers: Ecology, behaviour, conservation and management.* Oxford: Oxford University Press.
Ross, J., A.M. Tittensor, W.D. Hamilton, R.M. May, R.M. Anderson, J.H. Lawton, P. Holme, R.A. Brown, H.V. Thompson, and A. Gibbs. 1986. "The establishment and spread of myxomatosis and its effect on

rabbit populations [and discussion]." *Philosophical Transactions of the Royal Society of London. Series B, Biological Sciences* 314:599-606.
Rouco, C., J.A. Aguayo-Adan, S. Santoro, J. Abrantes, and M. Delibes-Mateos. 2019. "Worldwide rapid spread of the novel rabbit haemorrhagic disease virus (GI.2/RHDV2/b)." *Transbound Emerg Dis* 66 (4):1762-1764. doi: 10.1111/tbed.13189.
Rozycka, D., J.M. Lim, R.C. Trout, and S. Brooks. 2015. "Have feral boar significantly impacted hazel dormouse populations in Sussex, England?" *Folia Zoologica* 64 (4):337–341. doi: https://doi.org/10.25225/fozo.v64.i4.a8.2015.
RSPCA. 2022. "Road traffic accidents involving deer." https://www.rspca.org.uk/documents/1494939/7712578/Road+traffic+accidents+and+deer+-+2022+update.pdf. Date accessed: 30th April 2024.
Ruf, T., J. Fietz, W. Schlund, and C. Bieber. 2006. "High survival in poor years: life history tactics adapted to mast seeding in the edible dormouse." *Ecology* 87 (2):372-81. doi: 10.1890/05-0672.
Rushton, S.P. , P.W.W. Lurz, J. Gurnell, and R. Fuller. 2006. "Modelling the spatial dynamics of parapoxvirus disease in red and grey squirrels: a possible cause of the decline in the red squirrel in the UK?" *Journal of Applied Ecology* 37 (6):997-1012.
Rutter, S.M., and J. Langbein. 2005. Quantifying the damage wild deer cause to agricultural crops and pastures. Defra, Contract Report VC0327.
Sage, Rufus B., Kate Hollins, Catherine L. Gregory, Maureen I. A. Woodburn, and John P. Carroll. 2004. "Impact of roe deer *Capreolus capreolus* browsing on understorey vegetation in small farm woodlands." *Wildlife Biology* 10 (2):115-120. doi: 10.2981/wlb.2004.016.
Sage, Rufus B., Clare V. Turner, Maureen I.A. Woodburn, Andrew N. Hoodless, Roger A.H. Draycott, and Nicolas W. Sotherton. 2018. "Predation of released pheasants *Phasianus colchicus* on lowland farmland in the UK and the effect of predator control." *European Journal of Wildlife Research* 64 (2). doi: 10.1007/s10344-018-1174-1.
Sainsbury, A.W., R. Deaville, B. Lawson, W.A. Cooley, S.S. Farelly, M.J. Stack, P. Duff, C.J. McInnes, J. Gurnell, P.H. Russell, S.P. Rushton, D.U. Pfeiffer, P. Nettleton, and P.W. Lurz. 2008. "Poxviral disease in red squirrels *Sciurus vulgaris* in the UK: spatial and temporal trends of an emerging threat." *Ecohealth* 5 (3):305-16. doi: 10.1007/s10393-008-0191-z.
Sainsbury, K.A., R.F. Shore, H. Schofield, E. Croose, R.D. Campbell, and R.A. McDonald. 2019. "Recent history, current status, conservation and management of native mammalian carnivore species in Great Britain." *Mammal Review* 49 (2):171-88. doi: https://doi.org/10.1111/mam.12150.
Sala, Vittorio, and Eleonora De Faveri. 2016. "Epidemiology of Lyme Disease in Domestic and Wild Animals." *The Open Dermatology Journal* 10 (1):15-26. doi: 10.2174/1874372201610010015.
Salisbury, A.M., M. Begon, W. Dove, B. Niklasson, and J.P. Stewart. 2014. "Ljungan virus is endemic in rodents in the UK." *Arch Virol* 159 (3):547-51. doi: 10.1007/s00705-013-1731-6.
Santin-Duran, M., J.M. Alunda, E.P. Hoberg, and Fuente Concepcion de la. 2004. "Abomasal parasites in wild sympatric cervids, red deer, *Cervus elaphus* and fallow deer, *Dama dama*, from three localities across central and western Spain: relationship to host density and park management." *J Parasitol* 90 (6):1378-86. doi: 10.1645/GE-3376.
Santoro, M., C. Auriemma, M.G. Lucibelli, G. Borriello, N. D'Alessio, G. Sgroi, V. Veneziano, G. Galiero, and G. Fusco. 2019. "Molecular Detection of *Babesia* spp. (Apicomplexa: Piroplasma) in Free-Ranging Canids and Mustelids From Southern Italy." *Front Vet Sci* 6:269. doi: 10.3389/fvets.2019.00269.
Scaife, H.R., D. Cowan, J. Finney, S.F. Kinghorn-Perry, and B. Crook. 2006. "Wild rabbits (*Oryctolagus cuniculus*) as potential carriers of verocytotoxin-producing *Escherichia coli*." *Vet Rec* 159 (6):175-8. doi: 10.1136/vr.159.6.175.
Schatz, J., A. R. Fooks, L. McElhinney, D. Horton, J. Echevarria, S. Vazquez-Moron, E. A. Kooi, T. B. Rasmussen, T. Muller, and C. M. Freuling. 2013. "Bat rabies surveillance in Europe." *Zoonoses Public Health* 60 (1):22-34. doi: 10.1111/zph.12002.
Schilling, A.K., J. Del-Pozo, P.W.W. Lurz, K. Stevenson, C. Avanzi, C.M. Shuttleworth, S.T. Cole, and A.L. Meredith. 2019. "Leprosy in red squirrels in the UK." *Vet Rec* 184 (13):416. doi: 10.1136/vr.l1385.
Schmaljohn, C., and B. Hjelle. 1997. "Hantaviruses: a global disease problem." *Emerg Infect Dis* 3 (2):95-104. doi: 10.3201/eid0302.970202.

Schuchert, Pia, Craig M. Shuttleworth, Colin J. McInnes, David J. Everest, and Steven P. Rushton. 2014. "Landscape scale impacts of culling upon a European grey squirrel population: can trapping reduce population size and decrease the threat of squirrelpox virus infection for the native red squirrel?" *Biological Invasions* 16 (11):2381-2391. doi: 10.1007/s10530-014-0671-8.

Scott, Dawn M., Rowenna Baker, Alexandra Tomlinson, Maureen J. Berg, Naomi Charman, and Bryony A. Tolhurst. 2020. "Spatial distribution of sarcoptic mange (*Sarcoptes scabiei*) in urban foxes (*Vulpes vulpes*) in Great Britain as determined by citizen science." *Urban Ecosystems* 23 (5):1127-1140. doi: 10.1007/s11252-020-00985-5.

Selås, Vidar. 2006. "Does food competition from red fox (*Vulpes vulpes*) influence the breeding density of goshawk (*Accipiter gentilis*)? Evidence from a natural experiment." *Journal of Zoology* 246 (3):325-335. doi: 10.1111/j.1469-7998.1998.tb00162.x.

Senn, H.V., and J.M. Pemberton. 2008. "Variable extent of hybridization between invasive sika (*Cervus nippon*) and native red deer (*C. elaphus*) in a small geographical area." *Molecular Ecology* 18 (5):862-876.

Shaughnessy, L.J., L.A. Smith, J. Evans, D. Anderson, G. Caldow, G. Marion, J.C. Low, and M.R. Hutchings. 2013. "High prevalence of paratuberculosis in rabbits is associated with difficulties in controlling the disease in cattle." *Vet J* 198 (1):267-70. doi: 10.1016/j.tvjl.2013.08.030.

Shaw, R.F., R.J. Pakeman, M.R. Young, and G.R. Iason. 2013. "Microsite affects willow sapling recovery from bank vole (*Myodes glareolus*) herbivory, but does not affect grazing risk." *Ann Bot* 112 (4):731-9. doi: 10.1093/aob/mct126.

Shuttleworth, Craig M., Peter W.W. Lurz, Neville Geddes, and John Browne. 2012. "Integrating red squirrel (*Sciurus vulgaris*) habitat requirements with the management of pathogenic tree disease in commercial forests in the UK." *Forest Ecology and Management* 279:167-175. doi: 10.1016/j.foreco.2012.05.019.

Sibold, C., R. Ulrich, M. Labuda, Å. Lundkvist, H. Martens, M. Schütt, P. Gerke, K. Leitmeyer, H. Meisel, and D.H. Krüger. 2001. "Dobrava hantavirus causes hemorrhagic fever with renal syndrome in central Europe and is carried by two different *Apodemus* mice species." *Journal of Medical Virology* 63:158-167. doi: 10.1002/1096-9071(20000201)63:2%3c158::Aid-jmv1011%3e3.0.Co;2-%23.

Simpson, Callum, Alexander Gofton, Casey L. Taylor, Jenna P. Bytheway, Laura S. Grant, Dieter F. Hochuli, and Peter B. Banks. 2024. "Rat lungworm, Cryptosporidium and other zoonotic pathogens of *Rattus rattus* and native wildlife on Sydney's Northern beaches." *Australian Zoologist* 44 (1):231-251. doi: 10.7882/az.2024.020.

Simpson, V. 2010. "*Angiostrongylus vasorum* infection in a stoat." *Vet Rec* 166 (6):182. doi: 10.1136/vr.c666.

Simpson, V.R., J. Hargreaves, H.M. Butler, N.J. Davison, and D.J. Everest. 2013. "Causes of mortality and pathological lesions observed post-mortem in red squirrels (*Sciurus vulgaris*) in Great Britain." *BMC Vet Res* 9:229. doi: 10.1186/1746-6148-9-229.

Smith-Palmer, A., G. Hawkins, L. Browning, L. Allison, M. Hanson, R. Bruce, J. McElhiney, and J. Horne. 2018. "Outbreak of *Escherichia coli* O157 Phage Type 32 linked to the consumption of venison products." *Epidemiol Infect* 146 (15):1922-1927. doi: 10.1017/S0950268818001784.

Smith, G.C., D.G. Garthwaite, and A.J. Prickett. 2006. "Rabbit control in Great Britain." In *Advances in vertebrate pest management (vol. IV)*, edited by D.P. Cowan and C.J. Feare, 165-186. Fürth: Filander Verlag.

Smith, G.C., and D. Wilkinson. 2003. "Modeling control of rabies outbreaks in red fox populations to evaluate culling, vaccination, and vaccination combined with fertility control." *J Wildl Dis* 39 (2):278-86. doi: 10.7589/0090-3558-39.2.278.

Smith, H.V., J.S. Gray, and G. McKenzie. 1991. "A *Lyme borreliosis* human serosurvey of asymptomatic adults in Ireland." *Zentralbl Bakteriol* 275 (3):382-9. doi: 10.1016/s0934-8840(11)80304-x.

Smith, N.C., C. Goulart, J.A. Hayward, A. Kupz, C.M. Miller, and G.G. van Dooren. 2021. "Control of human toxoplasmosis." *Int J Parasitol* 51 (2-3):95-121. doi: 10.1016/j.ijpara.2020.11.001.

Smith, S.L., H.V. Senn, S. Perez-Espona, M.T. Wyman, E. Heap, and J.M. Pemberton. 2018. "Introgression of exotic *Cervus* (*nippon* and *canadensis*) into red deer (*Cervus elaphus*) populations in Scotland and the English Lake District." *Ecol Evol* 8 (4):2122-2134. doi: 10.1002/ece3.3767.

SNH. 2008. "Scottish Natural Heritage Hebridean Mink Project." www.snh.org.uk/pdfs/publications/wildlife/hebridean.pdf. Date Accessed: 18th November 2009.
Sotherton, N., S. Tapper, and A. Smith. 2009. "Hen harriers and red grouse: economic aspects of red grouse shooting and the implications for moorland conservation." *Journal of Applied Ecology* 46:955-960.
Southwell, R. M., K. Sherlock, and M. Baylis. 2020. "Cross-sectional study of British wild deer for evidence of Schmallenberg virus infection." *Vet Rec* 187 (8):e64. doi: 10.1136/vr.105869.
Stanbury, A.J., F. Burns, N.J. Aebischer, H. Baker, D.E. Balmer, A. Brown, T. Dunn, P. Lindley, M. Murphy, D.G. Noble, R. Owens, and L. Quinn. 2024. "The status of the UK's breeding seabirds: an addendum to the fifth Birds of Conservation Concern in the United Kingdom, Channel Islands and Isle of Man and second IUCN Red List assessment of extinction risk for Great Britain." *British Birds* 117:471-487.
Stapp, Paul. 2002. "Stable isotopes reveal evidence of predation by ship rats on seabirds on the Shiant Islands, Scotland." *Journal of Applied Ecology* 39 (5):831-840. doi: 10.1046/j.1365-2664.2002.00754.x.
Stejskal, V., P. Rodl, and R. Aulicky. 2016. "Pestilential activities of rodents at farms and in stores of agro-food commodities.". www.international-pest-control.
Stevenson-Holt, C.D., K. Watts, C.C. Bellamy, O.T. Nevin, and A.D. Ramsey. 2014. "Defining landscape resistance values in least-cost connectivity models for the invasive grey squirrel: a comparison of approaches using expert-opinion and habitat suitability modelling." *PLoS One* 9 (11):e112119. doi: 10.1371/journal.pone.0112119.
Stone, E.L., G. Jones, and S. Harris. 2013. "Mitigating the effect of development on bats in England with derogation licensing." *Conservation Biology* 27:1324-1334.
Strachan, C., and D.J. Jefferies. 1996. "An assessment of the diet of feral American mink, *Mustela vison*, from scats collected where water voles, *Arvicola terrestris*, occur." *Naturalist* 121:73-81.
Swift, B.M.C., E.S. Barron, R. Christley, D. Corbetta, L. Grau-Roma, C. Jewell, C. O'Cathail, A. Mitchell, J. Phoenix, A. Prosser, C. Rees, M. Sorley, R. Verin, and M. Bennett. 2021. "Tuberculosis in badgers where the bovine tuberculosis epidemic is expanding in cattle in England." *Scientific Reports* 11:20995. doi: https://doi.org/10.1038/s41598-021-00473-6.
Symes, R., and D. Yalden. 2002. "The former status of the black rat (*Rattus rattus*) in England and Wales." *Mammal Review* 32:315-316.
Tabor, R. 1981. "General biology of feral cats." In *In: The ecology and control of feral cats*, 5-11. Potters Bar: UFAW.
Takeuchi-Storm, N., H. Mejer, M.N. Al-Sabi, C.S. Olsen, S.M. Thamsborg, and H.L. Enemark. 2015. "Gastrointestinal parasites of cats in Denmark assessed by necropsy and concentration McMaster technique." *Vet Parasitol* 214 (3-4):327-32. doi: 10.1016/j.vetpar.2015.06.033.
Tappin, S. 2014. "*Angiostrongylus vasorum* update." Vet Times. https://www.vettimes.co.uk.
Tarlinton, R., J. Daly, S. Dunham, and J. Kydd. 2012. "The challenge of Schmallenberg virus emergence in Europe." *Vet J* 194 (1):10-8. doi: 10.1016/j.tvjl.2012.08.017.
Taylor, C.S., R. Garcia Gato, J. Learmount, N.A. Aziz, C. Montgomery, H. Rose, C.L. Coulthwaite, J.W. McGarry, D.W. Forman, S. Allen, R. Wall, and E.R. Morgan. 2015. "Increased prevalence and geographic spread of the cardiopulmonary nematode *Angiostrongylus vasorum* in fox populations in Great Britain." *Parasitology* 142 (9):1190-5. doi: 10.1017/S0031182015000463.
Thirgood, S.J. 1995. "The effects of sex, season and habitat availability on patterns of habitat use by fallow deer." *Journal of Zoology* 235:645-659.
Thomason, A.G., M. Begon, J.E. Bradley, S. Paterson, and J.A. Jackson. 2017. "Endemic Hantavirus in Field Voles, Northern England." *Emerg Infect Dis* 23 (6):1033-1035. doi: 10.3201/eid2306.161607.
Thompson, H.V. 1962. "Wild mink in Britain." *New Scientist* 13:130-132.
Thompson, H.V. 1968. "British wild mink." *Annals of Applied Biology* 61:345-349.
Tolhurst, Bryony A., Richard J. Delahay, Neil J. Walker, Alastair I. Ward, and Timothy J. Roper. 2009. "Behaviour of badgers (*Meles meles*) in farm buildings: Opportunities for the transmission of *Mycobacterium bovis* to cattle?" *Applied Animal Behaviour Science* 117 (1-2):103-113. doi: 10.1016/j.applanim.2008.10.009.

Trewby, I.D., R. Young, R.A. McDonald, G.J. Wilson, J. Davison, N. Walker, A. Robertson, C.P. Doncaster, and R.J. Delahay. 2014. "Impacts of removing badgers on localised counts of hedgehogs." *PLoS One* 9 (4):e95477. doi: 10.1371/journal.pone.0095477.

Trout, R.C., and A.M. Tittensor. 2008. "Can predators regulate wild Rabbit *Oryctolagus cuniculus* population density in England and Wales?" *Mammal Review* 19 (4):153-173. doi: 10.1111/j.1365-2907.1989.tb00409.x.

Trout, Roger C., Sarah Brooks, and Pat Morris. 2015. "Nest box usage by old edible dormice (*Glis glis*) in breeding and non-breeding years." *Folia Zoologica* 64 (4):320-324. doi: 10.25225/fozo.v64.i4.a5.2015.

Tsoleridis, T., O. Onianwa, E. Horncastle, E. Dayman, M. Zhu, T. Danjittrong, M. Wachtl, J.M. Behnke, S. Chapman, V. Strong, P. Dobbs, J. K. Ball, R.E. Tarlinton, and C.P. McClure. 2016. "Discovery of Novel Alphacoronaviruses in European Rodents and Shrews." *Viruses* 8 (3):84. doi: 10.3390/v8030084.

Tubby, I. 2022. "Reducing the impact of deer on the natural environment – consultation opens." Forestry Commission blog. https://forestrycommission.blog.gov.uk/2022/08/04/reducing-the-impact-of-deer-on-the-natural-environment-consultation-opens/.

Twining, J.P., C. Lawton, A. White, E. Sheehy, K. Hobson, W.I. Montgomery, and X. Lambin. 2022. "Restoring vertebrate predator populations can provide landscape-scale biological control of established invasive vertebrates: Insights from pine marten recovery in Europe." *Glob Chang Biol* 28 (18):5368-5384. doi: 10.1111/gcb.16236.

van Soolingen, D., T. Hoofenboezem, P.E. de Haas, P.W.M. Hermans, M.A. Koedam, K.S. Teppema, P.J. Brennan, G.S. Besra, F. Portaels, J. Top, L.M. Schouls, and J.D.A. Van Embden. 1997. "A novel pathogenic taxon of the *Mycobacterium tuberculosis* complex, Canetti: characterization of an exceptional isolate from Africa." *International Journal of Systematic and Evolutionary Microbacteriology* 47:1236–1245.

Vapalahti, O., J. Mustonen, Å. Lundkvist, H. Henttonen, A. Plyusnin, and A. Vaheri. 2003. "Hantavirus Infections in Europe." *Lancet Infectious Diseases* 3:653–61. doi: https://doi.org/10.1016/s1473-3099(03)00774-6.

Vaughan, N., E.A. Lucas, S. Harris, and P.C.L. White. 2003. "Habitat associations of European hares *Lepus europaeus* in England and Wales: implications for farmland management." *Journal of Applied Ecology* 40:163-175.

Velarde, R., P. Cavadini, A. Neimanis, O. Cabezon, M. Chiari, A. Gaffuri, S. Lavin, G. Grilli, D. Gavier-Widen, A. Lavazza, and L. Capucci. 2017. "Spillover Events of Infection of Brown Hares (*Lepus europaeus*) with Rabbit Haemorrhagic Disease Type 2 Virus (RHDV2) Caused Sporadic Cases of an European Brown Hare Syndrome-Like Disease in Italy and Spain." *Transbound Emerg Dis* 64 (6):1750-1761. doi: 10.1111/tbed.12562.

Veronesi, F., G. Deak, and A. Diakou. 2023. "Wild Mesocarnivores as Reservoirs of Endoparasites Causing Important Zoonoses and Emerging Bridging Infections across Europe." *Pathogens* 12 (2). doi: 10.3390/pathogens12020178.

Wang, W., X.D. Lin, W.P. Guo, R.H. Zhou, M.R. Wang, C.Q. Wang, S. Ge, S.H. Mei, M.H. Li, M. Shi, E.C. Holmes, and Y.Z. Zhang. 2015. "Discovery, diversity and evolution of novel coronaviruses sampled from rodents in China." *Virology* 474:19-27. doi: 10.1016/j.virol.2014.10.017.

Ward, A.I., G.C. Smith, T.R. Etherington, and R.J. Delahay. 2009. "Estimating the risk of cattle exposure to tuberculosis posed by wild deer relative to badgers in England and Wales." *Journal of Wildlife Diseases* 45:1104–1120.

Ward, Alastair I. 2005. "Expanding ranges of wild and feral deer in Great Britain." *Mammal Review* 35 (2):165-173. doi: 10.1111/j.1365-2907.2005.00060.x.

Ward, Alastair I., and Graham C. Smith. 2011. "Predicting the status of wild deer as hosts of *Mycobacterium bovis* infection in Britain." *European Journal of Wildlife Research* 58 (1):127-135. doi: 10.1007/s10344-011-0553-7.

Ward, Alastair I., Piran C. L. White, Andrew Smith, and Charles H. Critchley. 2004. "Modelling the cost of roe deer browsing damage to forestry." *Forest Ecology and Management* 191 (1-3):301-310. doi: 10.1016/j.foreco.2003.12.018.

Ward, L.K., R.T. Clarke, and A.S. Cooke. 1994. Long term scrub succession deflected by fallow deer at Castor

Hanglands National Nature Reserve. Annual Report of the Institute of Terrestrial Ecology (1993–4). NERC.

Warren, Philip K., and David Baines. 2002. "Dispersal, survival and causes of mortality in black grouse *Tetrao tetrix* in northern England." *Wildlife Biology* 8 (2):91-97. doi: 10.2981/wlb.2002.013.

Webster, J.P., and D.W. Macdonald. 1995. "Cryptosporidiosis reservoir in wild brown rats (*Rattus norvegicus*) in the UK." *Epidemiol Infect* 115 (1):207-9. doi: 10.1017/s0950268800058271.

Welch, D., B.W. Staines, D. Scott, D.D. French, and D.C. Catt. 1991. "Leader Browsing by Red and Roe Deer on Young Sitka Spruce Trees in Western Scotland I. Damage Rates and the Influence of Habitat Factors." *Forestry* 64 (1):61-82. doi: 10.1093/forestry/64.1.61.

Wells, B., H. Shaw, E. Hotchkiss, J. Gilray, R. Ayton, J. Green, F. Katzer, A. Wells, and E. Innes. 2015. "Prevalence, species identification and genotyping Cryptosporidium from livestock and deer in a catchment in the Cairngorms with a history of a contaminated public water supply." *Parasit Vectors* 8:66. doi: 10.1186/s13071-015-0684-x.

Wernike, K., C. Wylezich, D. Hoper, J. Schneider, P.W.W. Lurz, A. Meredith, E. Milne, M. Beer, and R.G. Ulrich. 2018. "Widespread occurrence of squirrel adenovirus 1 in red and grey squirrels in Scotland detected by a novel real-time PCR assay." *Virus Res* 257:113-118. doi: 10.1016/j.virusres.2018.09.007.

Westerman, P.R., J S. Wes, M J. Kropff, and W. Van Der Werf. 2003. "Annual losses of weed seeds due to predation in organic cereal fields." *Journal of Applied Ecology* 40 (5):824-836.

Whitby, H.L., G.S. Parry, M.Y. Jones, A. Powell, and D.W. Forman. 2010. Fisheries and predators in Wales: A preliminary consultation. Swansea University.

White, P.C.L., J.C.R. Smart, M. Böhm, J. Langbein, and A.I. Ward. 2004. "Economic impacts of wild deer in the east of England." http://www.woodlandforlife.net/wfl-woodbank/DisplayArticle_2333.html.

White, P.C.L., A.I. Ward, J.C.R. Smart, and N.P. Moore. 2004. Impacts of deer and deer management on woodland biodiversity in the English lowlands Report for the Woodland Trust.

Whitehead, G.K. 1993. *The Whitehead encyclopaedia of deer*. Shrewsbury, UK: Swan Hill Press.

Wildey, K. 2002. "UKs grain storage picture - production, storage and pests." In *Proceedings of the 2nd Meeting of the Working Group 4, Biocontrol of Arthropod Pests in Stored Products. Prague 2002 30-31st May*, edited by E. Zdarkova, M. Wakefield, J. Lukas and J. Hubert, 23-26. Prague: Research Institute of Crop Production.

Wilkinson, D., and G.C. Smith. 2008. "A preliminary survey for changes in urban Fox (*Vulpes vulpes*) densities in England and Wales, and implications for rabies control." *Mammal Review* 31 (1):107-110. doi: 10.1046/j.1365-2907.2001.00076.x.

Williams, F., R. Eschen, A. Harris, D. Djeddour, C. Pratt, R. Shaw, S. Varia, J. Lamontagne-Godwin, S. Thomas, and S. Murphy. 2010. The economic cost of invasive non-native species on Great Britain. CABI report.

Wilson, Andy M., Malcolm Ausden, and Tim P. Milsom. 2004. "Changes in breeding wader populations on lowland wet grasslands in England and Wales: causes and potential solutions." *Ibis* 146 (s2):32-40. doi: 10.1111/j.1474-919X.2004.00371.x.

Wilson, C.J. 2005. Feral wild boar in England: status, impact and management. A report on the behalf of Defra European Wildlife Division.

Wilson, C.J. 2014. "The establishment and distribution of feral wild boar (*Sus scrofa* L.) in England." *Wildlife Biology in Practice* 10:1-6.

Wilson, Charles John, Alan Britton, and Roger Symes. 2009. "An Assessment of Agricultural Damage Caused by Red Deer (*Cervus elaphus* L.)and Fallow Deer (*Dama dama* L.) in Southwest England." *Wildlife Biology in Practice* 5 (2). doi: 10.2461/wbp.2009.5.11.

Woods, Michael, Robbie A. McDonald, and Stephen Harris. 2003. "Predation of wildlife by domestic cats *Felis catus* in Great Britain." *Mammal Review* 33 (2):174-188. doi: 10.1046/j.1365-2907.2003.00017.x.

Yalden, D.W. 1998. "The past, present and future of deer in Britain." In *Population Ecology, Management and Welfare of Deer*, edited by C.R. Goldspink, S. King and R.J. Putman, 1-5. Manchester: The Manchester Metropolitan University.

Yamaguchi, N., D.W. Macdonald, W.C. Passanisi, D.A. Harbour, and C.D. Hopper. 1996. "Parasite prevalence in free-ranging farm cats, *Felis silvestris catus*." *Epidemiol Infect* 116 (2):217-23. doi: 10.1017/s0950268800052468.

Yang, X.L., B. Hu, B. Wang, M.N. Wang, Q. Zhang, W. Zhang, L.J. Wu, X.Y. Ge, Y.Z. Zhang, P. Daszak, L.F. Wang, and Z.L. Shi. 2015. "Isolation and Characterization of a Novel Bat Coronavirus Closely Related to the Direct Progenitor of Severe Acute Respiratory Syndrome Coronavirus." *J Virol* 90 (6):3253-6. doi: 10.1128/JVI.02582-15.

Yarto-Jaramillo, E. 2015. "Rodentia." In *Fowler's Zoo and Wild Animal Medicine, vol 8*, edited by R.E. Miller and M.E Fowler, 384–422. St Louis, Missouri, USA: Elsevier Saunders.

Young, R.P., J. Davison, I.D. Trewby, G.J. Wilson, R.J. Delahay, and C.P. Doncaster. 2006. "Abundance of hedgehogs (*Erinaceus europaeus*) in relation to the density and distribution of badgers (*Meles meles)*." *Journal of Zoology* 269 (3):349-356. doi: 10.1111/j.1469-7998.2006.00078.x.

Zanet, S., F. Occhibove, D. Capizzi, S. Fratini, F. Giannini, A. D. Hoida, P. Sposimo, F. Valentini, and E. Ferroglio. 2023. "Zoonotic Microparasites in Invasive Black Rats (*Rattus rattus*) from Small Islands in Central Italy." *Animals (Basel)* 13 (20). doi: 10.3390/ani13203279.

Zeale, M.R., E. Bennitt, S.E. Newson, C. Packman, W.J. Browne, S. Harris, G. Jones, and E. Stone. 2016. "Mitigating the Impact of Bats in Historic Churches: The Response of Natterer's Bats *Myotis nattereri* to Artificial Roosts and Deterrence." *PLoS One* 11 (1):e0146782. doi: 10.1371/journal.pone.0146782.

Zielonka, Natalia B., Robert W. Hawkes, Helen Jones, Robert J. Burnside, and Paul M. Dolman. 2020. "Placement, survival and predator identity of Eurasian Curlew *Numenius arquata* nests on lowland grass-heath." *Bird Study* 66 (4):471-483. doi: 10.1080/00063657.2020.1725421.

Zini, Valentina, Kristin Wäber, and Paul M. Dolman. 2022. "Relation of pine crop damage to species-specific density in a multi-ungulate assemblage." *European Journal of Forest Research* 141 (3):489-502. doi: 10.1007/s10342-022-01452-w.

Zonfrillo, B. 2000. Ailsa Craig: Before and after the eradication of rats in 1991. Ayrshire Bird Report 2000.

Zub, Karol, Pablo García-Díaz, Sarah Sankey, Robert Eisler, and Xavier Lambin. 2022. "Using a Modeling Approach to Inform Progress Towards Stoat Eradication From the Orkney Islands." *Frontiers in Conservation Science* 2. doi: 10.3389/fcosc.2021.780102.
