## Supplementary Material S3 for "Identifying and quantifying conflicts between humans and terrestrial mammals in Great Britain"

**Supplementary Table S3 to "Identifying and quantifying conflicts between humans and terrestrial mammals in Great Britain"**

List of species causing conflict, including order, status, population size, body mass, biomass and type of conflicts recorded over time (pre-2014 only; both pre-2014 and 2014 to present; 2014 to present only). The median impact score (1=minimal known impact to 5=highest known impact) and confidence rating (1=low; 2=medium; 3=high) are shown after each conflict [x,x]. The highest impact scores are shown in **bold**.

| Order | Species (status) | Population size[1] | Body mass (kg)[2] | Biomass (kg)[3] | No. of conflicts | Pre-2014 only conflicts | Both pre-2014 and 2014 to present conflicts | 2014 to present only conflicts |
|---|---|---|---|---|---|---|---|---|
| Artiodactyla | Roe deer (native) | 264,300 | 20 | 5,286,000 | 9 | - | Damage to forestry [4,3]<br>Damage to agricultural interests [4,3]<br>Damage to vehicles (RTAs) [4,3]<br>Health risks from RTAs [3,2]<br>Reservoir of disease [2,3]<br>bTB transmission [1,2]<br>Reservoir of zoonotic disease [2,2]<br>Damage to conservation habitats [3,2] | Threat to biodiversity [2,2] |
| | Red deer (native) | 345,900 | 75 | 25,942,500 | 10 | Competition with livestock [1,2] | Damage to forestry [4,3]<br>Damage to agricultural interests [3,3]<br>Damage to vehicles (RTAs) [3,3]<br>Health risks from RTAs [2,2]<br>Reservoir of disease [2,3]<br>bTB transmission [1,2]<br>Reservoir of zoonotic disease [2,2]<br>Damage to conservation habitats [3,2] | Threat to biodiversity [2,2] |
| | Sika deer (non-native) | 102,900 | 50 | 5,145,000 | 11 | Competition with livestock [1,3] | Damage to forestry [3,2]<br>Damage to agricultural interests [2,2]<br>Damage to vehicles (RTAs) [2,2]<br>Health risks from RTAs [2,2]<br>bTB transmission [1,1]<br>Reservoir of zoonotic disease [1,3]<br>Damage to conservation habitats [2,2]<br>Threat to genetic integrity [2,2] | Reservoir of disease [1,1]<br>Threat to biodiversity [2,2] |

[1] Population sizes derived from Mathews et al. (2018), except weasel (www.ptes.org), Exmoor/Dartmoor pony (www.visitdartmoor.co.uk & www.visitexmoor.co.uk), beaver (www.wildlifetrusts.org), Skomer vole (www.animalia.bio/skomer-vole), black rat (www.britishwildlifecentre.co.uk)
[2] Typical body weights taken from Harris & Yalden (2008), except beaver, Orkney vole and water vole (all www.mammal.org.uk), Exmoor/Dartmoor pony (www.rewildingbritain.org), Skomer vole (www.animalia.bio/skomer-vole)
[3] Biomass (kg) = population size X body mass (kg)

| | | | | | | | | |
|---|---|---|---|---|---|---|---|---|
| | Fallow deer (non-native, naturalised) | 263,700 | 50 | 13,185,000 | 9 | - | Damage to forestry [4,3]<br>Damage to agricultural interests [4,3]<br>Damage to vehicles (RTAs) [4,3]<br>Health risks from RTAs [2,2]<br>Reservoir of disease [2,3]<br>bTB transmission [2,2]<br>Reservoir of zoonotic disease [2,2]<br>Damage to conservation habitats [2,2] | Threat to biodiversity [2,2] |
| | Chinese water deer (non-native) | 3,600 | 15 | 54,000 | 7 | Health risks from RTAs [1,2]<br>Reservoir of disease [1,2]<br>Damage to conservation habitats [1,2] | Damage to forestry [1,3]<br>Damage to agricultural interests [2,3]<br>Damage to vehicles (RTAs) [2,3] | Reservoir of zoonotic disease [1,2] |
| | Muntjac (non-native) | 128,300 | 13 | 1,667,900 | 9 | - | Damage to forestry [2,2]<br>Damage to agricultural interests [2,2]<br>Damage to vehicles (RTAs) [3,3]<br>Health risks from RTAs [2,2]<br>Reservoir of disease [2,3]<br>bTB transmission [2,1]<br>Damage to conservation habitats [2,3] | Reservoir of zoonotic disease [2,2]<br>Public nuisance [2,2] |
| | Feral goat (non-native, naturalised) | 7,500 | 45 | 337,500 | 6 | Damage to forestry [2,2]<br>Competition with livestock [1,2]<br>Reservoir of disease [1,2]<br>Reservoir of zoonotic disease [1,3]<br>Damage to conservation habitats [2,2] | Public nuisance [2,3] | - |
| | Wild boar (native) | 2,650 | 80 | 212,000 | 7 | - | Damage to agricultural interests [2,2]<br>Damage to vehicles (RTAs) [2,3]<br>Health risks from RTAs [3,3]<br>Reservoir of disease [2,2]<br>bTB transmission [2,2]<br>Damage to conservation habitats [1,2]<br>Public nuisance [1,2] | - |

[1] Population sizes derived from Mathews et al. (2018), except weasel (www.ptes.org), Exmoor/Dartmoor pony (www.visitdartmoor.co.uk & www.visitexmoor.co.uk), beaver (www.wildlifetrusts.org), Skomer vole (www.animalia.bio/skomer-vole), black rat (www.britishwildlifecentre.co.uk)
[2] Typical body weights taken from Harris & Yalden (2008), except beaver, Orkney vole and water vole (all www.mammal.org.uk), Exmoor/Dartmoor pony (www.rewildingbritain.org), Skomer vole (www.animalia.bio/skomer-vole)
[3] Biomass (kg) = population size X body mass (kg)

| | | | | | | | | |
|---|---|---|---|---|---|---|---|
| Carnivora | Feral cat (feral) | Unknown | 4.4 | - | 3 | - | Reservoir of disease [2,2]<br>Threat to genetic integrity [3,2]<br>Competition with or predation of wildlife (inc. protected or conservation spp.) [2,2] | - |
| | Wildcat (native) | 200 | 4.5 | 900 | 1 | Predation of game species [1,3] | - | - |
| | Otter (native) | 11,000 | 8 | 88,000 | 2 | - | Predation/impact on commercial fisheries [2,2]<br>Competition with or predation of wildlife (inc. protected or conservation spp.) [2,2] | - |
| | Pine marten (native) | 3,739 | 1.5 | 5,609 | 4 | Predation of game species [1,2]<br>Predation of poultry [1,2]<br>Domestic nuisance [2,2] | - | Competition with or predation of wildlife (inc. protected or conservation spp.) [2,3] |
| | Stoat (native) | 437,600 | 0.25 | 109,400 | 3 | Reservoir of disease [1,2] | Predation of game species [2,2] | Competition with or predation of wildlife (inc. protected or conservation spp.) [2,2] |
| | Feral ferret (non-native) | Uknown | 0.9 | - | 3 | Predation of game species [1,1]<br>Predation of poultry [1,1] | Threat to genetic integrity [2,1] | - |
| | Weasel (native) | 450,000 | 0.9 | 405,000 | 1 | - | Predation of game species [1,2] | - |
| | Polecat (native) | 83,200 | 0.9 | 74,880 | 3 | Predation of game species [1,2]<br>Predation of poultry [1,2] | Reservoir of zoonotic disease [1,2] | - |
| | Badger (native) | 661,000 | 10.5 | 6,940,500 | 10 | Predation of game species [2,2] | **Damage to agricultural interests [5,3]**<br>Damage to vehicles (RTA) [4,2]<br>Damage to property [4,3]<br>Conflicts with construction, infrastructure and development [4,2]<br>Health risks from RTAs [2,2]<br>**bTB transmission [5,3]**<br>Competition with or predation of wildlife (inc. protected or conservation spp.) [3,3] | Damage to amenities and domestic grassland [2,2]<br>Reservoir of disease [2,3] |

[1] Population sizes derived from Mathews et al. (2018), except weasel (www.ptes.org), Exmoor/Dartmoor pony (www.visitdartmoor.co.uk & www.visitexmoor.co.uk), beaver (www.wildlifetrusts.org), Skomer vole (www.animalia.bio/skomer-vole), black rat (www.britishwildlifecentre.co.uk)
[2] Typical body weights taken from Harris & Yalden (2008), except beaver, Orkney vole and water vole (all www.mammal.org.uk), Exmoor/Dartmoor pony (www.rewildingbritain.org), Skomer vole (www.animalia.bio/skomer-vole)
[3] Biomass (kg) = population size X body mass (kg)

| | | | | | | | | |
|---|---|---|---|---|---|---|---|---|
| | American mink (non-native) | 121,900 | 0.8 | 97,520 | 6 | Predation of game species [2,1]<br>Predation/impact on commercial fisheries [2,2] | Predation of poultry [2,3]<br>Damage to amenities and domestic grassland [3,3]<br>Reservoir of zoonotic disease [2,2]<br>Competition with or predation of wildlife (inc. protected or conservation spp.) [3,3] | - |
| | Red fox (native) | 356,700 | 6 | 2,140,200 | 8 | Domestic nuisance [1,3] | Predation of game species [4,3]<br>Predation of livestock [4,3]<br>Reservoir of disease [2,3]<br>bTB transmission [1,3]<br>Reservoir of zoonotic disease [2,3]<br>Competition with or predation of wildlife (inc. protected or conservation spp.) [3,3] | Predation of domestic pets [2,3] |
| Chiroptera | Daubenton's bat (native) | 1,025,000 | 0.008 | 8,200 | 2 | - | Reservoir of disease [1,2]<br>Reservoir of zoonotic disease [2,3] | |
| | Natterer's bat (native) | 693,500 | 0.008 | 5,548 | 3 | | Conflicts with construction, infrastructure and development [3,3] | Domestic nuisance & hygiene[2,2] |
| | Serotine bat (native) | 15,050 | 0.032 | 482 | 3 | | Conflicts with construction, infrastructure and development [3,3] | Reservoir of zoonotic disease [2,3]<br>Domestic nuisance & hygiene[2,2] |
| | Common pipistrelle (native) | 2,745,297 | 0.006 | 16,472 | 1 | - | Conflicts with construction, infrastructure and development [3,3] | - |
| | Soprano pipistrelle (native) | 4,668,000 | 0.006 | 28,008 | 1 | - | Conflicts with construction, infrastructure and development [3,3] | - |
| | Brown long-eared bat (native) | 933,600 | 0.008 | 7,469 | 1 | - | Conflicts with construction, infrastructure and development [3,3] | - |
| Eulipotyphla | Hedgehog (native) | 521,800 | 0.5 | 260,900 | 4 | Predation of game species [2,2]<br>Reservoir of disease [1,2] | Reservoir of zoonotic disease [2,2]<br>Predation of important ground-nesting birds & island ecosystem damage [3,3] | - |
| | European mole (native) | 41,400,000 | 0.1 | 4,140,000 | 3 | Threat to livestock where ground is undermined [2,1] | Damage to agricultural interests [4,3]<br>Damage to amenities or domestic grassland [3,3] | - |

[1] Population sizes derived from Mathews et al. (2018), except weasel (www.ptes.org), Exmoor/Dartmoor pony (www.visitdartmoor.co.uk & www.visitexmoor.co.uk), beaver (www.wildlifetrusts.org), Skomer vole (www.animalia.bio/skomer-vole), black rat (www.britishwildlifecentre.co.uk)
[2] Typical body weights taken from Harris & Yalden (2008), except beaver, Orkney vole and water vole (all www.mammal.org.uk), Exmoor/Dartmoor pony (www.rewildingbritain.org), Skomer vole (www.animalia.bio/skomer-vole)
[3] Biomass (kg) = population size X body mass (kg)

| | | | | | | | | |
|---|---|---|---|---|---|---|---|---|
| | Common shrew (native) | 21,020,000 | 0.01 | 210,200 | 1 | - | - | Reservoir of zoonotic disease [1,2] |
| | Greater white-toothed shrew (non-native) | Unknown | 0.011 | - | 2 | - | - | Reservoir of zoonotic disease [1,2]<br>Threat to biodiversity [2,2] |
| Lagomorpha | Rabbit (non-native, naturalised) | 36,010,000 | 1.5 | 54,015,000 | 8 | Sustaining predators harmful to important conservation species [2,2] | **Damage to forestry [5,3]**<br>**Damage to agricultural interests [5,3]**<br>Damage to amenities and domestic grassland [4,3]<br>Conflicts with transport [3,3]<br>Reservoir of disease [3,3]<br>Reservoir of zoonotic disease [2,2] | Public nuisance [2,2] |
| | Brown hare (non-native, naturalised) | 579,000 | 3.3 | 1,910,700 | 5 | Damage to agricultural interests [2,2] | Damage to forestry [3,2]<br>Damage to property as a result of poaching [2,1] | Reservoir of disease [1,2]<br>Reservoir of zoonotic disease [1,2] |
| | Mountain hare (native) | 134,500 | 2.8 | 376,600 | 4 | Damage to forestry [2,2]<br>Damage to agricultural interests [2,2]<br>Damage to property as a result of poaching [1,2]<br>Reservoir of zoonotic disease [1,2] | - | - |
| Perissodactyla | Dartmoor/ Exmoor pony (feral) | 1,850 | 363 | 671,550 | 2 | - | Damage to vehicles (RTAs) [2,2]<br>Health risks from RTAs [1,1] | - |
| Rodentia | Red squirrel (native) | 287,090 | 0.2 | 57,418 | 2 | - | Reservoir of zoonotic disease [1,2] | Reservoir of disease [1,2] |
| | Grey squirrel (non-native) | 2,701,000 | 0.5 | 1,350,500 | 7 | Damage to agricultural interests [2,2] | **Damage to forestry [5,3]**<br>Damage to property [4,3]<br>Reservoir of disease [3,3]<br>Threat of branches falling to humans [1,2]Threat to biodiversity [4,3] | Reservoir of zoonotic disease [1,2] |

[1] Population sizes derived from Mathews et al. (2018), except weasel (www.ptes.org), Exmoor/Dartmoor pony (www.visitdartmoor.co.uk & www.visitexmoor.co.uk), beaver (www.wildlifetrusts.org), Skomer vole (www.animalia.bio/skomer-vole), black rat (www.britishwildlifecentre.co.uk)
[2] Typical body weights taken from Harris & Yalden (2008), except beaver, Orkney vole and water vole (all www.mammal.org.uk), Exmoor/Dartmoor pony (www.rewildingbritain.org), Skomer vole (www.animalia.bio/skomer-vole)
[3] Biomass (kg) = population size X body mass (kg)

| | | | | | | | | |
|---|---|---|---|---|---|---|---|---|
| | Eurasian beaver (native) | 2,000 | 25.25 | 50,500 | 6 | - | - | Damage to forestry [2,3]<br>Damage to agricultural interests [2,2]<br>Predation/impact on commercial fisheries [1,2]<br>Conflicts with construction, infrastructure and development [2,2]<br>Reservoir of disease [1,3]<br>Reservoir of zoonotic disease [1,3] |
| | Hazel dormouse (native) | 929,000 | 0.018 | 16,722 | 1 | - | Conflicts with construction, infrastructure and development [2,2] | - |
| | Edible dormouse (non-native) | 23,000 | 0.14 | 3,220 | 4 | Damage to agricultural interests [1,2] | Damage to forestry [3,3]<br>Damage to property [3,3]<br>Reservoir of zoonotic disease [1,2] | - |
| | Bank vole (native) | 27,420,000 | 0.02 | 548,400 | 3 | Damage to conservation habitats [1,3] | Damage to forestry [2,2] | Reservoir of zoonotic disease [2,2] |
| | Field vole (native) | 56,530,000 | 0.02 | 1,130,600 | 5 | Damage to agricultural interests [1,1] | Damage to forestry [3,2] | Sustaining predators harmful to game species [1,2]<br>Reservoir of disease [2,2]<br>Reservoir of zoonotic disease [2,2] |
| | Wood mouse (native) | 39,600,000 | 0.02 | 792,000 | 3 | Damage to agricultural interests [2,2] | Reservoir of disease [2,2] | Reservoir of zoonotic disease [2,2] |
| | Yellow-necked mouse (native) | 1,500,000 | 0.03 | 45,00 | 1 | - | - | Reservoir of zoonotic disease [1,1] |
| | Orkney vole (non-native, naturalised) | 1,500,000 | 0.045 | 67,500 | 1 | - | - | Sustaining predators harmful to game species [1,2] |
| | Skomer vole (non-native, naturalised) | 20,000 | 0.04 | 800 | 1 | - | - | Reservoir of zoonotic disease [1,2] |
| | Water vole (native) | 131,700 | 0.3 | 39,510 | 1 | - | - | Conflicts with construction, infrastructure and development [2,2] |

[1] Population sizes derived from Mathews et al. (2018), except weasel (www.ptes.org), Exmoor/Dartmoor pony (www.visitdartmoor.co.uk & www.visitexmoor.co.uk), beaver (www.wildlifetrusts.org), Skomer vole (www.animalia.bio/skomer-vole), black rat (www.britishwildlifecentre.co.uk)
[2] Typical body weights taken from Harris & Yalden (2008), except beaver, Orkney vole and water vole (all www.mammal.org.uk), Exmoor/Dartmoor pony (www.rewildingbritain.org), Skomer vole (www.animalia.bio/skomer-vole)
[3] Biomass (kg) = population size X body mass (kg)

| | | | | | | | | |
|---|---|---|---|---|---|---|---|---|
| | House mouse (non-native, naturalised) | 5,202,900 | 0.015 | 78,044 | 4 | - | Damage to agricultural interests [4,2]<br>Damage to property [4,3]<br>Reservoir of disease [2,3]<br>Reservoir of zoonotic disease [3,3] | |
| | Brown rat (non-native) | 7,070,000 | 0.25 | 1,767,500 | 7 | Predation of game species [2,2] | **Damage to agricultural interests [5,3]**<br>**Damage to property [5,3]**<br>Conflicts with transport [3,3]<br>Conflicts with utilities [4,3]<br>Reservoir of zoonotic disease [4,3]<br>Predation of important ground-nesting birds & island ecosystem damage[3,2] | - |
| | Black rat (non-native) | 1,300 | 0.175 | 228 | 2 | Reservoir of zoonotic disease [1,3]<br>Damage to island ecosystems [1,3] | | |

[1] Population sizes derived from Mathews et al. (2018), except weasel (www.ptes.org), Exmoor/Dartmoor pony (www.visitdartmoor.co.uk & www.visitexmoor.co.uk), beaver (www.wildlifetrusts.org), Skomer vole (www.animalia.bio/skomer-vole), black rat (www.britishwildlifecentre.co.uk)
[2] Typical body weights taken from Harris & Yalden (2008), except beaver, Orkney vole and water vole (all www.mammal.org.uk), Exmoor/Dartmoor pony (www.rewildingbritain.org), Skomer vole (www.animalia.bio/skomer-vole)
[3] Biomass (kg) = population size X body mass (kg)
