## Supplementary Material S4 for "Identifying and quantifying conflicts between humans and terrestrial mammals in Great Britain"

**Supplementary Table S4 to "Identifying and quantifying conflicts between humans and terrestrial mammals in Great Britain"**

Summary of model formulae applied when analysing the effect of non-native status and several independent covariates (population size (*circa* 2018), body mass (kg), and biomass (kg) on the impact scores (maximum or potential) of mammal species in Great Britain. Analyses were performed using generalised linear models (GLM) fitted using the *glmmTMB::glmmTMB* function (version 1.1.10; Brooks et al., 2017). Models were fitted with a gaussian error distribution, with "identity" link function, with impact scores log-transformed (log (n + 1)) to conform with model assumptions. Covariates were also log-transformed.

| Covariate | Model formulae |
|---|---|
| Population size (2018) | $\log(Score_i + 1) = \beta_0 + \beta_1 \cdot Nonnative\ status_i + \beta_2 \cdot \log(Population\ size_i) + \beta_3 \cdot (Nonnative\ status_i \text{ X } Population\ size_i)) + \varepsilon_i$ |
| Body mass (kg) | $\log(Score_i + 1) = \beta_0 + \beta_1 \cdot Nonnative\ status_i + \beta_2 \cdot \log(Body\ mass) + \beta_3 \cdot (Nonnative\ status_i \text{ X } Body\ mass)) + \varepsilon_i$ |
| Biomass (kg) | $\log(Score_i + 1) = \beta_0 + \beta_1 \cdot Nonnative\ status_i + \beta_2 \cdot \log(Biomass) + \beta_3 \cdot (Nonnative\ status_i \text{ X } Biomass)) + \varepsilon_i$ |

Note: *i* denotes the species-of-interest, and is applied when describing impact score (maximum or potential; $Score_i$), native/non-native status ($Nonnative\ status_i$) and population size ($Population\ size_i$), body mass ($Body\ mass_i$) and biomass ($Biomass_i$). $\beta_0$ denotes the model intercept, $\beta_1$ the effect of native/non-native status, $\beta_2$ the effect of population size, body mass, or biomass, $\beta_3$ the effect of an interaction between native/non-native status and either population size, body mass, or biomass, and $\varepsilon_i$ the error term (or residual) for species *i* , following a normal (i.e. gaussian) distribution $\varepsilon_i \sim N(0,\sigma^2)$.
