## Supplementary Table S5 for "Identifying and quantifying conflicts between humans and terrestrial mammals in Great Britain"

### Supplementary Tables 5a-5d to "Identifying and quantifying conflicts between humans and terrestrial mammals in Great Britain"

5a. Total number of mammal species causing conflict within the 15 economic subcategories and their median impact score. Status categories: Na=native; NN=non-native; Fe=feral.

| | Mammal species impact scores within each subcategory | | | | | Total per status |
|---|---|---|---|---|---|---|
| **Conflict (total mammal species)** | **Impact 5** | **Impact 4** | **Impact 3** | **Impact 2** | **Impact 1** | |
| Damage to forestry (15) | Grey squirrel, rabbit | Roe deer, red deer, fallow deer | Japanese sika, brown hare, edible dormouse, field vole | Muntjac, feral goat, mountain hare, beaver, bank vole | Chinese water deer | **Na - 6**<br>**NN - 9** |
| Damage to agricultural interests (19) | Badger, rabbit, brown rat | Roe deer, fallow deer, mole, house mouse | Red deer | Japanese sika, Chinese water deer, muntjac, wild boar, brown hare, mountain hare, grey squirrel, beaver, wood mouse | Edible dormouse, field vole | **Na - 9**<br>**NN - 10** |
| Damage to vehicles (RTAs) (9) | - | Roe deer, fallow deer, badger | Red deer, muntjac | Japanese sika, Chinese water deer, wild boar, Exmoor/Dartmoor pony | - | **Na - 4**<br>**NN - 4**<br>**Fe - 1** |
| Competition with livestock (3) | - | - | - | - | Red deer, Japanese sika, feral goat | **Na - 1**<br>**NN - 2** |
| Predation of game species (11) | - | Red fox | - | Stoat, badger, American mink, hedgehog, brown rat | Wild cat, pine marten, feral ferret, weasel, polecat | **Na - 8**<br>**NN - 3** |
| Predation/impact on commercial fisheries (3) | - | - | - | Otter, American mink | Beaver | **Na - 2**<br>**NN - 1** |
| Predation of poultry (4) | - | - | - | American mink | Pine marten, feral ferret, polecat | **Na - 2**<br>**NN - 2** |
| Damage to property (6) | Brown rat | Badger, grey squirrel, house mouse | Edible dormouse | Natterer's bat | - | **Na - 2**<br>**NN - 4** |
| Damage to amenities and domestic grassland (4) | - | Rabbit | American mink, mole | Badger | - | **Na - 2**<br>**NN=2** |
| Conflicts with construction, infrastructure and development (9) | - | Badger | Natterer's, serotine, common pipistrelle, soprano pipistrelle, brown long-eared bat | Beaver, hazel dormouse, water vole | - | **Na - 9** |
| Predation of livestock (1) | - | Red fox | - | - | - | **Na - 1** |
| Conflicts with transport (2) | - | - | Rabbit, brown rat | - | - | **NN - 2** |
| Sustaining predators harmful to game (2) | - | - | - | - | Field vole, Orkney vole | **Na - 1**<br>**NN - 1** |
| Damage to property as a result of poaching (2) | | | | Brown hare | Mountain hare | **Na - 1**<br>**NN - 1** |
| Conflicts with utilities (1) | - | Brown rat | - | - | - | **NN - 1** |
| **Total no. species per impact score** | **6** | **18** | **17** | **32** | **18** | **91** |

5b. Total number of mammal species causing conflict within the six health subcategories and their median impact score. Status categories: Na=native; NN=non-native; F=feral.

| | Mammal species impact scores within each subcategory | | | | | Total per status |
|---|---|---|---|---|---|---|
| **Conflict (total mammal species)** | **Impact 5** | **Impact 4** | **Impact 3** | **Impact 2** | **Impact 1** | |
| Health risks from RTAs (9) | - | - | Roe deer, wild boar | Red deer, Japanese sika, fallow deer, muntjac, badger | Chinese water deer, Exmoor/Dartmoor pony | **Na - 4**<br>**NN - 4**<br>**Fe - 1** |
| Reservoir of disease (22) | - | - | Rabbit, grey squirrel | Roe deer, red deer, fallow deer, muntjac, wild boar, feral cat, badger, red fox, field vole, wood mouse, house mouse | Japanese sika, Chinese water deer, feral goat, stoat, Daubenton's, hedgehog, brown hare, red squirrel, beaver | **Na - 12**<br>**NN - 9**<br>**Fe - 1** |
| bTB transmission (8) | Badger | - | - | Fallow deer, muntjac, wild boar | Roe deer, red deer, Japanese sika, red fox | **Na - 5**<br>**NN - 3** |
| Reservoir of zoonotic disease (30) | - | Brown rat | House mouse | Roe deer, red deer, fallow deer, muntjac, American mink, red fox, Daubenton's, serotine, hedgehog, rabbit, bank vole, field vole, wood mouse | Japanese sika, Chinese water deer, feral goat, polecat, common shrew, greater white-toothed shrew, brown hare, mountain hare, red squirrel, grey squirrel, beaver, edible dormouse, yellow-necked mouse, Skomer vole, black rat | **Na - 15**<br>**NN - 15** |
| Threat to livestock where ground is undermined (1) | - | - | - | Mole | - | **Na - 1** |
| Threats of branches falling (1) | - | - | - | - | Grey squirrel | **NN - 1** |
| **Total no. species per impact score** | **1** | **1** | **5** | **33** | **31** | **71** |

5c. Total number of mammal species causing conflict within the seven environmental subcategories and their median impact score. Status categories: Na=native; NN=non-native; F=feral.

| | Mammal species impact scores within each subcategory | | | | | Total per status |
|---|---|---|---|---|---|---|
| **Conflict (total mammal species)** | **Impact 5** | **Impact 4** | **Impact 3** | **Impact 2** | **Impact 1** | |
| Damage to conservation habitats (9) | - | - | Roe dee, red deer | Japanese sika, fallow deer, muntjac, feral goat | Chinese water deer, wild boar, bank vole | **Na - 4**<br>**NN - 5** |
| Threat to biodiversity (6) | - | Grey squirrel | - | Roe deer, red deer, Japanese sika, fallow deer, greater white-toothed shrew | - | **Na - 2**<br>**NN - 4** |
| Threat to genetic integrity (3) | - | - | Feral cat | Japanese sika, feral ferret | - | **NN - 2**<br>**Fe - 1** |
| Competition with or predation of wildlife (including protected or conservation species) (7) | - | - | Badger, American mink, red fox | Feral cat, otter, pine marten, stoat | - | **Na - 5**<br>**NN - 1**<br>**Fe - 1** |
| Predation of important ground-nesting birds and island ecosystem damage (2) | - | - | Hedgehog, brown rat | - | - | **Na - 1**<br>**NN - 1** |
| Sustaining predators harmful to important conservation species (1) | | | | Rabbit | - | **NN - 1** |
| Damage to island ecosystems (1) | - | - | - | - | Black rat | **NN - 1** |
| **Total no. species per impact score** | **0** | **1** | **8** | **16** | **4** | **29** |

5d. Total number of mammal species causing conflict within the four social subcategories and their median impact score. Status categories: Na=native; NN=non-native; F=feral.

| | Mammal species impact scores within each subcategory | | | | | Total per status |
|---|---|---|---|---|---|---|
| **Conflict (total mammal species)** | **Impact 5** | **Impact 4** | **Impact 3** | **Impact 2** | **Impact 1** | |
| Public nuisance (4) | - | - | - | Muntjac, feral goat, rabbit | Wild boar | **Na - 1**<br>**NN - 3** |
| Domestic nuisance (2) | - | - | - | Pine marten | Red fox | **Na - 2** |
| Predation of domestic pets (1) | - | - | - | Red fox | - | **Na - 1** |
| Domestic nuisance & hygiene (2) | - | - | - | Natterer's, serotine | - | **Na - 2** |
| **Total no. species per impact score** | **0** | **0** | **0** | **7** | **2** | **9** |
